# MODEL BASED OPTIMIZACION OF THE TERPENOIDS BIOSYNTHESIS IN *E. coli*

**DOI:** 10.1101/2021.08.11.455928

**Authors:** Fernando Álvarez-Vasquez, Carlos González-Alcón, Julia Gallego-Jara, Teresa de Diego, Manuel Cánovas, Néstor V. Torres

## Abstract

Terpenoids are a family of compounds with high industrial interest and the development of biotechnological production methods is essential to achieve more sustainable alternatives to traditional extraction and synthesis methods. The modification and engineering of the catalytic activity (kcat) have been shown to be a feasible strategy in the biotechnological realm. Accordingly, we introduce a novel optimization strategy based in the modification of the kcat of the enzymes and applied it to the maximization of the terpenoids synthesis in *E. coli*. This approach is fairly general and can be applied alone or in conjunction with classic optimization strategies such as the modification of enzymatic specific activities.

For this purpose we first build up a reliable dynamic mathematical model of the alternative mevalonate pathway synthesis leading terpenoids biosynthesis in *E. coli* through the methyl-D-erythritol 4-phosphate (MEP) pathway. This model includes the 2-C-methyl-D-erythritol 2, 4-diphosphate (MEC) pumps that mediate MEC extracellular extrusion. Although the physiological significance of the MEC extrusion is still discussed and their biological function is not clear, we find that this process is a must to guarantee bacteria homeostasis and cell viability.

We have identified the enzyme IspA as a bottleneck of the terpenoids biosynthesis, which is dual substrate for the two bisubstrate final reactions. Here are presented different ways to overcome this enzymatic flux restriction by modification of the IspA kcat or by introduction of new enzymes with parallel function.

Our results show that the MEP pathway optimized solutions for kcat can yield a maximum of 17.68 fold increment in the terpenoids biosynthetic flux when the kcat are modified. This maximal solution involves the modification of 8 kcat and the corresponding Km’s and Kd’s. Remarkably, this increase doesn’t imply a change in the total enzyme concentration of the cell. This is favorable output since enzyme overproduction can compromise cell functionality.

We also apply the overexpression of the enzyme activity approach and then we have compared and combined both strategies including scenarios as the deletion or import of the genes expressing enzymes not naturally present in *E. coli*.

A combined strategy of enzymes concentrations and kcat modifications with changes up ti six enzyme levels allows a flux increment of 21.22 fold the basal value. It involves the incorporation of two IspA with similar GPP and IPP bisubstrate functions than the native one but low affinity for the DMAPP as substrate.

## 1. INTRODUCTION

Terpenoids are a wide family of natural compounds, generally produced by plants, with a great structural variety and a large number of pharmacological, food or cosmetic applications [1]. The high industrial interest of these compounds has led to the search for alternative production methods to the extraction of their natural sources and chemical synthesis. The fermentation of microorganisms has become the most promising alternative due mainly to be a more sustainable production method than traditional [2-3]. One of the most important aspects of production methods based on microbial fermentation is the selection of the host microorganism. The most commonly used microorganisms in biotechnology are *Escherichia coli* (*E. coli*) and *Saccharomyces cerevisiae* (*S. cerevisiae*), prokaryotic and eukaryotic models, respectively [4].

The main advantages of *E. coli* as host are, fundamentally, its rapid growth, the simplicity of its genome, the great variety of techniques developed for its genetic manipulation, the diversity of culture media and the great knowledge about its metabolism [5]. The last one may be the most important, since it allows the optimization of production processes in order to obtain the best yields.

The use of rational optimization approaches in biotechnological settings is important not only to avoid the wasting of time and resources but principally because it can suggest optimization profiles that can enhance end product(s) of interest. In the case of the *E. coli* terpenoids production, the use of mathematical rational optimization routines have been scarcely addressed in the literature up to now [6-7]. In this work we are focused on this key biotechnological aspect. Thus, we introduce a mathematically optimization method to this biosynthetic pathway through the modification of the enzyme catalytic constants. We also consider the overexpression of the enzyme activity approach and then we have compared and combined both strategies including scenarios as the deletion or import of genes expressing enzymes not naturally present in *E. coli* [6].

What it is known about the terpenoids bioproduction is the excessive overexpression of some enzyme related genes, namely dxs, idi, ispD, and/or ispF can inhibit isoprenoid production [8-13]. It is thus of capital importance to know how much the gene(s) associated with the enzymes of interest must be modified in order to increase terpenoids production. Another topic of relevance regarding this issue is the well-known fact that any diversion of substrates out of the route of interest decreases the rate of synthesis. In our case the reported high accumulation of the MEC in the broth [13] seem to be caused by effluxes of MEC from the MEP pathway.

Although the transporters involved in the MEC extrusion are not well known, preliminary studies showed that the *fsr* efflux pump may have a role in this process [13]. The *fsr* efflux pump is part of the major facilitator superfamily driven by proton gradient[14-16]. Probably the MEC extrusion is performed by more than one pump as they can overlap their functionality if one pump is fully or partially exerted. This overlapping of functions is the case of the pumps involved in the antibiotics extrusion as they can accept diversity of antibiotics as substrate, [15]. Gram negative bacteria require energy for the extrusion process which is supplied by proton motive force generated by adenyl phosphates hydrolysis [17], Na^+^ or Ca^2+^ exchange and other methods. Only the diffusion processes which are either in equilibrium or approaching at the equilibrium does not require energy but their contribution at the MEC traffic is low and is not considered here.

Finally, the reported G3P feed forward activation of DXS [18] was not included in the model because the G3P was settled as an independent variable and by then not affecting the dynamics of the route.

## 2. MATERIALS AND METHODS

### Model design and assumptions

Figure 1 presents the proposed mechanistic model of the 4-diphosfocytidyl-2C-methyl-D-erytritol-2-phosphate (MEP) pathway in *E. coli*. When available, data were collected from *E. coli* wild type populations growing in continuous culture bioreactors during the idiophase. A key features of the model includes the formation of 2-C-methyl-D-erythritol 2,4-diphosphate (MEC, *X*_7_) and their posterior degradation [13, 19-21]. The model also incorporates the pump(s) that mediates the MEC extrusion. It includes the end of the pathway namely the IDI (*X*_15_) and IspA (*X*_11_ and *X*_16_) enzymes that regulates the key metabolites DMAPP (*X*_5_) and IPP(*X*_4_) that exert feedback inhibiting at the first enzyme of the route DXS (*X*_24_).

**Figure 1.**
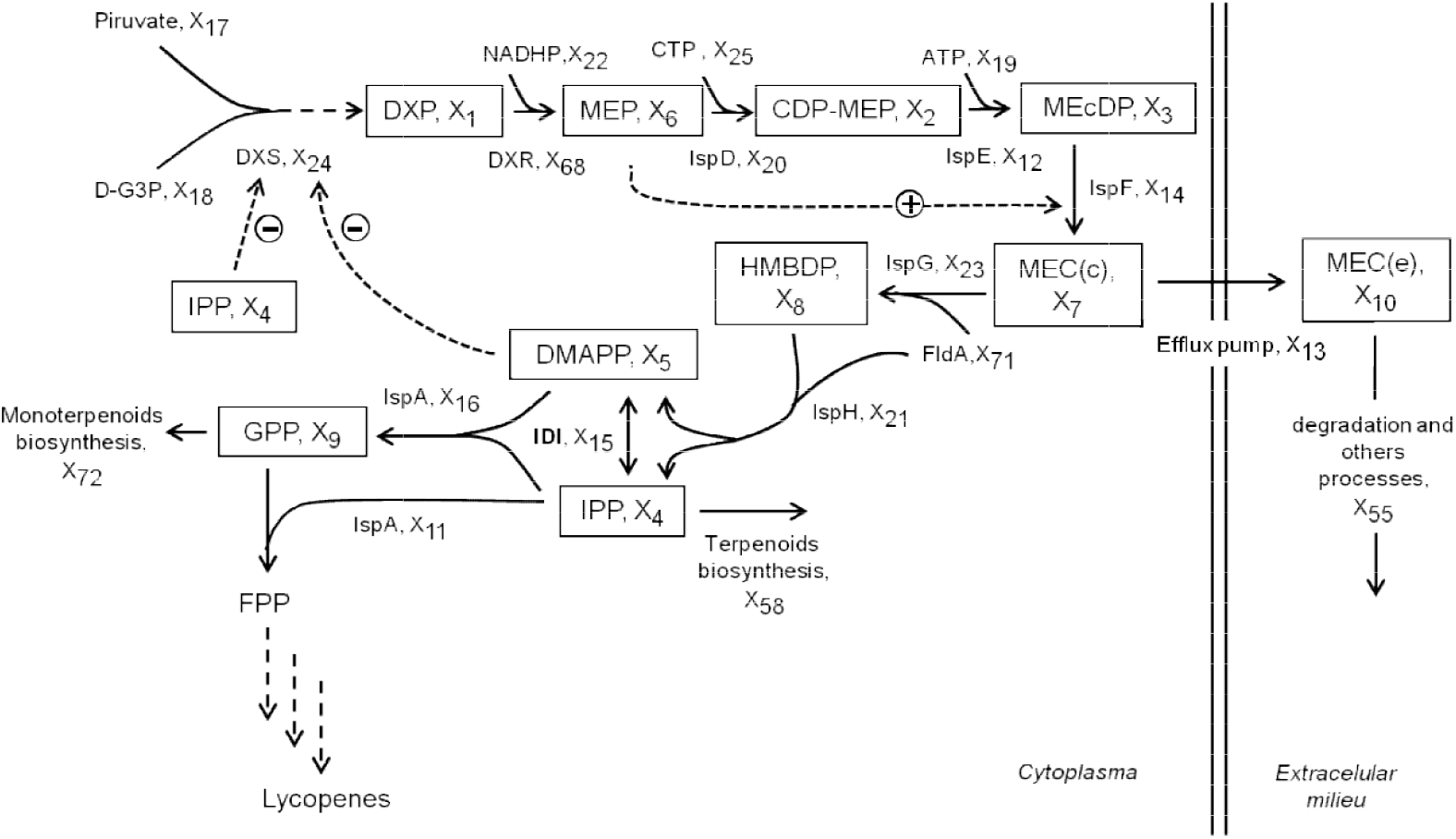
Mechanistic model of the 4-diphosfocytidyl-2C-methyl-D-erytritol-2-phosphate (MEP) isoprenoids biosynthesis pathway in *Escherichia coli*. Time dependent metabolites: (*X*_1_, DXP) 1-Deoxy-D-Xylulose 5-Phosphate, (*X*_2_, CDP-MEC) 4-diphosphocytidyl-2C-methyl D-erythritol, (*X*_3_, MEcDP) 4-diphosphocytidyl-2C-methyl D-erythritol 2-phosphate,(*X*_4_, IPP), Isopentenildiphoosphate, (*X*_5_, DMAPP) Dimethylallyl diphosphate, (*X*_6_, MEP) 2C-methyl-D-erythritol 4-phosphate, (*X*_7_, MEC(c)) Citosolic 4-difosfocitidil-2C-metil-D-eritritol-2-phosphate, (*X*_8_, HMBPP) Hydroxylmethylbutenyl diphosphate, (*X*_9_, GPP) Geranyl pyrophosphate, (*X*_10_, MEC(e)) 4-difosfocitidyl-2C-metil-D-eritritol-2-fosfato extracellular. Enzymes: (*X*_11_ and *X*_16_, IspA) farnesyl diphosphate synthase, (*X*_12_, IspE) MEcDP kinase, (*X*_13_) Efflux pump, (*X*_14_ IspF) MEC_(c)_ synthase, (*X*_15_, IDI) isopentenyl diphosphate isomerase, (*X*_20_, IspD) CDP-ME synthase, (*X*_21_, IspH) HMBDP reductase, (*X*_23_, IspG) HMBDP synthase, (*X*_24_, DXS) DXP synthase, (*X*_68_, DXR) DXP reductase, (*X*_55_). Degradation and biosynthetic processes: (*X*_58_) Terpenoids biosynthesis, (*X*_72_) Monoterpenoids biosynthesis. External metabolites: (*X*_17_) Pyruvate, (*X*_18_, D-G3P) D-glyceraldehyde 3-phosphate, (*X*_19_, ATP) Adenosine triphosphate, (*X*_22_, NADPH) Diphosphopyridindinucleotide phosphate reduced, (*X*_25_, CTP) Cytidine FldA) 5’-triphosphate, (*X*_71_, Flavodoxin-1

### 2.1. Model parameters

The model integrates all the available evidence on the metabolic steps, the kinetics of each process and the experimental values of intermediate concentrations (Tables S1.1 and S1.2).

### 2.2. Levels of metabolites external at the model

The model does not incorporate the central *E. coli* metabolism to describe dynamics of pyruvate, G3P, ATP, FldA, and NADPH. In the case of pyruvate there is evidence that *E. coli* chemostat culture supplemented with glucose maintains the intracellular levels inside a narrow range [22]. In the same vein it is assumed the G3P concentration is constant, since the supply of GAP and pyruvate for the MEP pathway in bacteria can be maintained through primary metabolism supply [23]. The level of D-G3P (*X*_18_) of 0.041 mM is equal at the reported Km of DXS for D-G3P assuming a maximal efficiency catalytic for the substrate-enzyme reaction [24]. As commented by Cleland, the Km of the enzymes is often useful to develop an idea of the physiological substrate concentration because: “While it is not an independent constant, it is a very useful one for the biochemist, since it is also a clue to the physiological level of the substrate. (A substrate concentration around Km utilizes most of the catalytic potential of the enzyme, while still maintaining proportional control; at high substrate levels the rate does not vary with substrate concentration, and one has no control” [24].

As a whole it is assumed all along the optimizations presented here that during continuous culture exponential growing phase and in presence of glucose the value of pyruvate, D-G3P, ATP, FldA, and NADPH does not change more than a 10% around their basal values.

### 2.3. Enzyme concentrations

The experimental data used were taken from the literature (see Tables S1.1 and S1.2). Enzymatic concentrations can be obtained from available *V*_*max*_’s or specific activities data. We decide use a direct approach for the enzyme levels calculation based in the number of enzyme molecules determinate by LC-MS/MS [25-26]. The concentrations (mM), were calculated as:

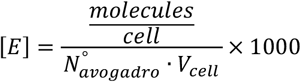

with the number of Avogadro 6.022140857 × 10^23^ molecules/mol and the *E. coli* volume 3.43 × 10^−1^ *L* ([27], Table 1 for complex medium). It is assumed that all the protein presents is in their active form i.e. not take into account the specific activity.

**Table 1.**
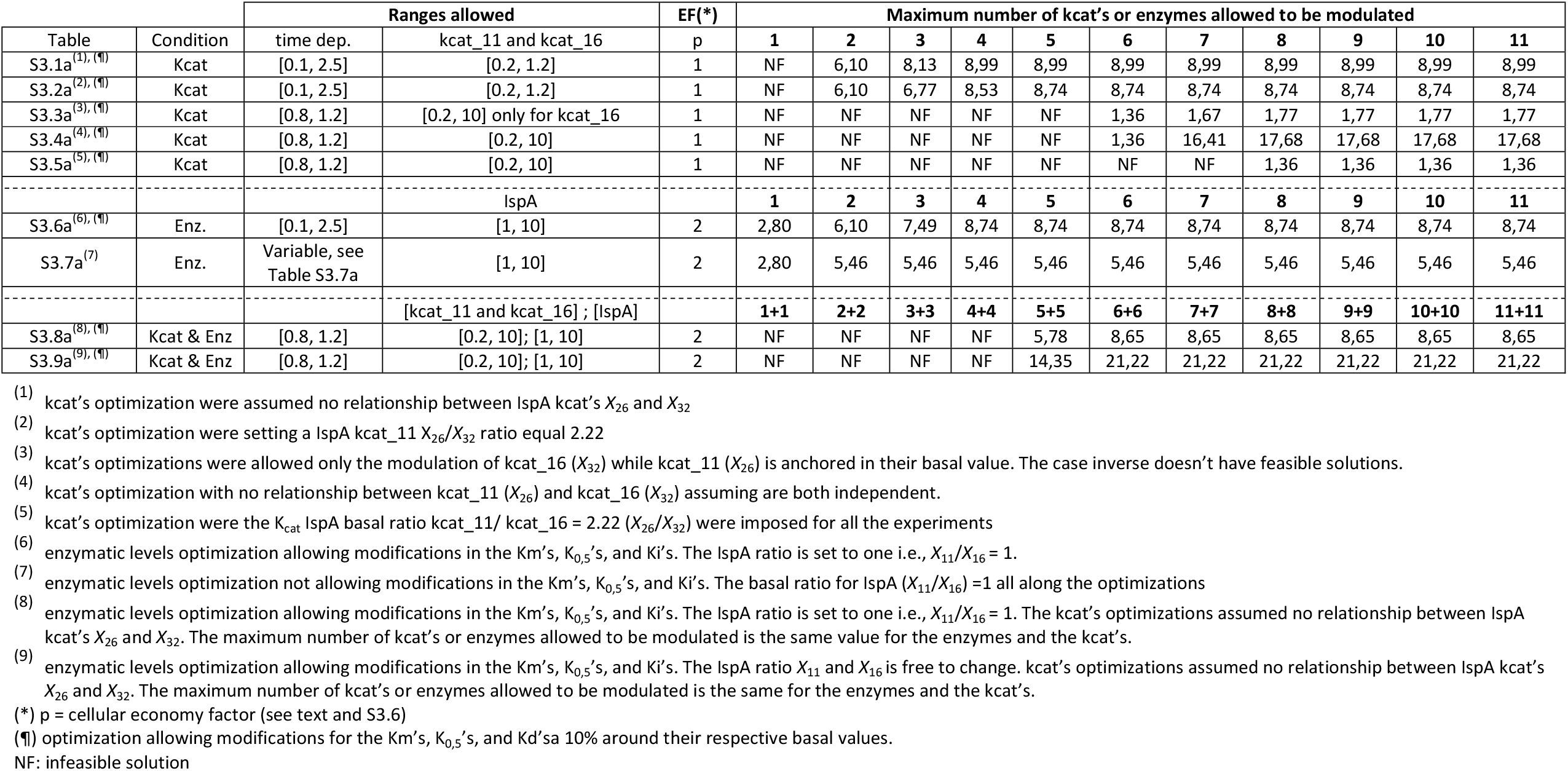
Summary of the optimization’s scenarios for the terpenoids flux maximization

### 2.4. Enzymatic mechanisms

Based in the reported 1-deoxy-D-xylulose 5-Phosphate synthase mechanism (DXS, *X*_24_) [28-30] their kinetics can be described as an ordered sequential mechanism in which the inhibitors IPP (*X*_4_) and DMAPP (*X*_5_) can compete with either the first substrate pyruvate for the free enzyme, or with the second substrate D-G3P for the complex pyruvate-enzyme, or both [31].

The feedback inhibition regulatory control mechanism exerted by IPP (*X*_4_) and DMAPP (*X*_5_) [23, 28] over the DXS thiamine diphosphate (ThDP) site [30] are in the low micromolar range (60–80 mM), having a significant binding ability to ThDP under physiological conditions [23].

Since the IPP (*X*_4_) and DMAPP (*X*_5_) inhibitions are reduced when one of them bind the DXS-ThDP [28] they were weighted according with the following expression:

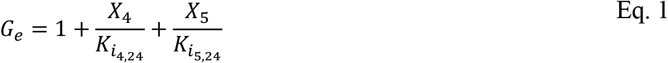

As commented above, the kinetics implemented was an ordered sequential mechanism with pyruvate binding first in and the inhibitor competing with either the first substrate for the free enzyme or with the second substrate for the complex enzyme-substrate ([31], Case VI).

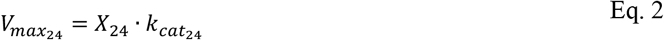

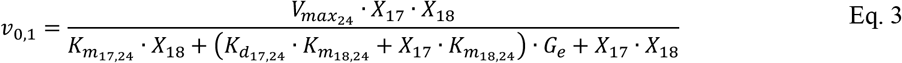

Finally, the model also includes the MEP (*X*_6_) nonessential activation over IspF (*X*_14_), a reaction that can take place even in absence of MEP [23].

### 2.5. V_max_ model assumptions

In the Biochemical System Theory (BST) formalism the enzymes had been represented assuming a direct proportionality between *V*_max_ or specific activity and its concentrations [32-36], since the *in vivo V*_max_ correlates with the *in vitro* value when measured under optimal conditions [37-39]. This direct proportionality assumption is mathematically correct within a range of enzyme concentration; however, it had been shown a maximum of enzyme functionality concentration is attained in the cell within a certain range of values [40].

To overcome this difficulty and for practical biotechnological applications we split the *V*_max_ in their two components named the enzymatic concentration and their kcat and treat them as two separated model parameters.

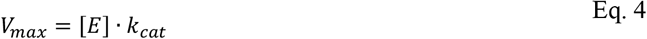

This procedure allows the variation of the kcat without affecting the total enzyme amount. Since we are assuming direct proportionality between *V*_max_ and enzyme concentration [E], in the power-law representation the kinetic order of [E] and kcat are both one [32, 35, 41-42].

The model finish with the IspA enzyme (*X*_11_) and (*X*_16_) in two consecutive bisubstrate reactions [7, 43], the first one IPP (*X*_4_) + DMAPP (*X*_5_) follow a ping-pong-bi-bi kinetics mechanism and the second one GPP (*X*_9_) + IPP (*X*_4_) follow an ordered bi-bi reaction, with initial GPP binding a terpenoids elongation rending farnesyl diphosphate [43-44]. For mathematical proposes we settled different numbers for the same IspA enzyme, In some experiments their values are similar but they also had been settle separately in order to fulfill the simulations presented along the manuscript (Table 1).

The IDI (*X*_15_) requires a divalent metal, either Mg^+2^ or Mn^+2^, for the turnover activity. The eukaryotic enzymes, with kcat between 1.8 to 11 s^-1^ have more active catalysts than *E. coli* isomerase with 0.33 s^-1^. However, all of them have similar catalytic efficiencies (kcat/Km) because the Michaelis constants of the bacterial isomerases are lower than those of their eukaryotic counterparts [45-47]. Similar catalytic efficiencies for the other isomerases suggest that the interconversion of IPP and DMAPP is not normally a rate-limiting reaction in isoprenoid metabolism [46].

## 3. THEORY/CALCULATION

### 3.1. Model representation

The model is represented by a set of power-law ordinary differential equations [48-51]. The S-system model consists in metabolic node flux aggregation and power-law approximations for the variables [52]. For each dependent variable we have an equation like

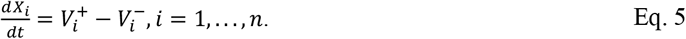

In this expression 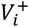 and 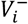 represent the aggregation of the inflows and outflows fluxes of each metabolic node respectively. After translating the above expression to the power law formalism we obtain:

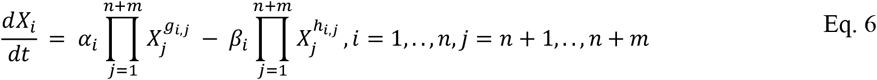

The indices *i*= 1,..,*n* refer to dependent variables, while *j* = *n* + 1,…, *n* + *m* refer to independent variables or parameters. The parameters *α*_*i*_ and *g*_*i,j*_ are the rate constants and kinetic orders associated with the rate law for net production of *X*_*i*_. Similarly, *β*_*i*_and *h*_*i,j*_ are associated with the rate law for net transformation of *X*_*i*_. The kinetic order parameters *g*_*i,j*_ and *h*_*i,j*_ are defined as follows:

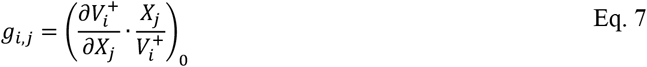

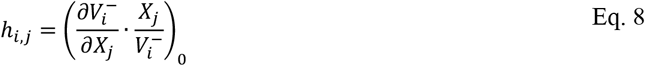

The supplementary materials include the model kinetics equations (S1.1), power-law equations (S1.3), and the S-system numeric model (S1.4).

### 3.2. Model verification

The model and optimal solutions were tested in their stability showing a stable steady state with oscillations (Table S1.3). The model robustness is checked by computing the sensitivities of variables (Tables S2.1, S2.3, S2.4, S2.5) and fluxes (Tables S2.2, S2.6, S2,7, S2.8). All of them show reasonable low values.

### 3.3. Optimization strategy

We aim to determinate profiles of enzyme concentrations and catalytic constants that maximize the biosynthetic flux through the terpenoids precursors (Fig. S1). Provided with this information we are in conditions to implement rational biotechnological changes to the increase the desired target flux. The optimization procedure is based on the Indirect Optimization Method [53-55]. Subsequently, based on this model we designed optimization strategies based on the modification of enzyme concentrations and/or their kcat’s.

Specifically the objective function was to maximize the flux synthesis of terpenoids *ν*_4,11_ through the IspA (*X*_11_) (see Fig. S1).

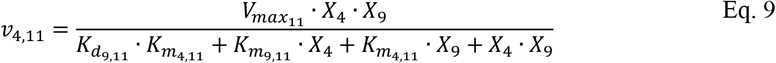

Where the 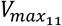 is given by 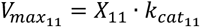. With 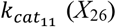, the 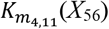 and 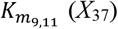 represents the *K*_*m*_ for IPP (*X*_4_) and GPP (*X*_9_) respectively and 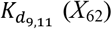 represents the dissociation constant for GPP.

The Eq. (9) in power-law terms read as

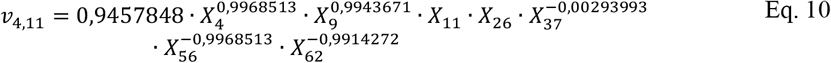

and after log-transforming Eq. 10 read as

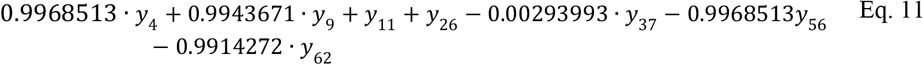

where *y*_*i*_ = *lnX*_*i*_ for *i* {4,9,11,26,37,56,62}

In order to guarantee cellular viability we imposed additional constraints in the form of ranges (between a minimum, LB, and a maximum, UB, bound) around their respective basal values for the metabolic intermediates, enzyme concentrations, K_m’s_ and K_d’s_ parameters.

Because the kcat compromises the enzymatic structure is predictable a concomitant change in the associated Km’s and/or Kd’s as in fact have been reported for various enzymes ([56], Table 1). This Km change can affect the model if the bioengineered enzyme is a mutant as well as if the enzyme inserted comes from another organism. To simulate this possible scenario we allowed variations of the Km’s and Kd’s when the corresponding kcat were allowed to optimize (Tables S3.1a and S3.4a). We permit variations in the Km’s and Kd’s of a 10% up-down their basal value, a variation that is frequently reported within the Km experimental error [29, 57-59].

Assuming unlimited supply of energy sources and redox agents, a probable scenario in bioreactors under continuous growing conditions, the external variables not controlled by the MEP model and principally coming from the central metabolism are pyruvate (*X*_17_), G3P (*X*_18_), ATP (*X*_19_), NADPH (*X*_22_), CPT (*X*_25_), and FLdA (*X*_71_) [23]. These were allowed to bascule a 10% around theirs basal values.

In the case of the enzymes optimization (*X*_11,_…,.*X*_16_, *X*_20_, *X*_21_, *X*_23_, *X*_24_, *X*_68_), the boundaries allowed were between 1 and 10 fold their basal value (Tables S3.5, S3.2, and S3.4a). The reason for the lower range of one (their basal value) and not lower is that a practical implementation of results with values lower than the basal enzyme concentration will be difficult to achieve.

For the kcat optimizations (*X*_26,_…,*X*_36_) they were allotted with upper bounds of 10-fold the basal value, as the enzymes, but the lower bounds were of 0.1 fold the basal value (Table S3.1a). A value under the basal one is not a problem theoretically because it is possible to insert mRNA of enzymes exerting similar function to the native one but with lower kcat. All these restrictions mentioned above are detailed in the supplementary section S3.

A final restriction was imposed at the model: an upper limit constraint to the total amount of enzyme, that we call *cellular economy factor* (p), (see [23, 32, 41]). This constraint is intended to maintain the cell viability in the optimum profile avoiding unrealistic total cell enzyme concentration (see S3.7 section for details).

## 4. RESULTS

From a set of initial values for the time-dependent metabolites we run our model up to reach an steady-state. Table S1.1 shows these initial and final values. The steady-state point was the initial point for the optimization experiments performed next. It can be observed that all the steady-state values are inside the referenced ranges which are an indication in favor of the model robustness.

The reliability assessment show that the model is stable (Table S1.3) and the analysis of its robustness indicates reasonable low values for their variables sensitivities (Tables S2.1, S2.3, S2.4, and S2.5) as well as for the fluxes sensitivities (Tables S2.2, S2.6, S2.7, and S2.8).

Table 1 presents the output optimization ratio increment 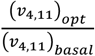 basal according to the number of enzymes, kcat or both allowed to be modulated. In all cases the optimized profiles were back at the original power-law S-system representation (see section S1.3) showed be stable by their corresponding negative eigen-values real parts. The optimized profiles back at the kinetic ODE’s equations lead lower values than their respective optimized outputs. This is caused principally for the readjusting of the time dependent variables present in the objective function to be maximized by the linear optimization program (not shown).

The kcat optimization approach shows to be more efficient because it allows to by-pass the end of the route bottleneck without requiring to incorporate mutants of the IspA enzyme (results in Table S3.2a) and produces bigger or similar results than the enzymatic levels modification (see Tables 3.6a and S3.7a). In addition, the kcat optimization it does not require enzymatic over-expression of enzymes in order to maximize the flux in terpenoids direction.

Table S3.1a shows the optimum profile for the kcat when simultaneously it is allowed to pivot 20% the corresponding Km’s and Kd’s and assuming no relationship between the IspA kcat_11 and kcat_16. In Table S3.1a, when up to 10 variables are allowed to be modulate a 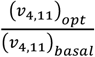 a flux ratio increment of 8.74 is reported for lycopenes direction. The interesting point is that this optimum profile indicates a decreasing of DXS kcat_24, IspD kcat_20 and IspF kcat_14 to a 29%, 40% and a 24% of their basal values, respectively, while IDI kcat_21 increase 10 fold its basal value, also IspG have a 59% of decrease in this case. All the above results at first glance are a counterintuitive result.

However it has been reported that a strong concomitant overexpression of four enzymes (DXS, IDI, IspD, and IspF) lead to the inhibition of the terpenoids production [13]. Also, an overexpression of IspG could be expected to bring to a concomitant decrease of the MEC efflux and to an increasing of the isoprenoid production. However, it has been reported that in *Mycobacterium tuberculosis* a combined increasing of DXS (*X*_24_) and IspG (*X*_23_) produce an accumulation of HMBDP (*X*_8_) and does not rise the terpenoids flux [60].

In Tables S3.1a and S3.2a the range of variation for the kcat’s during the optimization was settled between [0.2, 10] fold with the exception of the IspA kcat which were maintained within [0.2, 1.2]. The reason to keep the kcat IspA upper bound in 1.2 was to limit the maximum differences for the two IspA kcat to values lower than 300 fold because these two kcat are strongly affected during the optimizations with already basal fold differences important of kcat_11/ kcat_16 = 46.4 and probably this could be a limitation for future implementation of the results presented here. In the rest of the tables were the Ispa kcat (*X*_26_, *X*_32_) were modulated the range of variation was similar to the others kcat (Table 1).

The catalytic efficiencies of the optimum points in Table S3.1a are shown in Table S3.1b. Here it can be seen a generalized trend of lower values caused by an increasing in the Km’s in conjunction with a decreasing of the kcat. The sole exceptions at these low trend are the kcat_11/Km_4_11, kcat_11/Km_9_11 and IDI kcat_15/Km_5_15, which show kcat/Km ratios bigger than one. This last three exceptions are expected as is reported that a simultaneous decrement in the Km’s and increment in the kcat observed in mutants is able to increase the efficiency of the enzyme ([56], Table 1).

It is well known that the increasing in the specific constant (SC) enhance the reaction of the enzymes [24, 56]. But this is not the case for the MEC mechanism, in general the SC in the core of the model maintains the same or even decrease their value. This phenomenon can be appreciated in Tables S3.1b, S3.2b, S3.3b, S3.4b, S3.5b, S3.6b, S3.8b, and S3.9b. In the case of Tables S3.6b the changes are less pronounced because only was allowed modify the Km and Kd but not the kcat. In the same tables above mentioned there are three enzymes that enhance the SC and are the IDI (*X*_15_), the IspA (*X*_11_), and the efflux pump (*X*_13_) these three enzymes promote the increase of the flux *v*_4,11_ as the flux pump decrease the levels of metabolites in then pathway, and IspA (*X*_11_) is direct, the function of IDI is less clear as this reversible enzyme increase the conversion of DMAPP (*X*_5_) in IPP (*X*_4_) and vice versa. It is reported for the *in vitro* IDI a predominant formation of DMAPP (66%) with respect IPP (34%) at the equilibrium [47].

The reported increase in the terpenoids production after overexpression of IspG [13] no match with the results presented here, although in Zhou this trend is observed in a genetically modified strain (with overexpressed levels of dxs, idi, ispD and ispF).

The optimization profiles proposing MEC extrusion (Tables S3.1a, S3.3, S3.2, and S3.4a) are difficult to translate to a biotechnological real process since the MEC transports are still uncharacterized and there is a diversity of substrates for the transport systems [61]. It is thus cumbersome to pin-point a single transporter as a target to be modified.

It has been found that the *fsr* antiporter [13] is a putative MEC transporter. However, as commented above, MEC transporters modifications are difficult to implement since they comprise a synergistic and promiscuous family that can overtake the functions of other transporters in case one be impaired.

To analyze enzymatic insertions in *E. coli* the IspA kcat_11 (*X*_26_) and kcat_16 (*X*_32_) are allowed be modulated independent one of the other simulating two different enzymes (Tables S3.1a, S3.3a, and S3.4a). Also in Table S3.9a the IspA kcat (*X*_26_ and *X*_32_) and the enzymatic levels of IspA (*X*_16_ and *X*_11_) were treated independently between them. Additionally in the tables above mentioned the related Km’s and Kd’s were allowed bascules 20% around their basal value which implies additional enzymatic modifications. The only experiment that respects the WT strain profile allowing only increase in the enzymatic levels is represented in Table S3.7a.

In the single approach where the kcat (Tables S3.1a,…,S3.5a) or the enzymes (Tables 3.6a and 3.7a) were modulated by separate, the optimization with higher increase in the flux *v*_4,11_ is presented in Table S3.4a with a 17.68 fold increase in the basal flux direction lycopenes for six enzymes or kcat allowed be modulate. Please note that in Table S3.4a the range for the time dependent metabolites is restricted to 20% their respective basal values to guarantee the optimized metabolites be inside the ranges reported, this is probably the cause of many situation where the optimization rend non feasible solutions (Table 1).

The optimization mixed approach from Table S3.9a with not relationship between the enzymatic IspA (*X*_11_ and *X*_16_) nor between the IspA kcat (*X*_26_ and *X*_32_), present the higher levels of flux *v*_4,11_ direction lycopenes. In this experiment is reached an increment of the flux of 21.22 the basal value for the experiments equal or bigger than six enzymes and six kcat allowed to be modulated simultaneously. In general, can be observed that the IspA (*X*_16_) and the kcat_16 (*X*_32_) tend to decrease or be smaller than the IspA (*X*_11_) and kcat_11 (*X*_26_). This experiments cold be an indicative of the *E. coli* lycopene flux after two independents IspA insertions with different expression levels and specific activities in the bacteria flux via lycopenes.

Table S3.3a shows the optimized results when the IspA kcat_16 (*X*_32_) is allowed modulated keeping in their basal value the IspA kcat_11 (*X*_26_). In this case the increments of flux *v*_4,11_ where modest. The contrary situation i.e. allowing modulate IspA kcat_11 (*X*_26_) and keeping the basal value for IspA kcat_16 (*X*_32_) do no rend any feasible better solution.

## 5. DISCUSSION

Based on the information available, we have built an *E. coli* mathematical bottom-up model of the MEP biosynthetic route leading terpenoids. That includes MEC cell extrusion membrane transport. Since the mechanisms involved in the MEC extrusion are not yet fully characterized [13] here is described as a sole transmembrane protein pump in reference to the putative MEC (*X*_7_) transports candidates found in the literature.

The model was first tested in their stability and robustness and secondly was used for global rational optimizations aiming to define the optimum parameter profiles yielding the maximum flux direction terpenoids synthesis. The optimization profiles restrictions conditions guarantee cell viability avoiding the model parameters run out of physiological ranges.

We developed a new mathematical tool based on the modification of the kcat. This kcat based optimization strategy has the advantage that avoids the scenario of protein overproduction and thus the decreasing in the proteins functionality as consequence, between others, of increment of nonfunctional nascent unfolded proteins [40]. Moreover the kcat optimization opens the possibility to rational incorporation of other organism or mutant strains in the *E. coli* genome with minimal perturbation of the complex homeostasis perturbation that represent the change in the levels for the enzymes. Our results show that it is possible to obtain an increased direction terpenoids flux without the need of increasing the total amount of enzyme concentration, but simply by modulating their kcat (Tables 1, S3.1a, S3.1b).

The potential of the kcat based optimization strategy can be fully appreciated in Table 1 were can be seen a maximum predicted flux of 17.68 folds the basal value for the kcat + Km’s modulation (also see Table S3.4a). When the enzymes concentrations were allowed to be modified as in Table 1, S3.6a and S3.7a a 8.74 and 5.46 fold basal increase is obtained respectively. A dual change of the enzymatic level for IspA (*X*_11_ and *X*_16_) + IspA kcat (*X*_26_ and *X*_32_) + Km’s modifications can yield a flux increase of 21.22 fold the basal *ν*_4,11_ flux value (Table S3.9a) but this modifications imply different IspA for *X*_11_ and *X*_16_.

Interestingly, we found out that the kcat modification is a better strategy than the enzymes increase in order to decrease the MEC’s efflux pump (*X*_13_) (Tables 3.1a, SS3.2a, S3.3a, S3.4a, S3.5a, S3.8a, S3.9a) in comparison with the enzymatic levels modifications (Tables S3.6a and S3.7a).

In summary, to attain a precise evaluation of mechanisms proposed for terpenoids biosynthesis, further studies aimed at the elucidation of the processes involved with wild and recombinant strains are necessary. This information would allow the energetic (ATP), reductive (NADH, NADPH), and CoA dynamics to be accounted for. However, we think that the type of analysis carried-out in this work constitutes a key feature in any model building project. It is expected that the observed results will stimulate the next phase of model amendments and refinements using experimental work being undergone within our labs.

## 6. CONCLUSIONS

The *E*. coli route of synthesis of licopenes through the alternative mevalonate pathway was mathematically modeled and optimized. The model analysis showed that the MEC extrusion guarantees bacterial homeostasis and cell viability. In addition, it identified the enzyme IspA as a bottleneck of terpenoid biosynthesis. The application of the method allows different strategies to overcome the enzymatic flux restriction, by either modifying the IspA kcat or by introducing new enzymes with parallel functions. Through the kcat modification it is possible to increases significantly the terpenoid biosynthetic flux, through the modification of up to 8 kcat values and the corresponding Km and Kd, but without change the cell’s total enzyme concentration. The comparison of this strategy with the approach of overexpressingthe enzyme activity showed that the combination of both (overexpression of enzyme concentration and kcat modifications) is the most effective one.

## DECLARATIONS

a. Declaration of Competing Interest
b. Not applicable
c. Author contributions
d. FAV, CGA and NVT designed the model and the optimization strategy and wrote the corresponding parts of the manuscript. FAV also performed the model analyses and implemented the optimization strategy. JGJ, TdD and MC contributed to the experimental data and collaborated in the writing of the manuscript.

## Acknowledgments

At the professional writing services.

## Funding

This work was supported by grants from MICINN BIO2014-54411-C2-1-R, RTI2018-094393-B-C21-MCIU/AEI/FEDER, UE, which includes ERDF European cofunding and the Seneca Foundation CARM (19236/PI/14).

## ABREVIATIONS

*E. coli*: *Escherichia coli*
*S. cerevisiae*: *Saccharomyces cerevisiae*
MEP: 4-diphosphocytidyl-2C-methyl-D-erytritol-2-phosphate
BST: biochemical system theory
Kcat: catalytic constant
Km: Michaelis-Menten constant
Kd: dissociation constant
Ka: activation constant
DXP: 1-deoxy-D-xylulose 5-phosphate
CDP-MEC: 4-diphosphocytidyl-2C-methyl D-erythritol
MEcDP: 4-diphosphocytidyl-2C-methyl D-erythritol 2-phosphate
IPP: isopentenildiphosphate
DMAPP: dimethylallyl diphosphate
MEP: 2C-methyl-D-erythritol 4-phosphate
MEC(c): cytosolic 4-difosfocitidil-2C-metil-D-eritritol-2-phosphate
HMBPP: hydroxylmethylbutenyl diphosphate
GPP: geranyl pyrophosphate
MEC(e): 4-difosfocitidyl-metil-metil-D-difosyl-fosfatol-extracellular
IspA: farnesyl diphosphate synthase
IspE: MEcDP kinase
IspF: MEC_(c)_ synthase
IDI: isopentenyl diphosphate isomerase
IspD: CDP-ME synthase
IspH: HMBDP reductase
IspG: HMBDP synthase
DXS: DXP synthase
DXR: DXP reductase
D-G3P: dextro-glyceraldehyde 3-phosphate
ATP: adenosine triphosphate
NADPH: diphosphopyridindinucleotide phosphate reduced
CTP: cytidine 5’-triphosphate
FldA: flavodoxin-1
SC: specific constant

## Notes

### Competing Interest Statement

The authors have declared no competing interest.

## REFERENCES

1. Tholl D. Biosynthesis and biological functions of terpenoids in plants. Adv Biochem Eng Biotechnol. 2015;148:63–106; doi: 10.1007/10_2014_295.

2. Gallego Jara J, Lozano Terol G, Sola RA, Canovas Diaz M, de Diego T. Engineering of microbial cell factories for production of plant-based natural products. Elseiver, Academic Press; 2020.

3. Schempp FM, Drummond L, Buchhaupt M, Schrader J. Microbial Cell Factories for the Production of Terpenoid Flavor and Fragrance Compounds. J Agric Food Chem. 2018;66(10):2247–58; doi: 10.1021/acs.jafc.7b00473.

4. Moser S, Pichler H. Identifying and engineering the ideal microbial terpenoid production host. Appl Microbiol Biotechnol. 2019;103(14):5501–16; doi: 10.1007/s00253-019-09892-y 10.1007/s00253-019-09892-y [pii].

5. Theisen M, Liao JC. Industrial Biotechnology: Escherichia coli as a Host. In: Wittmann aL, editor. Industrial biotechnology. 2017. p. 182.

6. Li M, Nian R, Xian M, Zhang H. Metabolic engineering for the production of isoprene and isopentenol by Escherichia coli. Appl Microbiol Biotechnol. 2018;102(18):7725–38; doi: 10.1007/s00253-018-9200-510.1007/s00253-018-9200-5 [pii].

7. Ward VCA, Chatzivasileiou AO, Stephanopoulos G. Metabolic engineering of Escherichia coli for the production of isoprenoids. FEMS Microbiol Lett. 2018;365(10); doi: 4953741 [pii] 10.1093/femsle/fny079.

8. Ajikumar PK, Xiao WH, Tyo KE, Wang Y, Simeon F, Leonard E, et al. Isoprenoid pathway optimization for Taxol precursor overproduction in Escherichia coli. Science. 2010;330(6000):70–4; doi: 330/6000/70 [pii] 10.1126/science.1191652.

9. Kim SW, Keasling JD. Metabolic engineering of the nonmevalonate isopentenyl diphosphate synthesis pathway in Escherichia coli enhances lycopene production. Biotechnol Bioeng. 2001;72(4):408–15; doi: 10.1002/1097-0290(20000220)72:4<408::AID-BIT1003>3.0.CO;2-H [pii] 10.1002/1097-0290(20000220)72:4<408::aid-bit1003>3.0.co;2-h.

10. Martin VJ, Pitera DJ, Withers ST, Newman JD, Keasling JD. Engineering a mevalonate pathway in Escherichia coli for production of terpenoids. Nat Biotechnol. 2003;21(7):796–802; doi: 10.1038/nbt833nbt833 [pii].

11. Rodriguez-Villalon A, Perez-Gil J, Rodriguez-Concepcion M. Carotenoid accumulation in bacteria with enhanced supply of isoprenoid precursors by upregulation of exogenous or endogenous pathways. J Biotechnol. 2008;135(1):78–84; doi: S0168-1656(08)00094-1 [pii] 10.1016/j.jbiotec.2008.02.023.

12. Yoon SH, Lee YM, Kim JE, Lee SH, Lee JH, Kim JY, et al. Enhanced lycopene production in Escherichia coli engineered to synthesize isopentenyl diphosphate and dimethylallyl diphosphate from mevalonate. Biotechnol Bioeng. 2006;94(6):1025–32; doi: 10.1002/bit.20912.

13. Zhou K, Zou R, Stephanopoulos G, Too HP. Metabolite profiling identified methylerythritol cyclodiphosphate efflux as a limiting step in microbial isoprenoid production. PLoS One. 2012;7(11):e47513; doi: 10.1371/journal.pone.0047513PONE-D-12-14608 [pii].

14. Martinez-Jaramillo LM: Structural and functional study of efflux pumps involved in drug resistance. In: Agricultural Sciences. vol. Doctor in Biochemistry. Lyon, France: Université Claude Bernard; 2014.

15. Nagano K, Nikaido H. Kinetic behavior of the major multidrug efflux pump AcrB of Escherichia coli. Proc Natl Acad Sci U S A. 2009;106(14):5854–8; doi: 0901695106 [pii] 10.1073/pnas.0901695106.

16. Nikaido H, Normark S. Sensitivity of Escherichia coli to various beta-lactams is determined by the interplay of outer membrane permeability and degradation by periplasmic beta-lactamases: a quantitative predictive treatment. Molecular microbiology. 1987;1(1):29–36; doi: 10.1111/j.1365-2958.1987.tb00523.x.

17. Weingart H, Petrescu M, Winterhalter M. Biophysical characterization of in-and efflux in Gram-negative bacteria. Curr Drug Targets. 2008;9(9):789–96; doi: 10.2174/138945008785747752.

18. Patel H, Nemeria NS, Brammer LA, Freel Meyers CL, Jordan F. Observation of thiamin-bound intermediates and microscopic rate constants for their interconversion on 1-deoxy-D-xylulose 5-phosphate synthase: 600-fold rate acceleration of pyruvate decarboxylation by D-glyceraldehyde-3-phosphate. J Am Chem Soc. 2012;134(44):18374–9; doi: 10.1021/ja307315u.

19. Gracia JB. Regulación de la ruta del 1-deoxi-d-xylulose-5-fosfato (DXP) en Escherichia coli recombinante para la producción de licopeno. Facultad de Quimica Departamento de Bioquímica y Biología Molecular (B) e Inmunología. 2014;Máster en Química Fina y Molecular:54.

20. Wang XA, Kurra Y, Huang Y, Lee YJ, Liu WR. E1-catalyzed ubiquitin C-terminal amidation for the facile synthesis of deubiquitinase substrates. Chembiochem. 2014;15(1):37–41; doi: 10.1002/cbic.201300608.

21. Ye L, Lv X, Yu H. Engineering microbes for isoprene production. Metab Eng. 2016;38:125–38; doi: S1096-7176(16)30058-1 [pii] 10.1016/j.ymben.2016.07.005.

22. Yang YT, Bennett GN, San KY. The effects of feed and intracellular pyruvate levels on the redistribution of metabolic fluxes in Escherichia coli. Metab Eng. 2001;3(2):115–23; doi: 10.1006/mben.2000.0166S1096-7176(00)90166-6 [pii].

23. Banerjee A, Sharkey TD. Methylerythritol 4-phosphate (MEP) pathway metabolic regulation. Nat Prod Rep. 2014;31(8):1043–55; doi: 10.1039/c3np70124g.

24. Cleland WW. The Enzymes. In: Academic Press, Inc., New York and London; 1970. p. 1–65.

25. Soufi B, Krug K, Harst A, Macek B. Characterization of the E. coli proteome and its modifications during growth and ethanol stress. Front Microbiol. 2015;6:103; doi: 10.3389/fmicb.2015.00103.

26. Volke DC, Rohwer J, Fischer R, Jennewein S. Investigation of the methylerythritol 4-phosphate pathway for microbial terpenoid production through metabolic control analysis. Microb Cell Fact. 2019;18(1):192; doi: 10.1186/s12934-019-1235-5 10.1186/s12934-019-1235-5 [pii].

27. Volkmer B, Heinemann M. Condition-dependent cell volume and concentration of Escherichia coli to facilitate data conversion for systems biology modeling. PLoS One. 2011;6(7):e23126; doi: 10.1371/journal.pone.0023126PONE-D-10-04797 [pii].

28. Banerjee A, Wu Y, Banerjee R, Li Y, Yan H, Sharkey TD. Feedback inhibition of deoxy-D-xylulose-5-phosphate synthase regulates the methylerythritol 4-phosphate pathway. J Biol Chem. 2013;288(23):16926–36; doi: M113.464636 [pii] 10.1074/jbc.M113.464636.

29. Brammer LA, Smith JM, Wade H, Meyers CF. 1-Deoxy-D-xylulose 5-phosphate synthase catalyzes a novel random sequential mechanism. J Biol Chem. 2011;286(42):36522–31; doi: M111.259747 [pii] 10.1074/jbc.M111.259747.

30. White JK, Handa S, Vankayala SL, Merkler DJ, Woodcock HL. Thiamin Diphosphate Activation in 1-Deoxy-d-xylulose 5-Phosphate Synthase: Insights into the Mechanism and Underlying Intermolecular Interactions. J Phys Chem B. 2016;120(37):9922–34; doi: 10.1021/acs.jpcb.6b07248.

31. Cheng Y, Prusoff WH. Relationship between the inhibition constant (K1) and the concentration of inhibitor which causes 50 per cent inhibition (I50) of an enzymatic reaction. Biochem Pharmacol. 1973;22(23):3099–108; doi: 0006-2952(73)90196-2 [pii] 10.1016/0006-2952(73)90196-2.

32. Alvarez-Vasquez F, Gonzalez-Alcon C, Torres NV. Metabolism of citric acid production by Aspergillus niger: model definition, steady-state analysis and constrained optimization of citric acid production rate. Biotechnol Bioeng. 2000;70(1):82–108.

33. Alvarez-Vasquez F, Riezman H, Hannun YA, Voit EO. Mathematical Modeling and Validation of the Ergosterol Pathway in Saccharomyces cerevisiae. PLoS One. 2011;6(12):e28344; doi: 10.1371/journal.pone.0028344PONE-D-11-07746 [pii].

34. Alvarez-Vasquez F, Sims KJ, Cowart LA, Okamoto Y, Voit EO, Hannun YA. Simulation and validation of modelled sphingolipid metabolism in Saccharomyces cerevisiae. Nature. 2005;433(7024):425–30.

35. Shiraishi F, Savageau MA. The tricarboxylic acid cycle in Dictyostelium discoideum. I. Formulation of alternative kinetic representations. J Biol Chem. 1992;267(32):22912–8.

36. Sorribas A, Savageau MA. A comparison of variant theories of intact biochemical systems. I. Enzyme-enzyme interactions and biochemical systems theory. Math Biosci. 1989;94(2):161–93; doi: 0025-5564(89)90064-3 [pii].

37. Newsholme EA, Crabtree B. Maximum catalytic activity of some key enzymes in provision of physiologically useful information about metabolic fluxes. J Exp Zool. 1986;239(2):159–67; doi: 10.1002/jez.1402390203.

38. Newsholme EA, Crabtree B, Zammitt VA. Use of Enzyme Activities as Indices of Maximum Rates of Fuel Utilization. In: Ciba Foundation Symposium 73 - Trends in Enzyme Histochemistry and Cytochemistry. John Wiley & Sons, Ltd.; 2008. p. 245–58.

39. Suarez RK, Staples JF, Lighton JR, West TG. Relationships between enzymatic flux capacities and metabolic flux rates: nonequilibrium reactions in muscle glycolysis. Proc Natl Acad Sci U S A. 1997;94(13):7065–9.

40. Sharma AK, O’Brien EP. Increasing Protein Production Rates Can Decrease the Rate at Which Functional Protein Is Produced and Their Steady-State Levels. J Phys Chem B. 2017;121(28):6775–84; doi: 10.1021/acs.jpcb.7b01700.

41. Torres NV, Alvarez-Vasquez F, García J. Metabolism of citric acid production by A. niger: matematical modeling and optimization. In: Citric Acid: Occurrence, Biochemistry, Applications and Processing. Nova Science Publishers, Inc.; 2014. p. 129, 6×9 - (NBC-R).

42. Voit EO. Computational analysis of biochemical systems: a practical guide for biochemists and molecular biologists. New York: Cambridge University Press; 2000.

43. Ku B, Jeong JC, Mijts BN, Schmidt-Dannert C, Dordick JS. Preparation, characterization, and optimization of an in vitro C30 carotenoid pathway. Appl Environ Microbiol. 2005;71(11):6578–83; doi: 71/11/6578 [pii] 10.1128/AEM.71.11.6578-6583.2005.

44. Ramamoorthy G, Pugh ML, Tian BX, Phan RM, Perez LB, Jacobson MP, et al. Synthesis and enzymatic studies of bisubstrate analogues for farnesyl diphosphate synthase. J Org Chem. 2015;80(8):3902–13; doi: 10.1021/acs.joc.5b00202.

45. Durbecq V, Sainz G, Oudjama Y, Clantin B, Bompard-Gilles C, Tricot C, et al. Crystal structure of isopentenyl diphosphate:dimethylallyl diphosphate isomerase. Embo J. 2001;20(7):1530–7; doi: 10.1093/emboj/20.7.1530.

46. Hahn FM, Hurlburt AP, Poulter CD. Escherichia coli open reading frame 696 is idi, a nonessential gene encoding isopentenyl diphosphate isomerase. J Bacteriol. 1999;181(15):4499–504.

47. Jonnalagadda V, Toth K, Richard JP. Isopentenyl diphosphate isomerase catalyzed reactions in D2O: product release limits the rate of this sluggish enzyme-catalyzed reaction. J Am Chem Soc. 2012;134(15):6568–70; doi: 10.1021/ja302154k.

48. Savageau MA. Biochemical systems analysis. I. Some mathematical properties of the rate law for the component enzymatic reactions. J Theor Biol. 1969;25(3):365–9.

49. Savageau MA. Biochemical systems analysis. II. The steady-state solutions for an n-pool system using a power-law approximation. J Theor Biol. 1969;25(3):370–9.

50. Voit EO. Modelling metabolic networks using power-laws and S-systems. Essays in biochemistry. 2008;45:29–40; doi: BSE0450029 [pii] 10.1042/BSE0450029.

51. Voit EO, Qi Z, Miller GW. Steps of modeling complex biological systems. Pharmacopsychiatry. 2008;41 Suppl 1:S78–84; doi: 10.1055/s-2008-1080911.

52. Savageau MA. Biochemical systems analysis: a study of function and design in molecular biology. Reading, Mass.: Addison-Wesley Pub. Co., Advanced Book Program; 1976.

53. Petkov SB, Maranas CD. Quantitative assessment of uncertainty in the optimization of metabolic pathways. Biotechnol Bioeng. 1997;56(2):145–61; doi: 10.1002/(SICI)1097-0290(19971020)56:2<145::AID-BIT4>3.0.CO;2-P.

54. Torres NV, Voit EO, Glez-Alcon C, Rodriguez F. An indirect optimization method for biochemical systems: description of method and application to the maximization of the rate of ethanol, glycerol, and carbohydrate production in Saccharomyces cerevisiae. Biotechnol Bioeng. 1997;55(5):758–72; doi: 10.1002/(SICI)1097-0290(19970905)55:5<758::AID-BIT6>3.0.CO;2-A.

55. Torres NV, Voit EO, Gonzalez-Alcon C. Optimization of nonlinear biotechnological processes with linear programming: Application to citric acid production by Aspergillus niger. Biotechnol Bioeng. 1996;49(3):247–58; doi: 10.1002/(SICI)1097-0290(19960205)49:3<247::AID-BIT2>3.0.CO;2-K.

56. Jarboe LR, Liu P, Kautharapu KB, Ingram LO. Optimization of enzyme parameters for fermentative production of biorenewable fuels and chemicals. Comput Struct Biotechnol J. 2012;3:e201210005; doi: 10.5936/csbj.201210005CSBJ-3-e201210005 [pii].

57. Galian C, Manon F, Dezi M, Torres C, Ebel C, Levy D, et al. Optimized purification of a heterodimeric ABC transporter in a highly stable form amenable to 2-D crystallization. PLoS One. 2011;6(5):e19677; doi: 10.1371/journal.pone.0019677PONE-D-10-06141 [pii].

58. Xiao Y, Chu L, Sanakis Y, Liu P. Revisiting the IspH catalytic system in the deoxyxylulose phosphate pathway: achieving high activity. J Am Chem Soc. 2009;131(29):9931–3; doi: 10.1021/ja903778d.

59. Xiao Y, Zahariou G, Sanakis Y, Liu P. IspG enzyme activity in the deoxyxylulose phosphate pathway: roles of the iron-sulfur cluster. Biochemistry. 2009;48(44):10483–5; doi: 10.1021/bi901519q.

60. Brown AC, Eberl M, Crick DC, Jomaa H, Parish T. The nonmevalonate pathway of isoprenoid biosynthesis in Mycobacterium tuberculosis is essential and transcriptionally regulated by Dxs. J Bacteriol. 2010;192(9):2424–33; doi: JB.01402-09 [pii] 10.1128/JB.01402-09.

61. Moussatova A, Kandt C, O’Mara ML, Tieleman DP. ATP-binding cassette transporters in Escherichia coli. Biochim Biophys Acta. 2008;1778(9):1757–71; doi: S0005-2736(08)00194-6 [pii] 10.1016/j.bbamem.2008.06.009.

