## Supplemental: Fig S1, File 1, File 2, and File 3 for "MODEL BASED OPTIMIZACION OF THE TERPENOIDS BIOSYNTHESIS IN *E. coli*"

### Supplementary Figure 1


| | | Pag. |
|---|---|---|
| Figure S1 | Mechanistic fluxes ($\nu_{i,j}$) model of the 4-diphosfocytidyl-2C-methyl-D-erytritol-2-phosphate (MEP) isoprenoids biosynthesis pathway in *Escherichia coli*. ………… | 2 |

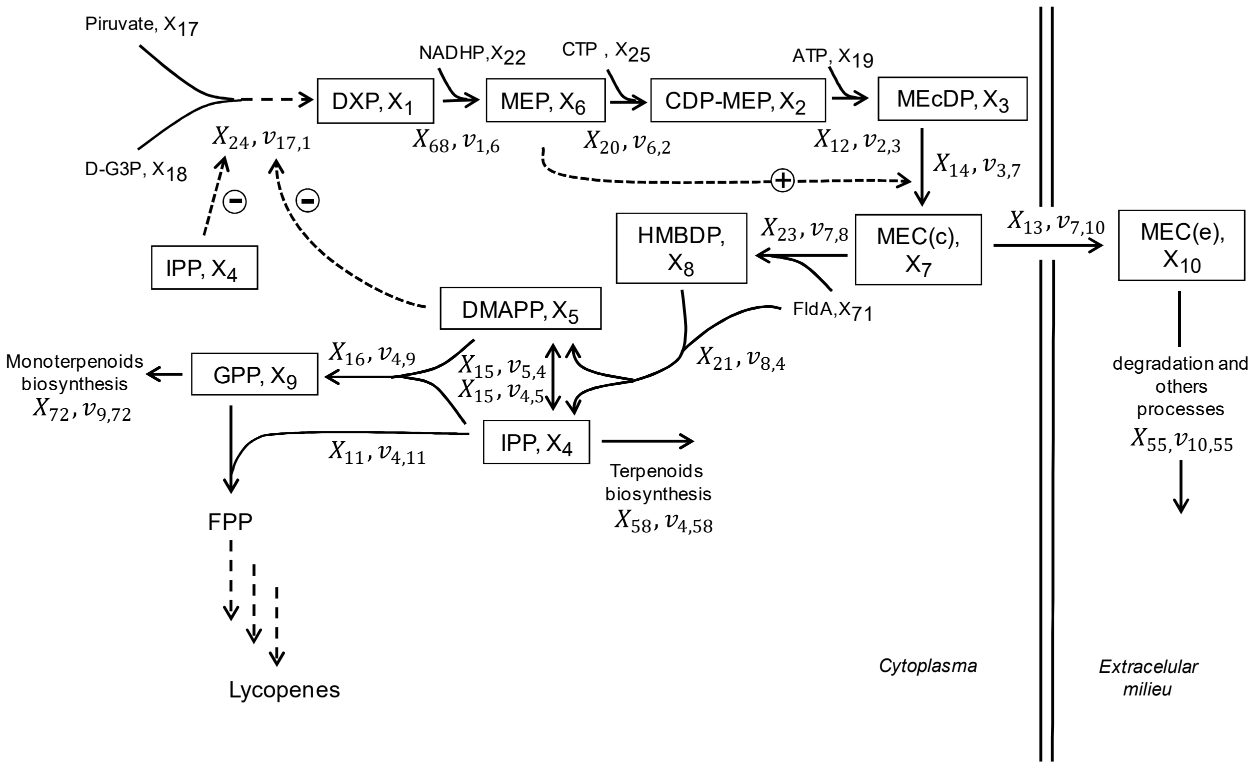


**Figure S1**. **Mechanistic fluxes ($v_{i,j}$) model of the 4-diphosfocytidyl-2C-methyl-D-erytritol-2-phosphate (MEP) isoprenoids biosynthesis pathway in *Escherichia coli.*** Time dependent metabolites: ($X_1$, DXP)1-Deoxy-D-Xylulose 5-Phosphate, ($X_2$, CDP-MEC)4-diphosphocytidyl-2C-methyl D-erythritol, ($X_3$, MEcDP) 4-diphosphocytidyl-2C-methyl D-erythritol 2-phosphate,($X_4$, IPP), Isopentenildiphoosphate, ($X_5$, DMAPP)Dimethylallyl diphosphate, ($X_6$, MEP)2C-methyl-D-erythritol 4-phosphate, ($X_7$, MEC(c))Citosolic 4-difosfocitidil-2C-metil-D-eritritol-2-phosphate, ($X_8$, HMBPP)Hydroxylmethylbutenyl diphosphate, ($X_9$, GPP)Geranyl pyrophosphate, ($X_{10}$, MEC(e))4-difosfocitidyl-2C-metil-D-eritritol-2-fosfato extracellular. Enzymes: ($X_{11}$ and $X_{16}$, IspA) farnesyl diphosphate synthase, ($X_{12}$, IspE) MEcDP kinase, ($X_{13}$) Efflux pump, ($X_{14}$IspF) $MEC_{(c)}$ synthase, ($X_{15}$, IDI) isopentenyldiphosphate isomerase, ($X_{20}$, IspD) CDP-ME synthase, ($X_{21}$, IspH) HMBDP reductase, ($X_{23}$, IspG) HMBDP synthase, ($X_{24}$, DXS) DXP synthase, ($X_{68}$, DXR) DXP reductase, ($X_{55}$). Degradation and biosynthetic processes: ($X_{58}$) Terpenoids biosynthesis, ($X_{72}$) Monoterpenoids biosynthesis. External metabolites: ($X_{17}$) Pyruvate, ($X_{18}$, D-G3P) D-glyceraldehyde 3-phosphate, ($X_{19}$, ATP) Adenosine triphosphate, ($X_{22}$, NADPH) Diphosphopyridindinucleotide phosphate reduced, ($X_{25}$, CTP) Cytidine 5'-triphosphate, ($X_{71}$, FldA) Flavodoxin-1

**Supplementary Files 1**


| | | Pag. |
|---|---|---|
| S1 | Mathematical model equations, structure, parameters, and stability. …………………… | 2 |
| S1.1 | Kinetics model equations. ……………………………………………………….. | 2 |
| S1.2 | Mass balance equations. ………………………………………………………… | 5 |
| S1.3 | Power-law numeric equations. …………………………………………………... | 6 |
| S1.4 | S-system numerical representation. …………………………………...……… | 7 |
| Table S1.1 | Initial and steady states values for the time-dependent variables. ………………. | 8 |
| Table S1.2 | Values for the time-independent parameters. ……….……………………..……….. | 9 |
| Table S1.3 | Eigen values. ……………………………………...…………….………….. | 12 |
| | References. ……………...………………………………………………….……. | 13 |

### S1. Mathematical model equations, structure, parameters, and stability

#### S1.1. *Kinetics equations*

a.- Binding of one inhibitor to one member of a dimer reduces the binding of a second inhibitor molecule to the other member of the dimer [1]

Kinetics from ([2], case VI)

$$G_e = 1 + \frac{X_4}{K_{i_{4,24}}} + \frac{X_5}{K_{i_{5,24}}}$$
$$v_{0,1} = \frac{V_{max_{24}} \cdot X_{17} \cdot X_{18}}{K_{m_{17,24}} \cdot X_{18} + \left(K_{d_{17,24}} \cdot K_{m_{18,24}} + X_{17} \cdot K_{m_{18,24}}\right) \cdot G_e + X_{17} \cdot X_{18}} \quad \text{Eqs. S1}$$

b.- For the fluxes: $v_{1,6}, v_{7,8}, v_{8,4}$ was used the random sequential mechanism kinetics from [2] with initial binding for NADPH [3], FldA, or GPP accordingly.

$$v_{1,6} = \frac{V_{max_{68}} \cdot X_1 \cdot X_{22}}{K_{d_{22,68}} \cdot K_{m_{1,68}} + K_{m_{1,68}} \cdot X_{22} + K_{m_{22,68}} \cdot X_1 + X_1 \cdot X_{22}} \quad \text{Eq. S3}$$

$$v_{7,8} = \frac{V_{max_{23}} \cdot X_7 \cdot X_{71}}{K_{d_{71,23}} \cdot K_{m_{7,23}} + K_{m_{7,23}} \cdot X_{71} + K_{m_{71,23}} \cdot X_7 + X_7 \cdot X_{71}} \quad \text{Eq. S4}$$

$$v_{8,4} = \frac{V_{max_{21}} \cdot X_8 \cdot X_{71}}{K_{d_{71,21}} \cdot K_{m_{8,21}} + K_{m_{8,21}} \cdot X_{71} + K_{m_{71,21}} \cdot X_8 + X_8 \cdot X_{71}} \quad \text{Eq. S5}$$

$$v_{4,11} = \frac{V_{max_{11}} \cdot X_4 \cdot X_9}{K_{d_{9,11}} \cdot K_{m_{4,11}} + K_{m_{9,11}} \cdot X_4 + K_{m_{4,11}} \cdot X_9 + X_4 \cdot X_9} \quad \text{Eq. S6}$$

c.- Equation bisubstrate. For the fluxes: $v_{4,9}$ ([4], Eq. 1), $v_{2,3}$ [5], and $v_{6,2}$

$$v_{4,9} = \frac{V_{max_{16}} \cdot X_4 \cdot X_5}{K_{m_{4,16}} \cdot X_5 + K_{m_{5,16}} \cdot X_4 + X_4 \cdot X_5} \qquad \text{Eq. S7}$$

$$v_{2,3} = \frac{V_{max_{12}} \cdot X_2 \cdot X_{19}}{K_{m_{19,12}} \cdot X_2 + K_{m_{2,12}} \cdot X_{19} + X_2 \cdot X_{19}} \qquad \text{Eq. S8}$$

$$v_{6,2} = \frac{V_{max_{20}} \cdot X_6 \cdot X_{25}}{K_{m_{6,20}} \cdot X_{25} + K_{m_{25,20}} \cdot X_6 + X_6 \cdot X_{25}} \qquad \text{Eq. S9}$$

d.- Flux $v_{4,5}$ described with a reversible kinetics ([6], Eq. 9)

$$v_{4,5} = \frac{V_{max_{15}} \cdot \left( \frac{X_4}{K_{m_{4,15}} - R_{4,5} \cdot \frac{X_5}{K_{m_{5,15}}}} \right)}{1 + \frac{X_4}{K_{m_{4,15}} + \frac{X_5}{K_{m_{5,15}}}}} \qquad \text{Eq. S10}$$

Where $R_{4\_5} = 0.9$ represent the IDI flux ratio in direction DMAPP ($X_5$) formation [7]. It is reported for the *in vitro* IDI a predominant formation of DMAPP (66%) with respect IPP (34%) at the equilibrium [8].

e.- Flux $v_{5,4}$ described with a reversible kinetics ([6], Eq. 9)

$$v_{5,4} = \frac{V_{max_{15}} \cdot \left( \frac{X_5}{K_{m_{5,15}} - R_{5,4} \cdot \frac{X_4}{K_{m_{4,15}}}} \right)}{1 + \frac{X_4}{K_{m_{4,15}} + \frac{X_5}{K_{m_{5,15}}}}} \qquad \text{Eq. S11}$$

Where $R_{5\_4} = 1 - R_{4\_5} = 0.1$ represent the IDI flux ratio in direction IPP ($X_4$) formation [7].

f.- Flux $v_{67,10}$ for MECc extrusion

$$v_{7,10} = \frac{V_{max_{13}} \cdot X_7^{hill_{7,13}}}{K_{m_{7,13}} + X_7^{hill_{7,13}}} \qquad \text{Eq. S12}$$

The effluxes $v_{4,58}$, $v_{9,72}$, and $v_{10,55}$ were described by mass actions first order rates as:

$$v_{4,58} = K_{4,58} \cdot X_4 \qquad \text{Eq. S13}$$

$$v_{9,72} = K_{9,72} \cdot X_9 \qquad \text{Eq. S14}$$

$$v_{10,55} = K_{10,55} \cdot X_{10} \qquad \text{Eq. S15}$$

g.- The $v_{3,7}$ flux is drive by IspF ($X_{14}$) enzyme. This enzyme report a non-essential activation kinetics by MEP ($X_6$) [9]. The non-essential activation kinetics equation was obtained from [10].

$$v_{3,7} = \frac{V_{max_{14}} \cdot \sigma_6 \cdot \left(\frac{1 + \beta_6 \cdot X_6}{\alpha_6 \cdot K_{a_{6.14}}}\right)}{\left(1 + \frac{X_6}{K_{a_{6.14}}} + \sigma_6 \cdot \left(1 + \left(\frac{X_6}{\alpha_6 \cdot K_{a_{6.14}}}\right)\right)\right)}$$

with:

$$v_6 = \frac{X_3}{K_{m_{3,14}}}$$
$$\alpha_6 = 1;\ 1 \le \alpha_6 \le \infty$$
$$\beta_6 = 2000;\ \beta_6 \ge 1 \qquad \text{Eq. S16}$$

With the $\alpha_6$ and $\beta_6$ parameters estimated as 1 and 2000 respectively in base of the time dependent kinetics behavior (not shown) as no data is currently available to our knowledge.

### S1.2. - *Mass balance equations.*

There are ten flux nodes in the MEP pathway (Figure 1). GMA representation as $v_{i,j}$ for $i,j = \{1..10\}$. At upper right is represented the S-system input and output fluxes aggregated by nodes as $V_i^+$ and $V_i^-$ for $i = \{1..10\}$.

$$\frac{dX_1}{dt} = v_{0,1} - v_{1,6} = V_1^+ - V_1^-$$
$$\frac{dX_2}{dt} = v_{6,2} - v_{2,3} = V_2^+ - V_2^-$$
$$\frac{dX_3}{dt} = v_{2,3} - v_{3,7} = V_3^+ - V_3^-$$
$$\frac{dX_4}{dt} = (v_{8,4} + v_{5,4}) - (v_{4,5} + v_{4,9} + v_{4,11} + v_{4,58}) = V_4^+ - V_4^-$$
$$\frac{dX_5}{dt} = (v_{8,4} + v_{4,5}) - (v_{5,4} + v_{4,9}) = V_5^+ - V_5^-$$
$$\frac{dX_6}{dt} = v_{1,6} - v_{6,2} = V_6^+ - V_6^-$$
$$\frac{dX_7}{dt} = v_{3,7} - (v_{7,8} + v_{7,10}) = V_7^+ - V_7^-$$
$$\frac{dX_8}{dt} = v_{7,8} - v_{8,4} = V_8^+ - V_8^-$$
$$\frac{dX_9}{dt} = v_{4,9} - (v_{4,11} + v_{9,72}) = V_9^+ - V_9^-$$
$$\frac{dX_{10}}{dt} = v_{7,10} - v_{10,53} = V_{10}^+ - V_{10}^-$$

Eqs. S17

#### S1.3.- *Power-law numeric equations*

$$v_{0,1} = 0{,}4204659 \cdot X_4^{-0{,}002963428} \cdot X_5^{-0{,}002984825} \cdot X_{17}^{0{,}03191547} \cdot X_{18}^{0{,}4947393} \cdot X_{24} \cdot X_{36} \cdot X_{48}^{-0{,}02414584} \cdot X_{49}^{-0{,}4947393} \cdot X_{57}^{0{,}002963428} \cdot X_{59}^{0{,}002984825} X_{64}^{-0{,}007769629}$$

$$v_{1,6} = 0{,}7969211 \cdot X_1^{0{,}9469133} \cdot X_{22}^{0{,}002843305} \cdot X_{27} \cdot X_{38}^{-0{,}9469133} \cdot X_{53}^{-0{,}000220277} \cdot X_{65}^{-0{,}002623028} \cdot X_{68}$$

$$v_{2,3} = 0{,}9158758 \cdot X_2^{0{,}9840265} \cdot X_{12} \cdot X_{19}^{0{,}001961656} \cdot X_{28} \cdot X_{39}^{-0{,}9840265} \cdot X_{50}^{-0{,}001961656}$$

$$v_{3,7} = 0{,}894294 \cdot X_3^{0{,}9954424} \cdot X_6^{0{,}9510551} \cdot X_{14} \cdot X_{30} \cdot X_{40}^{-0{,}9954424} \cdot X_{61}^{-0{,}9864621} \cdot X_{63}^{0{,}9866242} \cdot X_{69}^{-0{,}9510551}$$

$$v_{4,5} = 0{,}2412812 \cdot X_4^{2{,}17969} \cdot X_5^{-1{,}185707} \cdot X_{15} \cdot X_{31} \cdot X_{41}^{-2{,}17969} \cdot X_{42}^{1{,}185707}$$

$$v_{4,9} = 0{,}7526662 \cdot X_4^{0{,}05179843} \cdot X_5^{0{,}9324028} \cdot X_{16} \cdot X_{32} \cdot X_{46}^{-0{,}05179843} \cdot X_{47}^{-0{,}9324028}$$

$$v_{4,11} = 0{,}9457848 \cdot X_4^{0{,}9968513} \cdot X_9^{0{,}9943671} \cdot X_{11} \cdot X_{26} \cdot X_{37}^{-0{,}00293993} \cdot X_{56}^{-0{,}9968513} \cdot X_{62}^{-0{,}9914272}$$

$$v_{4,58} = X_4 \cdot X_{58}$$

$$v_{5,4} = 0{,}6648911 \cdot X_4^{-0{,}1754202} \cdot X_5^{1{,}104395} \cdot X_{15} \cdot X_{31} \cdot X_{41}^{0{,}1754202} \cdot X_{42}^{-1{,}104395}$$

$$v_{6,2} = 0{,}8026983 \cdot X_6^{0{,}9514521} \cdot X_{20} \cdot X_{25}^{0{,}01066371} \cdot X_{33} \cdot X_{43}^{-0{,}9514521} \cdot X_{54}^{-0{,}01066371}$$

$$v_{7,8} = 0{,}6374959 \cdot X_7^{0{,}9932275} \cdot X_{23} \cdot X_{35} \cdot X_{44}^{-0{,}9932275} \cdot X_{52}^{-0{,}0054125} \cdot X_{66}^{-0{,}1412736} \cdot X_{71}^{0{,}1466861}$$

$$v_{7,10} = 0{,}6519242 \cdot X_7^{0{,}5722953} \cdot X_{13} \cdot X_{29} \cdot X_{60}^{-0{,}8470297} \cdot X_{70}^{-5{,}94603}$$

$$v_{8,4} = 0{,}6419102 \cdot X_8^{0{,}994558} \cdot X_{21} \cdot X_{34} \cdot X_{45}^{-0{,}994558} \cdot X_{51}^{-0{,}004349214} \cdot X_{67}^{-0{,}1414628} \cdot X_{71}^{0{,}1458121}$$

$$v_{9,72} = X_9 \cdot X_{72}$$

$$v_{10,55} = X_{10} \cdot X_{55}$$

Eqs. S18

### S1.4.- *S-system numeric representation*

$$\frac{dX_1}{dt} = 0{,}4204659 \cdot X_4^{-0{,}002963428} \cdot X_5^{-0{,}002984825} \cdot X_{17}^{0{,}03191547} \cdot X_{18}^{0{,}4947393} \cdot X_{24} \cdot X_{36} \cdot X_{48}^{-0{,}02414584} \cdot X_{49}^{-0{,}4947393} \cdot X_{57}^{0{,}002963428} \cdot X_{59}^{0{,}002984825} X_{64}^{-0{,}007769629} - 0{,}7969211 \cdot X_1^{0{,}9469133} \cdot X_{22}^{0{,}002843305} \cdot X_{27} \cdot X_{38}^{-0{,}9469133} \cdot X_{53}^{-0{,}000220277} \cdot X_{65}^{-0{,}002623028} \cdot X_{68}$$

$$\frac{dX_2}{dt} = 0{,}8026983 \cdot X_6^{0{,}9514521} \cdot X_{20} \cdot X_{25}^{0{,}01066371} \cdot X_{33} \cdot X_{43}^{-0{,}9514521} \cdot X_{54}^{-0{,}01066371} - 0{,}9158758 \cdot X_2^{0{,}9840265} \cdot X_{12} \cdot X_{19}^{0{,}001961656} \cdot X_{28} \cdot X_{39}^{-0{,}9840265} \cdot X_{50}^{-0{,}001961656}$$

$$\frac{dX_3}{dt} = 0{,}9158758 \cdot X_2^{0{,}9840265} \cdot X_{12} \cdot X_{19}^{0{,}001961656} \cdot X_{28} \cdot X_{39}^{-0{,}9840265} \cdot X_{50}^{-0{,}001961656} - 0{,}894294 \cdot X_3^{0{,}9954424} \cdot X_6^{0{,}9510551} \cdot X_{14} \cdot X_{30} \cdot X_{40}^{-0{,}9954424} \cdot X_{61}^{-0{,}9864621} \cdot X_{63}^{0{,}9866242} \cdot X_{69}^{-0{,}9510551}$$

$$\frac{dX_4}{dt} = 1{,}232352 \cdot X_4^{-0{,}05945175} \cdot X_5^{0{,}3742912} \cdot X_8^{0{,}6574918} \cdot X_{15}^{0{,}3389105} \cdot X_{21}^{0{,}6610895} \cdot X_{31}^{0{,}3389105} \cdot X_{34}^{0{,}6610895} \cdot X_{41}^{0{,}05945175} \cdot X_{42}^{-0{,}3742912} \cdot X_{45}^{-0{,}6574918} \cdot X_{51}^{-0{,}00287522} \cdot X_{67}^{-0{,}0935196} \cdot X_{71}^{0{,}09639482} - 1{,}905929 \cdot X_4^{0{,}9262724} \cdot X_5^{0{,}05848977} \cdot X_9^{0{,}01279603} \cdot X_{11}^{0{,}01286852} \cdot X_{15}^{0{,}1164123} \cdot X_{16}^{0{,}4385913} \cdot X_{26}^{0{,}01286852} \cdot X_{31}^{0{,}1164123} \cdot X_{32}^{0{,}4385913} \cdot X_{37}^{--3{,}783254E-5} \cdot X_{41}^{-0{,}4585981} \cdot X_{42}^{0{,}350454} X_{46}^{-0{,}02271834} \cdot X_{47}^{-0{,}4089438} \cdot X_{56}^{-0{,}012828} \cdot X_{58}^{0{,}4321279} \cdot X_{62}^{-0{,}0127582}$$

$$\frac{dX_5}{dt} = 0{,}7488188 \cdot X_4^{0{,}5898355} \cdot X_5^{-0{,}4507437} \cdot X_8^{0{,}8456467} \cdot X_{15}^{0{,}1497261} \cdot X_{21}^{0{,}8502739} \cdot X_{31}^{0{,}1497261} \cdot X_{34}^{0{,}8502739} \cdot X_{41}^{-0{,}5898355} \cdot X_{42}^{0{,}4507437} \cdot X_{45}^{-0{,}8456467} \cdot X_{51}^{-0{,}003698024} \cdot X_{67}^{-0{,}1202822} \cdot X_{71}^{0{,}1239802} - 1{,}414424 \cdot X_4^{-0{,}04724543} \cdot X_5^{1{,}007374} \cdot X_{15}^{0{,}4358967} \cdot X_{16}^{0{,}5641033} \cdot X_{31}^{0{,}4358967} \cdot X_{32}^{0{,}5641033} \cdot X_{41}^{0{,}0764651} \cdot X_{42}^{-0{,}4814023} X_{46}^{-0{,}02921967} \cdot X_{47}^{-0{,}5259715}$$

$$\frac{dX_6}{dt} = 0{,}7969211 \cdot X_1^{0{,}9469133} \cdot X_{22}^{0{,}002843305} \cdot X_{27} \cdot X_{38}^{-0{,}9469133} \cdot X_{53}^{-0{,}000220277} \cdot X_{65}^{-0{,}002623028} \cdot X_{68} - 0{,}8026983 \cdot X_6^{0{,}9514521} \cdot X_{20} \cdot X_{25}^{0{,}01066371} \cdot X_{33} \cdot X_{43}^{-0{,}9514521} \cdot X_{54}^{-0{,}01066371}$$

$$\frac{dX_7}{dt} = 0{,}894294 \cdot X_3^{0{,}9954424} \cdot X_6^{0{,}9510551} \cdot X_{14} \cdot X_{30} \cdot X_{40}^{-0{,}9954424} \cdot X_{61}^{-0{,}9864621} \cdot X_{63}^{0{,}9866242} \cdot X_{69}^{-0{,}9510551} - 0{,}6623173 \cdot X_7^{0{,}8473563} \cdot X_{13}^{0{,}9977662} \cdot X_{23}^{0{,}002233782} \cdot X_{29}^{0{,}9977662} \cdot X_{35}^{0{,}002233782} \cdot X_{44}^{-0{,}002218653} \cdot X_{52}^{-1{,}209034E-5} \cdot X_{60}^{-0{,}8451376} \cdot X_{66}^{-0{,}0003155744} \cdot X_{70}^{-5{,}932748} \cdot X_{71}^{0{,}0003276647}$$

$$\frac{dX_8}{dt} = 0{,}6374959 \cdot X_7^{0{,}9932275} \cdot X_{23} \cdot X_{35} \cdot X_{44}^{-0{,}9932275} \cdot X_{52}^{-0{,}0054125} \cdot X_{66}^{-0{,}1412736} \cdot X_{71}^{0{,}1466861} - 0{,}6419102 \cdot X_8^{0{,}994558} \cdot X_{21} \cdot X_{34} \cdot X_{45}^{-0{,}994558} \cdot X_{51}^{-0{,}004349214} \cdot X_{67}^{-0{,}1414628} \cdot X_{71}^{0{,}1458121}$$

$$\frac{dX_9}{dt} = 0{,}7526662 \cdot X_4^{0{,}05179843} \cdot X_5^{0{,}9324028} \cdot X_{16} \cdot X_{32} \cdot X_{46}^{-0{,}05179843} \cdot X_{47}^{-0{,}9324028} - 1{,}139748 \cdot X_4^{0{,}02924818} \cdot X_9^{0{,}9998347} \cdot X_{11}^{0{,}02934057} \cdot X_{26}^{0{,}02934057} \cdot X_{37}^{-8{,}625921E-5} \cdot X_{56}^{-0{,}02924818} \cdot X_{62}^{-0{,}02908904} \cdot X_{72}^{0{,}9706594}$$

$$\frac{dX_{10}}{dt} = 0{,}6519242 \cdot X_7^{0{,}5722953} \cdot X_{13} \cdot X_{29} \cdot X_{60}^{-0{,}8470297} \cdot X_{70}^{-5{,}94603} - X_{10} \cdot X_{55}$$

Eqs. S19

**Table S1.1**. Initial and steady states values for the model time-dependent variables. The steady-state values were the used at the initial values in the simulation and optimization runs.

| | | ODE's initial values (mM) | References | ODE steady-state values (mM) |
|---|---|---|---|---|
| 1-Deoxy-D-Xylulose 5-Phosphate (DXP) | $X_1$ | 7e-3 | 0.01-2 μM. [11] (*) | 2.519342e-3 |
| 4-diphosphocytidyl-2C-methyl D-erythritol (CDP-MEC) | $X_2$ | 1e-4 | 0.01-0.5 μM. [11] | 2.135892e-3 |
| 4-diphosphocytidyl-2C-methyl D-erythritol 2-phosphate (MEcDP) | $X_3$ | 5e-4 | 0.1-1 μM. [12] | 1.552085e-3 |
| Isopentenildiphoosphate (IPP), | $X_4$ | 2.5e-5 | 0.025 μM. [13] | 3.965051e-4 |
| Dimethylallyl diphosphate, (DMAPP) | $X_5$ | 2.5e-5 | [14] | 4.964621e-4 |
| 2C-methyl-D-erythritol 4-phosphate (MEP) | $X_6$ | 1e-5 | 0.01-2 μM. [11] | 1.250263e-4 |
| Citosolic 4-difosfocitidil-2C-metil-D-eritritol-2-phosphate (MEC(c)) | $X_7$ | 3e-4 | 0.01-2 μM. [11] | 8.939506e-4 |
| Hydroxylmethylbutenyl diphosphate (HMBPP) | $X_8$ | 5e-5 | 0.01-2 μM. [11] | 3.843069e-5 |
| Geranyl pyrophosphate (GPP) | $X_9$ | 1.5e-3 | ([15], Fig 3B) | 3.906279e-4 |
| 4-difosfocitidyl-2C-metil-D-eritritol-2-fosfato extracelular (MEC(e)) | $X_{10}$ | 0.03 | 37-42 μM. [11] | 0.02709473 |

(*) The data from [11] were obtained from *E. coli* strains genetically engineering by over expressing enzymes of the MEC pathway: DXS ($X_{24}$), IspD ($X_{20}$), IspF ($X_{14}$), and IDI ($X_{15}$)

**Table S1.2.** Values for the time-independent parameters.

| **Enzymes** | | **mM** | **References and comments** |
|---|---|---|---|
| farnesyl diphosphate synthase (IspA) | $X_{11}$ | 1.632863e-4 | 354 molecules/cell ([16], P22939, Table S2)(1) |
| MEcDP kinase (IspE, MCS) | $X_{12}$ | 9.594226e-5 | 208±22 molecules/cell, ([17], Table 1) |
| Efflux pump | $X_{13}$ | 1.476035e-5 | 32 molecules/cell for antiporter *fsr*, ([16], P52067, Table S2) |
| $MEC_{(c)}$ synthase (IspF, HDS) | $X_{14}$ | 5.119995e-5 | 111±13 molecules/cell([17],Table 1) |
| isopentenyl diphosphate isomerase (IDI) | $X_{15}$ | 2.767565e-6 | 6 molecules/cell ([17], Table 1. < 10 molecules/cell) |
| farnesyl diphosphate synthase (IspA) | $X_{16}$ | 1.632863e-4 | 354 molecules/cell ([16], P22939, Table S2) |
| CDP-ME synthase (IspD, CMK) | $X_{20}$ | 2.306304e-5 | 50±8 molecules/cell([17], Table 1) |
| HMBDP reductase (IspH) | $X_{21}$ | 1.005549e-4 | 218±22 molecules/cell ([17], Table 1) |
| HMBDP synthase (IspG) | $X_{23}$ | 9.409721e-5 | 204±25 molecules/cell ([17], Table 1) |
| DXP synthase (DXS) | $X_{24}$ | 2.813691e-5 | 64±4 molecules/cell ([17], Table 1) |
| DXP reductase (DXR) | $X_{68}$ | 1.476035e-5 | 32±5 molecules/cell ([17], Table 1) |

| **External model metabolites** | | **mM** | |
|---|---|---|---|
| Pyruvate | $X_{17}$ | **2.67** | ([18], Table V) |
| | | 7.5 | [19]. |
| D-glyceraldehyde 3-phosphate (D-G3P) | $X_{18}$ | **0.041** | Assumed equal at the Km of DXS as a clue of the physiological level, see [20] |
| | | 0.218 | [18], Table V) for G3P |
| | | 1.1 | [21], Table 1) for G3P |
| Adenosine triphosphate (ATP) | $X_{19}$ | **3** | ([22], Table 3) |
| | | 1.5 | [23], [24] |
| Diphosphopyridindinucleotide phosphate, reduced (NADPH) | $X_{22}$ | **0.12** | ([25], Table 1) |
| | | 1.25 | [26] |
| Cytidine 5'-triphosphate (CTP) | $X_{25}$ | **2.7** | ([25], Table 1) |
| | | 0.515 | ([22], Table 3) |
| Flavodoxin-1(FldA) | $X_{71}$ | 3.0153e-3 | 6537 molecules/cell ([16], P61949, Table S2) |

| $\mathbf{k_{cat's}}$ | | $\mathbf{min^{-1}}$ | |
|---|---|---|---|
| kcat_11 | $X_{26}$ | 1.73 | Obtained in base the $V_{max's}$ relationship for GPP and FPP enzymatic IspA products ([15], Table 2). As $X_{11}$ and $X_{16}$ correspond at the same IspA: kcat_11= $V_{max11}/V_{max16} \times$ kcat_16= 23.8/10.7×0.78 $min^{-1}$= 1.73 $min^{-1}$ |
| kcat_12 | $X_{28}$ | **1010** | ([17], Table 1) data based in [27] |
| | | 24600 | [352.7 $s^{-1}$, 487 $s^{-1}$] ([28], Table 1) for different species (not included *E. coli*) |
| kcat_13 | $X_{29}$ | **600** | 10 $s^{-1}$ for Nitrocefin [29] |
| | | 282 | [30] ABC transporter |

| | | | |
|---|---|---|---|
| | | 9 | [31] *E. coli* Penicillin-Binding-Protein-5 |
| kcat_14 | $X_{30}$ | **80.7** | ([9], Table 1 with MEP activation) |
| | | 162 | ([32], Table 1) |
| | | 60.6 | ([9], Table 1) |
| kcat_15 | $X_{31}$ | 19.8 | [33] |
| kcat_16 | $X_{32}$ | **0.78** | 0.013 ± 0.001 $s^{-1}$ [34]. Undecaprenyl pyrophosphate synthase (UPPS) in absence of Triton (3) |
| | | 150 | 2.5 ± 0.1 $s^{-1}$[34]. Undecaprenyl pyrophosphate synthase (UPPS) in presence of Triton |
| | | 96 | [35] |
| kcat_20 | $X_{33}$ | **1554** | ([36], Table 1) |
| | | 1686 | 25.9 ±5.9 $s^{-1}$ ([36], Table 1) |
| | | 1008 | [37] |
| kcat_21 | $X_{34}$ | **27.6** | 0.46 $s^{-1}$ ([38] , Table 1. NADH as agent redox) |
| | | 1125 | [39], using MDQ) |
| kcat_23 | $X_{35}$ | 23.7 | [40] |
| kcat_24 | $X_{36}$ | **100.3** | [41] |
| | | 153.6 | 153.6 ± 6.8 $min^{-1}$[42] |
| kcat_68 | $X_{27}$ | **1740** | 29 ± 1 s-1 [43], Table 1) |
| | | 6960 | 116 ± 8 s-1 [3] |

| **Km, $K_{0.5}$ , and Ki** | | **mM** | |
|---|---|---|---|
| Km_1_68 | $X_{38}$ | **0.045** | [43], Table 1) |
| | | 0.115 | [3] |
| Km_2_12 | $X_{39}$ | 0.15 | ([27], Fig. 5) |
| Km_3_14 | $X_{40}$ | **0.339** | ([9], Table 1, no activation by MEP) |
| | | 0.0938 | ([9], Table 1, with MEP activation) |
| | | 0.41 | ([32], Table 1) |
| Km_4_15 | $X_{41}$ | 0.0095 | [33; 44] |
| Km_5_15 | $X_{42}$ | 0.0143 | [33] |
| Km_6_20 | $X_{43}$ | **0.00314** | [45] |
| | | 0.2 | [46] |
| Km_7_23 | $X_{44}$ | **0.56** | ([47], Table 3) |
| | | 0.311 | [40] |
| Km_8_21 | $X_{45}$ | 0.03 | ([39], Table 3) |
| Km_4_11 | $X_{56}$ | 0.0103 | ([15], Table 2) |
| Km_9_11 | $X_{37}$ | 0.0055 | ([15], Table 2) |
| Km_4_16 | $X_{46}$ | **0.0013** | ([15], Table 2) |
| | | 0.0041 | [34] |
| | | 0.00013 | [35] |
| Km_5_16 | $X_{47}$ | **0.0293** | ([15], Table 2) |
| | | 0.0005 | ([35], avian) |
| Km_17_24 | $X_{48}$ | **0.134** | ([41], Table 2) |
| | | 0.0487 | [42] |
| Km_18_24 | $X_{49}$ | **0.041** | ([41], Table 2) |
| | | 0.0235 | [42] |
| Km_19_12 | $X_{50}$ | 0.42 | ([27], Fig. 5) |
| Km_71_21 | $X_{51}$ | 0.012 | [48] |
| Km_71_23 | $X_{52}$ | 0.012 | [48] |
| Km_22_68 | $X_{53}$ | 0.0005 | [3; 43] |
| Km_25_20 | $X_{54}$ | **0.76** | ([36], Table 1) |

| | | | |
|---|---|---|---|
| | | 0.131 | [45] |
| K_10_55 | $X_{55}$ | 0.05 mM/min | Estimated flux of degradation and /or transformation |
| K_4_58 | $X_{58}$ | 0.005 mM/min | Estimated flux of biosynthesis of diverse monoterpenoids |
| K_9_72 | $X_{72}$ | 5e-3 mM/min | Estimated flux of biosynthesis of diverse terpenoids |
| hill_7_13 | $X_{70}$ | 1 | [49], Nitrocefin with no positive cooperativity |
| Ki_4_24 | $X_{57}$ | 0.0654 | [1]from *Populustrichocarpa* |
| Ki_5_24 | $X_{59}$ | 0.0813 | [1]from *Populustrichocarpa* |
| Km_7_13 | $X_{60}$ | **0.00495** | 4.95±1.14 μM [29] for Nitrocefin |
| | | 8.8e-4 | [49], Table 2) for Ampicillin |
| alpha_6_14 | $X_{61}$ | 1 | ([50], Eq. 2.8). Estimated inside the α ranges for non-essential activation |
| Kd_9_11 | $X_{62}$ | 0.0714 | [15] |
| betha_6_14 | $X_{63}$ | 2000 | ([50], Eq. 2.8). Estimated inside the β ranges for non-essential activation |
| Kd_17_24 | $X_{64}$ | 0.0426 | [42], Fig 3A) |
| Kd_22_68 | $X_{65}$ | 0.00033 | Estimated as 2/3×Km_22_68 (2) |
| Kd_71_23 | $X_{66}$ | 0.0005 | [48] |
| Kd_71_21 | $X_{67}$ | 0.0005 | [48] |
| Kd_22_23 | $X_{68}$ | 1.476e-5 | Estimated as 2/3×Km_22_23 (2) |

(*) Adimensional value

(1) The concentrations, in mM, were calculated as: $[E] = \frac{molecules/cell}{N^{\circ}_{avogadro} \cdot V_{cell}} \times 1000$

with the number of Avogadro $6.022140857 \times 10^{23}$molecules⁄mol and the *E. coli* volume $3.43e^{-15}L$ ([51], $3.43 \pm 1.2\ fL$ from Table 1 for complex medium)

(2) Estimated as Kd < Km because Kd = $K_{bak}/K_{for}$ and Km = $(K_{bak}+Kcat)/K_{for} \Rightarrow$ Kd ≅ Km-Kcat/$K_{for}$

(3) To resemble the cell *in vivo* conditions it was decided to use the IspA for $k_{cat\_16}$ value in absence of triton 0.013 $s^{-1}$ a far lower than the 150 $s^{-1}$ in presence of triton [34]. The $k_{cat\_11}$ was obtained in base the IspA products $V_{max}$ relationship for GPP and FPP ([15], Table 2)

When more than a value is indicated for a parameter then the value in bold is the implemented

**Table S1.3. Eigen values of the S-system representation for the model at *steady* state.** All the eigenvalues have only negative real parts indicating that the nominal state is locally stable and that the system will return to its original state after small perturbations. No imaginary parts present which indicate the absence of oscillations.

| Real parts | Im |
|---|---|
| - 0,05 | 0 |
| -10,33265 | 0 |
| -1,286988 | 0 |
| -0,8708976 | 0 |
| -0,6253112 | 0 |
| -0,5105081 | 0 |
| -0,07841364 | 0 |
| -0,006611018 | 0 |
| -0,005379767 | 0 |
| -0,0150956 | 0 |

### References

[1] A. Banerjee, Y. Wu, R. Banerjee, Y. Li, H. Yan, and T.D. Sharkey, Feedback inhibition of deoxy-D-xylulose-5-phosphate synthase regulates the methylerythritol 4-phosphate pathway. J Biol Chem 288 (2013) 16926-36.

[2] Y. Cheng, and W.H. Prusoff, Relationship between the inhibition constant (K1) and the concentration of inhibitor which causes 50 per cent inhibition (I50) of an enzymatic reaction. Biochem Pharmacol 22 (1973) 3099-108.

[3] A.T. Koppisch, D.T. Fox, B.S. Blagg, and C.D. Poulter, *E. coli* MEP synthase: steady-state kinetic analysis and substrate binding. Biochemistry 41 (2002) 236-43.

[4] Y. Matsue, H. Mizuno, T. Tomita, T. Asami, M. Nishiyama, and T. Kuzuyama, The herbicide ketoclomazone inhibits 1-deoxy-D-xylulose 5-phosphate synthase in the 2-C-methyl-D-erythritol 4-phosphate pathway and shows antibacterial activity against *Haemophilus influenzae*. J Antibiot (Tokyo) 63 (2010) 583-8.

[5] V.K. Singh, and I. Ghosh, Methylerythritol phosphate pathway to isoprenoids: kinetic modeling and in silico enzyme inhibitions in *Plasmodium falciparum*. FEBS Lett 587 (2013) 2806-17.

[6] A.K. Groen, R. van der Meer, H.V. Westerhoff, R.J.A. Wanders, T.P.M. Akerboom, and J.M. Tagerg, In: Metabolic compartmentation, Academic Press, London ; New York, 1982.

[7] T. Grawert, I. Span, W. Eisenreich, F. Rohdich, J. Eppinger, A. Bacher, and M. Groll, Probing the reaction mechanism of IspH protein by x-ray structure analysis. Proc Natl Acad Sci U S A 107 (2010) 1077-81.

[8] V. Jonnalagadda, K. Toth, and J.P. Richard, Isopentenyl diphosphate isomerase catalyzed reactions in D2O: product release limits the rate of this sluggish enzyme-catalyzed reaction. J Am Chem Soc 134 (2012) 6568-70.

[9] J.K. Bitok, and C.F. Meyers, 2C-Methyl-d-erythritol 4-phosphate enhances and sustains cyclodiphosphate synthase IspF activity. ACS Chem Biol 7 (2012) 1702-10.

[10] K. Brix, and W. Stöcker, Proteases : structure and function.

Ch. 2 - Kinetics of the Interaction of Peptidases with Substrates and Modifiers

Antonio Baici, Marko Novinec and Brigita Lenar*ci*c.

[11] K. Zhou, R. Zou, G. Stephanopoulos, and H.P. Too, Metabolite profiling identified methylerythritol cyclodiphosphate efflux as a limiting step in microbial isoprenoid production. PLoS One 7 (2012) e47513.

[12] K. Zhou, R. Zou, G. Stephanopoulos, and H.P. Too, Enhancing solubility of deoxyxylulose phosphate pathway enzymes for microbial isoprenoid production. Microb Cell Fact 11 (2012) 148.

[13] J.B. Gracia, Regulación de la ruta del 1-deoxi-d-xylulose-5-fosfato (DXP) en *Escherichia coli* recombinante para la producción de licopeno. Facultad de Quimica. Departamento de Bioquímica y Biología Molecular (B) e Inmunología. Máster en Química Fina y Molecular (2014) 54.

[14] V.J. Martin, D.J. Pitera, S.T. Withers, J.D. Newman, and J.D. Keasling, Engineering a mevalonate pathway in *Escherichia coli* for production of terpenoids. Nat Biotechnol 21 (2003) 796-802.

[15] B. Ku, J.C. Jeong, B.N. Mijts, C. Schmidt-Dannert, and J.S. Dordick, Preparation, characterization, and optimization of an in vitro C30 carotenoid pathway. Appl Environ Microbiol 71 (2005) 6578-83.

[16] B. Soufi, K. Krug, A. Harst, and B. Macek, Characterization of the *E. coli* proteome and its modifications during growth and ethanol stress. Front Microbiol 6 (2015) 103.

[17] D.C. Volke, J. Rohwer, R. Fischer, and S. Jennewein, Investigation of the methylerythritol 4-phosphate pathway for microbial terpenoid production through metabolic control analysis. Microb Cell Fact 18 (2019) 192.

[18] C. Chassagnole, N. Noisommit-Rizzi, J.W. Schmid, K. Mauch, and M. Reuss, Dynamic modeling of the central carbon metabolism of *Escherichia coli*. Biotechnol Bioeng 79 (2002) 53-73.

[19] Y.T. Yang, G.N. Bennett, and K.Y. San, The effects of feed and intracellular pyruvate levels on the redistribution of metabolic fluxes in *Escherichia coli*. Metab Eng 3 (2001) 115-23.

[20] W.W. Cleland, The Enzymes, Academic Press, Inc., New York and London, 1970, pp. 1-65.

[21] M.M. Zhu, P.D. Lawman, and D.C. Cameron, Improving 1,3-propanediol production from glycerol in a metabolically engineered *Escherichia coli* by reducing accumulation of sn-glycerol-3-phosphate. Biotechnol Prog 18 (2002) 694-9.

[22] M.H. Buckstein, J. He, and H. Rubin, Characterization of nucleotide pools as a function of physiological state in *Escherichia coli*. J Bacteriol 190 (2008) 718-26.

[23] D.A. Schneider, and R.L. Gourse, Relationship between growth rate and ATP concentration in *Escherichia coli*: a bioassay for available cellular ATP. J Biol Chem 279 (2004) 8262-8.

[24] D.R. Lasko, and D.I. Wang, On-line monitoring of intracellular ATP concentration in *Escherichia coli* fermentations. Biotechnol Bioeng 52 (1996) 364-72.

[25] B.D. Bennett, E.H. Kimball, M. Gao, R. Osterhout, S.J. Van Dien, and J.D. Rabinowitz, Absolute metabolite concentrations and implied enzyme active site occupancy in *Escherichia coli*. Nat Chem Biol 5 (2009) 593-9.

[26] Y. Zhou, L. Wang, F. Yang, X. Lin, S. Zhang, and Z.K. Zhao, Determining the extremes of the cellular NAD(H) level by using an *Escherichia coli* NAD(+)-auxotrophic mutant. Appl Environ Microbiol 77 (2011) 6133-40.

[27] C. Bernal, C. Palacin, A. Boronat, and S. Imperial, A colorimetric assay for the determination of 4-diphosphocytidyl-2-C-methyl-D-erythritol 4-phosphate synthase activity. Anal Biochem 337 (2005) 55-61.

[28] H. Eoh, P. Narayanasamy, A.C. Brown, T. Parish, P.J. Brennan, and D.C. Crick, Expression and characterization of soluble 4-diphosphocytidyl-2-C-methyl-D-erythritol kinase from bacterial pathogens. Chem Biol 16 (2009) 1230-9.

[29] K. Nagano, and H. Nikaido, Kinetic behavior of the major multidrug efflux pump AcrB of *Escherichia coli*. Proc Natl Acad Sci U S A 106 (2009) 5854-8.

[30] C. Galian, F. Manon, M. Dezi, C. Torres, C. Ebel, D. Levy, and J.M. Jault, Optimized purification of a heterodimeric ABC transporter in a highly stable form amenable to 2-D crystallization. PLoS One 6 (2011) e19677.

[31] W. Zhang, Q. Shi, S.O. Meroueh, S.B. Vakulenko, and S. Mobashery, Catalytic mechanism of penicillin-binding protein 5 of *Escherichia coli*. Biochemistry 46 (2007) 10113-21.

[32] J.G. Geist, S. Lauw, V. Illarionova, B. Illarionov, M. Fischer, T. Grawert, F. Rohdich, W. Eisenreich, J. Kaiser, M. Groll, C. Scheurer, S. Wittlin, J.L. Alonso-Gomez, W.B. Schweizer, A. Bacher, and F.

Diederich, Thiazolopyrimidine inhibitors of 2-methylerythritol 2,4-cyclodiphosphate synthase (IspF) from *Mycobacterium tuberculosis* and *Plasmodium falciparum*. ChemMedChem 5 (2010) 1092-101.

[33] F.M. Hahn, A.P. Hurlburt, and C.D. Poulter, *Escherichia coli* open reading frame 696 is idi, a nonessential gene encoding isopentenyl diphosphate isomerase. J Bacteriol 181 (1999) 4499-504.

[34] J.J. Pan, S.T. Chiou, and P.H. Liang, Product distribution and pre-steady-state kinetic analysis of *Escherichia coli* undecaprenyl pyrophosphate synthase reaction. Biochemistry 39 (2000) 10936-42.

[35] G. Ramamoorthy, M.L. Pugh, B.X. Tian, R.M. Phan, L.B. Perez, M.P. Jacobson, and C.D. Poulter, Synthesis and enzymatic studies of bisubstrate analogues for farnesyl diphosphate synthase. J Org Chem 80 (2015) 3902-13.

[36] S.B. Richard, A.M. Lillo, C.N. Tetzlaff, M.E. Bowman, J.P. Noel, and D.E. Cane, Kinetic analysis of *Escherichia coli* 2-C-methyl-D-erythritol-4-phosphate cytidyltransferase, wild type and mutants, reveals roles of active site amino acids. Biochemistry 43 (2004) 12189-97.

[37] D.E. Cane, C. Chow, A. Lillo, and I. Kang, Molecular cloning, expression and characterization of the first three genes in the mevalonate-independent isoprenoid pathway in *Streptomyces coelicolor*. Bioorg Med Chem 9 (2001) 1467-77.

[38] W. Wang, and E. Oldfield, Bioorganometallic chemistry with IspG and IspH: structure, function, and inhibition of the [Fe(4)S(4)] proteins involved in isoprenoid biosynthesis. Angew Chem Int Ed Engl 53 (2014) 4294-310.

[39] Y. Xiao, L. Chu, Y. Sanakis, and P. Liu, Revisiting the IspH catalytic system in the deoxyxylulose phosphate pathway: achieving high activity. J Am Chem Soc 131 (2009) 9931-3.

[40] Y. Xiao, G. Zahariou, Y. Sanakis, and P. Liu, IspG enzyme activity in the deoxyxylulose phosphate pathway: roles of the iron-sulfur cluster. Biochemistry 48 (2009) 10483-5.

[41] J. Querol-Audí, A. Boronat, J. Centelles, and S. Imperial, Catalytically Important Residues in *E. coli* 1-Deoxy-D-Xylulose 5-Phosphate Synthase. Journal of Biosciences and Medicines 2 (2014) 30-35.

[42] L.A. Brammer, J.M. Smith, H. Wade, and C.F. Meyers, 1-Deoxy-D-xylulose 5-phosphate synthase catalyzes a novel random sequential mechanism. J Biol Chem 286 (2011) 36522-31.

[43] D.T. Fox, and C.D. Poulter, Mechanistic studies with 2-C-methyl-D-erythritol 4-phosphate synthase from *Escherichia coli*. Biochemistry 44 (2005) 8360-8.

[44] J. de Ruyck, V. Durisotti, Y. Oudjama, and J. Wouters, Structural role for Tyr-104 in *Escherichia coli* isopentenyl-diphosphate isomerase: site-directed mutagenesis, enzymology, and protein crystallography. J Biol Chem 281 (2006) 17864-9.

[45] F. Rohdich, J. Wungsintaweekul, M. Fellermeier, S. Sagner, S. Herz, K. Kis, W. Eisenreich, A. Bacher, and M.H. Zenk, Cytidine 5'-triphosphate-dependent biosynthesis of isoprenoids: YgbP protein of *Escherichia coli* catalyzes the formation of 4-diphosphocytidyl-2-C-methylerythritol. Proc Natl Acad Sci U S A 96 (1999) 11758-63.

[46] M. Tang, S.I. Odejinmi, Y.M. Allette, H. Vankayalapati, and K. Lai, Identification of novel small molecule inhibitors of 4-diphosphocytidyl-2-C-methyl-D-erythritol (CDP-ME) kinase of Gram-negative bacteria. Bioorg Med Chem 19 (2011) 5886-95.

[47] F. Zepeck, T. Grawert, J. Kaiser, N. Schramek, W. Eisenreich, A. Bacher, and F. Rohdich, Biosynthesis of isoprenoids. purification and properties of IspG protein from *Escherichia coli*. J Org Chem 70 (2005) 9168-74.

[48] J.T. Wan, and J.T. Jarrett, Electron acceptor specificity of ferredoxin (flavodoxin):NADP+ oxidoreductase from *Escherichia coli*. Arch Biochem Biophys 406 (2002) 116-26.

[49] S.P. Lim, and H. Nikaido, Kinetic parameters of efflux of penicillins by the multidrug efflux transporter AcrAB-TolC of *Escherichia coli*. Antimicrob Agents Chemother 54 (2010) 1800-6.

[50] A. Baici, M. Novinec, and B. Lenarcic, Kinetics of the Interaction of Peptidases with Substrates and Modifiers. in: K. Brix, and W. Stoker, (Eds.), Proteases: Structure and Function, 2013, pp. 37-84.

[51] B. Volkmer, and M. Heinemann, Condition-dependent cell volume and concentration of *Escherichia coli* to facilitate data conversion for systems biology modeling. PLoS One 6 (2011) e23126.

**Supplementary File 2**


| | | Pag. |
|---|---|---|
| S2 | Sensitivity analysis. ………………………………………...………………… | 2 |
| Table S2.1 | Summary sensitivity of the metabolites at the model parameters perturbation. … | 8 |
| Table S2.2 | Summary sensitivity of the fluxes at the model parameters perturbations………. | 9 |
| Table S2.3 | Metabolites sensitivities Log Gains for the independent variables. ……..………. | 10 |
| Table S2.4 | Rate constants sensitivities for the independent variables. …………………...… | 12 |
| Table S2.5 | Kinetic orders sensitivities for the independent variables. ………………..…..… | 13 |
| Table S2.6 | Logarithmic gains sensitivities for the fluxes. …………………..……………… | 17 |
| Table S2.7 | Rate constants sensitivities for the fluxes. ………………………………………. | 19 |
| Table S2.8 | Kinetic orders sensitivities for the fluxes. ………………………………………. | 20 |
| | References. ……………...…………………………………………….…………. | 24 |

### *S2.- Sensitivity analysis*

The sensitivity analysis indicates if the model is able to tolerate structural changes [1; 2].

In all cases sensitivity with a magnitude greater than 1 implies amplification of the original alteration; a magnitude less than 1 indicates attenuation. A positive sign for the sensitivity indicates that the changes are in the same direction: both increase in value or both decreases. A negative sign indicates that the changes are in the opposite direction. In a good model the parameter sensitivities are in general small. Even with slightly altered parameters, the model will continue to exhibit essentially the same structure and behavior. On the contrary, section models with unusually high parameter sensitivities coul be ill determined and the pattern of the sensitivity profile can be used to suggest portions of the model that need further attention.

In an S-system there are two types of parameters for analyzing the system sensitivity: the rate constants and the kinetic orders (ko's), which are defined as follows:

*Rate constant sensitivities:*

A rate constant sensitivity coefficient is defined as the ratio of a relative change in a dependent concentration $X_i$ to a relative change in a rate constant, $\alpha_j$ or $\beta_j$. They can be determined by differentiation of the explicit steady state solution.

$$S(X_i, \alpha_j) = \left(\frac{\partial X_i}{\partial \alpha_j}\frac{\alpha_j}{X_i}\right)_0 = \frac{\partial(\log X_i)}{\partial(\log \alpha_j)} \qquad \text{Eq. S20}$$

$$S(X_i, \beta_j) = \left(\frac{\partial X_i}{\partial \beta_j}\frac{\beta_j}{X_i}\right)_0 = \frac{\partial(\log X_i)}{\partial(\log \beta_j)} \qquad \text{Eq. S21}$$

Similarly the ratio of a relative change in a dependent flux to relative change in a rate-constant parameter is defined as:

$$S(V_i, \alpha_j) = \left(\frac{\partial V_i}{\partial \alpha_j}\frac{\alpha_j}{V_i}\right)_0 = \frac{\partial(\log V_i)}{\partial(\log \alpha_j)} \qquad \text{Eq. S22}$$

$$S(V_i, \beta_j) = \left(\frac{\partial V_i}{\partial \beta_j}\frac{\beta_j}{X_i}\right)_0 = \frac{\partial(\log V_i)}{\partial(\log \beta_j)} \qquad \text{Eq. S23}$$

the subscript 0 refers at the steady state.

Rate constant sensitivities are systemic properties that in general depend upon all the ko's of a system, i.e., they are properties of the integrated system and not just of its isolated components.

$S(X_i, \alpha_j)$ and $S(X_i, \beta_j)$ sensitivities are equal in magnitude, but negative in sign, as consequence only the $S(X_i, \alpha_j)$ is presented (Table S2.4).

The results summary sensitivities for metabolites and fluxes are showed in Tables S1.4 and S1.5 respectively.

Regarding the $|S(X_i, \alpha_j)|$ sensitivities (Table S2.1 and S2.4) from a total number of 100 values a 65% are below 1 with no the biggest sensitivity values (above or equal 10). The dependent variables most affected by changes in the rate constants are those related with the end part of the metabolic route are $X_4, X_5, X_8$ and $X_9$.

The values for the $|S(V_i, \alpha_j)|$ sensitivities (Tables S2.2 and S2.7) a 78% lower than one they are also more evenly distributed, although the fluxes most affected by changes in the rate constants are those related with the end part of the metabolic route are $V_4, V_5, V_8$ and $V_9$.

*Kinetic orders sensitivities:*

A kinetic order (ko) sensitivity coefficient is defined as the ratio of a relative change in a dependent concentration $X_i$ to a relative change in a ko parameter, $g_{j,k}$ or $h_{j,k}$. It can be determined by differentiation of the explicit steady state solution, with respect to the parameter in question. The corresponding expressions are:

$$S(X_i, g_{j,k}) = \left(\frac{\partial X_i}{\partial g_{j,k}} \frac{g_{j,k}}{X_i}\right)_0 = \frac{\partial(\log X_i)}{\partial(\log g_{j,k})} \quad \text{Eq. S24}$$

$$S(X_i, h_{j,k}) = \left(\frac{\partial X_i}{\partial h_{j,k}} \frac{h_{j,k}}{X_i}\right)_0 = \frac{\partial(\log X_i)}{\partial(\log h_{j,k})} \quad \text{Eq. S25}$$

The ratio of a relative change in a dependent flux to a relative change in a ko is similarly defined as:

$$S(V_i, g_{j,k}) = \left(\frac{\partial V_i}{\partial g_{j,k}} \frac{g_{j,k}}{V_i}\right)_0 = \frac{\partial(\log V_i)}{\partial(\log g_{j,k})} \quad \text{Eq. S26}$$

$$S(V_i, h_{j,k}) = \left(\frac{\partial V_i}{\partial h_{j,k}} \frac{h_{j,k}}{V_i}\right)_0 = \frac{\partial(\log V_i)}{\partial(\log h_{j,k})} \quad \text{Eq. S27}$$

The ko sensitivities can be calculated by differentiation of the explicit solution. Like the rate constant sensitivities, these ko's sensitivities are properties of the integrated system and not of its isolated components. Similarly the sensitivities are a function of both rate constants and ko's.

Tables S2.1 and S2.5 shown the sensitivity profile for the ko's. Most of the dependent variable sensitivities, 1180 out of 1640 (71.95%) have absolute values smaller than one. Only 83 from 1640 (5.06 %) have absolute values bigger than 10.

The highest sensitivities are those related with the synthesis and degradation of DXP ($X_1$) the premier metabolite de le route, the degradation of DMAPP ($X_8$) by IspH ($X_{21}$) and synthesis and degradation of MEP ($X_6$) by IspD ($X_{20}$) as well as the extrusion of $MEC_{(c)}$ ($X_7$) through the efflux pump ($X_{13}$). For instance, some of the kinetics orders that more affect the metabolites above mentioned are, $g_{1,24}$,

$h_{1,27}$, $h_{1,68}$, $h_{6,6}$, $h_{6,20}$, $g_{6,68}$, $h_{3,12}$ and $h_{3,14}$ between others. They values with absolute value sensitivities equal or bigger than 10 are highlighted in bold in Table S2.5.

Some kinetics orders associated with the synthesis or degradation of MEC(e) $X_{10}$ have so small value over the core metabolites of the MEP pathway $(X_1..X_9)$ that are represented as cero.

Table S2.2 present the model fluxes kinetics orders sensitivities statistics from Table S2.8. Table S2.8 shows similarities with respect at the metabolites kinetics orders with higher sensibilities (Table S2.5) but their values are in average smaller. In Table S2.2 can be seen that the 71.95% (1180 from 1640) have absolute values smaller than one. On the other hand a 5.06% (83 from 1640) presents absolute values sensitivities bigger than 10.

In Table S2.8 can be appreciated that the kinetics orders directly associated with the flux MEC(e) $X_{10}$ have an small influence over the rest of the model fluxes with the exception of the ko's directly related with the flux thought MEC(c) ($X_7$).Some high absolute values ko's are well dispersed between all the fluxes for example $h_{2,12,}$ $g_{3,12,}$ $h_{3,14,}$ $h_{6,20}$ $h_{6,68}$ affecting principally the fluxes through the nodes $V_4$, $V_5$, $V_8$, and $V_9$. Interestingly, the kinetics order $g_{1,24}$ indicates that a change in one direction of this ko will have a contrary general effect over all the model fluxes.

*Logarithmic gains.*

There are two main kinds of logarithmic gains: concentration and flux logarithmic gains. The concentration logarithmic gain is defined as:

$$L(X_i, X_j) = \left(\frac{\partial X_i}{\partial X_j}\frac{X_j}{X_i}\right)_0 = \frac{\partial(\log X_i)}{\partial(\log X_j)} \qquad \text{Eq. S28}$$

where $X_i$ stands for a dependent variable (the metabolite concentration) and $X_j$ for an independent variable (usually an enzyme activity or transport step). In a similar fashion, one obtains the logarithmic gains in the dependent fluxes through the pools of the system:

$$L(V_i, X_j) = \left(\frac{\partial V_i}{\partial X_j}\frac{X_j}{V_i}\right)_0 = \frac{\partial(\log V_i)}{\partial(\log X_j)} \qquad \text{Eq. S29}$$

where $V_i$ represents a given flux and $X_i$ and $X_j$ are defined as before. These sensitivity coefficients represent the finite percentage change in a dependent concentration $X_i$ or flux $V_i$, resulting from an infinitesimal change in an independent concentration $X_j$while all other independent concentrations and parameters are held constant. A logarithmic gain with a magnitude greater than one implies amplification of the original signal; a magnitude less than one indicates attenuation. A positive sign for the logarithmic gain indicates that the changes are in the same direction, both increase in value or both decrease. A negative sign indicates that the changes are in the opposite direction. The influence of a given independent variable on a particular dependent variable is given by the magnitude of the corresponding logarithmic gain.

The influence of the independent variables on the dependent variables and fluxes through the pools are resumed in terms of their percents in Tables S2.1 and S2.1.

Table S2.1shows that most of the absolute values log gains, 573 from 620 (92.42%) are below one with no log-gain bigger than 10. It is an important highlight that a percent important of the log-gains have values equal cero (48.87%) which indicates than almost half of the model is not affected by infinitesimal changes in independent variables from others parts of the model.

The highest logarithmic gains (Table S2.3) are expressed for DXS $(X_{24})$ and $K_{cat_{24}}(X_{36})$, both directly related with the synthesis of DXP $(X_1)$, the first metabolite of the route. The perturbation of any of $(X_{24})$ or $(X_{36})$ affects in general all the dependent variables of the model. Also the extrusion pump $(X_{13})$ has had a general effect in the rest of the model but in this last case with log-gains negatives.

The influence of the time independent variables on the fluxes through the pools presents the statistic shown in Table S2.2. Most of the logarithmic gains absolute values, 573 from 620 (92.42%) are below one and only 47 of them (7.58%) have values bigger or equal to one.

The percent of log-gains are quite balanced between positives and negatives values. For the case of the log gains with absolute values lesser than one the values are a 24.96% for the negatives and a 26.18% for the log gains with absolute positives one's. In case of the log-gains with absolute value bigger or equal to one the log-gains negatives represent a 44.68% and the positives represent a 55.32% (see Table S2.2). This indicate an equilibrium of "forces" predominant in case of a model perturbation as can be the case for example of a change in the enzymatic activity of some enzyme of the model.

*Dynamic behavior*. Finally, we consider the dynamic that characterize the transient responses from the initial values to the steady state as consequence of a global perturbation at time cero in the time dependent variables. Such analysis presents a consistency and reliability of the mathematical representation as they in all cases are stable (not shown).

The examination of the mode's robustness in the previous section suggests that the model for the isopropenes production in *E. coli* cell recycle bioreactor is well determined. These results are supported by the steady state analysis of the systemic behavior seen in the sensitivity and logarithmic gain factors compendium (Tables S2.1 and S2.2) and the predictions of the dynamic behavior (Figs. 2a-d and S3.1, S3.2, S3.3). Thus, this model provides a robust basis for optimization analysis.

**Table S2.1.** Sensitivity of the metabolites at the model parameters perturbations [(1)]

| | Total | % of Total | % of Total < 0 | % of Total = 0 | % of Total > 0 |
|---|---|---|---|---|---|
| **All** | | | | | |
| Log gains | 620 | 100 | 26,45 | 45,16 | 28,39 |
| Rate constants | 100 | 100 | 28 | 14 | 58 |
| Kinetic orders | 1640 | 100 | 43,84 | 11,89 | 44,27 |
| **Absolute value < 1** | | | | | |
| Log gains | 573 | 92,42 | 24,96 | 48,87 | 26,18 |
| Rate constants | 65 | 65 | 43,08 | 21,54 | 35,38 |
| Kinetic orders | 1180 | 71,95 | 41,44 | 16,53 | 42,03 |
| **Absolute value ≥ 1** | | | | | |
| Log gains | 47 | 7,58 | 44,68 | N.A.[(2)] | 55,32 |
| Rate constants | 35 | 35 | 0 | N.A. | 100 |
| Kinetic orders | 460 | 28,05 | 50 | N.A. | 50 |
| **Absolute value ≥ 10** | | | | | |
| Log gains | 0 | 0 | 0 | N.A. | 0 |
| Rate constants | 0 | 0 | 0 | N.A. | 0 |
| Kinetic orders | 83 | 5,06 | 50,6 | N.A. | 49.4 |

(1) Statistics from the raw data from Tables S2.3, S2.4, and S2.5

(2) N.A. : not applies

**Table S2.2.** Sensitivities for the fluxes at the model parameters perturbations [(1)]

| | Total | % of Total | % of Total < 0 | % of Total = 0 | % of Total > 0 |
|---|---|---|---|---|---|
| **All** | | | | | |
| Log gains | 620 | 100 | 26,45 | 45,16 | 28,39 |
| Rate constants | 100 | 100 | 28 | 14 | 58 |
| Kinetic orders | 1640 | 100 | 43,84 | 11,89 | 44,27 |
| **Absolute value < 1** | | | | | |
| Log gains | 573 | 92,42 | 24,96 | 48,87 | 26,18 |
| Rate constants | 64 | 64 | 43,75 | 21,88 | 34,38 |
| Kinetic orders | 1180 | 71,95 | 41,44 | 16,53 | 42,03 |
| **Absolute value ≥ 1** | | | | | |
| Log gains | 47 | 7,58 | 44,68 | N.A.[(2)] | 55,32 |
| Rate constants | 36 | 36 | 0 | N.A. | 100 |
| Kinetic orders | 460 | 28,05 | 50 | N.A. | 50 |
| **Absolute value ≥ 10** | | | | | |
| Log gains | 0 | 0 | 0 | N.A. | 0 |
| Rate constants | 0 | 0 | 0 | N.A. | 0 |
| Kinetic orders | 83 | 5,06 | 50,6 | N.A. | 49,4 |

(1) Statistics from the raw data Tables S2.6, S2.7, and S2.8

(2) N.A. : not applies

**Table S2.3.**Metabolites sensitivities Log Gains for the independent variables

| | $X_1$ | $X_2$ | $X_3$ | $X_4$ | $X_5$ | $X_6$ | $X_7$ | $X_8$ | $X_9$ | $X_{10}$ |
|---|---|---|---|---|---|---|---|---|---|---|
| $X_{11}$ | 0,00007 | 0,00006 | 0 | -0,01457 | -0,00632 | 0,00007 | 0,00007 | 0,00007 | -0,03557 | 0,00006 |
| $X_{12}$ | 0 | -1,01623 | 0 | 0 | 0 | 0 | 0 | 0 | 0 | 0 |
| $X_{13}$ | 0,00728 | 0,007 | 0 | -1,14109 | -1,17584 | 0,00724 | -1,16937 | -1,16781 | -1,12228 | 0,00951 |
| $X_{14}$ | 0 | 0 | -1,00458 | 0 | 0 | 0 | 0 | 0 | 0 | 0 |
| $X_{15}$ | -0,00024 | -0,00023 | 0 | 0,19061 | -0,11313 | -0,00024 | -0,00027 | -0,00027 | -0,1012 | -0,00023 |
| $X_{16}$ | 0,0042 | 0,00404 | 0 | -0,66223 | -0,6735 | 0,00418 | 0,00469 | 0,00468 | 0,35716 | 0,00397 |
| $X_{17}$ | 0,03347 | 0,03221 | 0,00001 | 0,0365 | 0,03761 | 0,03331 | 0,0374 | 0,03735 | 0,0359 | 0,03168 |
| $X_{18}$ | 0,51887 | 0,4993 | 0,00021 | 0,5658 | 0,58304 | 0,51639 | 0,57983 | 0,57905 | 0,55648 | 0,49113 |
| $X_{19}$ | 0 | -0,00199 | 0 | 0 | 0 | 0 | 0 | 0 | 0 | 0 |
| $X_{20}$ | 0 | 0 | 1,00416 | 0 | 0 | -1,05103 | 0 | 0 | 0 | 0 |
| $X_{21}$ | 0 | 0 | 0 | 0 | 0 | 0 | 0 | -1,00547 | 0 | 0 |
| $X_{22}$ | -0,003 | 0 | 0 | 0 | 0 | 0 | 0 | 0 | 0 | 0 |
| $X_{23}$ | -0,00621 | -0,00597 | 0 | 0,97312 | 1,00277 | -0,00618 | -0,00957 | 0,99591 | 0,95708 | -0,00811 |
| $X_{24}$ | 1,04877 | 1,00921 | 0,00042 | 1,14364 | 1,17848 | 1,04377 | 1,17199 | 1,17042 | 1,12479 | 0,99271 |
| $X_{25}$ | 0 | 0 | 0,01071 | 0 | 0 | -0,01121 | 0 | 0 | 0 | 0 |
| $X_{26}$ | 0,00007 | 0,00006 | 0 | -0,01457 | -0,00632 | 0,00007 | 0,00007 | 0,00007 | -0,03557 | 0,00006 |
| $X_{27}$ | -1,05606 | 0 | 0 | 0 | 0 | 0 | 0 | 0 | 0 | 0 |
| $X_{28}$ | 0 | -1,01623 | 0 | 0 | 0 | 0 | 0 | 0 | 0 | 0 |
| $X_{29}$ | 0,00728 | 0,007 | 0 | -1,14109 | -1,17584 | 0,00724 | -1,16937 | -1,16781 | -1,12228 | 0,00951 |
| $X_{30}$ | 0 | 0 | -1,00458 | 0 | 0 | 0 | 0 | 0 | 0 | 0 |
| $X_{31}$ | -0,00024 | -0,00023 | 0 | 0,19061 | -0,11313 | -0,00024 | -0,00027 | -0,00027 | -0,1012 | -0,00023 |
| $X_{32}$ | 0,0042 | 0,00404 | 0 | -0,66223 | -0,6735 | 0,00418 | 0,00469 | 0,00468 | 0,35716 | 0,00397 |
| $X_{33}$ | 0 | 0 | 1,00416 | 0 | 0 | -1,05103 | 0 | 0 | 0 | 0 |
| $X_{34}$ | 0 | 0 | 0 | 0 | 0 | 0 | 0 | -1,00547 | 0 | 0 |
| $X_{35}$ | -0,00621 | -0,00597 | 0 | 0,97312 | 1,00277 | -0,00618 | -0,00957 | 0,99591 | 0,95708 | -0,00811 |
| $X_{36}$ | 1,04877 | 1,00921 | 0,00042 | 1,14364 | 1,17848 | 1,04377 | 1,17199 | 1,17042 | 1,12479 | 0,99271 |
| $X_{37}$ | 0 | 0 | 0 | 0,00004 | 0,00002 | 0 | 0 | 0 | 0,0001 | 0 |
| $X_{38}$ | 1 | 0 | 0 | 0 | 0 | 0 | 0 | 0 | 0 | 0 |
| $X_{39}$ | 0 | 1 | 0 | 0 | 0 | 0 | 0 | 0 | 0 | 0 |
| $X_{40}$ | 0 | 0 | 1 | 0 | 0 | 0 | 0 | 0 | 0 | 0 |
| $X_{41}$ | -0,00056 | -0,00054 | 0 | 0,4438 | -0,26341 | -0,00056 | -0,00062 | -0,00062 | -0,23564 | -0,00053 |
| $X_{42}$ | 0,00078 | 0,00075 | 0 | -0,62088 | 0,36851 | 0,00078 | 0,00087 | 0,00087 | 0,32966 | 0,00074 |
| $X_{43}$ | 0 | 0 | -0,95541 | 0 | 0 | 1 | 0 | 0 | 0 | 0 |
| $X_{44}$ | 0,00616 | 0,00593 | 0 | -0,96653 | -0,99597 | 0,00613 | 0,00951 | -0,98917 | -0,9506 | 0,00805 |
| $X_{45}$ | 0 | 0 | 0 | 0 | 0 | 0 | 0 | 1 | 0 | 0 |
| $X_{46}$ | -0,00022 | -0,00021 | 0 | 0,0343 | 0,03489 | -0,00022 | -0,00024 | -0,00024 | -0,0185 | -0,00021 |
| $X_{47}$ | -0,00391 | -0,00376 | 0 | 0,61746 | 0,62797 | -0,00389 | -0,00437 | -0,00437 | -0,33301 | -0,0037 |
| $X_{48}$ | -0,02532 | -0,02437 | -0,00001 | -0,02761 | -0,02846 | -0,0252 | -0,0283 | -0,02826 | -0,02716 | -0,02397 |
| $X_{49}$ | -0,51887 | -0,4993 | -0,00021 | -0,5658 | -0,58304 | -0,51639 | -0,57983 | -0,57905 | -0,55648 | -0,49113 |
| $X_{50}$ | 0 | 0,00199 | 0 | 0 | 0 | 0 | 0 | 0 | 0 | 0 |
| $X_{51}$ | 0 | 0 | 0 | 0 | 0 | 0 | 0 | 0,00437 | 0 | 0 |
| $X_{52}$ | 0,00003 | 0,00003 | 0 | -0,00527 | -0,00543 | 0,00003 | 0,00005 | -0,00539 | -0,00518 | 0,00004 |
| $X_{53}$ | 0,00023 | 0 | 0 | 0 | 0 | 0 | 0 | 0 | 0 | 0 |
| $X_{54}$ | 0 | 0 | -0,01071 | 0 | 0 | 0,01121 | 0 | 0 | 0 | 0 |
| $X_{55}$ | 0 | 0 | 0 | 0 | 0 | 0 | 0 | 0 | 0 | -1 |
| $X_{56}$ | -0,00007 | -0,00006 | 0 | 0,01452 | 0,0063 | -0,00007 | -0,00007 | -0,00007 | 0,03546 | -0,00006 |
| $X_{57}$ | 0,00311 | 0,00299 | 0 | 0,00339 | 0,00349 | 0,00309 | 0,00347 | 0,00347 | 0,00333 | 0,00294 |
| $X_{58}$ | 0,00227 | 0,00218 | 0 | -0,50398 | -0,21873 | 0,00226 | 0,00253 | 0,00253 | -0,21535 | 0,00215 |

| | | | | | | | | | | |
|---|---|---|---|---|---|---|---|---|---|---|
| $X_{59}$ | 0,00313 | 0,00301 | 0 | 0,00341 | 0,00352 | 0,00312 | 0,0035 | 0,00349 | 0,00336 | 0,00296 |
| $X_{60}$ | -0,00616 | -0,00593 | 0 | 0,96653 | 0,99597 | -0,00613 | 0,99049 | 0,98917 | 0,9506 | -0,00805 |
| $X_{61}$ | 0 | 0 | 0,99098 | 0 | 0 | 0 | 0 | 0 | 0 | 0 |
| $X_{62}$ | -0,00006 | -0,00006 | 0 | 0,01445 | 0,00627 | -0,00006 | -0,00007 | -0,00007 | 0,03527 | -0,00006 |
| $X_{63}$ | 0 | 0 | -0,99114 | 0 | 0 | 0 | 0 | 0 | 0 | 0 |
| $X_{64}$ | -0,00815 | -0,00784 | 0 | -0,00889 | -0,00916 | -0,00811 | -0,00911 | -0,00909 | -0,00874 | -0,00771 |
| $X_{65}$ | 0,00277 | 0 | 0 | 0 | 0 | 0 | 0 | 0 | 0 | 0 |
| $X_{66}$ | 0,00088 | 0,00084 | 0 | -0,13748 | -0,14166 | 0,00087 | 0,00135 | -0,1407 | -0,13521 | 0,00115 |
| $X_{67}$ | 0 | 0 | 0 | 0 | 0 | 0 | 0 | 0,14224 | 0 | 0 |
| $X_{68}$ | -1,05606 | 0 | 0 | 0 | 0 | 0 | 0 | 0 | 0 | 0 |
| $X_{69}$ | 0 | 0 | 0,95541 | 0 | 0 | 0 | 0 | 0 | 0 | 0 |
| $X_{70}$ | -0,04327 | -0,04164 | -0,00002 | 6,78493 | 6,9916 | -0,04307 | 6,95312 | 6,94382 | 6,67309 | -0,05653 |
| $X_{71}$ | -0,00091 | -0,00088 | 0 | 0,14274 | 0,14709 | -0,00091 | -0,0014 | -0,00052 | 0,14039 | -0,00119 |
| $X_{72}$ | -0,00007 | -0,00006 | 0 | 0,01449 | 0,00629 | -0,00006 | -0,00007 | -0,00007 | -0,96463 | -0,00006 |

**Table S2.4.** Rate constants sensitivities

| | $X_1$ | $X_2$ | $X_3$ | $X_4$ | $X_5$ | $X_6$ | $X_7$ | $X_8$ | $X_9$ | $X_{10}$ |
|---|---|---|---|---|---|---|---|---|---|---|
| $\alpha_1$ | 1,04877 | 1,00921 | 0,00042 | 1,14364 | 1,17848 | 1,04377 | 1,17199 | 1,17042 | 1,12479 | 0,99271 |
| $\alpha_2$ | -0,00729 | 1,00921 | 1,00458 | 1,14364 | 1,17848 | -0,00726 | 1,17199 | 1,17042 | 1,12479 | 0,99271 |
| $\alpha_3$ | -0,00729 | -0,00702 | 1,00458 | 1,14364 | 1,17848 | -0,00726 | 1,17199 | 1,17042 | 1,12479 | 0,99271 |
| $\alpha_4$ | -0,00525 | -0,00505 | 0 | 1,16627 | 0,50617 | -0,00522 | -0,00586 | -0,00585 | 0,49834 | -0,00497 |
| $\alpha_5$ | -0,00324 | -0,00312 | 0 | 0,24071 | 0,78889 | -0,00322 | -0,00362 | -0,00362 | 0,74111 | -0,00307 |
| $\alpha_6$ | -0,00729 | 1,00921 | 0,00042 | 1,14364 | 1,17848 | 1,04377 | 1,17199 | 1,17042 | 1,12479 | 0,99271 |
| $\alpha_7$ | -0,00729 | -0,00702 | 0 | 1,14364 | 1,17848 | -0,00726 | 1,17199 | 1,17042 | 1,12479 | 0,99271 |
| $\alpha_8$ | -0,00622 | -0,00599 | 0 | 0,97568 | 1,0054 | -0,00619 | -0,00695 | 0,99853 | 0,9596 | -0,00589 |
| $\alpha_9$ | 0,00007 | 0,00006 | 0 | -0,01493 | -0,00648 | 0,00007 | 0,00008 | 0,00007 | 0,99379 | 0,00006 |
| $\alpha_{10}$ | 0 | 0 | 0 | 0 | 0 | 0 | 0 | 0 | 0 | 1 |

**Table S2.5.** Kinetic orders sensitivities for the independent variables.

| | $X_1$ | $X_2$ | $X_3$ | $X_4$ | $X_5$ | $X_6$ | $X_7$ | $X_8$ | $X_9$ | $X_{10}$ |
|---|---|---|---|---|---|---|---|---|---|---|
| g(1,4) | 0,02434 | 0,02343 | 0,00001 | 0,02655 | 0,02735 | 0,02423 | 0,0272 | 0,02717 | 0,02611 | 0,02304 |
| g(1,5) | 0,02382 | 0,02292 | 0,00001 | 0,02597 | 0,02676 | 0,0237 | 0,02661 | 0,02658 | 0,02554 | 0,02254 |
| g(1,17) | 0,03287 | 0,03163 | 0,00001 | 0,03585 | 0,03694 | 0,03272 | 0,03673 | 0,03669 | 0,03525 | 0,03112 |
| g(1,18) | -1,65736 | -1,59485 | -0,00066 | -1,80728 | -1,86233 | -1,64945 | -1,85208 | -1,8496 | -1,77749 | -1,56877 |
| g(1,24) | **-10,98945** | **-10,57498** | -0,00436 | **-11,98356** | **-12,34857** | **-10,93703** | **-12,28062** | **-12,26419** | **-11,78602** | **-10,40205** |
| g(1,36) | 4,8329 | 4,65063 | 0,00192 | 5,27009 | 5,43061 | 4,80985 | 5,40073 | 5,3935 | 5,18322 | 4,57458 |
| g(1,48) | 0,0509 | 0,04898 | 0,00002 | 0,0555 | 0,05719 | 0,05066 | 0,05688 | 0,0568 | 0,05459 | 0,04818 |
| g(1,49) | 1,65736 | 1,59485 | 0,00066 | 1,80728 | 1,86233 | 1,64945 | 1,85208 | 1,8496 | 1,77749 | 1,56877 |
| g(1,57) | -0,00848 | -0,00816 | 0 | -0,00924 | -0,00952 | -0,00844 | -0,00947 | -0,00946 | -0,00909 | -0,00802 |
| g(1,59) | -0,00786 | -0,00756 | 0 | -0,00857 | -0,00883 | -0,00782 | -0,00878 | -0,00877 | -0,00843 | -0,00744 |
| g(1,64) | 0,02572 | 0,02475 | 0,00001 | 0,02804 | 0,0289 | 0,02559 | 0,02874 | 0,0287 | 0,02758 | 0,02434 |
| h(1,1) | 5,94243 | 5,71831 | 0,00236 | 6,47998 | 6,67736 | 5,91408 | 6,64061 | 6,63173 | 6,37317 | 5,6248 |
| h(1,22) | 0,00632 | 0,00608 | 0 | 0,00689 | 0,0071 | 0,00629 | 0,00707 | 0,00706 | 0,00678 | 0,00598 |
| h(1,27) | -7,82554 | -7,53039 | -0,00311 | -8,53344 | -8,79336 | -7,78821 | -8,74497 | -8,73327 | -8,39277 | -7,40725 |
| h(1,38) | -3,07967 | -2,96352 | -0,00122 | -3,35826 | -3,46055 | -3,06498 | -3,44151 | -3,43691 | -3,30291 | -2,91506 |
| h(1,53) | -0,00176 | -0,00169 | 0 | -0,00191 | -0,00197 | -0,00175 | -0,00196 | -0,00196 | -0,00188 | -0,00166 |
| h(1,65) | -0,02203 | -0,02119 | -0,00001 | -0,02402 | -0,02475 | -0,02192 | -0,02461 | -0,02458 | -0,02362 | -0,02085 |
| h(1,68) | **11,66605** | **11,22606** | 0,00463 | **12,72137** | **13,10885** | **11,6104** | **13,03671** | **13,01927** | **12,51167** | **11,04248** |
| g(2,6) | 0,06237 | -8,62947 | -8,58981 | -9,77892 | **-10,07678** | 0,06207 | **-10,02132** | **-10,00792** | -9,61772 | -8,48836 |
| g(2,20) | 0,07788 | **-10,77566** | **-10,72613** | **-12,21098** | **-12,58291** | 0,07751 | **-12,51367** | **-12,49693** | **-12,00969** | **-10,59945** |
| g(2,25) | -0,00008 | 0,01069 | 0,01064 | 0,01211 | 0,01248 | -0,00008 | 0,01241 | 0,0124 | 0,01191 | 0,01051 |
| g(2,33) | -0,0536 | 7,4163 | 7,38221 | 8,40415 | 8,66013 | -0,05334 | 8,61247 | 8,60095 | 8,26561 | 7,29502 |
| g(2,43) | -0,04 | 5,53425 | 5,50882 | 6,27141 | 6,46244 | -0,03981 | 6,42687 | 6,41827 | 6,16803 | 5,44375 |
| g(2,54) | -0,00002 | 0,00295 | 0,00294 | 0,00335 | 0,00345 | -0,00002 | 0,00343 | 0,00343 | 0,00329 | 0,00291 |
| h(2,2) | -0,04413 | 6,1064 | 6,07834 | 6,91977 | 7,13055 | -0,04392 | 7,09131 | 7,08182 | 6,80571 | 6,00655 |
| h(2,12) | -0,06748 | 9,33701 | 9,2941 | **10,5807** | **10,90298** | -0,06716 | **10,84298** | **10,82847** | **10,40628** | 9,18432 |
| h(2,19) | 0,00002 | -0,00217 | -0,00216 | -0,00246 | -0,00254 | 0,00002 | -0,00253 | -0,00252 | -0,00242 | -0,00214 |
| h(2,28) | 0,05046 | -6,98145 | -6,94936 | -7,91137 | -8,15235 | 0,05022 | -8,10748 | -8,09664 | -7,78096 | -6,86728 |
| h(2,39) | 0,01362 | -1,88402 | -1,87536 | -2,13497 | -2,2 | 0,01355 | -2,18789 | -2,18496 | -2,09977 | -1,85321 |
| h(2,50) | 0,00001 | -0,00172 | -0,00171 | -0,00195 | -0,00201 | 0,00001 | -0,00199 | -0,00199 | -0,00191 | -0,00169 |
| g(3,2) | 0,04413 | 0,04247 | -6,07834 | -6,91977 | -7,13055 | 0,04392 | -7,09131 | -7,08182 | -6,80571 | -6,00655 |
| g(3,12) | 0,06748 | 0,06494 | -9,2941 | **-10,5807** | **-10,90298** | 0,06716 | **-10,84298** | **-10,82847** | **-10,40628** | -9,18432 |
| g(3,19) | -0,00002 | -0,00002 | 0,00216 | 0,00246 | 0,00254 | -0,00002 | 0,00253 | 0,00252 | 0,00242 | 0,00214 |
| g(3,28) | -0,05046 | -0,04855 | 6,94936 | 7,91137 | 8,15235 | -0,05022 | 8,10748 | 8,09664 | 7,78096 | 6,86728 |
| g(3,39) | -0,01362 | -0,0131 | 1,87536 | 2,13497 | 2,2 | -0,01355 | 2,18789 | 2,18496 | 2,09977 | 1,85321 |
| g(3,50) | -0,00001 | -0,00001 | 0,00171 | 0,00195 | 0,00201 | -0,00001 | 0,00199 | 0,00199 | 0,00191 | 0,00169 |
| h(3,3) | -0,04696 | -0,04519 | 6,46814 | 7,36354 | 7,58782 | -0,04674 | 7,54607 | 7,53597 | 7,24215 | 6,39174 |
| h(3,6) | -0,06234 | -0,05999 | 8,58623 | 9,77484 | **10,07257** | -0,06204 | **10,01714** | **10,00374** | 9,61371 | 8,48482 |
| h(3,14) | -0,07206 | -0,06934 | 9,92498 | **11,29891** | **11,64307** | -0,07172 | **11,579** | **11,56351** | **11,11266** | 9,80775 |
| h(3,30) | 0,03203 | 0,03082 | -4,41083 | -5,02143 | -5,17438 | 0,03187 | -5,1459 | -5,13902 | -4,93866 | -4,35873 |
| h(3,40) | 0,00785 | 0,00756 | -1,08175 | -1,2315 | -1,26901 | 0,00782 | -1,26203 | -1,26034 | -1,2112 | -1,06898 |
| h(3,61) | 0 | 0 | 0 | 0 | 0 | 0 | 0 | 0 | 0 | 0 |
| h(3,63) | 0,0547 | 0,05264 | -7,53355 | -8,57643 | -8,83766 | 0,05444 | -8,78903 | -8,77727 | -8,43506 | -7,44457 |
| h(3,69) | 0,03945 | 0,03796 | -5,43333 | -6,18547 | -6,37388 | 0,03926 | -6,3388 | -6,33032 | -6,08351 | -5,36915 |
| g(4,4) | -0,00244 | -0,00235 | 0 | 0,5431 | 0,23571 | -0,00243 | -0,00273 | -0,00273 | 0,23206 | -0,00231 |
| g(4,5) | 0,01494 | 0,01437 | 0,00001 | -3,32109 | -1,44138 | 0,01487 | 0,01669 | 0,01667 | -1,41908 | 0,01414 |
| g(4,8) | 0,03506 | 0,03374 | 0,00001 | -7,79595 | -3,38351 | 0,0349 | 0,03918 | 0,03913 | -3,33115 | 0,03319 |
| g(4,15) | 0,02275 | 0,02189 | 0,00001 | -5,05839 | -2,19539 | 0,02264 | 0,02542 | 0,02539 | -2,16141 | 0,02153 |
| g(4,21) | 0,03192 | 0,03072 | 0,00001 | -7,09701 | -3,08017 | 0,03177 | 0,03567 | 0,03562 | -3,0325 | 0,03021 |

| | | | | | | | | | | |
|---|---|---|---|---|---|---|---|---|---|---|
| g(4,31) | -0,00531 | -0,00511 | 0 | 1,18013 | 0,51219 | -0,00528 | -0,00593 | -0,00592 | 0,50426 | -0,00502 |
| g(4,34) | -0,01151 | -0,01107 | 0 | 2,55807 | 1,11023 | -0,01145 | -0,01286 | -0,01284 | 1,09305 | -0,01089 |
| g(4,41) | 0,00145 | 0,0014 | 0 | -0,32287 | -0,14013 | 0,00145 | 0,00162 | 0,00162 | -0,13796 | 0,00137 |
| g(4,42) | -0,00834 | -0,00802 | 0 | 1,85414 | 0,80472 | -0,0083 | -0,00932 | -0,00931 | 0,79226 | -0,00789 |
| g(4,45) | -0,01209 | -0,01164 | 0 | 2,68888 | 1,167 | -0,01204 | -0,01351 | -0,0135 | 1,14894 | -0,01145 |
| g(4,51) | -0,00007 | -0,00006 | 0 | 0,01483 | 0,00644 | -0,00007 | -0,00007 | -0,00007 | 0,00634 | -0,00006 |
| g(4,67) | -0,00373 | -0,00359 | 0 | 0,82903 | 0,35981 | -0,00371 | -0,00417 | -0,00416 | 0,35424 | -0,00353 |
| g(4,71) | 0,00293 | 0,00282 | 0 | -0,65251 | -0,2832 | 0,00292 | 0,00328 | 0,00328 | -0,27881 | 0,00278 |
| h(4,4) | -0,03806 | -0,03662 | -0,00002 | 8,4617 | 3,67246 | -0,03788 | -0,04253 | -0,04247 | 3,61562 | -0,03602 |
| h(4,5) | -0,00233 | -0,00225 | 0 | 0,51898 | 0,22524 | -0,00232 | -0,00261 | -0,0026 | 0,22176 | -0,00221 |
| h(4,9) | -0,00053 | -0,00051 | 0 | 0,11712 | 0,05083 | -0,00052 | -0,00059 | -0,00059 | 0,05004 | -0,0005 |
| h(4,11) | -0,00059 | -0,00057 | 0 | 0,13087 | 0,0568 | -0,00059 | -0,00066 | -0,00066 | 0,05592 | -0,00056 |
| h(4,15) | -0,00781 | -0,00752 | 0 | 1,73751 | 0,75409 | -0,00778 | -0,00873 | -0,00872 | 0,74242 | -0,0074 |
| h(4,16) | -0,02006 | -0,0193 | -0,00001 | 4,46044 | 1,93587 | -0,01997 | -0,02242 | -0,02239 | 1,90591 | -0,01899 |
| h(4,26) | 0,00004 | 0,00004 | 0 | -0,00824 | -0,00358 | 0,00004 | 0,00004 | 0,00004 | -0,00352 | 0,00004 |
| h(4,31) | 0,00182 | 0,00175 | 0 | -0,40536 | -0,17593 | 0,00181 | 0,00204 | 0,00203 | -0,17321 | 0,00173 |
| h(4,32) | -0,00057 | -0,00055 | 0 | 0,12709 | 0,05516 | -0,00057 | -0,00064 | -0,00064 | 0,05431 | -0,00054 |
| h(4,37) | 0 | 0 | 0 | -0,00023 | -0,0001 | 0 | 0 | 0 | -0,0001 | 0 |
| h(4,41) | 0,0112 | 0,01078 | 0 | -2,49052 | -1,08091 | 0,01115 | 0,01252 | 0,0125 | -1,06418 | 0,0106 |
| h(4,42) | -0,00781 | -0,00751 | 0 | 1,73606 | 0,75347 | -0,00777 | -0,00873 | -0,00871 | 0,7418 | -0,00739 |
| h(4,46) | 0,00079 | 0,00076 | 0 | -0,17608 | -0,07642 | 0,00079 | 0,00088 | 0,00088 | -0,07524 | 0,00075 |
| h(4,47) | 0,00757 | 0,00729 | 0 | -1,68368 | -0,73073 | 0,00754 | 0,00846 | 0,00845 | -0,71942 | 0,00717 |
| h(4,56) | 0,00031 | 0,0003 | 0 | -0,06846 | -0,02971 | 0,00031 | 0,00034 | 0,00034 | -0,02925 | 0,00029 |
| h(4,58) | -0,01201 | -0,01156 | 0 | 2,67024 | 1,15891 | -0,01195 | -0,01342 | -0,0134 | 1,14097 | -0,01137 |
| h(4,62) | 0,00018 | 0,00017 | 0 | -0,03927 | -0,01705 | 0,00018 | 0,0002 | 0,0002 | -0,01678 | 0,00017 |
| g(5,4) | 0,01497 | 0,0144 | 0,00001 | -1,11209 | -3,64473 | 0,0149 | 0,01673 | 0,01671 | -3,424 | 0,01417 |
| g(5,5) | -0,01111 | -0,01069 | 0 | 0,82545 | 2,7053 | -0,01106 | -0,01242 | -0,0124 | 2,54147 | -0,01052 |
| g(5,8) | 0,02786 | 0,02681 | 0,00001 | -2,06946 | -6,78239 | 0,02772 | 0,03113 | 0,03109 | -6,37164 | 0,02637 |
| g(5,15) | 0,00621 | 0,00597 | 0 | -0,46123 | -1,51161 | 0,00618 | 0,00694 | 0,00693 | -1,42007 | 0,00588 |
| g(5,21) | 0,02536 | 0,0244 | 0,00001 | -1,88393 | -6,17432 | 0,02524 | 0,02834 | 0,0283 | -5,8004 | 0,024 |
| g(5,31) | -0,00145 | -0,00139 | 0 | 0,1076 | 0,35266 | -0,00144 | -0,00162 | -0,00162 | 0,3313 | -0,00137 |
| g(5,34) | -0,00914 | -0,0088 | 0 | 0,67905 | 2,2255 | -0,0091 | -0,01021 | -0,0102 | 2,09072 | -0,00865 |
| g(5,41) | -0,0089 | -0,00856 | 0 | 0,66112 | 2,16672 | -0,00886 | -0,00994 | -0,00993 | 2,0355 | -0,00842 |
| g(5,42) | 0,0062 | 0,00597 | 0 | -0,46084 | -1,51035 | 0,00617 | 0,00693 | 0,00692 | -1,41888 | 0,00587 |
| g(5,45) | -0,00961 | -0,00925 | 0 | 0,71377 | 2,3393 | -0,00956 | -0,01074 | -0,01072 | 2,19763 | -0,00909 |
| g(5,51) | -0,00005 | -0,00005 | 0 | 0,00394 | 0,0129 | -0,00005 | -0,00006 | -0,00006 | 0,01212 | -0,00005 |
| g(5,67) | -0,00296 | -0,00285 | 0 | 0,22007 | 0,72124 | -0,00295 | -0,00331 | -0,00331 | 0,67756 | -0,0028 |
| g(5,71) | 0,00233 | 0,00224 | 0 | -0,17321 | -0,56768 | 0,00232 | 0,00261 | 0,0026 | -0,5333 | 0,00221 |
| h(5,4) | 0,0012 | 0,00115 | 0 | -0,08908 | -0,29194 | 0,00119 | 0,00134 | 0,00134 | -0,27426 | 0,00113 |
| h(5,5) | -0,02483 | -0,0239 | -0,00001 | 1,84481 | 6,04613 | -0,02471 | -0,02775 | -0,02771 | 5,67996 | -0,0235 |
| h(5,15) | -0,01807 | -0,01739 | -0,00001 | 1,34277 | 4,40074 | -0,01799 | -0,0202 | -0,02017 | 4,13423 | -0,01711 |
| h(5,16) | -0,01594 | -0,01534 | -0,00001 | 1,18404 | 3,88053 | -0,01586 | -0,01781 | -0,01779 | 3,64552 | -0,01509 |
| h(5,31) | 0,00422 | 0,00406 | 0 | -0,31327 | -1,0267 | 0,0042 | 0,00471 | 0,00471 | -0,96452 | 0,00399 |
| h(5,32) | -0,00045 | -0,00044 | 0 | 0,03374 | 0,11057 | -0,00045 | -0,00051 | -0,00051 | 0,10387 | -0,00043 |
| h(5,41) | -0,00115 | -0,00111 | 0 | 0,08571 | 0,28089 | -0,00115 | -0,00129 | -0,00129 | 0,26388 | -0,00109 |
| h(5,42) | 0,00663 | 0,00638 | 0 | -0,49219 | -1,61308 | 0,00659 | 0,0074 | 0,00739 | -1,51539 | 0,00627 |
| h(5,46) | 0,00063 | 0,00061 | 0 | -0,04674 | -0,15318 | 0,00063 | 0,0007 | 0,0007 | -0,14391 | 0,0006 |
| h(5,47) | 0,00602 | 0,00579 | 0 | -0,44694 | -1,46478 | 0,00599 | 0,00672 | 0,00671 | -1,37607 | 0,00569 |
| g(6,1) | 0,04133 | -5,71831 | -0,00236 | -6,47998 | -6,67736 | -5,91408 | -6,64061 | -6,63173 | -6,37317 | -5,6248 |
| g(6,22) | 0,00004 | -0,00608 | 0 | -0,00689 | -0,0071 | -0,00629 | -0,00707 | -0,00706 | -0,00678 | -0,00598 |
| g(6,27) | -0,05442 | 7,53039 | 0,00311 | 8,53344 | 8,79336 | 7,78821 | 8,74497 | 8,73327 | 8,39277 | 7,40725 |
| g(6,38) | -0,02142 | 2,96352 | 0,00122 | 3,35826 | 3,46055 | 3,06498 | 3,44151 | 3,43691 | 3,30291 | 2,91506 |
| g(6,53) | -0,00001 | 0,00169 | 0 | 0,00191 | 0,00197 | 0,00175 | 0,00196 | 0,00196 | 0,00188 | 0,00166 |
| g(6,65) | -0,00015 | 0,02119 | 0,00001 | 0,02402 | 0,02475 | 0,02192 | 0,02461 | 0,02458 | 0,02362 | 0,02085 |

| | | | | | | | | | | |
|---|---|---|---|---|---|---|---|---|---|---|
| g(6,68) | 0,08113 | **-11,22606** | -0,00463 | **-12,72137** | **-13,10885** | **-11,6104** | **-13,03671** | **-13,01927** | **-12,51167** | **-11,04248** |
| h(6,6) | -0,06237 | 8,62947 | 0,00356 | 9,77892 | **10,07678** | 8,92492 | **10,02132** | **10,00792** | 9,61772 | 8,48836 |
| h(6,20) | -0,07788 | 10,77566 | 0,00444 | **12,21098** | **12,58291** | **11,14458** | **12,51367** | **12,49693** | **12,00969** | **10,59945** |
| h(6,25) | 0,00008 | -0,01069 | 0 | -0,01211 | -0,01248 | -0,01106 | -0,01241 | -0,0124 | -0,01191 | -0,01051 |
| h(6,33) | 0,0536 | -7,4163 | -0,00306 | -8,40415 | -8,66013 | -7,67021 | -8,61247 | -8,60095 | -8,26561 | -7,29502 |
| h(6,43) | 0,04 | -5,53425 | -0,00228 | -6,27141 | -6,46244 | -5,72373 | -6,42687 | -6,41827 | -6,16803 | -5,44375 |
| h(6,54) | 0,00002 | -0,00295 | 0 | -0,00335 | -0,00345 | -0,00305 | -0,00343 | -0,00343 | -0,00329 | -0,00291 |
| g(7,3) | 0,04696 | 0,04519 | 0,00002 | -7,36354 | -7,58782 | 0,04674 | -7,54607 | -7,53597 | -7,24215 | -6,39174 |
| g(7,6) | 0,06234 | 0,05999 | 0,00002 | -9,77484 | **-10,07257** | 0,06204 | **-10,01714** | **-10,00374** | -9,61371 | -8,48482 |
| g(7,14) | 0,07206 | 0,06934 | 0,00003 | **-11,29891** | **-11,64307** | 0,07172 | **-11,579** | **-11,56351** | **-11,11266** | -9,80775 |
| g(7,30) | -0,03203 | -0,03082 | -0,00001 | 5,02143 | 5,17438 | -0,03187 | 5,1459 | 5,13902 | 4,93866 | 4,35873 |
| g(7,40) | -0,00785 | -0,00756 | 0 | 1,2315 | 1,26901 | -0,00782 | 1,26203 | 1,26034 | 1,2112 | 1,06898 |
| g(7,61) | 0 | 0 | 0 | 0 | 0 | 0 | 0 | 0 | 0 | 0 |
| g(7,63) | -0,0547 | -0,05264 | -0,00002 | 8,57643 | 8,83766 | -0,05444 | 8,78903 | 8,77727 | 8,43506 | 7,44457 |
| g(7,69) | -0,03945 | -0,03796 | -0,00002 | 6,18547 | 6,37388 | -0,03926 | 6,3388 | 6,33032 | 6,08351 | 5,36915 |
| h(7,7) | -0,04339 | -0,04175 | -0,00002 | 6,80275 | 7,00995 | -0,04318 | 6,97138 | 6,96205 | 6,69061 | 5,90496 |
| h(7,13) | -0,08095 | -0,0779 | -0,00003 | **12,69295** | **13,07957** | -0,08057 | **13,00759** | **12,99019** | **12,48372** | **11,01782** |
| h(7,23) | -0,00015 | -0,00015 | 0 | 0,02368 | 0,02441 | -0,00015 | 0,02427 | 0,02424 | 0,02329 | 0,02056 |
| h(7,29) | 0,04655 | 0,0448 | 0,00002 | -7,29945 | -7,52179 | 0,04633 | -7,48039 | -7,47039 | -7,17913 | -6,33611 |
| h(7,35) | 0,00005 | 0,00005 | 0 | -0,00809 | -0,00833 | 0,00005 | -0,00829 | -0,00828 | -0,00795 | -0,00702 |
| h(7,44) | 0,00001 | 0,00001 | 0 | -0,00147 | -0,00152 | 0,00001 | -0,00151 | -0,00151 | -0,00145 | -0,00128 |
| h(7,52) | 0 | 0 | 0 | -0,00006 | -0,00006 | 0 | -0,00006 | -0,00006 | -0,00006 | -0,00005 |
| h(7,60) | 0,03272 | 0,03149 | 0,00001 | -5,13072 | -5,287 | 0,03257 | -5,2579 | -5,25087 | -5,04614 | -4,4536 |
| h(7,66) | 0,00002 | 0,00002 | 0 | -0,00274 | -0,00283 | 0,00002 | -0,00281 | -0,00281 | -0,0027 | -0,00238 |
| h(7,70) | 0 | 0 | 0 | 0 | 0 | 0 | 0 | 0 | 0 | 0 |
| h(7,71) | -0,00001 | -0,00001 | 0 | 0,00217 | 0,00224 | -0,00001 | 0,00223 | 0,00223 | 0,00214 | 0,00189 |
| g(8,7) | 0,04339 | 0,04175 | 0,00002 | -6,80275 | -7,00995 | 0,04318 | 0,04848 | -6,96205 | -6,69061 | 0,04107 |
| g(8,23) | 0,05769 | 0,05552 | 0,00002 | -9,0457 | -9,32123 | 0,05742 | 0,06447 | -9,25753 | -8,89659 | 0,05461 |
| g(8,35) | -0,0197 | -0,01895 | -0,00001 | 3,08849 | 3,18256 | -0,0196 | -0,02201 | 3,16081 | 3,03758 | -0,01864 |
| g(8,44) | -0,00358 | -0,00345 | 0 | 0,56189 | 0,579 | -0,00357 | -0,004 | 0,57504 | 0,55262 | -0,00339 |
| g(8,52) | -0,00015 | -0,00014 | 0 | 0,02336 | 0,02407 | -0,00015 | -0,00017 | 0,0239 | 0,02297 | -0,00014 |
| g(8,66) | -0,00668 | -0,00643 | 0 | 1,04769 | 1,0796 | -0,00665 | -0,00747 | 1,07223 | 1,03042 | -0,00632 |
| g(8,71) | 0,0053 | 0,0051 | 0 | -0,83067 | -0,85597 | 0,00527 | 0,00592 | -0,85012 | -0,81698 | 0,00501 |
| h(8,8) | -0,06292 | -0,06055 | -0,00002 | 9,86541 | **10,16591** | -0,06262 | -0,07031 | **10,09644** | 9,70279 | -0,05956 |
| h(8,21) | -0,05728 | -0,05512 | -0,00002 | 8,98094 | 9,25449 | -0,057 | -0,06401 | 9,19125 | 8,8329 | -0,05422 |
| h(8,34) | 0,02065 | 0,01987 | 0,00001 | -3,23712 | -3,33572 | 0,02055 | 0,02307 | -3,31293 | -3,18376 | 0,01954 |
| h(8,45) | 0,0217 | 0,02088 | 0,00001 | -3,40266 | -3,5063 | 0,0216 | 0,02425 | -3,48234 | -3,34657 | 0,02054 |
| h(8,51) | 0,00012 | 0,00012 | 0 | -0,01877 | -0,01934 | 0,00012 | 0,00013 | -0,01921 | -0,01846 | 0,00011 |
| h(8,67) | 0,00669 | 0,00644 | 0 | -1,04909 | -1,08105 | 0,00666 | 0,00748 | -1,07366 | -1,0318 | 0,00633 |
| h(8,71) | -0,00527 | -0,00507 | 0 | 0,82572 | 0,85087 | -0,00524 | -0,00588 | 0,84506 | 0,81211 | -0,00498 |
| g(9,4) | -0,00003 | -0,00003 | 0 | 0,00606 | 0,00263 | -0,00003 | -0,00003 | -0,00003 | -0,40321 | -0,00003 |
| g(9,5) | -0,00048 | -0,00046 | 0 | 0,10588 | 0,04595 | -0,00047 | -0,00053 | -0,00053 | -7,04965 | -0,00045 |
| g(9,16) | -0,00059 | -0,00056 | 0 | 0,13016 | 0,05649 | -0,00058 | -0,00065 | -0,00065 | -8,66583 | -0,00055 |
| g(9,32) | -0,00002 | -0,00002 | 0 | 0,00371 | 0,00161 | -0,00002 | -0,00002 | -0,00002 | -0,24692 | -0,00002 |
| g(9,46) | 0,00002 | 0,00002 | 0 | -0,00514 | -0,00223 | 0,00002 | 0,00003 | 0,00003 | 0,34208 | 0,00002 |
| g(9,47) | 0,00022 | 0,00021 | 0 | -0,04913 | -0,02132 | 0,00022 | 0,00025 | 0,00025 | 3,27109 | 0,00021 |
| h(9,4) | 0,00002 | 0,00001 | 0 | -0,00342 | -0,00148 | 0,00002 | 0,00002 | 0,00002 | 0,22767 | 0,00001 |
| h(9,9) | 0,00053 | 0,00051 | 0 | -0,11712 | -0,05083 | 0,00052 | 0,00059 | 0,00059 | 7,79771 | 0,0005 |
| h(9,11) | 0,00002 | 0,00002 | 0 | -0,00382 | -0,00166 | 0,00002 | 0,00002 | 0,00002 | 0,25426 | 0,00002 |
| h(9,26) | 0 | 0 | 0 | 0,00024 | 0,0001 | 0 | 0 | 0 | -0,01601 | 0 |
| h(9,37) | 0 | 0 | 0 | 0,00001 | 0 | 0 | 0 | 0 | -0,00045 | 0 |
| h(9,56) | -0,00001 | -0,00001 | 0 | 0,002 | 0,00087 | -0,00001 | -0,00001 | -0,00001 | -0,133 | -0,00001 |
| h(9,62) | -0,00001 | 0 | 0 | 0,00115 | 0,0005 | -0,00001 | -0,00001 | -0,00001 | -0,0763 | 0 |
| h(9,72) | 0,00035 | 0,00033 | 0 | -0,07676 | -0,03332 | 0,00034 | 0,00039 | 0,00039 | 5,11091 | 0,00033 |

| | | | | | | | | | | |
|---|---|---|---|---|---|---|---|---|---|---|
| g(10,7) | 0 | 0 | 0 | 0 | 0 | 0 | 0 | 0 | 0 | -5,94603 |
| g(10,13) | 0 | 0 | 0 | 0 | 0 | 0 | 0 | 0 | 0 | **-11,12357** |
| g(10,29) | 0 | 0 | 0 | 0 | 0 | 0 | 0 | 0 | 0 | 6,39693 |
| g(10,60) | 0 | 0 | 0 | 0 | 0 | 0 | 0 | 0 | 0 | 4,49635 |
| g(10,70) | 0 | 0 | 0 | 0 | 0 | 0 | 0 | 0 | 0 | 0 |
| h(10,10) | 0 | 0 | 0 | 0 | 0 | 0 | 0 | 0 | 0 | 3,60842 |
| h(10,55) | 0 | 0 | 0 | 0 | 0 | 0 | 0 | 0 | 0 | 2,99573 |

In bold the kinetics order sensibilities with absolute value ≥ 10

**Table S2.6.** Logarithmic gains sensitivities for the fluxes.

| | $V_1$ | $V_2$ | $V_3$ | $V_4$ | $V_5$ | $V_6$ | $V_7$ | $V_8$ | $V_9$ | $V_{10}$ |
|---|---|---|---|---|---|---|---|---|---|---|
| $X_{11}$ | 0,00007 | 0,00006 | 0 | -0,01457 | -0,00632 | 0,00007 | 0,00007 | 0,00007 | -0,03557 | 0,00006 |
| $X_{12}$ | 0 | -1,01623 | 0 | 0 | 0 | 0 | 0 | 0 | 0 | 0 |
| $X_{13}$ | 0,00728 | 0,007 | 0 | -1,14109 | -1,17584 | 0,00724 | -1,16937 | -1,16781 | -1,12228 | 0,00951 |
| $X_{14}$ | 0 | 0 | -1,00458 | 0 | 0 | 0 | 0 | 0 | 0 | 0 |
| $X_{15}$ | -0,00024 | -0,00023 | 0 | 0,19061 | -0,11313 | -0,00024 | -0,00027 | -0,00027 | -0,1012 | -0,00023 |
| $X_{16}$ | 0,0042 | 0,00404 | 0 | -0,66223 | -0,6735 | 0,00418 | 0,00469 | 0,00468 | 0,35716 | 0,00397 |
| $X_{17}$ | 0,03347 | 0,03221 | 0,00001 | 0,0365 | 0,03761 | 0,03331 | 0,0374 | 0,03735 | 0,0359 | 0,03168 |
| $X_{18}$ | 0,51887 | 0,4993 | 0,00021 | 0,5658 | 0,58304 | 0,51639 | 0,57983 | 0,57905 | 0,55648 | 0,49113 |
| $X_{19}$ | 0 | -0,00199 | 0 | 0 | 0 | 0 | 0 | 0 | 0 | 0 |
| $X_{20}$ | 0 | 0 | 1,00416 | 0 | 0 | -1,05103 | 0 | 0 | 0 | 0 |
| $X_{21}$ | 0 | 0 | 0 | 0 | 0 | 0 | 0 | -1,00547 | 0 | 0 |
| $X_{22}$ | -0,003 | 0 | 0 | 0 | 0 | 0 | 0 | 0 | 0 | 0 |
| $X_{23}$ | -0,00621 | -0,00597 | 0 | 0,97312 | 1,00277 | -0,00618 | -0,00957 | 0,99591 | 0,95708 | -0,00811 |
| $X_{24}$ | 1,04877 | 1,00921 | 0,00042 | 1,14364 | 1,17848 | 1,04377 | 1,17199 | 1,17042 | 1,12479 | 0,99271 |
| $X_{25}$ | 0 | 0 | 0,01071 | 0 | 0 | -0,01121 | 0 | 0 | 0 | 0 |
| $X_{26}$ | 0,00007 | 0,00006 | 0 | -0,01457 | -0,00632 | 0,00007 | 0,00007 | 0,00007 | -0,03557 | 0,00006 |
| $X_{27}$ | -1,05606 | 0 | 0 | 0 | 0 | 0 | 0 | 0 | 0 | 0 |
| $X_{28}$ | 0 | -1,01623 | 0 | 0 | 0 | 0 | 0 | 0 | 0 | 0 |
| $X_{29}$ | 0,00728 | 0,007 | 0 | -1,14109 | -1,17584 | 0,00724 | -1,16937 | -1,16781 | -1,12228 | 0,00951 |
| $X_{30}$ | 0 | 0 | -1,00458 | 0 | 0 | 0 | 0 | 0 | 0 | 0 |
| $X_{31}$ | -0,00024 | -0,00023 | 0 | 0,19061 | -0,11313 | -0,00024 | -0,00027 | -0,00027 | -0,1012 | -0,00023 |
| $X_{32}$ | 0,0042 | 0,00404 | 0 | -0,66223 | -0,6735 | 0,00418 | 0,00469 | 0,00468 | 0,35716 | 0,00397 |
| $X_{33}$ | 0 | 0 | 1,00416 | 0 | 0 | -1,05103 | 0 | 0 | 0 | 0 |
| $X_{34}$ | 0 | 0 | 0 | 0 | 0 | 0 | 0 | -1,00547 | 0 | 0 |
| $X_{35}$ | -0,00621 | -0,00597 | 0 | 0,97312 | 1,00277 | -0,00618 | -0,00957 | 0,99591 | 0,95708 | -0,00811 |
| $X_{36}$ | 1,04877 | 1,00921 | 0,00042 | 1,14364 | 1,17848 | 1,04377 | 1,17199 | 1,17042 | 1,12479 | 0,99271 |
| $X_{37}$ | 0 | 0 | 0 | 0,00004 | 0,00002 | 0 | 0 | 0 | 0,0001 | 0 |
| $X_{38}$ | 1 | 0 | 0 | 0 | 0 | 0 | 0 | 0 | 0 | 0 |
| $X_{39}$ | 0 | 1 | 0 | 0 | 0 | 0 | 0 | 0 | 0 | 0 |
| $X_{40}$ | 0 | 0 | 1 | 0 | 0 | 0 | 0 | 0 | 0 | 0 |
| $X_{41}$ | -0,00056 | -0,00054 | 0 | 0,4438 | -0,26341 | -0,00056 | -0,00062 | -0,00062 | -0,23564 | -0,00053 |
| $X_{42}$ | 0,00078 | 0,00075 | 0 | -0,62088 | 0,36851 | 0,00078 | 0,00087 | 0,00087 | 0,32966 | 0,00074 |
| $X_{43}$ | 0 | 0 | -0,95541 | 0 | 0 | 1 | 0 | 0 | 0 | 0 |
| $X_{44}$ | 0,00616 | 0,00593 | 0 | -0,96653 | -0,99597 | 0,00613 | 0,00951 | -0,98917 | -0,9506 | 0,00805 |
| $X_{45}$ | 0 | 0 | 0 | 0 | 0 | 0 | 0 | 1 | 0 | 0 |
| $X_{46}$ | -0,00022 | -0,00021 | 0 | 0,0343 | 0,03489 | -0,00022 | -0,00024 | -0,00024 | -0,0185 | -0,00021 |
| $X_{47}$ | -0,00391 | -0,00376 | 0 | 0,61746 | 0,62797 | -0,00389 | -0,00437 | -0,00437 | -0,33301 | -0,0037 |
| $X_{48}$ | -0,02532 | -0,02437 | -0,00001 | -0,02761 | -0,02846 | -0,0252 | -0,0283 | -0,02826 | -0,02716 | -0,02397 |
| $X_{49}$ | -0,51887 | -0,4993 | -0,00021 | -0,5658 | -0,58304 | -0,51639 | -0,57983 | -0,57905 | -0,55648 | -0,49113 |
| $X_{50}$ | 0 | 0,00199 | 0 | 0 | 0 | 0 | 0 | 0 | 0 | 0 |
| $X_{51}$ | 0 | 0 | 0 | 0 | 0 | 0 | 0 | 0,00437 | 0 | 0 |
| $X_{52}$ | 0,00003 | 0,00003 | 0 | -0,00527 | -0,00543 | 0,00003 | 0,00005 | -0,00539 | -0,00518 | 0,00004 |
| $X_{53}$ | 0,00023 | 0 | 0 | 0 | 0 | 0 | 0 | 0 | 0 | 0 |
| $X_{54}$ | 0 | 0 | -0,01071 | 0 | 0 | 0,01121 | 0 | 0 | 0 | 0 |
| $X_{55}$ | 0 | 0 | 0 | 0 | 0 | 0 | 0 | 0 | 0 | -1 |
| $X_{56}$ | -0,00007 | -0,00006 | 0 | 0,01452 | 0,0063 | -0,00007 | -0,00007 | -0,00007 | 0,03546 | -0,00006 |
| $X_{57}$ | 0,00311 | 0,00299 | 0 | 0,00339 | 0,00349 | 0,00309 | 0,00347 | 0,00347 | 0,00333 | 0,00294 |
| $X_{58}$ | 0,00227 | 0,00218 | 0 | -0,50398 | -0,21873 | 0,00226 | 0,00253 | 0,00253 | -0,21535 | 0,00215 |

| | | | | | | | | | | |
|---|---|---|---|---|---|---|---|---|---|---|
| $X_{59}$ | 0,00313 | 0,00301 | 0 | 0,00341 | 0,00352 | 0,00312 | 0,0035 | 0,00349 | 0,00336 | 0,00296 |
| $X_{60}$ | -0,00616 | -0,00593 | 0 | 0,96653 | 0,99597 | -0,00613 | 0,99049 | 0,98917 | 0,9506 | -0,00805 |
| $X_{61}$ | 0 | 0 | 0,99098 | 0 | 0 | 0 | 0 | 0 | 0 | 0 |
| $X_{62}$ | -0,00006 | -0,00006 | 0 | 0,01445 | 0,00627 | -0,00006 | -0,00007 | -0,00007 | 0,03527 | -0,00006 |
| $X_{63}$ | 0 | 0 | -0,99114 | 0 | 0 | 0 | 0 | 0 | 0 | 0 |
| $X_{64}$ | -0,00815 | -0,00784 | 0 | -0,00889 | -0,00916 | -0,00811 | -0,00911 | -0,00909 | -0,00874 | -0,00771 |
| $X_{65}$ | 0,00277 | 0 | 0 | 0 | 0 | 0 | 0 | 0 | 0 | 0 |
| $X_{66}$ | 0,00088 | 0,00084 | 0 | -0,13748 | -0,14166 | 0,00087 | 0,00135 | -0,1407 | -0,13521 | 0,00115 |
| $X_{67}$ | 0 | 0 | 0 | 0 | 0 | 0 | 0 | 0,14224 | 0 | 0 |
| $X_{68}$ | -1,05606 | 0 | 0 | 0 | 0 | 0 | 0 | 0 | 0 | 0 |
| $X_{69}$ | 0 | 0 | 0,95541 | 0 | 0 | 0 | 0 | 0 | 0 | 0 |
| $X_{70}$ | -0,04327 | -0,04164 | -0,00002 | 6,78493 | 6,9916 | -0,04307 | 6,95312 | 6,94382 | 6,67309 | -0,05653 |
| $X_{71}$ | -0,00091 | -0,00088 | 0 | 0,14274 | 0,14709 | -0,00091 | -0,0014 | -0,00052 | 0,14039 | -0,00119 |
| $X_{72}$ | -0,00007 | -0,00006 | 0 | 0,01449 | 0,00629 | -0,00006 | -0,00007 | -0,00007 | -0,96463 | -0,00006 |

**Table S2.7.** Rate constants sensitivities for the fluxes

| | $V_1$ | $V_2$ | $V_3$ | $V_4$ | $V_5$ | $V_6$ | $V_7$ | $V_8$ | $V_9$ | $V_{10}$ |
|---|---|---|---|---|---|---|---|---|---|---|
| $\alpha_1$ | 1,04877 | 1,00921 | 0,00042 | 1,14364 | 1,17848 | 1,04377 | 1,17199 | 1,17042 | 1,12479 | 0,99271 |
| $\alpha_2$ | -0,00729 | 1,00921 | 1,00458 | 1,14364 | 1,17848 | -0,00726 | 1,17199 | 1,17042 | 1,12479 | 0,99271 |
| $\alpha_3$ | -0,00729 | -0,00702 | 1,00458 | 1,14364 | 1,17848 | -0,00726 | 1,17199 | 1,17042 | 1,12479 | 0,99271 |
| $\alpha_4$ | -0,00525 | -0,00505 | 0 | 1,16627 | 0,50617 | -0,00522 | -0,00586 | -0,00585 | 0,49834 | -0,00497 |
| $\alpha_5$ | -0,00324 | -0,00312 | 0 | 0,24071 | 0,78889 | -0,00322 | -0,00362 | -0,00362 | 0,74111 | -0,00307 |
| $\alpha_6$ | -0,00729 | 1,00921 | 0,00042 | 1,14364 | 1,17848 | 1,04377 | 1,17199 | 1,17042 | 1,12479 | 0,99271 |
| $\alpha_7$ | -0,00729 | -0,00702 | 0 | 1,14364 | 1,17848 | -0,00726 | 1,17199 | 1,17042 | 1,12479 | 0,99271 |
| $\alpha_8$ | -0,00622 | -0,00599 | 0 | 0,97568 | 1,0054 | -0,00619 | -0,00695 | 0,99853 | 0,9596 | -0,00589 |
| $\alpha_9$ | 0,00007 | 0,00006 | 0 | -0,01493 | -0,00648 | 0,00007 | 0,00008 | 0,00007 | 0,99379 | 0,00006 |
| $\alpha_{10}$ | 0 | 0 | 0 | 0 | 0 | 0 | 0 | 0 | 0 | 1 |

**Table S2.8.** Kinetic orders sensitivities for the fluxes

| | $V_1$ | $V_2$ | $V_3$ | $V_4$ | $V_5$ | $V_6$ | $V_7$ | $V_8$ | $V_9$ | $V_{10}$ |
|---|---|---|---|---|---|---|---|---|---|---|
| g(1,4) | 0,02434 | 0,02343 | 0,00001 | 0,02655 | 0,02735 | 0,02423 | 0,0272 | 0,02717 | 0,02611 | 0,02304 |
| g(1,5) | 0,02382 | 0,02292 | 0,00001 | 0,02597 | 0,02676 | 0,0237 | 0,02661 | 0,02658 | 0,02554 | 0,02254 |
| g(1,17) | 0,03287 | 0,03163 | 0,00001 | 0,03585 | 0,03694 | 0,03272 | 0,03673 | 0,03669 | 0,03525 | 0,03112 |
| g(1,18) | -1,65736 | -1,59485 | -0,00066 | -1,80728 | -1,86233 | -1,64945 | -1,85208 | -1,8496 | -1,77749 | -1,56877 |
| g(1,24) | **-10,98945** | **-10,57498** | -0,00436 | **-11,98356** | **-12,34857** | **-10,93703** | **-12,28062** | **-12,26419** | **-11,78602** | **-10,40205** |
| g(1,36) | 4,8329 | 4,65063 | 0,00192 | 5,27009 | 5,43061 | 4,80985 | 5,40073 | 5,3935 | 5,18322 | 4,57458 |
| g(1,48) | 0,0509 | 0,04898 | 0,00002 | 0,0555 | 0,05719 | 0,05066 | 0,05688 | 0,0568 | 0,05459 | 0,04818 |
| g(1,49) | 1,65736 | 1,59485 | 0,00066 | 1,80728 | 1,86233 | 1,64945 | 1,85208 | 1,8496 | 1,77749 | 1,56877 |
| g(1,57) | -0,00848 | -0,00816 | 0 | -0,00924 | -0,00952 | -0,00844 | -0,00947 | -0,00946 | -0,00909 | -0,00802 |
| g(1,59) | -0,00786 | -0,00756 | 0 | -0,00857 | -0,00883 | -0,00782 | -0,00878 | -0,00877 | -0,00843 | -0,00744 |
| g(1,64) | 0,02572 | 0,02475 | 0,00001 | 0,02804 | 0,0289 | 0,02559 | 0,02874 | 0,0287 | 0,02758 | 0,02434 |
| h(1,1) | 5,94243 | 5,71831 | 0,00236 | 6,47998 | 6,67736 | 5,91408 | 6,64061 | 6,63173 | 6,37317 | 5,6248 |
| h(1,22) | 0,00632 | 0,00608 | 0 | 0,00689 | 0,0071 | 0,00629 | 0,00707 | 0,00706 | 0,00678 | 0,00598 |
| h(1,27) | -7,82554 | -7,53039 | -0,00311 | -8,53344 | -8,79336 | -7,78821 | -8,74497 | -8,73327 | -8,39277 | -7,40725 |
| h(1,38) | -3,07967 | -2,96352 | -0,00122 | -3,35826 | -3,46055 | -3,06498 | -3,44151 | -3,43691 | -3,30291 | -2,91506 |
| h(1,53) | -0,00176 | -0,00169 | 0 | -0,00191 | -0,00197 | -0,00175 | -0,00196 | -0,00196 | -0,00188 | -0,00166 |
| h(1,65) | -0,02203 | -0,02119 | -0,00001 | -0,02402 | -0,02475 | -0,02192 | -0,02461 | -0,02458 | -0,02362 | -0,02085 |
| h(1,68) | **11,66605** | **11,22606** | 0,00463 | **12,72137** | **13,10885** | **11,6104** | **13,03671** | **13,01927** | **12,51167** | **11,04248** |
| g(2,6) | 0,06237 | -8,62947 | -8,58981 | -9,77892 | **-10,07678** | 0,06207 | **-10,02132** | **-10,00792** | -9,61772 | -8,48836 |
| g(2,20) | 0,07788 | **-10,77566** | **-10,72613** | **-12,21098** | **-12,58291** | 0,07751 | **-12,51367** | **-12,49693** | **-12,00969** | **-10,59945** |
| g(2,25) | -0,00008 | 0,01069 | 0,01064 | 0,01211 | 0,01248 | -0,00008 | 0,01241 | 0,0124 | 0,01191 | 0,01051 |
| g(2,33) | -0,0536 | 7,4163 | 7,38221 | 8,40415 | 8,66013 | -0,05334 | 8,61247 | 8,60095 | 8,26561 | 7,29502 |
| g(2,43) | -0,04 | 5,53425 | 5,50882 | 6,27141 | 6,46244 | -0,03981 | 6,42687 | 6,41827 | 6,16803 | 5,44375 |
| g(2,54) | -0,00002 | 0,00295 | 0,00294 | 0,00335 | 0,00345 | -0,00002 | 0,00343 | 0,00343 | 0,00329 | 0,00291 |
| h(2,2) | -0,04413 | 6,1064 | 6,07834 | 6,91977 | 7,13055 | -0,04392 | 7,09131 | 7,08182 | 6,80571 | 6,00655 |
| h(2,12) | -0,06748 | 9,33701 | 9,2941 | **10,5807** | **10,90298** | -0,06716 | **10,84298** | **10,82847** | **10,40628** | 9,18432 |
| h(2,19) | 0,00002 | -0,00217 | -0,00216 | -0,00246 | -0,00254 | 0,00002 | -0,00253 | -0,00252 | -0,00242 | -0,00214 |
| h(2,28) | 0,05046 | -6,98145 | -6,94936 | -7,91137 | -8,15235 | 0,05022 | -8,10748 | -8,09664 | -7,78096 | -6,86728 |
| h(2,39) | 0,01362 | -1,88402 | -1,87536 | -2,13497 | -2,2 | 0,01355 | -2,18789 | -2,18496 | -2,09977 | -1,85321 |
| h(2,50) | 0,00001 | -0,00172 | -0,00171 | -0,00195 | -0,00201 | 0,00001 | -0,00199 | -0,00199 | -0,00191 | -0,00169 |
| g(3,2) | 0,04413 | 0,04247 | -6,07834 | -6,91977 | -7,13055 | 0,04392 | -7,09131 | -7,08182 | -6,80571 | -6,00655 |
| g(3,12) | 0,06748 | 0,06494 | -9,2941 | **-10,5807** | **-10,90298** | 0,06716 | **-10,84298** | **-10,82847** | **-10,40628** | -9,18432 |
| g(3,19) | -0,00002 | -0,00002 | 0,00216 | 0,00246 | 0,00254 | -0,00002 | 0,00253 | 0,00252 | 0,00242 | 0,00214 |
| g(3,28) | -0,05046 | -0,04855 | 6,94936 | 7,91137 | 8,15235 | -0,05022 | 8,10748 | 8,09664 | 7,78096 | 6,86728 |
| g(3,39) | -0,01362 | -0,0131 | 1,87536 | 2,13497 | 2,2 | -0,01355 | 2,18789 | 2,18496 | 2,09977 | 1,85321 |
| g(3,50) | -0,00001 | -0,00001 | 0,00171 | 0,00195 | 0,00201 | -0,00001 | 0,00199 | 0,00199 | 0,00191 | 0,00169 |
| h(3,3) | -0,04696 | -0,04519 | 6,46814 | 7,36354 | 7,58782 | -0,04674 | 7,54607 | 7,53597 | 7,24215 | 6,39174 |
| h(3,6) | -0,06234 | -0,05999 | 8,58623 | 9,77484 | **10,07257** | -0,06204 | **10,01714** | **10,00374** | 9,61371 | 8,48482 |
| h(3,14) | -0,07206 | -0,06934 | 9,92498 | **11,29891** | **11,64307** | -0,07172 | **11,579** | **11,56351** | **11,11266** | 9,80775 |
| h(3,30) | 0,03203 | 0,03082 | -4,41083 | -5,02143 | -5,17438 | 0,03187 | -5,1459 | -5,13902 | -4,93866 | -4,35873 |
| h(3,40) | 0,00785 | 0,00756 | -1,08175 | -1,2315 | -1,26901 | 0,00782 | -1,26203 | -1,26034 | -1,2112 | -1,06898 |
| h(3,61) | 0 | 0 | 0 | 0 | 0 | 0 | 0 | 0 | 0 | 0 |
| h(3,63) | 0,0547 | 0,05264 | -7,53355 | -8,57643 | -8,83766 | 0,05444 | -8,78903 | -8,77727 | -8,43506 | -7,44457 |
| h(3,69) | 0,03945 | 0,03796 | -5,43333 | -6,18547 | -6,37388 | 0,03926 | -6,3388 | -6,33032 | -6,08351 | -5,36915 |
| g(4,4) | -0,00244 | -0,00235 | 0 | 0,5431 | 0,23571 | -0,00243 | -0,00273 | -0,00273 | 0,23206 | -0,00231 |
| g(4,5) | 0,01494 | 0,01437 | 0,00001 | -3,32109 | -1,44138 | 0,01487 | 0,01669 | 0,01667 | -1,41908 | 0,01414 |
| g(4,8) | 0,03506 | 0,03374 | 0,00001 | -7,79595 | -3,38351 | 0,0349 | 0,03918 | 0,03913 | -3,33115 | 0,03319 |
| g(4,15) | 0,02275 | 0,02189 | 0,00001 | -5,05839 | -2,19539 | 0,02264 | 0,02542 | 0,02539 | -2,16141 | 0,02153 |
| g(4,21) | 0,03192 | 0,03072 | 0,00001 | -7,09701 | -3,08017 | 0,03177 | 0,03567 | 0,03562 | -3,0325 | 0,03021 |

| | | | | | | | | | | |
|---|---|---|---|---|---|---|---|---|---|---|
| g(4,31) | -0,00531 | -0,00511 | 0 | 1,18013 | 0,51219 | -0,00528 | -0,00593 | -0,00592 | 0,50426 | -0,00502 |
| g(4,34) | -0,01151 | -0,01107 | 0 | 2,55807 | 1,11023 | -0,01145 | -0,01286 | -0,01284 | 1,09305 | -0,01089 |
| g(4,41) | 0,00145 | 0,0014 | 0 | -0,32287 | -0,14013 | 0,00145 | 0,00162 | 0,00162 | -0,13796 | 0,00137 |
| g(4,42) | -0,00834 | -0,00802 | 0 | 1,85414 | 0,80472 | -0,0083 | -0,00932 | -0,00931 | 0,79226 | -0,00789 |
| g(4,45) | -0,01209 | -0,01164 | 0 | 2,68888 | 1,167 | -0,01204 | -0,01351 | -0,0135 | 1,14894 | -0,01145 |
| g(4,51) | -0,00007 | -0,00006 | 0 | 0,01483 | 0,00644 | -0,00007 | -0,00007 | -0,00007 | 0,00634 | -0,00006 |
| g(4,67) | -0,00373 | -0,00359 | 0 | 0,82903 | 0,35981 | -0,00371 | -0,00417 | -0,00416 | 0,35424 | -0,00353 |
| g(4,71) | 0,00293 | 0,00282 | 0 | -0,65251 | -0,2832 | 0,00292 | 0,00328 | 0,00328 | -0,27881 | 0,00278 |
| h(4,4) | -0,03806 | -0,03662 | -0,00002 | 8,4617 | 3,67246 | -0,03788 | -0,04253 | -0,04247 | 3,61562 | -0,03602 |
| h(4,5) | -0,00233 | -0,00225 | 0 | 0,51898 | 0,22524 | -0,00232 | -0,00261 | -0,0026 | 0,22176 | -0,00221 |
| h(4,9) | -0,00053 | -0,00051 | 0 | 0,11712 | 0,05083 | -0,00052 | -0,00059 | -0,00059 | 0,05004 | -0,0005 |
| h(4,11) | -0,00059 | -0,00057 | 0 | 0,13087 | 0,0568 | -0,00059 | -0,00066 | -0,00066 | 0,05592 | -0,00056 |
| h(4,15) | -0,00781 | -0,00752 | 0 | 1,73751 | 0,75409 | -0,00778 | -0,00873 | -0,00872 | 0,74242 | -0,0074 |
| h(4,16) | -0,02006 | -0,0193 | -0,00001 | 4,46044 | 1,93587 | -0,01997 | -0,02242 | -0,02239 | 1,90591 | -0,01899 |
| h(4,26) | 0,00004 | 0,00004 | 0 | -0,00824 | -0,00358 | 0,00004 | 0,00004 | 0,00004 | -0,00352 | 0,00004 |
| h(4,31) | 0,00182 | 0,00175 | 0 | -0,40536 | -0,17593 | 0,00181 | 0,00204 | 0,00203 | -0,17321 | 0,00173 |
| h(4,32) | -0,00057 | -0,00055 | 0 | 0,12709 | 0,05516 | -0,00057 | -0,00064 | -0,00064 | 0,05431 | -0,00054 |
| h(4,37) | 0 | 0 | 0 | -0,00023 | -0,0001 | 0 | 0 | 0 | -0,0001 | 0 |
| h(4,41) | 0,0112 | 0,01078 | 0 | -2,49052 | -1,08091 | 0,01115 | 0,01252 | 0,0125 | -1,06418 | 0,0106 |
| h(4,42) | -0,00781 | -0,00751 | 0 | 1,73606 | 0,75347 | -0,00777 | -0,00873 | -0,00871 | 0,7418 | -0,00739 |
| h(4,46) | 0,00079 | 0,00076 | 0 | -0,17608 | -0,07642 | 0,00079 | 0,00088 | 0,00088 | -0,07524 | 0,00075 |
| h(4,47) | 0,00757 | 0,00729 | 0 | -1,68368 | -0,73073 | 0,00754 | 0,00846 | 0,00845 | -0,71942 | 0,00717 |
| h(4,56) | 0,00031 | 0,0003 | 0 | -0,06846 | -0,02971 | 0,00031 | 0,00034 | 0,00034 | -0,02925 | 0,00029 |
| h(4,58) | -0,01201 | -0,01156 | 0 | 2,67024 | 1,15891 | -0,01195 | -0,01342 | -0,0134 | 1,14097 | -0,01137 |
| h(4,62) | 0,00018 | 0,00017 | 0 | -0,03927 | -0,01705 | 0,00018 | 0,0002 | 0,0002 | -0,01678 | 0,00017 |
| g(5,4) | 0,01497 | 0,0144 | 0,00001 | -1,11209 | -3,64473 | 0,0149 | 0,01673 | 0,01671 | -3,424 | 0,01417 |
| g(5,5) | -0,01111 | -0,01069 | 0 | 0,82545 | 2,7053 | -0,01106 | -0,01242 | -0,0124 | 2,54147 | -0,01052 |
| g(5,8) | 0,02786 | 0,02681 | 0,00001 | -2,06946 | -6,78239 | 0,02772 | 0,03113 | 0,03109 | -6,37164 | 0,02637 |
| g(5,15) | 0,00621 | 0,00597 | 0 | -0,46123 | -1,51161 | 0,00618 | 0,00694 | 0,00693 | -1,42007 | 0,00588 |
| g(5,21) | 0,02536 | 0,0244 | 0,00001 | -1,88393 | -6,17432 | 0,02524 | 0,02834 | 0,0283 | -5,8004 | 0,024 |
| g(5,31) | -0,00145 | -0,00139 | 0 | 0,1076 | 0,35266 | -0,00144 | -0,00162 | -0,00162 | 0,3313 | -0,00137 |
| g(5,34) | -0,00914 | -0,0088 | 0 | 0,67905 | 2,2255 | -0,0091 | -0,01021 | -0,0102 | 2,09072 | -0,00865 |
| g(5,41) | -0,0089 | -0,00856 | 0 | 0,66112 | 2,16672 | -0,00886 | -0,00994 | -0,00993 | 2,0355 | -0,00842 |
| g(5,42) | 0,0062 | 0,00597 | 0 | -0,46084 | -1,51035 | 0,00617 | 0,00693 | 0,00692 | -1,41888 | 0,00587 |
| g(5,45) | -0,00961 | -0,00925 | 0 | 0,71377 | 2,3393 | -0,00956 | -0,01074 | -0,01072 | 2,19763 | -0,00909 |
| g(5,51) | -0,00005 | -0,00005 | 0 | 0,00394 | 0,0129 | -0,00005 | -0,00006 | -0,00006 | 0,01212 | -0,00005 |
| g(5,67) | -0,00296 | -0,00285 | 0 | 0,22007 | 0,72124 | -0,00295 | -0,00331 | -0,00331 | 0,67756 | -0,0028 |
| g(5,71) | 0,00233 | 0,00224 | 0 | -0,17321 | -0,56768 | 0,00232 | 0,00261 | 0,0026 | -0,5333 | 0,00221 |
| h(5,4) | 0,0012 | 0,00115 | 0 | -0,08908 | -0,29194 | 0,00119 | 0,00134 | 0,00134 | -0,27426 | 0,00113 |
| h(5,5) | -0,02483 | -0,0239 | -0,00001 | 1,84481 | 6,04613 | -0,02471 | -0,02775 | -0,02771 | 5,67996 | -0,0235 |
| h(5,15) | -0,01807 | -0,01739 | -0,00001 | 1,34277 | 4,40074 | -0,01799 | -0,0202 | -0,02017 | 4,13423 | -0,01711 |
| h(5,16) | -0,01594 | -0,01534 | -0,00001 | 1,18404 | 3,88053 | -0,01586 | -0,01781 | -0,01779 | 3,64552 | -0,01509 |
| h(5,31) | 0,00422 | 0,00406 | 0 | -0,31327 | -1,0267 | 0,0042 | 0,00471 | 0,00471 | -0,96452 | 0,00399 |
| h(5,32) | -0,00045 | -0,00044 | 0 | 0,03374 | 0,11057 | -0,00045 | -0,00051 | -0,00051 | 0,10387 | -0,00043 |
| h(5,41) | -0,00115 | -0,00111 | 0 | 0,08571 | 0,28089 | -0,00115 | -0,00129 | -0,00129 | 0,26388 | -0,00109 |
| h(5,42) | 0,00663 | 0,00638 | 0 | -0,49219 | -1,61308 | 0,00659 | 0,0074 | 0,00739 | -1,51539 | 0,00627 |
| h(5,46) | 0,00063 | 0,00061 | 0 | -0,04674 | -0,15318 | 0,00063 | 0,0007 | 0,0007 | -0,14391 | 0,0006 |
| h(5,47) | 0,00602 | 0,00579 | 0 | -0,44694 | -1,46478 | 0,00599 | 0,00672 | 0,00671 | -1,37607 | 0,00569 |
| g(6,1) | 0,04133 | -5,71831 | -0,00236 | -6,47998 | -6,67736 | -5,91408 | -6,64061 | -6,63173 | -6,37317 | -5,6248 |
| g(6,22) | 0,00004 | -0,00608 | 0 | -0,00689 | -0,0071 | -0,00629 | -0,00707 | -0,00706 | -0,00678 | -0,00598 |
| g(6,27) | -0,05442 | 7,53039 | 0,00311 | 8,53344 | 8,79336 | 7,78821 | 8,74497 | 8,73327 | 8,39277 | 7,40725 |
| g(6,38) | -0,02142 | 2,96352 | 0,00122 | 3,35826 | 3,46055 | 3,06498 | 3,44151 | 3,43691 | 3,30291 | 2,91506 |
| g(6,53) | -0,00001 | 0,00169 | 0 | 0,00191 | 0,00197 | 0,00175 | 0,00196 | 0,00196 | 0,00188 | 0,00166 |
| g(6,65) | -0,00015 | 0,02119 | 0,00001 | 0,02402 | 0,02475 | 0,02192 | 0,02461 | 0,02458 | 0,02362 | 0,02085 |

| | | | | | | | | | | |
|---|---|---|---|---|---|---|---|---|---|---|
| g(6,68) | 0,08113 | **-11,22606** | -0,00463 | **-12,72137** | **-13,10885** | **-11,6104** | **-13,03671** | **-13,01927** | **-12,51167** | **-11,04248** |
| h(6,6) | -0,06237 | 8,62947 | 0,00356 | 9,77892 | **10,07678** | 8,92492 | **10,02132** | **10,00792** | 9,61772 | 8,48836 |
| h(6,20) | -0,07788 | **10,77566** | 0,00444 | **12,21098** | **12,58291** | **11,14458** | **12,51367** | **12,49693** | **12,00969** | **10,59945** |
| h(6,25) | 0,00008 | -0,01069 | 0 | -0,01211 | -0,01248 | -0,01106 | -0,01241 | -0,0124 | -0,01191 | -0,01051 |
| h(6,33) | 0,0536 | -7,4163 | -0,00306 | -8,40415 | -8,66013 | -7,67021 | -8,61247 | -8,60095 | -8,26561 | -7,29502 |
| h(6,43) | 0,04 | -5,53425 | -0,00228 | -6,27141 | -6,46244 | -5,72373 | -6,42687 | -6,41827 | -6,16803 | -5,44375 |
| h(6,54) | 0,00002 | -0,00295 | 0 | -0,00335 | -0,00345 | -0,00305 | -0,00343 | -0,00343 | -0,00329 | -0,00291 |
| g(7,3) | 0,04696 | 0,04519 | 0,00002 | -7,36354 | -7,58782 | 0,04674 | -7,54607 | -7,53597 | -7,24215 | -6,39174 |
| g(7,6) | 0,06234 | 0,05999 | 0,00002 | **-9,77484** | **-10,07257** | 0,06204 | **-10,01714** | **-10,00374** | -9,61371 | -8,48482 |
| g(7,14) | 0,07206 | 0,06934 | 0,00003 | **-11,29891** | **-11,64307** | 0,07172 | **-11,579** | **-11,56351** | **-11,11266** | -9,80775 |
| g(7,30) | -0,03203 | -0,03082 | -0,00001 | 5,02143 | 5,17438 | -0,03187 | 5,1459 | 5,13902 | 4,93866 | 4,35873 |
| g(7,40) | -0,00785 | -0,00756 | 0 | 1,2315 | 1,26901 | -0,00782 | 1,26203 | 1,26034 | 1,2112 | 1,06898 |
| g(7,61) | 0 | 0 | 0 | 0 | 0 | 0 | 0 | 0 | 0 | 0 |
| g(7,63) | -0,0547 | -0,05264 | -0,00002 | 8,57643 | 8,83766 | -0,05444 | 8,78903 | 8,77727 | 8,43506 | 7,44457 |
| g(7,69) | -0,03945 | -0,03796 | -0,00002 | 6,18547 | 6,37388 | -0,03926 | 6,3388 | 6,33032 | 6,08351 | 5,36915 |
| h(7,7) | -0,04339 | -0,04175 | -0,00002 | 6,80275 | 7,00995 | -0,04318 | 6,97138 | 6,96205 | 6,69061 | 5,90496 |
| h(7,13) | -0,08095 | -0,0779 | -0,00003 | **12,69295** | **13,07957** | -0,08057 | **13,00759** | **12,99019** | **12,48372** | **11,01782** |
| h(7,23) | -0,00015 | -0,00015 | 0 | 0,02368 | 0,02441 | -0,00015 | 0,02427 | 0,02424 | 0,02329 | 0,02056 |
| h(7,29) | 0,04655 | 0,0448 | 0,00002 | -7,29945 | -7,52179 | 0,04633 | -7,48039 | -7,47039 | -7,17913 | -6,33611 |
| h(7,35) | 0,00005 | 0,00005 | 0 | -0,00809 | -0,00833 | 0,00005 | -0,00829 | -0,00828 | -0,00795 | -0,00702 |
| h(7,44) | 0,00001 | 0,00001 | 0 | -0,00147 | -0,00152 | 0,00001 | -0,00151 | -0,00151 | -0,00145 | -0,00128 |
| h(7,52) | 0 | 0 | 0 | -0,00006 | -0,00006 | 0 | -0,00006 | -0,00006 | -0,00006 | -0,00005 |
| h(7,60) | 0,03272 | 0,03149 | 0,00001 | -5,13072 | -5,287 | 0,03257 | -5,2579 | -5,25087 | -5,04614 | -4,4536 |
| h(7,66) | 0,00002 | 0,00002 | 0 | -0,00274 | -0,00283 | 0,00002 | -0,00281 | -0,00281 | -0,0027 | -0,00238 |
| h(7,70) | 0 | 0 | 0 | 0 | 0 | 0 | 0 | 0 | 0 | 0 |
| h(7,71) | -0,00001 | -0,00001 | 0 | 0,00217 | 0,00224 | -0,00001 | 0,00223 | 0,00223 | 0,00214 | 0,00189 |
| g(8,7) | 0,04339 | 0,04175 | 0,00002 | -6,80275 | -7,00995 | 0,04318 | 0,04848 | -6,96205 | -6,69061 | 0,04107 |
| g(8,23) | 0,05769 | 0,05552 | 0,00002 | -9,0457 | -9,32123 | 0,05742 | 0,06447 | -9,25753 | -8,89659 | 0,05461 |
| g(8,35) | -0,0197 | -0,01895 | -0,00001 | 3,08849 | 3,18256 | -0,0196 | -0,02201 | 3,16081 | 3,03758 | -0,01864 |
| g(8,44) | -0,00358 | -0,00345 | 0 | 0,56189 | 0,579 | -0,00357 | -0,004 | 0,57504 | 0,55262 | -0,00339 |
| g(8,52) | -0,00015 | -0,00014 | 0 | 0,02336 | 0,02407 | -0,00015 | -0,00017 | 0,0239 | 0,02297 | -0,00014 |
| g(8,66) | -0,00668 | -0,00643 | 0 | 1,04769 | 1,0796 | -0,00665 | -0,00747 | 1,07223 | 1,03042 | -0,00632 |
| g(8,71) | 0,0053 | 0,0051 | 0 | -0,83067 | -0,85597 | 0,00527 | 0,00592 | -0,85012 | -0,81698 | 0,00501 |
| h(8,8) | -0,06292 | -0,06055 | -0,00002 | 9,86541 | **10,16591** | -0,06262 | -0,07031 | **10,09644** | 9,70279 | -0,05956 |
| h(8,21) | -0,05728 | -0,05512 | -0,00002 | 8,98094 | 9,25449 | -0,057 | -0,06401 | 9,19125 | 8,8329 | -0,05422 |
| h(8,34) | 0,02065 | 0,01987 | 0,00001 | -3,23712 | -3,33572 | 0,02055 | 0,02307 | -3,31293 | -3,18376 | 0,01954 |
| h(8,45) | 0,0217 | 0,02088 | 0,00001 | -3,40266 | -3,5063 | 0,0216 | 0,02425 | -3,48234 | -3,34657 | 0,02054 |
| h(8,51) | 0,00012 | 0,00012 | 0 | -0,01877 | -0,01934 | 0,00012 | 0,00013 | -0,01921 | -0,01846 | 0,00011 |
| h(8,67) | 0,00669 | 0,00644 | 0 | -1,04909 | -1,08105 | 0,00666 | 0,00748 | -1,07366 | -1,0318 | 0,00633 |
| h(8,71) | -0,00527 | -0,00507 | 0 | 0,82572 | 0,85087 | -0,00524 | -0,00588 | 0,84506 | 0,81211 | -0,00498 |
| g(9,4) | -0,00003 | -0,00003 | 0 | 0,00606 | 0,00263 | -0,00003 | -0,00003 | -0,00003 | -0,40321 | -0,00003 |
| g(9,5) | -0,00048 | -0,00046 | 0 | 0,10588 | 0,04595 | -0,00047 | -0,00053 | -0,00053 | -7,04965 | -0,00045 |
| g(9,16) | -0,00059 | -0,00056 | 0 | 0,13016 | 0,05649 | -0,00058 | -0,00065 | -0,00065 | -8,66583 | -0,00055 |
| g(9,32) | -0,00002 | -0,00002 | 0 | 0,00371 | 0,00161 | -0,00002 | -0,00002 | -0,00002 | -0,24692 | -0,00002 |
| g(9,46) | 0,00002 | 0,00002 | 0 | -0,00514 | -0,00223 | 0,00002 | 0,00003 | 0,00003 | 0,34208 | 0,00002 |
| g(9,47) | 0,00022 | 0,00021 | 0 | -0,04913 | -0,02132 | 0,00022 | 0,00025 | 0,00025 | 3,27109 | 0,00021 |
| h(9,4) | 0,00002 | 0,00001 | 0 | -0,00342 | -0,00148 | 0,00002 | 0,00002 | 0,00002 | 0,22767 | 0,00001 |
| h(9,9) | 0,00053 | 0,00051 | 0 | -0,11712 | -0,05083 | 0,00052 | 0,00059 | 0,00059 | 7,79771 | 0,0005 |
| h(9,11) | 0,00002 | 0,00002 | 0 | -0,00382 | -0,00166 | 0,00002 | 0,00002 | 0,00002 | 0,25426 | 0,00002 |
| h(9,26) | 0 | 0 | 0 | 0,00024 | 0,0001 | 0 | 0 | 0 | -0,01601 | 0 |
| h(9,37) | 0 | 0 | 0 | 0,00001 | 0 | 0 | 0 | 0 | -0,00045 | 0 |
| h(9,56) | -0,00001 | -0,00001 | 0 | 0,002 | 0,00087 | -0,00001 | -0,00001 | -0,00001 | -0,133 | -0,00001 |
| h(9,62) | -0,00001 | 0 | 0 | 0,00115 | 0,0005 | -0,00001 | -0,00001 | -0,00001 | -0,0763 | 0 |
| h(9,72) | 0,00035 | 0,00033 | 0 | -0,07676 | -0,03332 | 0,00034 | 0,00039 | 0,00039 | 5,11091 | 0,00033 |

| | | | | | | | | | | |
|---|---|---|---|---|---|---|---|---|---|---|
| g(10,7) | 0 | 0 | 0 | 0 | 0 | 0 | 0 | 0 | 0 | -5,94603 |
| g(10,13) | 0 | 0 | 0 | 0 | 0 | 0 | 0 | 0 | 0 | **-11,12357** |
| g(10,29) | 0 | 0 | 0 | 0 | 0 | 0 | 0 | 0 | 0 | 6,39693 |
| g(10,60) | 0 | 0 | 0 | 0 | 0 | 0 | 0 | 0 | 0 | 4,49635 |
| g(10,70) | 0 | 0 | 0 | 0 | 0 | 0 | 0 | 0 | 0 | 0 |
| h(10,10) | 0 | 0 | 0 | 0 | 0 | 0 | 0 | 0 | 0 | 3,60842 |
| h(10,55) | 0 | 0 | 0 | 0 | 0 | 0 | 0 | 0 | 0 | 2,99573 |

In bold the kinetics order sensibilities with absolute value ≥ 10

### References

[1] T.C. Ni, and M.A. Savageau, Model assessment and refinement using strategies from biochemical systems theory: application to metabolism in human red blood cells. J Theor Biol 179 (1996) 329-68.

[2] F. Shiraishi, and M.A. Savageau, The tricarboxylic acid cycle in *Dictyostelium discoideum*. II. Evaluation of model consistency and robustness. J Biol Chem 267 (1992) 22919-25.

**Supplementary File 3**


| | | Pag |
|---|---|---|
| S3 | Model optimization. ……………………………………………………...………… | 3 |
| S3.1 | Flux maximization thought IspA ($X_{11}$). ……………………………………………… | 3 |
| S3.2 | Steady state restrictions. …………………………………………………...…… | 3 |
| S3.3 | External model metabolites restrictions. …………….…………….………… | 4 |
| S3.4 | Independent variables boundaries. ……………………………………………… | 5 |
| S3.5 | Enzymes modification restriction. ………………………………………………. | 6 |
| S3.6 | Total enzyme restriction: The cellular economy factor. ………………………… | 7 |
| Table S3.1a | Optimization of the *K*cat's allowing modifications for the Km's, $K_{0,5}$'s, and Ki's. Assuming no relationship between IspA kcat's $X_{26}$ and $X_{32}$.…………………...… | 8 |
| Table S3.1b | Specific constants ($K_{cat}$ /Km) ratios for data from Table 3.1a. ……………………. | 13 |
| Table S3.2a | Optimization of the *K*cat's allowing modifications for the Km's, $K_{0,5}$'s, and Ki's. Setting a IspA kcat_11 $X_{26}/X_{32}$ ratio equal 2.22. …………………………………. | 15 |
| Table S3.2b | Specific constants ($K_{cat}$ /Km) ratios for data from Table 3.2a. ……………………. | 19 |
| Table S3.3a | Optimization of the *k*cat's allowing modifications in the Km's, Kd's, Ki's, and $K_{0,5}$'s. For the IspA kcat is allowed only the modulation of kcat_16 ($X_{32}$) while the kcat_11 ($X_{26}$) is anchored in their basal value.………………………………… | 21 |
| Table S3.3b | Specific constants ($K_{cat}$ /Km) ratios for data from Table 3.3a. ……………………. | 23 |
| Table S3.4a | Optimization of the *K*cat's allowing modifications in the Km's, Kd's, Ki's, and $K_{0,5}$'s. No relationship between kcat_11 ($X_{26}$) and kcat_16 ($X_{32}$). ………………… | 24 |
| Table S3.4b | Specific constants ($K_{cat}$ /Km) ratios for data from Table 3.4a. …………………….. | 26 |
| Table S3.5a | Optimization of the *K*cat's allowing modifications in the Km's, Kd's, Ki's, and $K_{0,5}$'s. The$K_{cat}$ IspA basal ratio of kcat_11/ kcat_16 = 2.22 ($X_{26}/X_{32}$) was imposed for all the experiments. ……………………………………………………………….. | 27 |
| Table S3.5b | Specific constants ($K_{cat}$ /Km) ratios for data from Table 3.7a. …………………… | 29 |
| Table S3.6a | Optimization of the enzimatic levels allowing modifications in the Km's, $K_{0,5}$'s, and Ki's. The IspA ratio is set to one i.e., $X_{11}/X_{16}$ = 1. …….……….…….……..... | 30 |
| Table S3.6b | Specific constants ($K_{cat}$ /Km) ratios for data from Table 3.3a. …………………… | 34 |
| Table S3.7a | Optimization of the enzimatic levels not allowing modifications in the Km's, $K_{0,5}$'s, and Ki's. The basal ratio for IspA ($X_{11}/X_{16}$) is set to one all along the optimizations. ……………………………………………………………………….. | 36 |
| Table S3.8a | Optimization of the *K*cat's allowing modifications for the Km's, $K_{0,5}$'s, and Ki's. Assuming no relationship betweenIspA kcat's $X_{26}$ and $X_{32}$. Optimization of the enzimatic levels allowing modifications in the Km's, $K_{0,5}$'s, and Ki's. The IspA ratio is set to one i.e., $X_{11}/X_{16}$ = 1.………………...…………………………….. | 39 |
| Table S3.8b | Specific constants ($K_{cat}$ /Km) ratios for data from Table 3.8a. …………………… | 42 |
| Table S3.9a | Optimization of the *K*cat's allowing modifications for the Km's, $K_{0,5}$'s, and Ki's. Assuming no relationship betweenIspA kcat's $X_{26}$ and $X_{32}$. Optimization of the enzimatic levels allowing modifications in the Km's, $K_{0,5}$'s, and Ki's. The IspA $X_{11}$ and $X_{16}$ are independent between them.………………...…………………… | 43 |

Table S3.9b Specific constants ($K_{cat}$ /Km) ratios for data from Table 3.9a. …………………… 46
S3.7 Optimizations equations. ………..…………………………..……………………...... 47
References. ……………...……………………………………………………………. 57

### S3.- Model optimization

The 61 independent variables can be grouped in 8 blocks named: 1) enzymatic concentrations ($X_{11}$... $X_{16}$, $X_{20}$, $X_{21}$,$X_{23}$,$X_{24}$, $X_{68}$), 2) catalytic turnovers ($X_{26}$…$X_{36}$), 3) Km's ($X_{37}$, $X_{54}$, $X_{56}$,$X_{60}$), 4) Kd's ($X_{62}$, $X_{64}$….$X_{67}$), 5) metabolic pools constants ($X_{17}$…$X_{19}$, $X_{25}$, $X_{71}$), 6) flux model divertissement rate constant ($X_{55}$, $X_{58}$, and $X_{72}$), 7) constants of enzymatic activation and inhibition ($X_{57}$, $X_{59}$, $X_{69}$), and 8) intrinsic enzymatic mathematical constants ($X_{61}$, $X_{63}$, $X_{70}$).

From these above mentioned blocks the numbered from 1 to 6 are the one's allowed to be modified during the set of the performed optimizations. Blocks 7 and 8 were not allowed be modified because is our thinking they are more difficult to be performed operationally as a battery of enzymatic modifications of these parameters are not easily be biotechnologically implemented.

#### S3.1. *Flux maximization through the IspA* ($X_{11}$).

The objective function was maximizing flux direction terpenoids. In our model this flux corresponds at:

$$v_{4,11} = 0.7646372 \cdot X_4^{0.996861} \cdot X_9^{0.994367} \cdot X_{11} \cdot X_{26} \cdot X_{37}^{-0.002939} \cdot X_{56}^{-0.996851} \cdot X_{62}^{-0.991427}$$

Eq. S30

Afterlog-transforming Eq. S30 read as

$$0.9968512 \cdot y4 + 0.9943671 \cdot y9 + y11 + y26 - 0.0029399 \cdot y37 - 0.9968512 \cdot y56 - 0.9914271 \cdot y62\,;$$

where $y_i = ln\, X_i$ , for $i$ = {4,9,11,26,37,56,62} Eq. S31

#### S3.2.- *Steady state restrictions*

The second step was the formulation of the steady state S-system equations following the log-transformation and posterior $K$ translation of the power-law elements. As example the first equation for $V_1^+$ (S1.2, S1.4, S1.5) takes the following shape:

$$0.0319154 \cdot y17 - 0.9469132 \cdot y1 + 0.4947393 \cdot y18 - 0.0028433 \cdot y22 + y24 - y27 + y36 + 0.9469132 \cdot y38 - 0.0029634 \cdot y4 - 0.0241458 \cdot y48 - 0.4947393 \cdot y49 - 0.0029848 \cdot y5 + 0.0002202 \cdot y53 + 0.0029634 \cdot y57 + 0.0029848 \cdot y59 - 0.0077696 \cdot y64 + 0.0026230 \cdot y65 - y68 = -0.6393922 - 1.88163e^{-16} \cdot K$$

where $y_i = ln\, X_i + K$ for $i = \{4,5,18,22,24,27,36,48,49,53,57,59,64,65,68\}$ and $K = 50$ Eq. S32

The rest of the S-system log-linearized equations are represented in the appendix equations S3.5

Please note that the constant $K = 50$ was added at the logarithmic transformation simply to translate the log-model at the positive Cartesian space as specific requirement of the optimization software [1].

**S3.3.- *External model metabolites restrictions.***

Limits for the concentration of metabolites: restrictions of homeostasis. These restrictions determine how many the enzymes concentrations were allowed to vary.

It is also necessary to limit the range of variation of the concentration of the time dependent metabolites. Over expression of enzymes is associated with increased concentrations of intermediates something that is self-controlled by the cell and frequently out of the hands of the biotechnological process. Therefore, the metabolites were allowed to vary by 20% around their basal value, a sufficiently small range to ensure that there are no changes in other processes that affect the yield of lycopens. In other cases the range allowed was increased to [0.1, 2.5] for test the power of the optimization when the time dependent variables range of variation is increased (Table 1)

$$y_i \leq ln\,(X_i \cdot UB_i) + K$$
$$y_i \geq ln\,(X_i \cdot LB_i) + K$$

where $y_i = ln\, X_i + K$ for $i = \{1\ldots10\}$, and $K = 50$ Eqs. S34

For example the first time dependent variable DXP ($X_1$) have a concentration at steady state of $2.519343e^{-3}$ mM (Table S1.1). After log transformation and translation we obtain the range for their optimization as:

$$y_1 \leq ln\,(2.519343e^{-3} \cdot UB_1)\; + K$$
$$y_1 \geq ln\,(2.519343e^{-3} \cdot LB_1)\; + K$$

with $LB_1 = 0.1, UB_1 = 2.5$ and $K = 50$

**S3.4.- *Independent variables boundaries.***

The independent parameters are restricted according the following generic equation

$$y_i - (ln\,(X_i \cdot UB_i)\; + K) \cdot W_i - (ln\,(X_i)\; + K) \cdot M_i \leq 0$$
$$y_i - (ln\,(X_i \cdot LB_i)\; + K) \cdot W_i - (ln\,(X_i)\; + K) \cdot M_i \geq 0 \qquad \text{Eqs. S35}$$

where $y_i = ln\, X_i\; + K$

$W_i + M_i = 1$ in which $W_i$ and $M_i$can take the form of the binary values 0 or 1, for $i = \{11...72\}$ and

For the case of IspA ($X_{11}$) we have

$$y_{11} - (ln\,(1.632863e^{-4} \cdot UB_{11})\; + K) \cdot W_{11} - (ln\,(1.632863e^{-4})\; + K) \cdot M_{11} \leq 0$$
$$y_{11} - (ln\,(1.632863e^{-4} \cdot LB_{11})\; + K) \cdot W_{11} - (ln\,(1.632863e^{-4})\; + K) \cdot M_{11} \geq 0$$

with $y_{11} = ln\, X_{11}\; + K$ and $K$ =50

For the energetic sources:

Pyruvate ($X_{17}$) and G3P ($X_{18}$), ATP ($X_{19}$), adenine nucleotides ($X_{22}$) and CTP ($X_{25}$) were allowed to vary by 10% around the corresponding basal values. This is consistent with the small range of variation reported for such variables in related systems [2; 3]

For the optimizations of the enzymes the range was chose between1 and 10 times its basal value; this range of variation is consistent with those obtained by means of recombinant DNA techniques [4; 5].

The optimization biotechnological implementation for MEC membrane pumps are difficult to implement. The reason of this difficulty are diverse: first the MEC extrusion pump(s) are not yet well determinate and until our knowledge exist only candidates as antiporters (*sfr*) and ATP driven candidates [6], second in many cases these pumps work synergistically with overlapping functions in case one is impaired and board range of substrates as reported for the antibiotics pumps [7; 8]. Because the above reasons, the MEC transporter ($X_{13}$) and their corresponding Kcat ($X_{29}$) optimizations results need to be taken with caution.

#### S3.5.- *Enzymes modification restriction*

In case of exert optimizations allowing the variation of the Km's and Kd's, they can be optimized only if the corresponding enzyme or Kcat is also allowed to bascule around their basal value:

$$y_i - (ln\,(X_i \cdot UB_i)\; + K) \cdot W_j - (ln\,(X_i)\; + K) \cdot M_j \leq 0$$
$$y_i - (ln\,(X_i \cdot LB_i)\; + K) \cdot W_j - (ln\,(X_i)\; + K) \cdot M_j \geq 0$$
$$W_i + M_i = 1$$

with

$i = \{X_{37}, X_{54}, X_{56}, X_{60}, X_{62}, X_{64} \ldots X_{67}\}$ for the Km's and Kcat's

$j = \{X_{11} \ldots X_{16}, X_{20}, X_{21}, X_{23}, X_{24}, X_{68}\}$ for the enzyme's concentrations

or

$j = \{X_{26} \ldots X_{36}\}$ for the Kcat's Eqs. S36

where $y_i$ is the $X_i$ log transformation of the corresponding Km and/or Kd. The binary tandem $\{W_j, M_j\}$ correspond at the associated $X_j$ enzyme or Kcat. Only in case $W_j = 1$ the $y_i$ enzyme or Kd is modified.

For example, in case of optimization of the Kcat's allowing modification of the Km_9_11 ($X_{37}$), Km_4_11 ($X_{56}$) and Kd_9_11 ($X_{62}$) they are modified only if Kcat_11 ($X_{26}$) is also allowed to be modified (Table S3.1):

$$y_{37} - (ln\,(5.5e^{-3} \cdot UB_{37}) + K) \cdot W_{26} - (ln\,(5.5e^{-3}) + K) \cdot M_{26} \leq 0$$
$$y_{37} - (ln\,(5.5e^{-3} \cdot LB_{37}) + K) \cdot W_{26} - (ln\,(5.5e^{-3}) + K) \cdot M_{26} \geq 0$$
$$y_{56} - (ln\,(1.03e^{-2} \cdot UB_{56}) + K) \cdot W_{26} - (ln\,(1.03e^{-2}) + K) \cdot M_{26} \leq 0$$
$$y_{56} - (ln\,(1.03e^{-2} \cdot LB_{56}) + K) \cdot W_{26} - (ln\,(1.03e^{-2}) + K) \cdot M_{26} \geq 0$$
$$y_{62} - (ln\,(7.14e^{-2} \cdot UB_{62}) + K) \cdot W_{26} - (ln\,(7.14e^{-2}) + K) \cdot M_{26} \leq 0$$
$$y_{62} - (ln\,(7.14e^{-2} \cdot LB_{62}) + K) \cdot W_{26} - (ln\,(7.14e^{-2}) + K) \cdot M_{26} \geq 0$$

with $W_{26} + M_{26} = 1$. When $W_{26} = 1$ the boundary ranges for the above enzymes are allowed be modified during the optimization process.

Please note that $X_{37}$, $X_{56}$, and $X_{62}$ also have their corresponding restrictions as describes in Eqs. S35. At the end it is the optimization program which decides if the parameter is going to be modified or not. These restrictions only allow the Km's and Kd's modifications but does not force at the parameters to be modified.

**S3.6.-*Total enzyme restriction: The cellular economy factor***

As mentioned above, the upper limit for the total concentration of enzymes must be carefully controlled as a whole, since in the case of overproduction of proteins we could reach a point of cellular infeasibility. It is for this reason that the cell economy factor was established as a restriction to the maximum total concentration of enzyme. This factor was set at 1 or 2 times the total value of basal enzymatic concentration and was expressed as:

$$\sum_i X_i \leq X_t \qquad \text{Eq. S35}$$

where $X_t$ is:

$$X_t = p \cdot \sum_i X_{i_0}, \qquad p = \{1,2\} \qquad \text{Eq. S36}$$

Transforming (Eq. 25) to power law formalism:

$$X_t \geq \gamma_t \prod_i X_i^{f_{i,t}} \qquad \text{Eq.S37}$$

In this equation $\gamma$ is a parameter analogous to the previously defined BST rate constants $\alpha$ and $\beta$. Similarly the ko's $f_{i,t}$ is similar at the BST kinetic orders $g_{i,j}$ and $h_{i,j}$ [9].

Log-linearizing Eq S37 is obtained the equation S36 for $p = \{1,2\}$

$$0.2774294 \cdot y11 + 0.1630094 \cdot y12 + 0.0250783 \cdot y13 + 0.0869905 \cdot y14 + 0.0047021 \cdot y15 + 0.0391849 \cdot y20 + 0.1708463 \cdot y21 + 0.1598746 \cdot y23 + 0.0478056 \cdot y24 + 0.0250783 \cdot y68 \leq = 1.0 \cdot K + @log(p) - 9.3790079$$

where $y_i = ln\, X_i$ for $i = \{11,12,13,14,15,20,21,23,24,68\}$ and $K = 50$ Eq. S38

Please note than Eq. S38 does not include $y_{16}$ as IspA is already present once as $y_{11}$.

**Table S3.1a**. Optimization of the *K*cat's allowing modifications for the Km's, $K_{0,5}$'s, and Ki's. Assuming no relationship between IspA kcat's $X_{26}$ and $X_{32}$.

| | | | | | | Opt./Basal | | | | |
|---|---|---|---|---|---|---|---|---|---|---|
| | | | Basal | LB | UB | 2 | 3 | 4 | 5 | 6 |
| Time-dependent | DXP (mM) | $X_1$ | 0,002519342 | 0,1 | 2,5 | 0,63 | 0,65 | 0,63 | 0,63 | 0,63 |
| | CDP_MEP | $X_2$ | 0,002135892 | 0,1 | 2,5 | 0,85 | 0,87 | 0,85 | 0,85 | 0,85 |
| | MEcDP | $X_3$ | 0,001552085 | 0,1 | 2,5 | 1,24 | 1,24 | 1,24 | 1,24 | 1,24 |
| | IPP | $X_4$ | 0,000396505 | 0,1 | 2,5 | 2,50 | 2,50 | 2,50 | 2,50 | 2,50 |
| | DMAPP | $X_5$ | 0,000496462 | 0,1 | 2,5 | 1,18 | 1,19 | 1,15 | 1,12 | 1,12 |
| | MEP | $X_6$ | 0,000125026 | 0,1 | 2,5 | 0,64 | 0,66 | 0,64 | 0,64 | 0,64 |
| | $MEC_{(c)}$ | $X_7$ | 0,000893951 | 0,1 | 2,5 | 0,38 | 0,40 | 0,38 | 0,38 | 0,38 |
| | HMBPP | $X_8$ | 3,84307E-05 | 0,1 | 2,5 | 0,39 | 0,40 | 0,39 | 0,38 | 0,38 |
| | GPP | $X_9$ | 0,000390628 | 0,1 | 2,5 | 2,50 | 2,50 | 2,50 | 2,50 | 2,50 |
| | $MEC_{(e)}$ | $X_{10}$ | 0,02709473 | 0,1 | 2,5 | 1,16 | 1,19 | 1,16 | 1,16 | 1,16 |
| | kcat_11 ($min^{-1}$) | $X_{26}$ | 1,7316 | 0,2 | 1,2 | 1 | 1 | 1,14 | 1,17 | 1,17 |
| | kcat_12 | $X_{28}$ | 1010 | 0,2 | 10 | 1 | 1 | 1 | 1 | 1 |
| | kcat_13 | $X_{29}$ | 600 | 0,2 | 10 | 1 | 1 | 1 | 1 | 1 |
| | kcat_14 | $X_{30}$ | 80,7 | 0,2 | 10 | 1 | 1 | 1 | 1 | 1 |
| | kcat_15 | $X_{31}$ | 19,8 | 0,2 | 10 | 9,31 | 10 | 10 | 10 | 10 |
| | kcat_16 | $X_{32}$ | 0,78 | 0,2 | 1,2 | 1 | 1 | 1,14 | 1,17 | 1,17 |
| | kcat_20 | $X_{33}$ | 1554 | 0,2 | 10 | 1 | 1 | 1 | 1 | 1 |
| | kcat_21 | $X_{34}$ | 27,6 | 0,2 | 10 | 1 | 1 | 1 | 1 | 1 |
| | kcat_23 | $X_{35}$ | 23,7 | 0,2 | 10 | 1 | 1 | 1 | 1,08 | 1,08 |
| | kcat_24 | $X_{36}$ | 100,3 | 0,2 | 10 | 0,2 | 0,2 | 0,2 | 0,2 | 0,2 |
| | kcat_68 | $X_{27}$ | 1740 | 0,2 | 10 | 1 | 1 | 1 | 1 | 1 |
| | kcat_11/ kcat_16 | $X_{26}/X_{32}$ | 2.22 | - | - | 2,22 | 2,66 | 4,81 | 4,81 | 4,81 |
| Time-independent | Piruvate (mM) | $X_{17}$ | 2,67 | 0,9 | 1,1 | 0,9 | 0,9 | 0,9 | 0,9 | 0,9 |
| | G3P | $X_{18}$ | 0,041 | 0,9 | 1,1 | 0,9 | 0,9 | 0,9 | 0,9 | 0,9 |
| | ATP | $X_{19}$ | 3 | 0,9 | 1,1 | 0,9 | 0,9 | 0,9 | 0,9 | 0,9 |
| | NADPH | $X_{22}$ | 0,12 | 0,9 | 1,1 | 0,9 | 0,9 | 0,9 | 0,9 | 0,9 |
| | CTP | $X_{25}$ | 2,7 | 0,9 | 1,1 | 0,9 | 0,9 | 0,9 | 0,9 | 0,9 |
| | FldA | $X_{71}$ | 0,003015 | 0,9 | 1,1 | 0,9 | 0,9 | 0,9 | 0,9 | 0,9 |
| | Km_1_68 (mM) | $X_{38}$ | 0,045 | 0,9 | 1,1 | 1 | 1 | 1 | 1 | 1 |
| | Km_2_12 | $X_{39}$ | 0,15 | 0,9 | 1,1 | 1 | 1 | 1 | 1 | 1 |
| | Km_3_14 | $X_{40}$ | 0,339 | 0,9 | 1,1 | 1 | 1 | 1 | 1 | 1 |
| | Km_4_15 | $X_{41}$ | 0,0095 | 0,9 | 1,1 | 1,1 | 1,1 | 1,1 | 1,1 | 1,1 |
| | Km_4_16 | $X_{46}$ | 0,0013 | 0,9 | 1,1 | 1 | 1 | 1,1 | 1,1 | 1,1 |
| | Km_4_11 | $X_{56}$ | 0,0103 | 0,9 | 1,1 | 1 | 0,9 | 0,9 | 0,9 | 0,9 |
| | Km_5_15 | $X_{42}$ | 0,0143 | 0,9 | 1,1 | 0,9 | 0,90 | 0,9 | 0,9 | 0,9 |
| | Km_5_16 | $X_{47}$ | 0,0293 | 0,9 | 1,1 | 1 | 1 | 1,1 | 1,1 | 1,1 |
| | Km_6_20 | $X_{43}$ | 0,00314 | 0,9 | 1,1 | 1 | 1 | 1 | 1 | 1 |
| | Km_7_13 | $X_{60}$ | 0,00495 | 0,9 | 1,1 | 1 | 1 | 1 | 1 | 1 |
| | Km_7_23 | $X_{44}$ | 0,56 | 0,9 | 1,1 | 1 | 1 | 1 | 1,1 | 1,1 |

| | | | | | | | | | | |
|---|---|---|---|---|---|---|---|---|---|---|
| | Km_8_21 | $X_{45}$ | 0,03 | 0,9 | 1,1 | 1 | 1 | 1 | 1 | 1 |
| | Km_9_11 | $X_{37}$ | 0,0055 | 0,9 | 1,1 | 1 | 0,9 | 0,9 | 0,9 | 0,9 |
| | Km_17_24 | $X_{48}$ | 0,134 | 0,9 | 1,1 | 1,1 | 1.1 | 1.1 | 1,1 | 1,1 |
| | Km_18_24 | $X_{49}$ | 0,041 | 0,9 | 1,1 | 1,1 | 1,05 | 1,1 | 1,1 | 1,1 |
| | Km_19_12 | $X_{50}$ | 0,42 | 0,9 | 1,1 | 1 | 1 | 1 | 1 | 1 |
| | Km_71_21 | $X_{51}$ | 0,012 | 0,9 | 1,1 | 1 | 1 | 1 | 1 | 1 |
| | Km_71_23 | $X_{52}$ | 0,012 | 0,9 | 1,1 | 1 | 1 | 1 | 0,9 | 0,9 |
| | Km_22_68 | $X_{53}$ | 0,0005 | 0,9 | 1,1 | 1 | 1 | 1 | 1 | 1 |
| | Km_25_20 | $X_{54}$ | 0,76 | 0,9 | 1,1 | 1 | 1 | 1 | 1 | 1 |
| Time-independent | K_10_55 (mM) | $X_{55}$ | 0,05 | 0,9 | 1,1 | 0,9 | 0,9 | 0,9 | 0,9 | 0,9 |
| | K_4_58 | $X_{58}$ | 0,005 | 0,9 | 1,1 | 1,01 | 1,1 | 0,98 | 0,9 | 0,9 |
| | K_9_72 | $X_{72}$ | 0,005 | 0,9 | 1,1 | 1,1 | 1,1 | 1,1 | 1,1 | 1,1 |
| | Ki_4_24 | $X_{57}$ | 0,0654 | 1 | 1 | 1 | 1 | 1 | 1 | 1 |
| | Ki_5_24 | $X_{59}$ | 0,0813 | 1 | 1 | 1 | 1 | 1 | 1 | 1 |
| | Ka_6_14 | $X_{69}$ | 0,00339 | 0,9 | 1,1 | 1 | 1 | 1 | 1 | 1 |
| | $\alpha_6$ (adimensional) | $X_{61}$ | 1 | 0,9 | 1,1 | 1 | 1 | 1 | 1 | 1 |
| | $\beta_6$(adimensional) | $X_{63}$ | 2000 | 0,9 | 1,1 | 1 | 1 | 1 | 1 | 1 |
| | Kd_9_11 | $X_{62}$ | 0,0714 | 0,9 | 1,1 | 1 | 1 | 0,9 | 0,9 | 0,9 |
| | Kd_17_24 | $X_{64}$ | 0,0426 | 0,9 | 1,1 | 1,1 | 0,9 | 1,1 | 1,1 | 1,1 |
| | Kd_22_68 | $X_{65}$ | 0,000333333 | 0,9 | 1,1 | 1 | 1 | 1 | 1 | 1 |
| | Kd_71_23 | $X_{66}$ | 0,0005 | 0,9 | 1,1 | 1 | 1 | 1 | 1,1 | 1,1 |
| | Kd_71_21 | $X_{67}$ | 0,0005 | 0,9 | 1,1 | 1 | 1 | 1 | 1 | 1 |
| | hill_7_13 (adimensional) | $X_{70}$ | 1 | 0,9 | 1,1 | 1 | 1 | 1 | 1 | 1 |
| | $(v_{4,11})_{opt} / (v_{4,11})_{basal}$ | | | | | 6,10 | 6,77 | 8,53 | 8,74 | 8,74 |

***a***) Time dependent metabolites ($X_1$... $X_{10}$) ranges between 0.1 and 2.5 times its respective basal values

***b***) External variables ($X_{17}$, $X_{18}$, $X_{22}$, $X_{25}$, and $X_{71}$) allowed varying a 10% around their respective basal values.

***c***) The kcat's ($X_{27}$...$X_{31}$, $X_{33}$... $X_{36}$) ranges between 0.2 and 10 times their basal values at exception of IspA kcat's$X_{26}$ and $X_{32}$ with upper level 1.2 times their respective basal values

***d***) Cellular economy factor p = 1

***e***) The $v_{9,11}$basal flux was:

$$v_{9,11} = \frac{V_{max_{11}} \times X_4 \times X_9}{\left(K_{m_{4,11}} \times X_9 + K_{m_{9,11}} \times X_4 + X_4 \times X_9 + K_{d_{9,11}} \times K_{m_{9,11}}\right)} = 2.83 \times 10^{-4}\ mM \cdot min^{-1}$$

**Table S3.1a**. (cont…)

| | | | Basal | LB | UB | Opt./Basal 7 | 8 | 9 | 10 | 11 |
|---|---|---|---|---|---|---|---|---|---|---|
| Time-dependent | DXP (mM) | $X_1$ | 0,002519342 | 0,1 | 2,5 | 1,74 | 1,74 | 0,17 | 0,1 | 2,50 |
| | CDP_MEP | $X_2$ | 0,002135892 | 0,1 | 2,5 | 2,26 | 2,26 | 2,5 | 2,5 | 2,50 |
| | MEcDP | $X_3$ | 0,001552085 | 0,1 | 2,5 | 1,24 | 1,24 | 2,5 | 2,5 | 2,50 |
| | IPP | $X_4$ | 0,000396505 | 0,1 | 2,5 | 2,5 | 2,5 | 2,5 | 2,5 | 2,50 |
| | DMAPP | $X_5$ | 0,000496462 | 0,1 | 2,5 | 1,12 | 1,12 | 1,12 | 1,12 | 1,12 |
| | MEP | $X_6$ | 0,000125026 | 0,1 | 2,5 | 1,76 | 1,76 | 1,76 | 2,5 | 0,14 |
| | $MEC_{(c)}$ | $X_7$ | 0,000893951 | 0,1 | 2,5 | 1,2 | 1,2 | 1,2 | 0,67 | 0,10 |
| | HMBPP | $X_8$ | 3,84307E-05 | 0,1 | 2,5 | 0,37 | 0,37 | 1,65 | 1,65 | 2,08 |
| | GPP | $X_9$ | 0,000390628 | 0,1 | 2,5 | 2,5 | 2,5 | 2,5 | 2,5 | 2,50 |
| | $MEC_{(e)}$ | $X_{10}$ | 0,02709473 | 0,1 | 2,5 | 2,5 | 2,5 | 2,5 | 1,87 | 2,50 |
| | kcat_11 ($min^{-1}$) | $X_{26}$ | 1,7316 | 0,2 | 1,2 | 1,17 | 1,17 | 1,17 | 1,17 | 1,20 |
| | kcat_12 | $X_{28}$ | 1010 | 0,2 | 10 | 1 | 1 | 0,99 | 0,61 | 1 |
| | kcat_13 | $X_{29}$ | 600 | 0,2 | 10 | 1 | 1 | 1 | 1 | 7,52 |
| | kcat_14 | $X_{30}$ | 80,7 | 0,2 | 10 | 1 | 1 | 0,55 | 0,24 | 5 |
| | kcat_15 | $X_{31}$ | 19,8 | 0,2 | 10 | 10 | 10 | 10 | 10 | 10,00 |
| | kcat_16 | $X_{32}$ | 0,78 | 0,2 | 1,2 | 1,17 | 1,17 | 1,17 | 1,17 | 0,96 |
| | kcat_20 | $X_{33}$ | 1554 | 0,2 | 10 | 1 | 1 | 1 | 0,40 | 10 |
| | kcat_21 | $X_{34}$ | 27,6 | 0,2 | 10 | 1 | 1 | 0,2 | 0,2 | 0,20 |
| | kcat_23 | $X_{35}$ | 23,7 | 0,2 | 10 | 0,33 | 0,33 | 0,33 | 0,59 | 3 |
| | kcat_24 | $X_{36}$ | 100,3 | 0,2 | 10 | 0,47 | 0,47 | 0,47 | 0,29 | 0,52 |
| | kcat_68 | $X_{27}$ | 1740 | 0,2 | 10 | 1 | 1 | 10 | 10 | 1 |
| | kcat_11/ kcat_16 | $X_{26}/X_{32}$ | 2.22 | - | - | 4,81 | 2,28 | 2,28 | 2,28 | 2,78 |
| Time-independent | Piruvate (mM) | $X_{17}$ | 2,67 | 0,9 | 1,1 | 0,9 | 0,9 | 0,9 | 0,9 | 0,90 |
| | G3P | $X_{18}$ | 0,041 | 0,9 | 1,1 | 0,9 | 0,9 | 0,9 | 0,9 | 0,90 |
| | ATP | $X_{19}$ | 3 | 0,9 | 1,1 | 0,9 | 0,9 | 1,1 | 1,1 | 1,10 |
| | NADPH | $X_{22}$ | 0,12 | 0,9 | 1,1 | 0,9 | 0,9 | 0,9 | 0,9 | 1,10 |
| | CTP | $X_{25}$ | 2,7 | 0,9 | 1,1 | 0,9 | 0,9 | 0,9 | 1,1 | 1,10 |
| | FldA | $X_{71}$ | 0,003015 | 0,9 | 1,1 | 1,1 | 1,1 | 1,1 | 1,1 | 1,10 |
| | Km_1_68 (mM) | $X_{38}$ | 0,045 | 0,9 | 1,1 | 1 | 1 | 1,1 | 1,1 | 0,9 |
| | Km_2_12 | $X_{39}$ | 0,15 | 0,9 | 1,1 | 1 | 1 | 1,1 | 1,1 | 0,9 |
| | Km_3_14 | $X_{40}$ | 0,339 | 0,9 | 1,1 | 1 | 1 | 1,1 | 1,1 | 0,9 |
| | Km_4_15 | $X_{41}$ | 0,0095 | 0,9 | 1,1 | 1,1 | 1,1 | 1,1 | 1,1 | 1,1 |
| | Km_4_16 | $X_{46}$ | 0,0013 | 0,9 | 1,1 | 1,1 | 1,1 | 1,1 | 1,1 | 0,9 |
| | Km_4_11 | $X_{56}$ | 0,0103 | 0,9 | 1,1 | 0,9 | 0,9 | 0,9 | 0,9 | 0,9 |
| | Km_5_15 | $X_{42}$ | 0,0143 | 0,9 | 1,1 | 0,9 | 0,9 | 0,9 | 0,9 | 0,9 |
| | Km_5_16 | $X_{47}$ | 0,0293 | 0,9 | 1,1 | 1,1 | 1,1 | 1,1 | 1,1 | 0,9 |
| | Km_6_20 | $X_{43}$ | 0,00314 | 0,9 | 1,1 | 1 | 1 | 1 | 0,9 | 0,9 |
| | Km_7_13 | $X_{60}$ | 0,00495 | 0,9 | 1,1 | 1 | 1 | 1 | 1 | 0,9 |
| | Km_7_23 | $X_{44}$ | 0,56 | 0,9 | 1,1 | 1,1 | 1,1 | 1,1 | 1,1 | 0,9 |
| | Km_8_21 | $X_{45}$ | 0,03 | 0,9 | 1,1 | 1 | 1 | 0,9 | 0,9 | 1,1 |
| | Km_9_11 | $X_{37}$ | 0,0055 | 0,9 | 1,1 | 0,9 | 0,9 | 0,9 | 0,9 | 0,9 |
| | Km_17_24 | $X_{48}$ | 0,134 | 0,9 | 1,1 | 1,1 | 1,1 | 1,1 | 1,1 | 1,1 |
| | Km_18_24 | $X_{49}$ | 0,041 | 0,9 | 1,1 | 0,9 | 0,9 | 0,9 | 0,9 | 1,1 |

| | | | | | | | | | | |
|---|---|---|---|---|---|---|---|---|---|---|
| | Km_19_12 | $X_{50}$ | 0,42 | 0,9 | 1,1 | 1 | 1 | 1,1 | 1,1 | 0,9 |
| | Km_71_21 | $X_{51}$ | 0,012 | 0,9 | 1,1 | 1 | 1 | 0,9 | 0,9 | 1,1 |
| | Km_71_23 | $X_{52}$ | 0,012 | 0,9 | 1,1 | 0,96 | 0,96 | 0,9 | 0,9 | 0,9 |
| | Km_22_68 | $X_{53}$ | 0,0005 | 0,9 | 1,1 | 1 | 1 | 1,1 | 1,1 | 0,9 |
| | Km_25_20 | $X_{54}$ | 0,76 | 0,9 | 1,1 | 1 | 1 | 1 | 1,1 | 0,9 |
| Time-independent | K_10_55 (mM) | $X_{55}$ | 0,05 | 0,9 | 1,1 | 1,1 | 1,1 | 1,1 | 0,9 | 1,1 |
| | K_4_58 | $X_{58}$ | 0,005 | 0,9 | 1,1 | 0,9 | 0,9 | 0,9 | 0,9 | 0,9 |
| | K_9_72 | $X_{72}$ | 0,005 | 0,9 | 1,1 | 1,1 | 1,1 | 1,1 | 1,1 | 1,1 |
| | Ki_4_24 | $X_{57}$ | 0,0654 | 1 | 1 | 1 | 1 | 1 | 1 | 1 |
| | Ki_5_24 | $X_{59}$ | 0,0813 | 1 | 1 | 1 | 1 | 1 | 1 | 1 |
| | Ka_6_14 | $X_{69}$ | 0,00339 | 0,9 | 1,1 | 1 | 1 | 1 | 1 | 1 |
| | $\alpha_6$ (adimensional) | $X_{61}$ | 1 | 0,9 | 1,1 | 1 | 1 | 1 | 1 | 1 |
| | $\beta_6$(adimensional) | $X_{63}$ | 2000 | 0,9 | 1,1 | 1 | 1 | 1 | 1 | 1 |
| | Kd_9_11 | $X_{62}$ | 0,0714 | 0,9 | 1,1 | 0,9 | 0,9 | 0,9 | 0,9 | 1,1 |
| | Kd_17_24 | $X_{64}$ | 0,0426 | 0,9 | 1,1 | 0,9 | 0,9 | 0,9 | 0,9 | 0,9 |
| | Kd_22_68 | $X_{65}$ | 0,000333333 | 0,9 | 1,1 | 1 | 1 | 1,1 | 1,1 | 1,1 |
| | Kd_71_23 | $X_{66}$ | 0,0005 | 0,9 | 1,1 | 0,96 | 0,96 | 0,9 | 0,9 | 0,9 |
| | Kd_71_21 | $X_{67}$ | 0,0005 | 0,9 | 1,1 | 1 | 1 | 0,9 | 0,9 | 0,9 |
| | hill_7_13(adimensional) | $X_{70}$ | 1 | 0,9 | 1,1 | 1 | 1 | 1 | 1 | 1,1 |
| | $(v_{4,11})_{opt}/(v_{4,11})_{basal}$ | | | | | 8,74 | 8,74 | 8,74 | 8,74 | 8,74 |

**Table S3.1b.** Specific constants (kcat/Km) ratios for data from Table S3.1a

| | | | Opt./Basal | | | | |
|---|---|---|---|---|---|---|---|
| **kcat / Km** | **Model symbols** | **Basal** | **2** | **3** | **4** | **5** | **6** |
| kcat_11/Km_4_11 | $X_{26}/ X_{56}$ | 168,12 | 1 | 1 | 1,33 | 1,33 | 1,33 |
| kcat_11/Km_9_11 | $X_{26} / X_{37}$ | 314,84 | 1 | 1 | 1,33 | 1,33 | 1,33 |
| kcat_12 / Km_2_12 | $X_{28} / X_{39}$ | 6733,33 | 1 | 1 | 1 | 1 | 1 |
| kcat_12 / Km_19_12 | $X_{28} / X_{50}$ | 2404,76 | 1 | 1 | 1 | 1 | 1 |
| kcat_13 / Km_7_13 | $X_{29} / X_{60}$ | 121212,12 | 1 | 1 | 1 | 1 | 0,98 |
| kcat_14 / Km_3_14 | $X_{30} / X_{40}$ | 238,05 | 1 | 1 | 1 | 1 | 1 |
| kcat_15 / Km_4_15 | $X_{31} / X_{41}$ | 2084,21 | 9,37 | 9,37 | 0,72 | 0,72 | 0,82 |
| kcat_15 / Km_5_15 | $X_{31} / X_{42}$ | 1384,62 | 11,11 | 11,11 | 0,88 | 0,88 | 1,00 |
| kcat_16 / Km_4_16 | $X_{32} / X_{46}$ | 600,00 | 1 | 1 | 0,50 | 0,50 | 0,50 |
| kcat_16 / Km_5_16 | $X_{32}/ X_{47}$ | 26,62 | 1 | 1 | 0,50 | 0,50 | 0,50 |
| kcat_20 / Km_6_20 | $X_{33} / X_{43}$ | 494904,46 | 1 | 1 | 1 | 1 | 1 |
| kcat_20 / Km_25_20 | $X_{33} / X_{54}$ | 2044,74 | 1 | 1 | 1 | 1 | 1 |
| kcat_21 / Km_8_21 | $X_{34} / X_{45}$ | 920,00 | 1 | 1 | 1 | 0,18 | 0,18 |
| kcat_21 / Km_71_21 | $X_{34} / X_{51}$ | 2300,00 | 1 | 1 | 1 | 0,22 | 0,22 |
| kcat_23 / Km_7_23 | $X_{35} / X_{44}$ | 42,32 | 1 | 1 | 1 | 1 | 1 |
| kcat_23 / Km_71_23 | $X_{35} / X_{52}$ | 1975,00 | 1 | 1 | 1 | 1 | 1 |
| kcat_24 / Km_17_24 | $X_{36} / X_{48}$ | 748,51 | 0,18 | 0,18 | 0,18 | 0,18 | 0,18 |
| kcat_24/ Km_18_24 | $X_{36} / X_{49}$ | 2446,34 | 0,18 | 0,18 | 0,18 | 0,18 | 0,18 |
| kcat_68/ Km_1_68 | $X_{27} / X_{38}$ | 38666,67 | 1 | 1 | 1 | 1 | 1 |
| kcat_68/ Km_22_68 | $X_{27} / X_{53}$ | 3480000,00 | 1 | 1 | 1 | 1 | 1 |

**Table S3.1b.** (cont..)

| kcat / Km | Model symbols | Basal | Opt./Basal 7 | 8 | 9 | 10 | 11 |
|---|---|---|---|---|---|---|---|
| kcat_11/Km_4_11 | $X_{26}/ X_{56}$ | 168,12 | 1,33 | 1,33 | 1,33 | 1,33 | 1,33 |
| kcat_11/Km_9_11 | $X_{26} / X_{37}$ | 314,84 | 1,33 | 1,33 | 1,33 | 1,33 | 1,33 |
| kcat_12 / Km_2_12 | $X_{28} / X_{39}$ | 6733,33 | 1 | 1 | 1 | 1 | 0,91 |
| kcat_12 / Km_19_12 | $X_{28} / X_{50}$ | 2404,76 | 1 | 1 | 1 | 1 | 0,91 |
| kcat_13 / Km_7_13 | $X_{29} / X_{60}$ | 121212,12 | 2,65 | 2,65 | 2,65 | 2,65 | 8,35 |
| kcat_14 / Km_3_14 | $X_{30} / X_{40}$ | 238,05 | 1 | 1 | 1 | 1 | 5,52 |
| kcat_15 / Km_4_15 | $X_{31} / X_{41}$ | 2084,21 | 0,67 | 9,09 | 9,09 | 9,09 | 9,09 |
| kcat_15 / Km_5_15 | $X_{31} / X_{42}$ | 1384,62 | 0,82 | 11,11 | 11,11 | 11,11 | 11,11 |
| kcat_16 / Km_4_16 | $X_{32} / X_{46}$ | 600,00 | 0,50 | 1,06 | 1,06 | 1,06 | 1,06 |
| kcat_16 / Km_5_16 | $X_{32}/ X_{47}$ | 26,62 | 0,50 | 1,06 | 1,06 | 1,06 | 1,06 |
| kcat_20 / Km_6_20 | $X_{33} / X_{43}$ | 494904,46 | 1 | 1 | 1 | 1 | 11,11 |
| kcat_20 / Km_25_20 | $X_{33} / X_{54}$ | 2044,74 | 1 | 1 | 1 | 1 | 11,11 |
| kcat_21 / Km_8_21 | $X_{34} / X_{45}$ | 920,00 | 0,18 | 0,18 | 0,18 | 0,18 | 0,18 |
| kcat_21 / Km_71_21 | $X_{34} / X_{51}$ | 2300,00 | 0,22 | 0,22 | 0,22 | 0,22 | 0,18 |
| kcat_23 / Km_7_23 | $X_{35} / X_{44}$ | 42,32 | 1 | 1 | 1 | 1 | 3,55 |
| kcat_23 / Km_71_23 | $X_{35} / X_{52}$ | 1975,00 | 1 | 1 | 1 | 1 | 3,55 |
| kcat_24 / Km_17_24 | $X_{36} / X_{48}$ | 748,51 | 0,43 | 0,43 | 0,43 | 0,43 | 0,48 |
| kcat_24/ Km_18_24 | $X_{36} / X_{49}$ | 2446,34 | 0,53 | 0,53 | 0,53 | 0,53 | 0,48 |
| kcat_68/ Km_1_68 | $X_{27} / X_{38}$ | 38666,67 | 1 | 1 | 1 | 1 | 0,71 |
| kcat_68/ Km_22_68 | $X_{27} / X_{53}$ | 3480000,00 | 1 | 1 | 1 | 1 | 0,71 |

**Table S3.2a**. Optimization of the *K*cat's allowing modifications for the Km's, $K_{0,5}$'s, and Ki's. Setting the IspA kcat_11 ratio $X_{26}/X_{32} = 2.22$ for all the experiments.

| | | | Basal | LB | UB | Opt./Basal | | | | | |
|---|---|---|---|---|---|---|---|---|---|---|---|
| | | | | | | 1 | 2 | 3 | 4 | 5 | 6 |
| Time-dependent | DXP (mM) | $X_1$ | 0,002519342 | 0,1 | 2,5 | - | 0,63 | 0,65 | 0,63 | 0,63 | 0,63 |
| | CDP_MEP | $X_2$ | 0,002135892 | 0,1 | 2,5 | - | 0,85 | 0,87 | 0,85 | 0,85 | 0,85 |
| | MEcDP | $X_3$ | 0,001552085 | 0,1 | 2,5 | - | 1,24 | 1,24 | 1,24 | 1,24 | 1,24 |
| | IPP | $X_4$ | 0,000396505 | 0,1 | 2,5 | - | 2,50 | 2,50 | 2,50 | 2,50 | 2,50 |
| | DMAPP | $X_5$ | 0,000496462 | 0,1 | 2,5 | - | 1,18 | 1,19 | 1,15 | 1,12 | 1,12 |
| | MEP | $X_6$ | 0,000125026 | 0,1 | 2,5 | - | 0,64 | 0,66 | 0,64 | 0,64 | 0,64 |
| | $MEC_{(c)}$ | $X_7$ | 0,000893951 | 0,1 | 2,5 | - | 0,38 | 0,40 | 0,38 | 0,38 | 0,38 |
| | HMBPP | $X_8$ | 3,84307E-05 | 0,1 | 2,5 | - | 0,39 | 0,40 | 0,39 | 0,38 | 0,38 |
| | GPP | $X_9$ | 0,000390628 | 0,1 | 2,5 | - | 2,50 | 2,50 | 2,50 | 2,50 | 2,50 |
| | $MEC_{(e)}$ | $X_{10}$ | 0,02709473 | 0,1 | 2,5 | - | 1,16 | 1,19 | 1,16 | 1,16 | 1,16 |
| Time-independent | kcat_11 ($min^{-1}$) | $X_{26}$ | 1,7316 | 0,2 | 1,2 | - | 1 | 1 | 1,14 | 1,17 | 1,17 |
| | kcat_12 | $X_{28}$ | 1010 | 0,2 | 10 | - | 1 | 1 | 1 | 1 | 1 |
| | kcat_13 | $X_{29}$ | 600 | 0,2 | 10 | - | 1 | 1 | 1 | 1 | 1 |
| | kcat_14 | $X_{30}$ | 80,7 | 0,2 | 10 | - | 1 | 1 | 1 | 1 | 1 |
| | kcat_15 | $X_{31}$ | 19,8 | 0,2 | 10 | - | 9,31 | 10 | 10 | 10 | 10 |
| | kcat_16 | $X_{32}$ | 0,78 | 0,2 | 1,2 | - | **1** | **1** | 1,14 | 1,17 | 1,17 |
| | kcat_20 | $X_{33}$ | 1554 | 0,2 | 10 | - | 1 | 1 | 1 | 1 | 1 |
| | kcat_21 | $X_{34}$ | 27,6 | 0,2 | 10 | - | 1 | 1 | 1 | 1 | 1 |
| | kcat_23 | $X_{35}$ | 23,7 | 0,2 | 10 | - | 1 | 1 | 1 | 1,08 | 1,08 |
| | kcat_24 | $X_{36}$ | 100,3 | 0,2 | 10 | - | 0,2 | 0,2 | 0,2 | 0,2 | 0,2 |
| | kcat_68 | $X_{27}$ | 1740 | 0,2 | 10 | - | 1 | 1 | 1 | 1 | 1 |
| | kcat_11/ kcat_16 | $X_{26}/X_{32}$ | 2.22 | - | - | - | 2.22 | 2.22 | 2.22 | 2.22 | 2.22 |
| | Piruvate (mM) | $X_{17}$ | 2,67 | 0,9 | 1,1 | - | 0,9 | 0,9 | 0,9 | 0,9 | 0,9 |
| | G3P | $X_{18}$ | 0,041 | 0,9 | 1,1 | - | 0,9 | 0,9 | 0,9 | 0,9 | 0,9 |
| | ATP | $X_{19}$ | 3 | 0,9 | 1,1 | - | 0,9 | 0,9 | 0,9 | 0,9 | 0,9 |
| | NADPH | $X_{22}$ | 0,12 | 0,9 | 1,1 | - | 0,9 | 0,9 | 0,9 | 0,9 | 0,9 |
| | CTP | $X_{25}$ | 2,7 | 0,9 | 1,1 | - | 0,9 | 0,9 | 0,9 | 0,9 | 0,9 |
| | FldA | $X_{71}$ | 0,003015 | 0,9 | 1,1 | - | 0,9 | 0,9 | 0,9 | 0,9 | 0,9 |
| | Km_1_68 (mM) | $X_{38}$ | 0,045 | 0,9 | 1,1 | - | 1 | 1 | 1 | 1 | 1 |
| | Km_2_12 | $X_{39}$ | 0,15 | 0,9 | 1,1 | - | 1 | 1 | 1 | 1 | 1 |
| | Km_3_14 | $X_{40}$ | 0,339 | 0,9 | 1,1 | - | 1 | 1 | 1 | 1 | 1 |
| | Km_4_15 | $X_{41}$ | 0,0095 | 0,9 | 1,1 | - | 1,1 | 1,1 | 1,1 | 1,1 | 1,1 |
| | Km_4_16 | $X_{46}$ | 0,0013 | 0,9 | 1,1 | - | 1 | 1 | 1,1 | 1,1 | 1,1 |
| | Km_4_11 | $X_{56}$ | 0,0103 | 0,9 | 1,1 | - | 1 | 0,9 | 0,9 | 0,9 | 0,9 |
| | Km_5_15 | $X_{42}$ | 0,0143 | 0,9 | 1,1 | - | 0,9 | 0,90 | 0,9 | 0,9 | 0,9 |
| | Km_5_16 | $X_{47}$ | 0,0293 | 0,9 | 1,1 | - | 1 | 1 | 1,1 | 1,1 | 1,1 |
| | Km_6_20 | $X_{43}$ | 0,00314 | 0,9 | 1,1 | - | 1 | 1 | 1 | 1 | 1 |
| | Km_7_13 | $X_{60}$ | 0,00495 | 0,9 | 1,1 | - | 1 | 1 | 1 | 1 | 1 |
| | Km_7_23 | $X_{44}$ | 0,56 | 0,9 | 1,1 | - | 1 | 1 | 1 | 1,1 | 1,1 |

| | | | | | | | | | | | |
|---|---|---|---|---|---|---|---|---|---|---|---|
| | Km_8_21 | $X_{45}$ | 0,03 | 0,9 | 1,1 | - | 1 | 1 | 1 | 1 | 1 |
| | Km_9_11 | $X_{37}$ | 0,0055 | 0,9 | 1,1 | - | 1 | 0,9 | 0,9 | 0,9 | 0,9 |
| | Km_17_24 | $X_{48}$ | 0,134 | 0,9 | 1,1 | - | 1,1 | 1,1 | 1,1 | 1,1 | 1,1 |
| | Km_18_24 | $X_{49}$ | 0,041 | 0,9 | 1,1 | - | 1,1 | 1,05 | 1,1 | 1,1 | 1,1 |
| | Km_19_12 | $X_{50}$ | 0,42 | 0,9 | 1,1 | - | 1 | 1 | 1 | 1 | 1 |
| | Km_71_21 | $X_{51}$ | 0,012 | 0,9 | 1,1 | - | 1 | 1 | 1 | 1 | 1 |
| | Km_71_23 | $X_{52}$ | 0,012 | 0,9 | 1,1 | - | 1 | 1 | 1 | 0,9 | 0,9 |
| | Km_22_68 | $X_{53}$ | 0,0005 | 0,9 | 1,1 | - | 1 | 1 | 1 | 1 | 1 |
| | Km_25_20 | $X_{54}$ | 0,76 | 0,9 | 1,1 | - | 1 | 1 | 1 | 1 | 1 |
| Time-independent | K_10_55(mM) | $X_{55}$ | 0,05 | 0,9 | 1,1 | - | 0,9 | 0,9 | 0,9 | 0,9 | 0,9 |
| | K_4_58 | $X_{58}$ | 0,005 | 0,9 | 1,1 | - | 1,01 | 1,1 | 0,98 | 0,9 | 0,9 |
| | K_9_72 | $X_{72}$ | 0,005 | 0,9 | 1,1 | - | 1,1 | 1,1 | 1,1 | 1,1 | 1,1 |
| | Ki_4_24 | $X_{57}$ | 0,0654 | 1 | 1 | - | 1 | 1 | 1 | 1 | 1 |
| | Ki_5_24 | $X_{59}$ | 0,0813 | 1 | 1 | - | 1 | 1 | 1 | 1 | 1 |
| | Ka_6_14 | $X_{69}$ | 0,00339 | 0,9 | 1,1 | - | 1 | 1 | 1 | 1 | 1 |
| | $\alpha_6$ (adimensional) | $X_{61}$ | 1 | 0,9 | 1,1 | - | 1 | 1 | 1 | 1 | 1 |
| | $\beta_6$(adimensional) | $X_{63}$ | 2000 | 0,9 | 1,1 | - | 1 | 1 | 1 | 1 | 1 |
| | Kd_9_11 | $X_{62}$ | 0,0714 | 0,9 | 1,1 | - | 1 | 1 | 0,9 | 0,9 | 0,9 |
| | Kd_17_24 | $X_{64}$ | 0,0426 | 0,9 | 1,1 | - | 1,1 | 0,9 | 1,1 | 1,1 | 1,1 |
| | Kd_22_68 | $X_{65}$ | 0,000333333 | 0,9 | 1,1 | - | 1 | 1 | 1 | 1 | 1 |
| | Kd_71_23 | $X_{66}$ | 0,0005 | 0,9 | 1,1 | - | 1 | 1 | 1 | 1,1 | 1,1 |
| | Kd_71_21 | $X_{67}$ | 0,0005 | 0,9 | 1,1 | - | 1 | 1 | 1 | 1 | 1 |
| | hill_7_13 (adimensional) | $X_{70}$ | 1 | 0,9 | 1,1 | - | 1 | 1 | 1 | 1 | 1 |
| | $(v_{4,11})_{opt}/\ (v_{4,11})_{basal}$ | | | | | NF | 6,10 | 6,77 | 8,53 | 8,74 | 8,74 |

***a***) Time dependent metabolites ($X_1$... $X_{10}$) ranges between 0.1 and 2.5 times its respective basal values

***b***) External variables ($X_{17}$, $X_{18}$, $X_{22}$, $X_{25}$, and $X_{71}$) allowed varying a 10% around their respective basal values.

***c***) The kcat's ($X_{27}$...$X_{31}$, $X_{33}$... $X_{36}$) ranges between 0.2 and 10 times their basal values at exception of IspA kcat's$X_{26}$ and $X_{32}$ with upper level 1.2 times their respective basal values

***d***) The Km's, Kd's, Ki's, and $K_{0,5}$'s variables are allowed be modified around ± 10 % their corresponding basal values.

***e***) Cellular economy factor p = 1

***f***) The $v_{9,11}$ basal flux was:

$$v_{9,11} = \frac{V_{max_{11}} \times X_4 \times X_9}{\left(K_{m_{4,11}} \times X_9 + K_{m_{9,11}} \times X_4 + X_4 \times X_9 + K_{d_{9,11}} \times K_{m_{9,11}}\right)} = 2.83 \times 10^{-4}\ mM \cdot min^{-1}$$

***g***) NF: no feasible solution

**Table S3.2a**. (cont…)

| | | | | | | Opt./Basal | | | | |
|---|---|---|---|---|---|---|---|---|---|---|
| | | | **Basal** | **LB** | **UB** | **7** | **8** | **9** | **10** | **11** |
| Time-dependent | DXP (mM) | $X_{1}$ | 0,002519342 | 0,1 | 2,5 | 1,74 | 1,74 | 0,17 | 0,10 | 2,50 |
| | CDP_MEP | $X_{2}$ | 0,002135892 | 0,1 | 2,5 | 2,26 | 2,26 | 2,50 | 2,50 | 2,50 |
| | MEcDP | $X_{3}$ | 0,001552085 | 0,1 | 2,5 | 1,24 | 1,24 | 2,50 | 2,50 | 2,50 |
| | IPP | $X_{4}$ | 0,000396505 | 0,1 | 2,5 | 2,50 | 2,50 | 2,50 | 2,50 | 2,50 |
| | DMAPP | $X_{5}$ | 0,000496462 | 0,1 | 2,5 | 1,12 | 1,12 | 1,12 | 1,12 | 1,12 |
| | MEP | $X_{6}$ | 0,000125026 | 0,1 | 2,5 | 1,76 | 1,76 | 1,76 | 2,50 | 0,14 |
| | $MEC_{(c)}$ | $X_{7}$ | 0,000893951 | 0,1 | 2,5 | 1,20 | 1,20 | 1,20 | 0,67 | 0,10 |
| | HMBPP | $X_{8}$ | 3,84307E-05 | 0,1 | 2,5 | 0,37 | 0,37 | 1,65 | 1,65 | 2,08 |
| | GPP | $X_{9}$ | 0,000390628 | 0,1 | 2,5 | 2,50 | 2,50 | 2,50 | 2,50 | 2,50 |
| | $MEC_{(e)}$ | $X_{10}$ | 0,02709473 | 0,1 | 2,5 | 2,50 | 2,50 | 2,50 | 1,87 | 2,50 |
| | kcat_11 ($min^{-1}$) | $X_{26}$ | 1,7316 | 0,2 | 1,2 | 1,17 | 1,17 | 1,17 | 1,17 | 1,17 |
| | kcat_12 | $X_{28}$ | 1010 | 0,2 | 10 | 1 | 1 | 0,99 | 0,61 | 0,82 |
| | kcat_13 | $X_{29}$ | 600 | 0,2 | 10 | 1 | 1 | 1 | 1 | 7,52 |
| | kcat_14 | $X_{30}$ | 80,7 | 0,2 | 10 | 1 | 1 | 0,55 | 0,24 | 4,97 |
| | kcat_15 | $X_{31}$ | 19,8 | 0,2 | 10 | 10 | 10 | 10 | 10 | 10 |
| | kcat_16 | $X_{32}$ | 0,78 | 0,2 | 1,2 | 1,17 | 1,17 | 1,17 | 1,17 | 1,17 |
| | kcat_20 | $X_{33}$ | 1554 | 0,2 | 10 | 1 | 1 | 1 | 0,40 | 10 |
| | kcat_21 | $X_{34}$ | 27,6 | 0,2 | 10 | 1 | 1 | 0,2 | 0,2 | 0,20 |
| | kcat_23 | $X_{35}$ | 23,7 | 0,2 | 10 | 0,33 | 0,33 | 0,33 | 0,59 | 3,19 |
| | kcat_24 | $X_{36}$ | 100,3 | 0,2 | 10 | 0,47 | 0,47 | 0,47 | 0,29 | 0,52 |
| | kcat_68 | $X_{27}$ | 1740 | 0,2 | 10 | 1 | 1 | 10 | 10 | 0,64 |
| | kcat_11/ kcat_16 | $X_{26}/X_{32}$ | 2.22 | - | - | 2.22 | 2.22 | 2.22 | 2.22 | 1,17 |
| Time-independent | Piruvate (mM) | $X_{17}$ | 2,67 | 0,9 | 1,1 | 0,9 | 0,9 | 0,9 | 0,9 | 0,90 |
| | G3P | $X_{18}$ | 0,041 | 0,9 | 1,1 | 0,9 | 0,9 | 0,9 | 0,9 | 0,90 |
| | ATP | $X_{19}$ | 3 | 0,9 | 1,1 | 0,9 | 0,9 | 1,1 | 1,1 | 1,1 |
| | NADPH | $X_{22}$ | 0,12 | 0,9 | 1,1 | 0,9 | 0,9 | 0,9 | 0,9 | 1,1 |
| | CTP | $X_{25}$ | 2,7 | 0,9 | 1,1 | 0,9 | 0,9 | 0,9 | 1,1 | 1,1 |
| | FldA | $X_{71}$ | 0,003015 | 0,9 | 1,1 | 1,1 | 1,1 | 1,1 | 1,1 | 1,1 |
| | Km_1_68 (mM) | $X_{38}$ | 0,045 | 0,9 | 1,1 | 1 | 1 | 1,1 | 1,1 | 0,9 |
| | Km_2_12 | $X_{39}$ | 0,15 | 0,9 | 1,1 | 1 | 1 | 1,1 | 1,1 | 0,90 |
| | Km_3_14 | $X_{40}$ | 0,339 | 0,9 | 1,1 | 1 | 1 | 1,1 | 1,1 | 0,90 |
| | Km_4_15 | $X_{41}$ | 0,0095 | 0,9 | 1,1 | 1,1 | 1,1 | 1,1 | 1,1 | 1,10 |
| | Km_4_16 | $X_{46}$ | 0,0013 | 0,9 | 1,1 | 1,1 | 1,1 | 1,1 | 1,1 | 1,10 |
| | Km_4_11 | $X_{56}$ | 0,0103 | 0,9 | 1,1 | 0,9 | 0,9 | 0,9 | 0,9 | 0,90 |
| | Km_5_15 | $X_{42}$ | 0,0143 | 0,9 | 1,1 | 0,9 | 0,9 | 0,9 | 0,9 | 0,90 |
| | Km_5_16 | $X_{47}$ | 0,0293 | 0,9 | 1,1 | 1,1 | 1,1 | 1,1 | 1,1 | 1,10 |
| | Km_6_20 | $X_{43}$ | 0,00314 | 0,9 | 1,1 | 1 | 1 | 1 | 0,9 | 0,90 |
| | Km_7_13 | $X_{60}$ | 0,00495 | 0,9 | 1,1 | 1 | 1 | 1 | 1 | 0,90 |
| | Km_7_23 | $X_{44}$ | 0,56 | 0,9 | 1,1 | 1,1 | 1,1 | 1,1 | 1,1 | 0,90 |
| | Km_8_21 | $X_{45}$ | 0,03 | 0,9 | 1,1 | 1 | 1 | 0,9 | 0,9 | 1,10 |
| | Km_9_11 | $X_{37}$ | 0,0055 | 0,9 | 1,1 | 0,9 | 0,9 | 0,9 | 0,9 | 0,90 |

| | | | | | | | | | | |
|---|---|---|---|---|---|---|---|---|---|---|
| | Km_17_24 | $X_{48}$ | 0,134 | 0,9 | 1,1 | 1,1 | 1,1 | 1,1 | 1,1 | 1,10 |
| | Km_18_24 | $X_{49}$ | 0,041 | 0,9 | 1,1 | 0,9 | 0,9 | 0,9 | 0,9 | 1,10 |
| | Km_19_12 | $X_{50}$ | 0,42 | 0,9 | 1,1 | 1 | 1 | 1,1 | 1,1 | 0,90 |
| | Km_71_21 | $X_{51}$ | 0,012 | 0,9 | 1,1 | 1 | 1 | 0,9 | 0,9 | 1,10 |
| | Km_71_23 | $X_{52}$ | 0,012 | 0,9 | 1,1 | 0,96 | 0,96 | 0,9 | 0,9 | 0,90 |
| | Km_22_68 | $X_{53}$ | 0,0005 | 0,9 | 1,1 | 1 | 1 | 1,1 | 1,1 | 0,90 |
| | Km_25_20 | $X_{54}$ | 0,76 | 0,9 | 1,1 | 1 | 1 | 1 | 1,1 | 0,90 |
| | K_10_55 (mM) | $X_{55}$ | 0,05 | 0,9 | 1,1 | 1,1 | 1,1 | 1,1 | 0,9 | 1,10 |
| | K_4_58 | $X_{58}$ | 0,005 | 0,9 | 1,1 | 0,9 | 0,9 | 0,9 | 0,9 | 0,90 |
| | K_9_72 | $X_{72}$ | 0,005 | 0,9 | 1,1 | 1,1 | 1,1 | 1,1 | 1,1 | 1,10 |
| | Ki_4_24 | $X_{57}$ | 0,0654 | 1 | 1 | 1 | 1 | 1 | 1 | 1 |
| Time-independent | Ki_5_24 | $X_{59}$ | 0,0813 | 1 | 1 | 1 | 1 | 1 | 1 | 1 |
| | Ka_6_14 | $X_{69}$ | 0,00339 | 0,9 | 1,1 | 1 | 1 | 1 | 1 | 1 |
| | $\alpha_6$ (adimensional) | $X_{61}$ | 1 | 0,9 | 1,1 | 1 | 1 | 1 | 1 | 1 |
| | $\beta_6$(adimensional) | $X_{63}$ | 2000 | 0,9 | 1,1 | 1 | 1 | 1 | 1 | 1 |
| | Kd_9_11 | $X_{62}$ | 0,0714 | 0,9 | 1,1 | 0,9 | 0,9 | 0,9 | 0,9 | 1,10 |
| | Kd_17_24 | $X_{64}$ | 0,0426 | 0,9 | 1,1 | 0,9 | 0,9 | 0,9 | 0,9 | 0,9 |
| | Kd_22_68 | $X_{65}$ | 0,000333333 | 0,9 | 1,1 | 1 | 1 | 1,1 | 1,1 | 1,10 |
| | Kd_71_23 | $X_{66}$ | 0,0005 | 0,9 | 1,1 | 0,96 | 0,96 | 0,9 | 0,9 | 0,90 |
| | Kd_71_21 | $X_{67}$ | 0,0005 | 0,9 | 1,1 | 1 | 1 | 0,9 | 0,9 | 0,90 |
| | hill_7_13(adimensional) | $X_{70}$ | 1 | 0,9 | 1,1 | 1 | 1 | 1 | 1 | 1,10 |
| | $(\nu_{4,11})_{opt}/(\nu_{4,11})_{basal}$ | | | | | 8,74 | 8,74 | 8,74 | 8,74 | 8,74 |

**Table S3.2b.** Specific constants (kcat/Km) fold change respect their basal values for data from Table S3.2a

| | | | Opt./Basal | | | | |
|---|---|---|---|---|---|---|---|
| **kcat / Km** | **Model symbols** | **Basal** | **2** | **3** | **4** | **5** | **6** |
| kcat_11/Km_4_11 | $X_{26}/ X_{56}$ | 168,12 | 1 | 1,11 | 1,27 | 1,30 | 1,30 |
| kcat_11/Km_9_11 | $X_{26} / X_{37}$ | 314,84 | 1 | 1,11 | 1,27 | 1,30 | 1,30 |
| kcat_12 / Km_2_12 | $X_{28} / X_{39}$ | 6733,33 | 1 | 1 | 1 | 1 | 1 |
| kcat_12 / Km_19_12 | $X_{28} / X_{50}$ | 2404,76 | 1 | 1 | 1 | 1 | 1 |
| kcat_13 / Km_7_13 | $X_{29} / X_{60}$ | 121212,12 | 1 | 1 | 1 | 1 | 1 |
| kcat_14 / Km_3_14 | $X_{30} / X_{40}$ | 238,05 | 1 | 1 | 1 | 1 | 1 |
| kcat_15 / Km_4_15 | $X_{31} / X_{41}$ | 2084,21 | 8,46 | 9,09 | 9,09 | 9,09 | 9,09 |
| kcat_15 / Km_5_15 | $X_{31} / X_{42}$ | 1384,62 | 10,35 | 11,11 | 11,11 | 11,11 | 11,11 |
| kcat_16 / Km_4_16 | $X_{32} / X_{46}$ | 600,00 | 1 | 1 | 1,04 | 1,06 | 1,06 |
| kcat_16 / Km_5_16 | $X_{32}/ X_{47}$ | 26,62 | 1 | 1 | 1,04 | 1,06 | 1,06 |
| kcat_20 / Km_6_20 | $X_{33} / X_{43}$ | 494904,46 | 1 | 1 | 1 | 1 | 1 |
| kcat_20 / Km_25_20 | $X_{33} / X_{54}$ | 2044,74 | 1 | 1 | 1 | 1 | 1 |
| kcat_21 / Km_8_21 | $X_{34} / X_{45}$ | 920,00 | 1 | 1 | 1 | 1 | 1 |
| kcat_21 / Km_71_21 | $X_{34} / X_{51}$ | 2300,00 | 1 | 1 | 1 | 1 | 1 |
| kcat_23 / Km_7_23 | $X_{35} / X_{44}$ | 42,32 | 1 | 1 | 1 | 0,98 | 0,98 |
| kcat_23 / Km_71_23 | $X_{35} / X_{52}$ | 1975,00 | 1 | 1 | 1 | 1,20 | 1,20 |
| kcat_24 / Km_17_24 | $X_{36} / X_{48}$ | 748,51 | 0,18 | 0,18 | 0,18 | 0,18 | 0,18 |
| kcat_24/ Km_18_24 | $X_{36} / X_{49}$ | 2446,34 | 0,18 | 0,19 | 0,18 | 0,18 | 0,18 |
| kcat_68/ Km_1_68 | $X_{27} / X_{38}$ | 38666,67 | 1 | 1 | 1 | 1 | 1 |
| kcat_68/ Km_22_68 | $X_{27} / X_{53}$ | 3480000,00 | 1 | 1 | 1 | 1 | 1 |

**Table S3.2b.** (cont..)

| kcat / Km | Model symbols | Basal | Opt./Basal 7 | 8 | 9 | 10 | 11 |
|---|---|---|---|---|---|---|---|
| kcat_11/Km_4_11 | $X_{26}$/ $X_{56}$ | 168,12 | 1,30 | 1,30 | 1,30 | 1,30 | 2,50 |
| kcat_11/Km_9_11 | $X_{26}$ / $X_{37}$ | 314,84 | 1,30 | 1,30 | 1,30 | 1,30 | 2,50 |
| kcat_12 / Km_2_12 | $X_{28}$ / $X_{39}$ | 6733,33 | 1 | 1 | 0,90 | 0,55 | 2,50 |
| kcat_12 / Km_19_12 | $X_{28}$ / $X_{50}$ | 2404,76 | 1 | 1 | 0,90 | 0,55 | 2,50 |
| kcat_13 / Km_7_13 | $X_{29}$ / $X_{60}$ | 121212,12 | 1 | 1 | 1 | 1 | 1,12 |
| kcat_14 / Km_3_14 | $X_{30}$ / $X_{40}$ | 238,05 | 1 | 1 | 0,50 | 0,22 | 0,14 |
| kcat_15 / Km_4_15 | $X_{31}$ / $X_{41}$ | 2084,21 | 9,09 | 9,09 | 9,09 | 9,09 | 0,10 |
| kcat_15 / Km_5_15 | $X_{31}$ / $X_{42}$ | 1384,62 | 11,11 | 11,11 | 11,11 | 11,11 | 2,08 |
| kcat_16 / Km_4_16 | $X_{32}$ / $X_{46}$ | 600,00 | 1,06 | 1,06 | 1,06 | 1,06 | 2,50 |
| kcat_16 / Km_5_16 | $X_{32}$/ $X_{47}$ | 26,62 | 1,06 | 1,06 | 1,06 | 1,06 | 2,50 |
| kcat_20 / Km_6_20 | $X_{33}$ / $X_{43}$ | 494904,46 | 1 | 1 | 1 | 0,44 | 1 |
| kcat_20 / Km_25_20 | $X_{33}$ / $X_{54}$ | 2044,74 | 1 | 1 | 1 | 0,36 | 1 |
| kcat_21 / Km_8_21 | $X_{34}$ / $X_{45}$ | 920,00 | 1 | 1 | 0,22 | 0,22 | 1 |
| kcat_21 / Km_71_21 | $X_{34}$ / $X_{51}$ | 2300,00 | 1 | 1 | 0,22 | 0,22 | 1 |
| kcat_23 / Km_7_23 | $X_{35}$ / $X_{44}$ | 42,32 | 0,30 | 0,30 | 0,30 | 0,53 | 1 |
| kcat_23 / Km_71_23 | $X_{35}$ / $X_{52}$ | 1975,00 | 0,35 | 0,35 | 0,37 | 0,65 | 1 |
| kcat_24 / Km_17_24 | $X_{36}$ / $X_{48}$ | 748,51 | 0,43 | 0,43 | 0,43 | 0,26 | 1 |
| kcat_24/ Km_18_24 | $X_{36}$ / $X_{49}$ | 2446,34 | 0,53 | 0,53 | 0,53 | 0,32 | 1 |
| kcat_68/ Km_1_68 | $X_{27}$ / $X_{38}$ | 38666,67 | 1 | 1 | 9,09 | 9,09 | 1 |
| kcat_68/ Km_22_68 | $X_{27}$ / $X_{53}$ | 3480000,00 | 1 | 1 | 9,09 | 9,09 | 1 |

**Table S3.3a**. Optimization of the *k*cat's allowing modifications in the Km's, Kd's, Ki's, and $K_{0,5}$'s. For the IspA kcat is allowed only the modulation of kcat_16 ($X_{32}$) while the kcat_11 ($X_{26}$) is anchored in their basal value.

| | | | | | | **Opt./Basal** | | | | | |
|---|---|---|---|---|---|---|---|---|---|---|---|
| | | | **Basal** | **LB** | **UB** | **6** | **7** | **8** | **9** | **10** | **11** |
| Time-dependent | DXP (mM) | $X_1$ | 0,002519342 | 0,8 | 1,2 | 0,80 | 0,80 | 0,80 | 0,80 | 0,80 | 0,80 |
| | CDP_MEP | $X_2$ | 0,002135892 | 0,8 | 1,2 | 1,07 | 1,07 | 1,07 | 1,07 | 1,07 | 0,80 |
| | MEcDP | $X_3$ | 0,001552085 | 0,8 | 1,2 | 1,20 | 1,20 | 0,80 | 1,20 | 1,20 | 0,80 |
| | IPP | $X_4$ | 0,000396505 | 0,8 | 1,2 | 1,14 | 1,14 | 1,20 | 1,20 | 1,20 | 1,20 |
| | DMAPP | $X_5$ | 0,000496462 | 0,8 | 1,2 | 1,20 | 1,20 | 0,80 | 1,20 | 1,20 | 1,20 |
| | MEP | $X_6$ | 0,000125026 | 0,8 | 1,2 | 0,84 | 0,84 | 0,81 | 0,84 | 0,84 | 0,80 |
| | $MEC_{(c)}$ | $X_7$ | 0,000893951 | 0,8 | 1,2 | 0,80 | 0,80 | 0,80 | 0,80 | 0,80 | 1,05 |
| | HMBPP | $X_8$ | 3,84307E-05 | 0,8 | 1,2 | 0,97 | 0,98 | 1,06 | 1,00 | 1,00 | 1,06 |
| | GPP | $X_9$ | 0,000390628 | 0,8 | 1,2 | 1,20 | 1,20 | 1,20 | 1,20 | 1,20 | 1,20 |
| | $MEC_{(e)}$ | $X_{10}$ | 0,02709473 | 0,8 | 1,2 | 1,20 | 1,20 | 1,20 | 1,20 | 1,20 | 1,17 |
| Time-independent | kcat_11 ($min^{-1}$) | $X_{26}$ | 1,7316 | **1** | **1** | **1** | **1** | **1** | **1** | **1** | **1** |
| | kcat_12 | $X_{28}$ | 1010 | 0,2 | 10 | 1 | 1 | 1 | 1 | 1 | 1 |
| | kcat_13 | $X_{29}$ | 600 | 0,2 | 10 | 0,62 | 0,62 | 0,62 | 0,62 | 0,62 | 0,39 |
| | kcat_14 | $X_{30}$ | 80,7 | 0,2 | 10 | 1 | 1 | 2 | 1 | 1 | 1 |
| | kcat_15 | $X_{31}$ | 19,8 | 0,2 | 10 | 1,00 | 1,00 | 5,98 | 0,68 | 0,68 | 1,33 |
| | kcat_16 | $X_{32}$ | 0,78 | **0,2** | **10** | **0,44** | **0,44** | **0,64** | **0,44** | **0,44** | **0,44** |
| | kcat_20 | $X_{33}$ | 1554 | 0,2 | 10 | 1 | 1 | 1 | 1 | 1 | 1 |
| | kcat_21 | $X_{34}$ | 27,6 | 0,2 | 10 | 0,20 | 0,20 | 0,20 | 0,20 | 0,20 | 0,20 |
| | kcat_23 | $X_{35}$ | 23,7 | 0,2 | 10 | 0,20 | 0,20 | 0,21 | 0,20 | 0,20 | 0,20 |
| | kcat_24 | $X_{36}$ | 100,3 | 0,2 | 10 | 0,23 | 0,23 | 0,23 | 0,23 | 0,23 | 0,20 |
| | kcat_68 | $X_{27}$ | 1740 | 0,2 | 10 | 1 | 1 | 1 | 1 | 1 | 1 |
| | kcat_11/ kcat_16 | $X_{26}/X_{32}$ | 2.22 | - | - | 5,09 | 5,06 | 3,47 | 5,07 | 5,07 | 5,07 |
| | Piruvate (mM) | $X_{17}$ | 2,67 | 0,9 | 1,1 | 0,90 | 0,90 | 0,90 | 0,90 | 0,90 | 0,90 |
| | G3P | $X_{18}$ | 0,041 | 0,9 | 1,1 | 0,90 | 0,90 | 0,90 | 0,90 | 0,90 | 0,90 |
| | ATP | $X_{19}$ | 3 | 0,9 | 1,1 | 0,90 | 0,90 | 0,90 | 0,90 | 0,90 | 0,90 |
| | NADPH | $X_{22}$ | 0,12 | 0,9 | 1,1 | 0,90 | 0,90 | 0,90 | 0,90 | 0,90 | 0,90 |
| | CTP | $X_{25}$ | 2,7 | 0,9 | 1,1 | 0,90 | 0,90 | 0,90 | 0,90 | 0,90 | 0,90 |
| | FldA | $X_{71}$ | 0,003015 | 0,9 | 1,1 | 0,90 | 0,90 | 0,90 | 0,90 | 0,90 | 0,90 |
| | Km_1_68 (mM) | $X_{38}$ | 0,045 | 0,9 | 1,1 | 1 | 1 | 1 | 1,1 | 1,1 | 0,9 |
| | Km_2_12 | $X_{39}$ | 0,15 | 0,9 | 1,1 | 1 | 1 | 1 | 1 | 1 | 0,9 |
| | Km_3_14 | $X_{40}$ | 0,339 | 0,9 | 1,1 | 1 | 1 | 1,1 | 1 | 1 | 0,9 |
| | Km_4_15 | $X_{41}$ | 0,0095 | 0,9 | 1,1 | 1 | 1 | 1,1 | 1,1 | 1,1 | 0,9 |
| | Km_4_16 | $X_{46}$ | 0,0013 | 0,9 | 1,1 | 0,9 | 0,9 | 0,9 | 0,9 | 0,9 | 0,9 |
| | Km_4_11 | $X_{56}$ | 0,0103 | 0,9 | 1,1 | 1 | 0,9 | 0,9 | 0,9 | 0,9 | 0,9 |
| | Km_5_15 | $X_{42}$ | 0,0143 | 0,9 | 1,1 | 1 | 1 | 1,1 | 0,9 | 0,9 | 0,9 |
| | Km_5_16 | $X_{47}$ | 0,0293 | 0,9 | 1,1 | 0,9 | 0,9 | 0,9 | 0,9 | 0,9 | 0,9 |
| | Km_6_20 | $X_{43}$ | 0,00314 | 0,9 | 1,1 | 1,1 | 1,1 | 1 | 1,1 | 1,1 | 0,9 |
| | Km_7_13 | $X_{60}$ | 0,00495 | 0,9 | 1,1 | 0,9 | 0,9 | 0,9 | 0,9 | 0,9 | 0,9 |

| | | | | | | | | | | | |
|---|---|---|---|---|---|---|---|---|---|---|---|
| | Km_7_23 | $X_{44}$ | 0,56 | 0,9 | 1,1 | 0,95 | 0,94 | 0,90 | 0,92 | 0,92 | 1,1 |
| | Km_8_21 | $X_{45}$ | 0,03 | 0,9 | 1,1 | 1,1 | 1,1 | 1,1 | 1,1 | 1,1 | 1,1 |
| | Km_9_11 | $X_{37}$ | 0,0055 | 0,9 | 1,1 | 1 | 0,9 | 0,9 | 0,9 | 0,9 | 0,9 |
| | Km_17_24 | $X_{48}$ | 0,134 | 0,9 | 1,1 | 0,9 | 1,1 | 1,1 | 1,1 | 1,1 | 1,1 |
| | Km_18_24 | $X_{49}$ | 0,041 | 0,9 | 1,1 | 0,9 | 0,9 | 0,9 | 0,9 | 0,9 | 1,1 |
| | Km_19_12 | $X_{50}$ | 0,42 | 0,9 | 1,1 | 1 | 1 | 1 | 1 | 1 | 0,9 |
| | Km_71_21 | $X_{51}$ | 0,012 | 0,9 | 1,1 | 1,1 | 1,1 | 1,1 | 1,1 | 1,1 | 1,1 |
| | Km_71_23 | $X_{52}$ | 0,012 | 0,9 | 1,1 | 0,9 | 0,9 | 0,9 | 0,9 | 0,9 | 1,1 |
| | Km_22_68 | $X_{53}$ | 0,0005 | 0,9 | 1,1 | 1 | 1 | 1 | 1,1 | 1,1 | 0,9 |
| | Km_25_20 | $X_{54}$ | 0,76 | 0,9 | 1,1 | 1,1 | 1,1 | 1 | 1,1 | 1,1 | 0,9 |
| | K_10_55 (mM) | $X_{55}$ | 0,05 | 0,9 | 1,1 | 1,10 | 1,10 | 1,10 | 1,10 | 1,10 | 0,90 |
| | K_4_58 | $X_{58}$ | 0,005 | 0,9 | 1,1 | 0,90 | 0,90 | 1,10 | 0,90 | 0,90 | 1,10 |
| | K_9_72 | $X_{72}$ | 0,005 | 0,9 | 1,1 | 1,10 | 1,10 | 1,10 | 1,10 | 1,10 | 1,10 |
| | Ki_4_24 | $X_{57}$ | 0,0654 | 1 | 1 | 1 | 1 | 1 | 1 | 1 | 1 |
| | Ki_5_24 | $X_{59}$ | 0,0813 | 1 | 1 | 1 | 1 | 1 | 1 | 1 | 1 |
| Time-independent | Ka_6_14 | $X_{69}$ | 0,00339 | 0,9 | 1,1 | 1 | 1 | 1 | 1 | 1 | 1 |
| | $\alpha_6$ (adimensional) | $X_{61}$ | 1 | 0,9 | 1,1 | 1 | 1 | 1 | 1 | 1 | 1 |
| | $\beta_6$(adimensional) | $X_{63}$ | 2000 | 0,9 | 1,1 | 1 | 1 | 1 | 1 | 1 | 1 |
| | Kd_9_11 | $X_{62}$ | 0,0714 | 0,9 | 1,1 | 1 | 0,9 | 0,9 | 0,9 | 0,9 | 0,9 |
| | Kd_17_24 | $X_{64}$ | 0,0426 | 0,9 | 1,1 | 0,9 | 0,9 | 0,9 | 0,9 | 0,9 | 1,1 |
| | Kd_22_68 | $X_{65}$ | 0,000333333 | 0,9 | 1,1 | 1 | 1 | 1 | 1,1 | 1,1 | 0,9 |
| | Kd_71_23 | $X_{66}$ | 0,0005 | 0,9 | 1,1 | 0,9 | 0,9 | 0,9 | 0,9 | 0,9 | 1,1 |
| | Kd_71_21 | $X_{67}$ | 0,0005 | 0,9 | 1,1 | 1,1 | 1,1 | 1,1 | 1,1 | 1,1 | 1,1 |
| | hill_7_13 (adimensional) | $X_{70}$ | 1 | 0,9 | 1,1 | 1 | 1 | 1 | 1 | 1 | 1 |
| | $(v_{4,11})_{opt}/\ (v_{4,11})_{basal}$ | | | | | 1,36 | 1,67 | 1,77 | 1,77 | 1,77 | 1,77 |

***a)*** Time dependent metabolites ($X_1$... $X_{10}$) ranges between 0.8 and 1.2 times its respective basal values

***b)*** External variables ($X_{17}$, $X_{18}$, $X_{22}$, $X_{25}$, and $X_{71}$) allowed varying a 10% around their respective basal values.

***c)*** The kcat's ($X_{27}$...$X_{31,}$$X_{33}$... $X_{36}$) ranges between 0.2 and 10 times their basal values at exception of IspA kcat_11($X_{26}$) which is anchored in their basal value.

***d)*** Cellular economy factor p = 1

***e)*** The option of the modulation of kcat_11 ($X_{26}$) while kcat_16 ($X_{32}$) is anchored in their basal value does not give feasible solutions.

***f)*** The $v_{9,11}$basal flux was calculated as:

$$v_{9,11} = \frac{V_{max_{11}} \times X_4 \times X_9}{\left(K_{m_{4,11}} \times X_9 + K_{m_{9,11}} \times X_4 + X_4 \times X_9 + K_{d_{9,11}} \times K_{m_{9,11}}\right)} = 2.83 \times 10^{-4}\ mM \cdot min^{-1}$$

**Table S3.3b.** Specific constants (kcat/Km) fold change respect their basal values for data from Table S3.3a

| | | | Opt./Basal | | | | | |
|---|---|---|---|---|---|---|---|---|
| **kcat / Km** | **Model symbols** | **Basal** | **6** | **7** | **8** | **9** | **10** | **11** |
| kcat_11/Km_4_11 | $X_{26}/X_{56}$ | 168,12 | 1 | 1,11 | 1,11 | 1,11 | 1,11 | 1,11 |
| kcat_11/Km_9_11 | $X_{26}/X_{37}$ | 314,84 | 1 | 1,11 | 1,11 | 1,11 | 1,11 | 1,11 |
| kcat_12 / Km_2_12 | $X_{28}/X_{39}$ | 6733,33 | 1 | 1 | 1 | 1 | 1 | 1,06 |
| kcat_12 / Km_19_12 | $X_{28}/X_{50}$ | 2404,76 | 1 | 1 | 1 | 1 | 1 | 1,06 |
| kcat_13 / Km_7_13 | $X_{29}/X_{60}$ | 121212,12 | 0,69 | 0,69 | 0,69 | 0,69 | 0,69 | 0,44 |
| kcat_14 / Km_3_14 | $X_{30}/X_{40}$ | 238,05 | 1 | 1 | 1,55 | 1 | 1 | 1,26 |
| kcat_15 / Km_4_15 | $X_{31}/X_{41}$ | 2084,21 | 1 | 1 | 5,44 | 0,62 | 0,62 | 1,47 |
| kcat_15 / Km_5_15 | $X_{31}/X_{42}$ | 1384,62 | 1 | 1 | 5,44 | 0,76 | 0,76 | 1,47 |
| kcat_16 / Km_4_16 | $X_{32}/X_{46}$ | 600,00 | 0,48 | 0,49 | 0,71 | 0,49 | 0,49 | 0,49 |
| kcat_16 / Km_5_16 | $X_{32}/X_{47}$ | 26,62 | 0,48 | 0,49 | 0,71 | 0,49 | 0,49 | 0,49 |
| kcat_20 / Km_6_20 | $X_{33}/X_{43}$ | 494904,46 | 0,96 | 0,96 | 1 | 0,96 | 0,96 | 0,81 |
| kcat_20 / Km_25_20 | $X_{33}/X_{54}$ | 2044,74 | 0,96 | 0,96 | 1 | 0,96 | 0,96 | 0,81 |
| kcat_21 / Km_8_21 | $X_{34}/X_{45}$ | 920,00 | 0,18 | 0,18 | 0,18 | 0,18 | 0,18 | 0,18 |
| kcat_21 / Km_71_21 | $X_{34}/X_{51}$ | 2300,00 | 0,18 | 0,18 | 0,18 | 0,18 | 0,18 | 0,18 |
| kcat_23 / Km_7_23 | $X_{35}/X_{44}$ | 42,32 | 0,21 | 0,21 | 0,23 | 0,22 | 0,22 | 0,18 |
| kcat_23 / Km_71_23 | $X_{35}/X_{52}$ | 1975,00 | 0,22 | 0,22 | 0,23 | 0,22 | 0,22 | 0,18 |
| kcat_24 / Km_17_24 | $X_{36}/X_{48}$ | 748,51 | 0,25 | 0,21 | 0,21 | 0,21 | 0,21 | 0,18 |
| kcat_24/ Km_18_24 | $X_{36}/X_{49}$ | 2446,34 | 0,25 | 0,25 | 0,25 | 0,25 | 0,25 | 0,18 |
| kcat_68/ Km_1_68 | $X_{27}/X_{38}$ | 38666,67 | 1 | 1 | 1 | 1 | 1 | 0,80 |
| kcat_68/ Km_22_68 | $X_{27}/X_{53}$ | 3480000,00 | 1 | 1 | 1 | 1 | 1 | 0,80 |

**Table S3.4a**. Optimization of the *K*cat's allowing modifications in the Km's, Kd's, Ki's, and $K_{0,5}$'s. No relationship between kcat_11 ($X_{26}$) and kcat_16 ($X_{32}$).

| | | | Basal | LB | UB | Opt./Basal 6 | 7 | 8 | 9 | 10 | 11 |
|---|---|---|---|---|---|---|---|---|---|---|---|
| Time-dependent | DXP (mM) | $X_1$ | 0,002519342 | 0,8 | 1,2 | 0,80 | 0,80 | 0,80 | 0,80 | 0,80 | 0,80 |
| | CDP_MEP | $X_2$ | 0,002135892 | 0,8 | 1,2 | 1,07 | 1,07 | 1,07 | 0,80 | 0,80 | 0,80 |
| | MEcDP | $X_3$ | 0,001552085 | 0,8 | 1,2 | 1,20 | 1,20 | 1,20 | 0,80 | 0,80 | 0,80 |
| | IPP | $X_4$ | 0,000396505 | 0,8 | 1,2 | 1,14 | 1,11 | 1,20 | 1,20 | 1,20 | 1,20 |
| | DMAPP | $X_5$ | 0,000496462 | 0,8 | 1,2 | 1,20 | 1,20 | 0,80 | 0,80 | 0,80 | 1,20 |
| | MEP | $X_6$ | 0,000125026 | 0,8 | 1,2 | 0,84 | 0,84 | 0,84 | 0,81 | 0,80 | 0,80 |
| | $MEC_{(c)}$ | $X_7$ | 0,000893951 | 0,8 | 1,2 | 0,80 | 0,80 | 0,80 | 0,80 | 0,80 | 1,12 |
| | HMBPP | $X_8$ | 3,84307E-05 | 0,8 | 1,2 | 0,97 | 1,04 | 1,14 | 1,14 | 1,14 | 1,14 |
| | GPP | $X_9$ | 0,000390628 | 0,8 | 1,2 | 1,20 | 1,20 | 1,20 | 1,20 | 1,20 | 1,20 |
| | $MEC_{(e)}$ | $X_{10}$ | 0,02709473 | 0,8 | 1,2 | 1,20 | 1,20 | 1,20 | 1,20 | 1,20 | 1,17 |
| Time-independent | kcat_11 ($min^{-1}$) | $X_{26}$ | 1,7316 | 0,2 | 10 | **1** | **10** | **10** | **10** | **10** | **10** |
| | kcat_12 | $X_{28}$ | 1010 | 0,2 | 10 | 1 | 1 | 1 | 1 | 1 | 1 |
| | kcat_13 | $X_{29}$ | 600 | 0,2 | 10 | 0,62 | 0,62 | 0,62 | 0,62 | 0,62 | 0,37 |
| | kcat_14 | $X_{30}$ | 80,7 | 0,2 | 10 | 1 | 1 | 1 | 2 | 2 | 1 |
| | kcat_15 | $X_{31}$ | 19,8 | 0,2 | 10 | 1,00 | 1,00 | 6,39 | 6,39 | 6,39 | 1,42 |
| | kcat_16 | $X_{32}$ | 0,78 | 0,2 | 10 | **0,44** | **0,47** | **0,68** | **0,68** | **0,68** | **0,47** |
| | kcat_20 | $X_{33}$ | 1554 | 0,2 | 10 | 1 | 1 | 1 | 1 | 1 | 1 |
| | kcat_21 | $X_{34}$ | 27,6 | 0,2 | 10 | 0,20 | 0,20 | 0,20 | 0,20 | 0,20 | 0,20 |
| | kcat_23 | $X_{35}$ | 23,7 | 0,2 | 10 | 0,20 | 0,20 | 0,20 | 0,20 | 0,20 | 0,20 |
| | kcat_24 | $X_{36}$ | 100,3 | 0,2 | 10 | 0,23 | 0,23 | 0,23 | 0,23 | 0,23 | 0,20 |
| | kcat_68 | $X_{27}$ | 1740 | 0,2 | 10 | 1 | 1 | 1 | 1 | 1 | 1 |
| | kcat_11/ kcat_16 | $X_{26}/X_{32}$ | 2.22 | - | - | 5,09 | 47,34 | 32,49 | 32,49 | 32,49 | 47,43 |
| | Piruvate (mM) | $X_{17}$ | 2,67 | 0,9 | 1,1 | 0,90 | 0,90 | 0,90 | 0,90 | 0,90 | 0,90 |
| | G3P | $X_{18}$ | 0,041 | 0,9 | 1,1 | 0,90 | 0,90 | 0,90 | 0,90 | 0,90 | 0,90 |
| | ATP | $X_{19}$ | 3 | 0,9 | 1,1 | 0,90 | 0,90 | 0,90 | 0,90 | 0,90 | 0,90 |
| | NADPH | $X_{22}$ | 0,12 | 0,9 | 1,1 | 0,90 | 0,90 | 0,90 | 0,90 | 0,90 | 0,90 |
| | CTP | $X_{25}$ | 2,7 | 0,9 | 1,1 | 0,90 | 0,90 | 0,90 | 0,90 | 0,90 | 0,90 |
| | FldA | $X_{71}$ | 0,003015 | 0,9 | 1,1 | 0,90 | 0,90 | 0,90 | 0,90 | 0,90 | 0,90 |
| | Km_1_68 (mM) | $X_{38}$ | 0,045 | 0,9 | 1,1 | 1 | 1 | 1 | 1 | 1 | 0,9 |
| | Km_2_12 | $X_{39}$ | 0,15 | 0,9 | 1,1 | 1 | 1 | 1 | 1,1 | 1,1 | 0,9 |
| | Km_3_14 | $X_{40}$ | 0,339 | 0,9 | 1,1 | 1 | 1 | 1 | 1,1 | 1,1 | 0,9 |
| | Km_4_15 | $X_{41}$ | 0,0095 | 0,9 | 1,1 | 1 | 1 | 1,1 | 1,1 | 1,1 | 0,9 |
| | Km_4_16 | $X_{46}$ | 0,0013 | 0,9 | 1,1 | 0,9 | 0,9 | 0,9 | 0,9 | 0,9 | 0,9 |
| | Km_4_11 | $X_{56}$ | 0,0103 | 0,9 | 1,1 | 1 | 0,9 | 0,9 | 0,9 | 0,9 | 0,9 |
| | Km_5_15 | $X_{42}$ | 0,0143 | 0,9 | 1,1 | 1 | 1 | 1,1 | 1,1 | 1,1 | 0,9 |
| | Km_5_16 | $X_{47}$ | 0,0293 | 0,9 | 1,1 | 0,9 | 0,9 | 0,9 | 0,9 | 0,9 | 0,9 |
| | Km_6_20 | $X_{43}$ | 0,00314 | 0,9 | 1,1 | 1,1 | 1,1 | 1,1 | 1 | 1,1 | 0,9 |
| | Km_7_13 | $X_{60}$ | 0,00495 | 0,9 | 1,1 | 0,9 | 0,9 | 0,9 | 0,9 | 0,9 | 0,9 |
| | Km_7_23 | $X_{44}$ | 0,56 | 0,9 | 1,1 | 0,95 | 0,9 | 0,9 | 0,9 | 0,9 | 1,1 |

| | | | | | | | | | | | |
|---|---|---|---|---|---|---|---|---|---|---|---|
| | Km_8_21 | $X_{45}$ | 0,03 | 0,9 | 1,1 | 1,1 | 1,1 | 1,1 | 1,1 | 1,1 | 1,1 |
| | Km_9_11 | $X_{37}$ | 0,0055 | 0,9 | 1,1 | 1 | 0,9 | 0,9 | 0,9 | 0,9 | 0,9 |
| | Km_17_24 | $X_{48}$ | 0,134 | 0,9 | 1,1 | 0,9 | 1,1 | 1,1 | 1,1 | 1,1 | 1,1 |
| | Km_18_24 | $X_{49}$ | 0,041 | 0,9 | 1,1 | 0,9 | 0,9 | 0,9 | 0,9 | 0,9 | 1,1 |
| | Km_19_12 | $X_{50}$ | 0,42 | 0,9 | 1,1 | 1 | 1 | 1 | 1,1 | 1,1 | 0,9 |
| | Km_71_21 | $X_{51}$ | 0,012 | 0,9 | 1,1 | 1,1 | 1,1 | 1,1 | 1,1 | 1,1 | 1,1 |
| | Km_71_23 | $X_{52}$ | 0,012 | 0,9 | 1,1 | 0,9 | 0,9 | 0,9 | 0,9 | 0,9 | 1,1 |
| | Km_22_68 | $X_{53}$ | 0,0005 | 0,9 | 1,1 | 1 | 1 | 1 | 1 | 1 | 0,9 |
| | Km_25_20 | $X_{54}$ | 0,76 | 0,9 | 1,1 | 1,1 | 1,1 | 1,1 | 1 | 1,1 | 0,9 |
| Time-independent | K_10_55 (mM) | $X_{55}$ | 0,05 | 0,9 | 1,1 | 1,10 | 1,10 | 1,10 | 1,10 | 1,10 | 0,90 |
| | K_4_58 | $X_{58}$ | 0,005 | 0,9 | 1,1 | 0,90 | 0,90 | 1,10 | 1,10 | 1,10 | 1,10 |
| | K_9_72 | $X_{72}$ | 0,005 | 0,9 | 1,1 | 1,10 | 1,10 | 1,10 | 1,10 | 1,10 | 1,10 |
| | Ki_4_24 | $X_{57}$ | 0,0654 | 1 | 1 | 1 | 1 | 1 | 1 | 1 | 1 |
| | Ki_5_24 | $X_{59}$ | 0,0813 | 1 | 1 | 1 | 1 | 1 | 1 | 1 | 1 |
| | Ka_6_14 | $X_{69}$ | 0,00339 | 0,9 | 1,1 | 1 | 1 | 1 | 1 | 1 | 1 |
| | $\alpha_6$ (adimensional) | $X_{61}$ | 1 | 0,9 | 1,1 | 1 | 1 | 1 | 1 | 1 | 1 |
| | $\beta_6$(adimensional) | $X_{63}$ | 2000 | 0,9 | 1,1 | 1 | 1 | 1 | 1 | 1 | 1 |
| | Kd_9_11 | $X_{62}$ | 0,0714 | 0,9 | 1,1 | 1 | 0,9 | 0,9 | 0,9 | 0,9 | 0,9 |
| | Kd_17_24 | $X_{64}$ | 0,0426 | 0,9 | 1,1 | 0,9 | 0,9 | 0,9 | 0,9 | 0,9 | 1,1 |
| | Kd_22_68 | $X_{65}$ | 0,000333333 | 0,9 | 1,1 | 1 | 1 | 1 | 1 | 1 | 0,9 |
| | Kd_71_23 | $X_{66}$ | 0,0005 | 0,9 | 1,1 | 0,9 | 0,9 | 0,9 | 0,9 | 0,9 | 1,1 |
| | Kd_71_21 | $X_{67}$ | 0,0005 | 0,9 | 1,1 | 1,1 | 1,1 | 1,1 | 1,1 | 1,1 | 1,1 |
| | hill_7_13 (adimensional) | $X_{70}$ | 1 | 0,9 | 1,1 | 1 | 1 | 1 | 1 | 1 | 1 |
| | $(v_{4,11})_{opt}/(v_{4,11})_{basal}$ | | | | | 1,36 | 16,41 | 17,68 | 17,68 | 17,68 | 17,68 |

***a***) Time dependent metabolites ($X_1 ... X_{10}$) ranges between 0.1 and 2.5 times its respective basal values

***b***) External variables ($X_{17}$, $X_{18}$, $X_{22}$, $X_{25}$, and $X_{71}$) allowed varying 10% aroundtheir respective basal values.

**c**) The kcat's ($X_{27}...X_{31}, X_{33}... X_{36}$) ranges between 0.2 and 10 times their basal values.

***d***) Cellular economy factor p = 1

***e***) The $v_{9,11}$basal flux was calculated as:

$$v_{9,11} = \frac{V_{max_{11}} \times X_4 \times X_9}{\left(K_{m_{4,11}} \times X_9 + K_{m_{9,11}} \times X_4 + X_4 \times X_9 + K_{d_{9,11}} \times K_{m_{9,11}}\right)} = 2.83 \times 10^{-4}\ mM \cdot min^{-1}$$

**Table S3.4b.** Specific constants (kcat/Km) fold change respect their basal values for data from Table S3.4a

| kcat / Km | Model symbols | Basal | Opt./Basal | | | | | |
|---|---|---|---|---|---|---|---|---|
| | | | 6 | 7 | 8 | 9 | 10 | 11 |
| kcat_11/Km_4_11 | $X_{26}/X_{56}$ | 168,12 | 1 | 11,11 | 11,11 | 11,11 | 11,11 | 11,11 |
| kcat_11/Km_9_11 | $X_{26}/X_{37}$ | 314,84 | 1 | 11,11 | 11,11 | 11,11 | 11,11 | 11,11 |
| kcat_12 / Km_2_12 | $X_{28}/X_{39}$ | 6733,33 | 1 | 1 | 1 | 1,33 | 1,33 | 1,06 |
| kcat_12 / Km_19_12 | $X_{28}/X_{50}$ | 2404,76 | 1 | 1 | 1 | 1,33 | 1,33 | 1,06 |
| kcat_13 / Km_7_13 | $X_{29}/X_{60}$ | 121212,12 | 1 | 1 | 1 | 1 | 1 | 0,41 |
| kcat_14 / Km_3_14 | $X_{30}/X_{40}$ | 238,05 | 1 | 1 | 1 | 1,55 | 1,57 | 1,26 |
| kcat_15 / Km_4_15 | $X_{31}/X_{41}$ | 2084,21 | 1 | 1 | 5,81 | 5,81 | 5,81 | 1,57 |
| kcat_15 / Km_5_15 | $X_{31}/X_{42}$ | 1384,62 | 1 | 1 | 5,81 | 5,81 | 5,81 | 1,57 |
| kcat_16 / Km_4_16 | $X_{32}/X_{46}$ | 600,00 | 0,48 | 0,52 | 0,76 | 0,76 | 0,76 | 0,52 |
| kcat_16 / Km_5_16 | $X_{32}/X_{47}$ | 26,62 | 0,48 | 0,52 | 0,76 | 0,76 | 0,76 | 0,52 |
| kcat_20 / Km_6_20 | $X_{33}/X_{43}$ | 494904,46 | 0,96 | 0,96 | 0,96 | 1 | 1,01 | 0,81 |
| kcat_20 / Km_25_20 | $X_{33}/X_{54}$ | 2044,74 | 0,96 | 0,96 | 0,96 | 1 | 1,01 | 0,81 |
| kcat_21 / Km_8_21 | $X_{34}/X_{45}$ | 920,00 | 0,18 | 0,18 | 0,18 | 0,18 | 0,18 | 0,18 |
| kcat_21 / Km_71_21 | $X_{34}/X_{51}$ | 2300,00 | 0,18 | 0,18 | 0,18 | 0,18 | 0,18 | 0,18 |
| kcat_23 / Km_7_23 | $X_{35}/X_{44}$ | 42,32 | 0,21 | 0,23 | 0,25 | 0,25 | 0,25 | 0,18 |
| kcat_23 / Km_71_23 | $X_{35}/X_{52}$ | 1975,00 | 0,22 | 0,23 | 0,25 | 0,25 | 0,25 | 0,18 |
| kcat_24 / Km_17_24 | $X_{36}/X_{48}$ | 748,51 | 0,25 | 0,21 | 0,21 | 0,21 | 0,21 | 0,18 |
| kcat_24/ Km_18_24 | $X_{36}/X_{49}$ | 2446,34 | 0,25 | 0,25 | 0,25 | 0,25 | 0,25 | 0,18 |
| kcat_68/ Km_1_68 | $X_{27}/X_{38}$ | 38666,67 | 1 | 1 | 1 | 1 | 1 | 0,80 |
| kcat_68/ Km_22_68 | $X_{27}/X_{53}$ | 3480000,00 | 1 | 1 | 1 | 1 | 1 | 0,80 |

**Table S3.5a**. Optimization of the *K*cat's allowing modifications in the Km's, Kd's, Ki's, and $K_{0,5}$'s. The $K_{cat}$ IspA basal ratio kcat_11/ kcat_16 = 2.22 ($X_{26}/X_{32}$) was imposed for all the experiments.

| | | | | | | Opt./Basal | | | |
|---|---|---|---|---|---|---|---|---|---|
| | | | Basal | LB | UB | 8 | 9 | 10 | 11 |
| Time-dependent | DXP (mM) | $X_1$ | 0,002519342 | 0,8 | 1.2 | 0,80 | 0,80 | 0,80 | 0,80 |
| | CDP_MEP | $X_2$ | 0,002135892 | 0,8 | 1.2 | 1,07 | 1,07 | 1,07 | 0,80 |
| | MEcDP | $X_3$ | 0,001552085 | 0,8 | 1.2 | 1,20 | 1,20 | 1,20 | 0,80 |
| | IPP | $X_4$ | 0,000396505 | 0,8 | 1.2 | 1,20 | 1,20 | 1,20 | 1,20 |
| | DMAPP | $X_5$ | 0,000496462 | 0,8 | 1.2 | 0,80 | 0,80 | 0,80 | 0,80 |
| | MEP | $X_6$ | 0,000125026 | 0,8 | 1.2 | 0,81 | 0,81 | 0,81 | 0,80 |
| | $MEC_{(c)}$ | $X_7$ | 0,000893951 | 0,8 | 1.2 | 0,80 | 0,80 | 0,80 | 1,04 |
| | HMBPP | $X_8$ | 3,84307E-05 | 0,8 | 1.2 | 1,06 | 1,06 | 1,06 | 1,06 |
| | GPP | $X_9$ | 0,000390628 | 0,8 | 1.2 | 1,20 | 1,20 | 1,20 | 1,20 |
| | $MEC_{(e)}$ | $X_{10}$ | 0,02709473 | 0,8 | 1.2 | 1,20 | 1,20 | 1,20 | 1,17 |
| | kcat_11 ($min^{-1}$) | $X_{26}$ | 1,7316 | 0,2 | 10 | 0,77 | 0,77 | 0,77 | 0,77 |
| | kcat_12 | $X_{28}$ | 1010 | 0,2 | 10 | 1 | 1,00 | 1,00 | 0,96 |
| | kcat_13 | $X_{29}$ | 600 | 0,2 | 10 | 0,62 | 0,62 | 0,62 | 0,40 |
| | kcat_14 | $X_{30}$ | 80,7 | 0,2 | 10 | 1,14 | 1,14 | 1,14 | 1,13 |
| | kcat_15 | $X_{31}$ | 19,8 | 0,2 | 10 | 5,94 | 5,94 | 5,94 | 4,93 |
| | kcat_16 | $X_{32}$ | 0,78 | 0,2 | 10 | 0,77 | 0,77 | 0,77 | 0,77 |
| | kcat_20 | $X_{33}$ | 1554 | 0,2 | 10 | 1 | 1 | 1 | 0,73 |
| | kcat_21 | $X_{34}$ | 27,6 | 0,2 | 10 | 0,20 | 0,20 | 0,20 | 0,2 |
| | kcat_23 | $X_{35}$ | 23,7 | 0,2 | 10 | 0,21 | 0,21 | 0,21 | 0,20 |
| | kcat_24 | $X_{36}$ | 100,3 | 0,2 | 10 | 0,23 | 0,23 | 0,23 | 0,20 |
| | kcat_68 | $X_{27}$ | 1740 | 0,2 | 10 | 1 | 1 | 1 | 0,72 |
| | kcat_11/ kcat_16 | $X_{26}/X_{32}$ | 2.22 | - | - | 2,22 | 2,22 | 2,22 | 2,22 |
| Time-independent | Piruvate (mM) | $X_{17}$ | 2,67 | 0,9 | 1,1 | 0,90 | 0,90 | 0,90 | 0,90 |
| | G3P | $X_{18}$ | 0,041 | 0,9 | 1,1 | 0,90 | 0,90 | 0,90 | 0,90 |
| | ATP | $X_{19}$ | 3 | 0,9 | 1,1 | 0,90 | 0,90 | 0,90 | 0,90 |
| | NADPH | $X_{22}$ | 0,12 | 0,9 | 1,1 | 0,90 | 0,90 | 0,90 | 0,90 |
| | CTP | $X_{25}$ | 2,7 | 0,9 | 1,1 | 0,90 | 0,90 | 0,90 | 0,90 |
| | FldA | $X_{71}$ | 0,003015 | 0,9 | 1,1 | 0,90 | 0,90 | 0,90 | 0,90 |
| | Km_1_68 (mM) | $X_{38}$ | 0,045 | 0,9 | 1,1 | 1 | 1 | 1 | 0,9 |
| | Km_2_12 | $X_{39}$ | 0,15 | 0,9 | 1,1 | 1 | 1 | 1 | 0,9 |
| | Km_3_14 | $X_{40}$ | 0,339 | 0,9 | 1,1 | 1,1 | 1,1 | 1,1 | 0,9 |
| | Km_4_15 | $X_{41}$ | 0,0095 | 0,9 | 1,1 | 1,1 | 1,1 | 1,1 | 0,9 |
| | Km_4_16 | $X_{46}$ | 0,0013 | 0,9 | 1,1 | 1,1 | 1,1 | 1,1 | 1,1 |
| | Km_4_11 | $X_{56}$ | 0,0103 | 0,9 | 1,1 | 0,9 | 0,9 | 0,9 | 0,9 |
| | Km_5_15 | $X_{42}$ | 0,0143 | 0,9 | 1,1 | 1,1 | 1,1 | 1,1 | 0,9 |
| | Km_5_16 | $X_{47}$ | 0,0293 | 0,9 | 1,1 | 1,1 | 1,1 | 1,1 | 1,1 |
| | Km_6_20 | $X_{43}$ | 0,00314 | 0,9 | 1,1 | 1 | 1 | 1 | 0,9 |
| | Km_7_13 | $X_{60}$ | 0,00495 | 0,9 | 1,1 | 0,9 | 0,9 | 0,9 | 0,9 |

| | | | | | | | | | |
|---|---|---|---|---|---|---|---|---|---|
| | Km_7_23 | $X_{44}$ | 0,56 | 0,9 | 1,1 | 0,9 | 0,9 | 0,9 | 1,1 |
| | Km_8_21 | $X_{45}$ | 0,03 | 0,9 | 1,1 | 1,1 | 1,1 | 1,1 | 1,1 |
| | Km_9_11 | $X_{37}$ | 0,0055 | 0,9 | 1,1 | 0,9 | 0,9 | 0,9 | 0,9 |
| | Km_17_24 | $X_{48}$ | 0,134 | 0,9 | 1,1 | 1,1 | 1,1 | 1,1 | 1,1 |
| | Km_18_24 | $X_{49}$ | 0,041 | 0,9 | 1,1 | 0,9 | 0,9 | 0,9 | 1,1 |
| | Km_19_12 | $X_{50}$ | 0,42 | 0,9 | 1,1 | 1 | 1 | 1 | 0,9 |
| | Km_71_21 | $X_{51}$ | 0,012 | 0,9 | 1,1 | 1,10 | 1,1 | 1,1 | 1,1 |
| | Km_71_23 | $X_{52}$ | 0,012 | 0,9 | 1,1 | 0,90 | 0,9 | 0,9 | 1,1 |
| | Km_22_68 | $X_{53}$ | 0,0005 | 0,9 | 1,1 | 1 | 1 | 1 | 0,9 |
| | Km_25_20 | $X_{54}$ | 0,76 | 0,9 | 1,1 | 1 | 1 | 1 | 0,9 |
| | K_10_55 (mM) | $X_{55}$ | 0,05 | 0,9 | 1,1 | 1,10 | 1,10 | 1,10 | 0,90 |
| | K_4_58 | $X_{58}$ | 0,005 | 0,9 | 1,1 | 1,10 | 1,10 | 1,10 | 1,10 |
| | K_9_72 | $X_{72}$ | 0,005 | 0,9 | 1,1 | 1,10 | 1,10 | 1,10 | 1,10 |
| | Ki_4_24 | $X_{57}$ | 0,0654 | 1 | 1 | 1 | 1 | 1 | 1 |
| | Ki_5_24 | $X_{59}$ | 0,0813 | 1 | 1 | 1 | 1 | 1 | 1 |
| Time-independent | Ka_6_14 | $X_{69}$ | 0,00339 | 0,9 | 1,1 | 1 | 1 | 1 | 1 |
| | $\alpha_6$ (adimensional) | $X_{61}$ | 1 | 0,9 | 1,1 | 1 | 1 | 1 | 1 |
| | $\beta_6$(adimensional) | $X_{63}$ | 2000 | 0,9 | 1,1 | 1 | 1 | 1 | 1 |
| | Kd_9_11 | $X_{62}$ | 0,0714 | 0,9 | 1,1 | 0,9 | 0,9 | 0,9 | 0,9 |
| | Kd_17_24 | $X_{64}$ | 0,0426 | 0,9 | 1,1 | 0,9 | 0,9 | 0,9 | 1,1 |
| | Kd_22_68 | $X_{65}$ | 0,000333333 | 0,9 | 1,1 | 1 | 1 | 1 | 0,9 |
| | Kd_71_23 | $X_{66}$ | 0,0005 | 0,9 | 1,1 | 0,90 | 0,9 | 0,9 | 1,1 |
| | Kd_71_21 | $X_{67}$ | 0,0005 | 0,9 | 1,1 | 1,1 | 1,1 | 1,1 | 1,1 |
| | hill_7_13 (adimensional) | $X_{70}$ | 1 | 0,9 | 1,1 | 1 | 1 | 1 | 1 |
| | $(v_{4,11})_{opt}/(v_{4,11})_{basal}$ | | | | | 1.36 | 1.36 | 1.36 | 1.36 |

***a)*** Time dependent metabolites ($X_1$... $X_{10}$) ranges between 0.8 and 1.2 times its respective basal values

***b)*** External variables ($X_{17}$, $X_{18}$, $X_{22}$, $X_{25}$, and $X_{71}$) allowed varying a 10% around their respective basal values.

***c)*** The kcat's ($X_{27}$...$X_{31}$, $X_{33}$... $X_{36}$) ranges between 0.2 and 10 times their basal values.

***d)***The $v_{9,11}$basal flux was calculated as:

$$v_{9,11} = \frac{V_{max_{11}} \times X_4 \times X_9}{\left(K_{m_{4,11}} \times X_9 + K_{m_{9,11}} \times X_4 + X_4 \times X_9 + K_{d_{9,11}} \times K_{m_{9,11}}\right)} = 2.83 \times 10^{-4}\ mM \cdot min^{-1}$$

**Table S3.5b.** Specific constants (kcat/Km) fold change respect their basal values for data from Table S3.5a

| | | | Opt./Basal | | | | |
|---|---|---|---|---|---|---|---|
| **kcat / Km** | **Model symbols** | **Basal** | **7** | **8** | **9** | **10** | **11** |
| kcat_11/Km_4_11 | $X_{26}/ X_{56}$ | 168,12 | 0,58 | 0,86 | 0,86 | 0,86 | 0,86 |
| kcat_11/Km_9_11 | $X_{26} / X_{37}$ | 314,84 | 0,58 | 0,86 | 0,86 | 0,86 | 0,86 |
| kcat_12 / Km_2_12 | $X_{28} / X_{39}$ | 6733,33 | 1 | 1 | 1 | 1 | 1,07 |
| kcat_12 / Km_19_12 | $X_{28} / X_{50}$ | 2404,76 | 1 | 1 | 1 | 1 | 1,07 |
| kcat_13 / Km_7_13 | $X_{29} / X_{60}$ | 121212,12 | 1 | 1 | 1 | 1 | 0,44 |
| kcat_14 / Km_3_14 | $X_{30} / X_{40}$ | 238,05 | 1 | 1,03 | 1,03 | 1,03 | 1,26 |
| kcat_15 / Km_4_15 | $X_{31} / X_{41}$ | 2084,21 | 1,00 | 5,40 | 5,40 | 5,40 | 5,47 |
| kcat_15 / Km_5_15 | $X_{31} / X_{42}$ | 1384,62 | 1,00 | 5,40 | 5,40 | 5,40 | 5,47 |
| kcat_16 / Km_4_16 | $X_{32} / X_{46}$ | 600,00 | 0,48 | 0,70 | 0,70 | 0,70 | 0,70 |
| kcat_16 / Km_5_16 | $X_{32}/ X_{47}$ | 26,62 | 0,48 | 0,70 | 0,70 | 0,70 | 0,70 |
| kcat_20 / Km_6_20 | $X_{33} / X_{43}$ | 494904,46 | 0,96 | 1 | 1 | 1 | 0,82 |
| kcat_20 / Km_25_20 | $X_{33} / X_{54}$ | 2044,74 | 0,96 | 1 | 1 | 1 | 0,82 |
| kcat_21 / Km_8_21 | $X_{34} / X_{45}$ | 920,00 | 0,18 | 0,18 | 0,18 | 0,18 | 0,18 |
| kcat_21 / Km_71_21 | $X_{34} / X_{51}$ | 2300,00 | 0,18 | 0,18 | 0,18 | 0,18 | 0,18 |
| kcat_23 / Km_7_23 | $X_{35} / X_{44}$ | 42,32 | 0,21 | 0,23 | 0,23 | 0,23 | 0,18 |
| kcat_23 / Km_71_23 | $X_{35} / X_{52}$ | 1975,00 | 0,22 | 0,23 | 0,23 | 0,23 | 0,18 |
| kcat_24 / Km_17_24 | $X_{36} / X_{48}$ | 748,51 | 0,25 | 0,21 | 0,21 | 0,21 | 0,18 |
| kcat_24/ Km_18_24 | $X_{36} / X_{49}$ | 2446,34 | 0,25 | 0,25 | 0,25 | 0,25 | 0,18 |
| kcat_68/ Km_1_68 | $X_{27} / X_{38}$ | 38666,67 | 1 | 1 | 1 | 1 | 0,80 |
| kcat_68/ Km_22_68 | $X_{27} / X_{53}$ | 3480000,00 | 1 | 1 | 1 | 1 | 0,80 |

**Table S3.6a**. Optimization of the enzimatic levels allowing modifications in the Km's, $K_{0,5}$'s, and Ki's. The IspA ratio is set to one i.e., $X_{11}/X_{16} = 1$ for all the experiments.

| | | | | | | Opt./Basal | | | | | |
|---|---|---|---|---|---|---|---|---|---|---|---|
| | | | Basal | LB | UB | 1 | 2 | 3 | 4 | 5 | 6 |
| Time-dependent | DXP (mM) | $X_1$ | 0,002519342 | 0,1 | **5,5** | 3,64 | 3,63 | 3,63 | 3,63 | 3,63 | 3,63 |
| | CDP_MEP | $X_2$ | 0,002135892 | 0,1 | **5,5** | 4,60 | 4,59 | 4,59 | 4,59 | 4,59 | 4,59 |
| | MEcDP | $X_3$ | 0,001552085 | 0,1 | 2,5 | 1,24 | 1,24 | 1,24 | 1,24 | 1,24 | 1,24 |
| | IPP | $X_4$ | 0,000396505 | 0,1 | 2,5 | 1,14 | 2,50 | 2,50 | 2,50 | 2,50 | 2,50 |
| | DMAPP | $X_5$ | 0,000496462 | 0,1 | 2,5 | 1,21 | 1,18 | 1,12 | 1,12 | 1,12 | 1,12 |
| | MEP | $X_6$ | 0,000125026 | 0,1 | **5,5** | 3,67 | 3,66 | 3,67 | 3,67 | 3,67 | 3,67 |
| | $MEC_{(c)}$ | $X_7$ | 0,000893951 | 0,1 | 2,5 | 0,28 | 0,37 | 0,37 | 0,37 | 0,37 | 0,37 |
| | HMBPP | $X_8$ | 3,84307E-05 | 0,1 | 2,5 | 0,28 | 0,38 | 0,38 | 0,38 | 0,38 | 0,38 |
| | GPP | $X_9$ | 0,000390628 | 0,1 | 2,5 | 2,50 | 2,50 | 2,50 | 2,50 | 2,50 | 2,50 |
| | $MEC_{(e)}$ | $X_{10}$ | 0,02709473 | 0,1 | **5,5** | 5,50 | 5,50 | 5,50 | 5,50 | 5,50 | 5,50 |
| Time-independent | **IspA** | $\mathbf{X_{11}}$ | 0,000163286 | 1 | 10 | 1 | 1 | 1 | 1 | 1,17 | 1,17 |
| | IspE, MCS | $X_{12}$ | 9,59423E-05 | 1 | 10 | 1 | 1 | 1 | 1 | 1 | 1 |
| | Effluxpump | $X_{13}$ | 1,47604E-05 | 1 | 10 | 6,35 | 4,99 | 4,96 | 4,94 | 4,94 | 4,94 |
| | IspF, HDS | $X_{14}$ | 5,12E-05 | 1 | 10 | 1 | 1 | 1 | 1 | 1 | 1 |
| | IDI | $X_{15}$ | 2,76757E-06 | 1 | 10 | 1 | 10 | 10 | 10 | 10 | 10 |
| | **IspA** | $\mathbf{X_{16}}$ | 0,000163286 | 1 | 10 | 1 | 1 | 1 | 1 | 1,17 | 1,17 |
| | IspD, CMK | $X_{20}$ | 2,31E-05 | 1 | 10 | 1 | 1 | 1 | 1 | 1 | 1 |
| | IspH | $X_{21}$ | 1,01E-04 | 1 | 10 | 1 | 1 | 1 | 1 | 1 | 1 |
| | IspG | $X_{23}$ | 9,40972E-05 | 1 | 10 | 1 | 1 | 1 | 1 | 1 | 1 |
| | DXS | $X_{24}$ | 2,81369E-05 | 1 | 10 | 1 | 1 | 1 | 1 | 1 | 1 |
| | DXR | $X_{68}$ | 1,47604E-05 | 1 | 10 | 1 | 1 | 1 | 1 | 1 | 1 |
| | IspA_11/IspA_16 | $X_{11} / X_{16}$ | 1 | - | - | 1 | 1 | 1 | 1 | 1 | 1 |
| | Piruvate (mM) | $X_{17}$ | 2,67 | 0,9 | 1,1 | 0,90 | 0,90 | 0,90 | 0,90 | 0,90 | 0,90 |
| | G3P | $X_{18}$ | 0,041 | 0,9 | 1,1 | 0,90 | 0,90 | 0,90 | 0,90 | 0,90 | 0,90 |
| | ATP | $X_{19}$ | 3 | 0,9 | 1,1 | 0,90 | 0,90 | 0,90 | 0,90 | 0,90 | 0,90 |
| | NADPH | $X_{22}$ | 0,12 | 0,9 | 1,1 | 0,90 | 0,90 | 0,90 | 0,90 | 0,90 | 0,90 |
| | CTP | $X_{25}$ | 2,7 | 0,9 | 1,1 | 0,90 | 0,90 | 0,90 | 0,90 | 0,90 | 0,90 |
| | FldA | $X_{71}$ | 0,003015 | 0,9 | 1,1 | 0,90 | 0,90 | 0,90 | 0,90 | 0,90 | 0,90 |
| | Km_1_68 (mM) | $X_{38}$ | 0,045 | 0,9 | 1,1 | 1 | 1 | 1 | 1 | 1 | 1 |
| | Km_2_12 | $X_{39}$ | 0,15 | 0,9 | 1,1 | 1 | 1 | 1 | 1 | 1 | 1 |
| | Km_3_14 | $X_{40}$ | 0,339 | 0,9 | 1,1 | 1 | 1 | 1 | 1 | 1 | 1 |
| | Km_4_15 | $X_{41}$ | 0,0095 | 0,9 | 1,1 | 1 | 1,02 | 1,10 | 1,10 | 1,10 | 1,10 |
| | Km_4_16 | $X_{46}$ | 0,0013 | 0,9 | 1,1 | 1 | 1 | 1,10 | 1,10 | 1,10 | 1,10 |
| | Km_4_11 | $X_{56}$ | 0,0103 | 0,9 | 1,1 | 1 | 1 | 0,90 | 0,90 | 0,90 | 0,90 |
| | Km_5_15 | $X_{42}$ | 0,0143 | 0,9 | 1,1 | 1 | 0,90 | 0,90 | 0,90 | 0,90 | 0,90 |
| | Km_5_16 | $X_{47}$ | 0,0293 | 0,9 | 1,1 | 1 | 1 | 0,94 | 1,10 | 1 | 1 |
| | Km_6_20 | $X_{43}$ | 0,00314 | 0,9 | 1,1 | 1 | 1 | 1 | 1 | 1 | 1 |
| | Km_7_13 | $X_{60}$ | 0,00495 | 0,9 | 1,1 | 0,9 | 0,90 | 0,90 | 0,90 | 0,90 | 0,90 |

Time-independent

| | | | | | | | | | | |
|---|---|---|---|---|---|---|---|---|---|---|
| Km_7_23 | $X_{44}$ | 0,56 | 0,9 | 1,1 | 1 | 1 | 1 | 1 | 1 | 1 |
| Km_8_21 | $X_{45}$ | 0,03 | 0,9 | 1,1 | 1 | 1 | 1 | 1 | 1 | 1 |
| Km_9_11 | $X_{37}$ | 0,0055 | 0,9 | 1,1 | 1 | 1 | 0,90 | 0,90 | 0,90 | 0,90 |
| Km_17_24 | $X_{48}$ | 0,134 | 0,9 | 1,1 | 1 | 1 | 1 | 1 | 1 | 1 |
| Km_18_24 | $X_{49}$ | 0,041 | 0,9 | 1,1 | 1 | 1 | 1 | 1 | 1 | 1 |
| Km_19_12 | $X_{50}$ | 0,42 | 0,9 | 1,1 | 1 | 1 | 1 | 1 | 1 | 1 |
| Km_71_21 | $X_{51}$ | 0,012 | 0,9 | 1,1 | 1 | 1 | 1 | 1 | 1 | 1 |
| Km_71_23 | $X_{52}$ | 0,012 | 0,9 | 1,1 | 1 | 1 | 1 | 1 | 1 | 1 |
| Km_22_68 | $X_{53}$ | 0,0005 | 0,9 | 1,1 | 1 | 1 | 1 | 1 | 1 | 1 |
| Km_25_20 | $X_{54}$ | 0,76 | 0,9 | 1,1 | 1 | 1 | 1 | 1 | 1 | 1 |
| K_10_55 (mM) | $X_{55}$ | 0,05 | 0,9 | 1,1 | 1,01 | 1,01 | 1,01 | 1,01 | 1,01 | 1,01 |
| K_4_58 | $X_{58}$ | 0,005 | 0,9 | 1,1 | 0,90 | 0,90 | 0,90 | 0,90 | 0,90 | 0,90 |
| K_9_72 | $X_{72}$ | 0,005 | 0,9 | 1,1 | 1,10 | 1,10 | 1,10 | 1,10 | 1,10 | 1,10 |
| Ki_4_24 | $X_{57}$ | 0,0654 | 1 | 1 | 1 | 1 | 1 | 1 | 1 | 1 |
| Ki_5_24 | $X_{59}$ | 0,0813 | 1 | 1 | 1 | 1 | 1 | 1 | 1 | 1 |
| Ka_6_14 | $X_{69}$ | 0,00339 | 0,9 | 1,1 | 1 | 1 | 1 | 1 | 1 | 1 |
| □$_6$ (adimensional) | $X_{61}$ | 1 | 0,9 | 1,1 | 1 | 1 | 1 | 1 | 1 | 1 |
| □$_6$(adimensional) | $X_{63}$ | 2000 | 0,9 | 1,1 | 1 | 1 | 1 | 1 | 1 | 1 |
| Kd_9_11 | $X_{62}$ | 0,0714 | 0,9 | 1,1 | 1 | 1 | 0,90 | 0,90 | 0,90 | 0,90 |
| Kd_17_24 | $X_{64}$ | 0,0426 | 0,9 | 1,1 | 1 | 1 | 1 | 1 | 1 | 1 |
| Kd_22_68 | $X_{65}$ | 0,000333333 | 0,9 | 1,1 | 1 | 1 | 1 | 1 | 1 | 1 |
| Kd_71_23 | $X_{66}$ | 0,0005 | 0,9 | 1,1 | 1 | 1 | 1 | 1 | 1 | 1 |
| Kd_71_21 | $X_{67}$ | 0,0005 | 0,9 | 1,1 | 1 | 1 | 1 | 1 | 1 | 1 |
| hill_7_13 (adimensional) | $X_{70}$ | 1 | 0,9 | 1,1 | 1 | 1 | 1 | 1 | 1 | 1 |
| $(v_{4,11})_{opt} / (v_{4,11})_{basal}$ | | | | | 2,80 | 6,10 | 7,49 | 8,74 | 8,74 | 8,74 |

***a*)** Time dependent metabolites ($X_1$... $X_{10}$) ranges between 0.1 and 2.5 times its respective basal values

***b*)** External variables ($X_{17}$, $X_{18}$, $X_{22}$, $X_{25}$, and $X_{71}$) allowed varying a 10% around their respective basal values.

***c*)** The Km's, Kd's, Ki's, and $K_{0,5}$'s variables are allowed be modified around ± 10 % their corresponding basal values.

***d*)** Cellular economy factor p = 2

***e*)** The $v_{9,11}$ basal flux was:

$$v_{9,11} = \frac{V_{max_{11}} \times X_4 \times X_9}{\left(K_{m_{4,11}} \times X_9 + K_{m_{9,11}} \times X_4 + X_4 \times X_9 + K_{d_{9,11}} \times K_{m_{9,11}}\right)} = 2.83 \times 10^{-4}\ mM \cdot min^{-1}$$

**Table S3.6a (cont…)**

| | | | Basal | LB | UB | Opt./Basal 7 | 8 | 9 | 10 | 11 |
|---|---|---|---|---|---|---|---|---|---|---|
| Time-dependent | DXP (mM) | $X_1$ | 0,002519342 | 0,1 | **5,5** | 3,63 | 3,63 | 3,63 | 3,63 | 3,59 |
| | CDP_MEP | $X_2$ | 0,002135892 | 0,1 | **5,5** | 4,59 | 4,59 | 4,59 | 4,59 | 5,50 |
| | MEcDP | $X_3$ | 0,001552085 | 0,1 | 2,5 | 1,24 | 1,24 | 1,24 | 1,24 | 2,50 |
| | IPP | $X_4$ | 0,000396505 | 0,1 | 2,5 | 2,50 | 2,50 | 2,50 | 2,50 | 2,50 |
| | DMAPP | $X_5$ | 0,000496462 | 0,1 | 2,5 | 1,12 | 1,12 | 1,12 | 1,12 | 1,12 |
| | MEP | $X_6$ | 0,000125026 | 0,1 | **5,5** | 3,67 | 3,67 | 3,67 | 3,67 | 1,73 |
| | $MEC_{(c)}$ | $X_7$ | 0,000893951 | 0,1 | 2,5 | 0,37 | 0,37 | 0,37 | 0,37 | 0,42 |
| | HMBPP | $X_8$ | 3,84307E-05 | 0,1 | 2,5 | 0,38 | 0,38 | 0,38 | 0,38 | 0,42 |
| | GPP | $X_9$ | 0,000390628 | 0,1 | 2,5 | 2,50 | 2,50 | 2,50 | 2,50 | 2,50 |
| | $MEC_{(e)}$ | $X_{10}$ | 0,02709473 | 0,1 | **5,5** | 5,50 | 5,50 | 5,50 | 5,50 | 5,50 |
| | **IspA** | $\mathbf{X_{11}}$ | 0,000163286 | 1 | 10 | 1,17 | 1,17 | 1,17 | 1,17 | 1,17 |
| | IspE, MCS | $X_{12}$ | 9,59423E-05 | 1 | 10 | 1 | 1 | 1 | 1 | 1 |
| | Effluxpump | $X_{13}$ | 1,47604E-05 | 1 | 10 | 4,94 | 4,94 | 4,94 | 4,94 | 4,93 |
| | IspF, HDS | $X_{14}$ | 5,12E-05 | 1 | 10 | 1 | 1 | 1 | 1 | 1 |
| | IDI | $X_{15}$ | 2,76757E-06 | 1 | 10 | 10,00 | 10,00 | 10,00 | 10,00 | 10,00 |
| | **IspA** | $\mathbf{X_{16}}$ | 0,000163286 | 1 | 10 | 1,17 | 1,17 | 1,17 | 1,17 | 1,17 |
| | IspD, CMK | $X_{20}$ | 2,31E-05 | 1 | 10 | 1 | 1 | 1 | 1 | 2,01 |
| | IspH | $X_{21}$ | 1,01E-04 | 1 | 10 | 1 | 1 | 1 | 1 | 1 |
| | IspG | $X_{23}$ | 9,40972E-05 | 1 | 10 | 1 | 1 | 1 | 1 | 1 |
| | DXS | $X_{24}$ | 2,81369E-05 | 1 | 10 | 1,00 | 1,00 | 1,00 | 1,00 | 1,00 |
| | DXR | $X_{68}$ | 1,47604E-05 | 1 | 10 | 1 | 1 | 1 | 1 | 1 |
| | IspA_11/IspA_16 | $X_{11}$ / $X_{16}$ | 1 | - | - | 1 | 1 | 1 | 1 | 1 |
| Time-independent | Piruvate (mM) | $X_{17}$ | 2,67 | 0,9 | 1,1 | 0,90 | 0,90 | 0,90 | 0,90 | 1,10 |
| | G3P | $X_{18}$ | 0,041 | 0,9 | 1,1 | 0,90 | 0,90 | 0,90 | 0,90 | 1,10 |
| | ATP | $X_{19}$ | 3 | 0,9 | 1,1 | 0,90 | 0,90 | 0,90 | 0,90 | 1,10 |
| | NADPH | $X_{22}$ | 0,12 | 0,9 | 1,1 | 0,90 | 0,90 | 0,90 | 0,90 | 0,90 |
| | CTP | $X_{25}$ | 2,7 | 0,9 | 1,1 | 0,90 | 0,90 | 0,90 | 0,90 | 1,10 |
| | FldA | $X_{71}$ | 0,003015 | 0,9 | 1,1 | 0,90 | 0,90 | 0,90 | 0,90 | 0,90 |
| | Km_1_68 (mM) | $X_{38}$ | 0,045 | 0,9 | 1,1 | 1,00 | 1,00 | 1,00 | 1,00 | 0,90 |
| | Km_2_12 | $X_{39}$ | 0,15 | 0,9 | 1,1 | 1,00 | 1,00 | 1,00 | 1,00 | 1,10 |
| | Km_3_14 | $X_{40}$ | 0,339 | 0,9 | 1,1 | 1,00 | 1,00 | 1,00 | 1,00 | 0,90 |
| | Km_4_15 | $X_{41}$ | 0,0095 | 0,9 | 1,1 | 1,10 | 1,10 | 1,10 | 1,10 | 1,10 |
| | Km_4_16 | $X_{46}$ | 0,0013 | 0,9 | 1,1 | 1,10 | 1,10 | 1,10 | 1,10 | 1,10 |
| | Km_4_11 | $X_{56}$ | 0,0103 | 0,9 | 1,1 | 0,90 | 0,90 | 0,90 | 0,90 | 0,90 |
| | Km_5_15 | $X_{42}$ | 0,0143 | 0,9 | 1,1 | 0,90 | 0,90 | 0,90 | 0,90 | 0,90 |
| | Km_5_16 | $X_{47}$ | 0,0293 | 0,9 | 1,1 | 1,10 | 1,10 | 1,10 | 1,10 | 1,10 |
| | Km_6_20 | $X_{43}$ | 0,00314 | 0,9 | 1,1 | 1,00 | 1,00 | 1,00 | 1,00 | 0,90 |
| | Km_7_13 | $X_{60}$ | 0,00495 | 0,9 | 1,1 | 0,90 | 0,90 | 0,90 | 0,90 | 0,90 |
| | Km_7_23 | $X_{44}$ | 0,56 | 0,9 | 1,1 | 1,00 | 1,00 | 1,00 | 1,00 | 1,10 |
| | Km_8_21 | $X_{45}$ | 0,03 | 0,9 | 1,1 | 1,00 | 1,00 | 1,00 | 1,00 | 1,10 |
| | Km_9_11 | $X_{37}$ | 0,0055 | 0,9 | 1,1 | 0,90 | 0,90 | 0,90 | 0,90 | 0,90 |
| | Km_17_24 | $X_{48}$ | 0,134 | 0,9 | 1,1 | 1,00 | 1,00 | 1,00 | 1,00 | 0,90 |
| | Km_18_24 | $X_{49}$ | 0,041 | 0,9 | 1,1 | 1,00 | 1,00 | 1,00 | 1,00 | 1,04 |

| | | | | | | | | | | |
|---|---|---|---|---|---|---|---|---|---|---|
| | Km_19_12 | $X_{50}$ | 0,42 | 0,9 | 1,1 | 1,00 | 1,00 | 1,00 | 1,00 | 0,90 |
| | Km_71_21 | $X_{51}$ | 0,012 | 0,9 | 1,1 | 1,00 | 1,00 | 1,00 | 1,00 | 1,10 |
| | Km_71_23 | $X_{52}$ | 0,012 | 0,9 | 1,1 | 1,00 | 1,00 | 1,00 | 1,00 | 1,10 |
| | Km_22_68 | $X_{53}$ | 0,0005 | 0,9 | 1,1 | 1,00 | 1,00 | 1,00 | 1,00 | 0,90 |
| | Km_25_20 | $X_{54}$ | 0,76 | 0,9 | 1,1 | 1,00 | 1,00 | 1,00 | 1,00 | 0,90 |
| Time-independent | K_10_55 (mM) | $X_{55}$ | 0,05 | 0,9 | 1,1 | 1,01 | 1,01 | 1,01 | 1,01 | 1,10 |
| | K_4_58 | $X_{58}$ | 0,005 | 0,9 | 1,1 | 0,90 | 0,90 | 0,90 | 0,90 | 0,90 |
| | K_9_72 | $X_{72}$ | 0,005 | 0,9 | 1,1 | 1,10 | 1,10 | 1,10 | 1,10 | 1,10 |
| | Ki_4_24 | $X_{57}$ | 0,0654 | 1 | 1 | 1,00 | 1,00 | 1,00 | 1,00 | 1,00 |
| | Ki_5_24 | $X_{59}$ | 0,0813 | 1 | 1 | 1,00 | 1,00 | 1,00 | 1,00 | 1,00 |
| | Ka_6_14 | $X_{69}$ | 0,00339 | 0,9 | 1,1 | 1,00 | 1,00 | 1,00 | 1,00 | 1,00 |
| | $\alpha_6$ (adimensional) | $X_{61}$ | 1 | 0,9 | 1,1 | 1,00 | 1,00 | 1,00 | 1,00 | 1,00 |
| | $\beta_6$(adimensional) | $X_{63}$ | 2000 | 0,9 | 1,1 | 1,00 | 1,00 | 1,00 | 1,00 | 1,00 |
| | Kd_9_11 | $X_{62}$ | 0,0714 | 0,9 | 1,1 | 0,90 | 0,90 | 0,90 | 0,90 | 0,90 |
| | Kd_17_24 | $X_{64}$ | 0,0426 | 0,9 | 1,1 | 1,00 | 1,00 | 1,00 | 1,00 | 0,90 |
| | Kd_22_68 | $X_{65}$ | 0,000333333 | 0,9 | 1,1 | 1,00 | 1,00 | 1,00 | 1,00 | 0,90 |
| | Kd_71_23 | $X_{66}$ | 0,0005 | 0,9 | 1,1 | 1,00 | 1,00 | 1,00 | 1,00 | 1,10 |
| | Kd_71_21 | $X_{67}$ | 0,0005 | 0,9 | 1,1 | 1,00 | 1,00 | 1,00 | 1,00 | 1,10 |
| | hill_7_13(adimensional) | $X_{70}$ | 1 | 0,9 | 1,1 | 1,00 | 1,00 | 1,00 | 1,00 | 1,00 |
| | $(v_{4,11})_{opt} / (v_{4,11})_{basal}$ | | | | | 8,74 | 8,74 | 8,74 | 8,74 | 8,74 |

**Table S3.6b.** Specific constants (kcat/Km) fold change respect their respective basal values for data from Table S3.6a

| | | | Opt./Basal | | | | | |
|---|---|---|---|---|---|---|---|---|
| **kcat / Km** | **Model symbols** | **Basal** | **1** | **2** | **3** | **4** | **5** | **6** |
| kcat_11/Km_4_11 | $X_{26}/ X_{56}$ | 168,12 | 1 | 1 | 1 | 1,11 | 1,11 | 1,11 |
| kcat_11/Km_9_11 | $X_{26} / X_{37}$ | 314,84 | 1 | 1 | 1 | 1,11 | 1,11 | 1,11 |
| kcat_12 / Km_2_12 | $X_{28} / X_{39}$ | 6733,33 | 1 | 1 | 1 | 1 | 1 | 1 |
| kcat_12 / Km_19_12 | $X_{28} / X_{50}$ | 2404,76 | 1 | 1 | 1 | 1 | 1 | 1 |
| kcat_13 / Km_7_13 | $X_{29} / X_{60}$ | 121212,12 | 1 | 1 | 1 | 1 | 1 | 1 |
| kcat_14 / Km_3_14 | $X_{30} / X_{40}$ | 238,05 | 1 | 1 | 0,91 | 0,91 | 0,91 | 0,91 |
| kcat_15 / Km_4_15 | $X_{31} / X_{41}$ | 2084,21 | 1 | 1 | 1,11 | 1,11 | 1,11 | 1,11 |
| kcat_15 / Km_5_15 | $X_{31} / X_{42}$ | 1384,62 | 1 | 1 | 1 | 0,91 | 0,91 | 0,91 |
| kcat_16 / Km_4_16 | $X_{32} / X_{46}$ | 600,00 | 1 | 1 | 1 | 0,91 | 0,91 | 0,91 |
| kcat_16 / Km_5_16 | $X_{32}/ X_{47}$ | 26,62 | 1 | 1 | 1 | 1 | 1 | 1 |
| kcat_20 / Km_6_20 | $X_{33} / X_{43}$ | 494904,46 | 1 | 1 | 1 | 1 | 1 | 1 |
| kcat_20 / Km_25_20 | $X_{33} / X_{54}$ | 2044,74 | 1 | 1 | 1 | 1 | 1 | 1 |
| kcat_21 / Km_8_21 | $X_{34} / X_{45}$ | 920,00 | 1 | 1 | 1 | 1 | 1 | 1 |
| kcat_21 / Km_71_21 | $X_{34} / X_{51}$ | 2300,00 | 1 | 1 | 1 | 1 | 1 | 1 |
| kcat_23 / Km_7_23 | $X_{35} / X_{44}$ | 42,32 | 1 | 1 | 1 | 1 | 1 | 1 |
| kcat_23 / Km_71_23 | $X_{35} / X_{52}$ | 1975,00 | 1 | 0,91 | 0,91 | 0,91 | 0,91 | 0,91 |
| kcat_24 / Km_17_24 | $X_{36} / X_{48}$ | 748,51 | 1 | 0,91 | 0,91 | 0,91 | 0,91 | 0,91 |
| kcat_24/ Km_18_24 | $X_{36} / X_{49}$ | 2446,34 | 1 | 1 | 1 | 1 | 1 | 1 |
| kcat_68/ Km_1_68 | $X_{27} / X_{38}$ | 38666,67 | 1 | 1 | 1 | 1 | 1 | 1 |
| kcat_68/ Km_22_68 | $X_{27} / X_{53}$ | 3480000,00 | 1 | 1 | 1 | 1,11 | 1,11 | 1,11 |

**Table S3.6b** (cont..)

| kcat / Km | Model symbols | Basal | Opt./Basal 7 | 8 | 9 | 10 | 11 |
|---|---|---|---|---|---|---|---|
| kcat_11/Km_4_11 | $X_{26}$/ $X_{56}$ | 168,12 | 1,11 | 1,11 | 1,11 | 1,11 | 1,11 |
| kcat_11/Km_9_11 | $X_{26}$ / $X_{37}$ | 314,84 | 1,11 | 1,11 | 1,11 | 1,11 | 1,11 |
| kcat_12 / Km_2_12 | $X_{28}$ / $X_{39}$ | 6733,33 | 1 | 1 | 1 | 1 | 1 |
| kcat_12 / Km_19_12 | $X_{28}$ / $X_{50}$ | 2404,76 | 1 | 1 | 1 | 1 | 1 |
| kcat_13 / Km_7_13 | $X_{29}$ / $X_{60}$ | 121212,12 | 1 | 1 | 1 | 1 | 1 |
| kcat_14 / Km_3_14 | $X_{30}$ / $X_{40}$ | 238,05 | 0,91 | 0,91 | 0,91 | 0,91 | 0,91 |
| kcat_15 / Km_4_15 | $X_{31}$ / $X_{41}$ | 2084,21 | 1,11 | 1,11 | 1,11 | 1,11 | 1,11 |
| kcat_15 / Km_5_15 | $X_{31}$ / $X_{42}$ | 1384,62 | 0,91 | 0,91 | 0,91 | 0,91 | 0,91 |
| kcat_16 / Km_4_16 | $X_{32}$ / $X_{46}$ | 600,00 | 0,91 | 0,91 | 0,91 | 0,91 | 0,91 |
| kcat_16 / Km_5_16 | $X_{32}$/ $X_{47}$ | 26,62 | 1 | 1 | 1 | 1 | 1 |
| kcat_20 / Km_6_20 | $X_{33}$ / $X_{43}$ | 494904,46 | 1 | 1 | 1 | 1 | 1 |
| kcat_20 / Km_25_20 | $X_{33}$ / $X_{54}$ | 2044,74 | 1 | 1 | 1 | 1 | 1 |
| kcat_21 / Km_8_21 | $X_{34}$ / $X_{45}$ | 920,00 | 1 | 1 | 1 | 1 | 1 |
| kcat_21 / Km_71_21 | $X_{34}$ / $X_{51}$ | 2300,00 | 1 | 1 | 1 | 1 | 1 |
| kcat_23 / Km_7_23 | $X_{35}$ / $X_{44}$ | 42,32 | 1 | 1 | 1 | 1 | 1 |
| kcat_23 / Km_71_23 | $X_{35}$ / $X_{52}$ | 1975,00 | 0,91 | 0,91 | 0,91 | 0,91 | 0,91 |
| kcat_24 / Km_17_24 | $X_{36}$ / $X_{48}$ | 748,51 | 0,91 | 0,91 | 0,91 | 0,91 | 0,91 |
| kcat_24/ Km_18_24 | $X_{36}$ / $X_{49}$ | 2446,34 | 1 | 1 | 1 | 1 | 1 |
| kcat_68/ Km_1_68 | $X_{27}$ / $X_{38}$ | 38666,67 | 1 | 1 | 1 | 1 | 1 |
| kcat_68/ Km_22_68 | $X_{27}$ / $X_{53}$ | 3480000,00 | 1,11 | 1,11 | 1,11 | 1,11 | 1,11 |

**Table S3.7a**. Optimization of the enzymatic levels not allowing modifications in the Km's, $K_{0,5}$'s, and Ki's. The basal ratio for IspA ($X_{11}/X_{16}$) was set to one all along the optimizations.

| | | | Basal | LB | UB | Opt./Basal 1 | 2 | 3 | 4 | 5 | 6 |
|---|---|---|---|---|---|---|---|---|---|---|---|
| Time-dependent | DXP (mM) | $X_1$ | 0,002519342 | 0,1 | **5,5** | 3,64 | 3,63 | 3,63 | 3,63 | 3,63 | 3,63 |
| | CDP_MEP | $X_2$ | 0,002135892 | 0,1 | **5,5** | 4,60 | 4,59 | 4,59 | 4,59 | 4,59 | 4,59 |
| | MEcDP | $X_3$ | 0,001552085 | 0,1 | 2,5 | 1,24 | 1,24 | 1,24 | 1,24 | 1,24 | 1,24 |
| | IPP | $X_4$ | 0,000396505 | 0,1 | 2,5 | 1,14 | 2,24 | 2,24 | 2,24 | 2,24 | 2,24 |
| | DMAPP | $X_5$ | 0,000496462 | 0,1 | 2,5 | 1,21 | 1,19 | 1,19 | 1,19 | 1,19 | 1,19 |
| | MEP | $X_6$ | 0,000125026 | 0,1 | **5,5** | 3,67 | 3,66 | 3,66 | 3,66 | 3,66 | 3,66 |
| | $MEC_{(c)}$ | $X_7$ | 0,000893951 | 0,1 | 2,5 | 0,27 | 0,36 | 0,36 | 0,36 | 0,36 | 0,36 |
| | HMBPP | $X_8$ | 3,84307E-05 | 0,1 | 2,5 | 0,28 | 0,36 | 0,36 | 0,36 | 0,36 | 0,36 |
| | GPP | $X_9$ | 0,000390628 | 0,1 | 2,5 | 2,50 | 2,50 | 2,50 | 2,50 | 2,50 | 2,50 |
| | $MEC_{(e)}$ | $X_{10}$ | 0,02709473 | 0,1 | **5,5** | 5,50 | 5,50 | 5,50 | 5,50 | 5,50 | 5,50 |
| Time-independent | **IspA** | $\boldsymbol{X_{11}}$ | 0,000163286 | 1 | 10 | 1 | 1 | 1 | 1 | 1 | 1 |
| | IspE, MCS | $X_{12}$ | 9,59423E-05 | 1 | 10 | 1 | 1 | 1 | 1 | 1 | 1 |
| | Effluxpump | $X_{13}$ | 1,47604E-05 | 1 | 10 | 7,12 | 5,64 | 5,64 | 5,64 | 5,64 | 5,64 |
| | IspF, HDS | $X_{14}$ | 5,12E-05 | 1 | 10 | 1 | 1 | 1 | 1 | 1 | 1 |
| | IDI | $X_{15}$ | 2,76757E-06 | 1 | 10 | 1 | 10 | 10 | 10 | 10 | 10 |
| | **IspA** | $\boldsymbol{X_{16}}$ | 0,000163286 | 1 | 10 | 1 | 1 | 1 | 1 | 1 | 1 |
| | IspD, CMK | $X_{20}$ | 2,31E-05 | 1 | 10 | 1 | 1 | 1 | 1 | 1 | 1 |
| | IspH | $X_{21}$ | 1,01E-04 | 1 | 10 | 1 | 1 | 1 | 1 | 1 | 1 |
| | IspG | $X_{23}$ | 9,40972E-05 | 1 | 10 | 1 | 1 | 1 | 1 | 1 | 1 |
| | DXS | $X_{24}$ | 2,81369E-05 | 1 | 10 | 1 | 1 | 1 | 1 | 1 | 1 |
| | DXR | $X_{68}$ | 1,47604E-05 | 1 | 10 | 1 | 1 | 1 | 1 | 1 | 1 |
| | IspA_11/IspA_16 | $X_{11} / X_{16}$ | 1 | - | - | 1 | 1 | 1 | 1 | 1 | 1 |
| | Piruvate (mM) | $X_{17}$ | 2,67 | 0,9 | 1,1 | 0,90 | 0,90 | 0,90 | 0,90 | 0,90 | 0,90 |
| | G3P | $X_{18}$ | 0,041 | 0,9 | 1,1 | 0,90 | 0,90 | 0,90 | 0,90 | 0,90 | 0,90 |
| | ATP | $X_{19}$ | 3 | 0,9 | 1,1 | 0,90 | 0,90 | 0,90 | 0,90 | 0,90 | 0,90 |
| | NADPH | $X_{22}$ | 0,12 | 0,9 | 1,1 | 0,90 | 0,90 | 0,90 | 0,90 | 0,90 | 0,90 |
| | CTP | $X_{25}$ | 2,7 | 0,9 | 1,1 | 1,10 | 1,10 | 1,10 | 1,10 | 1,10 | 1,10 |
| | FldA | $X_{71}$ | 0,003015 | 0,9 | 1,1 | 1,10 | 0,90 | 0,90 | 0,90 | 0,90 | 0,90 |
| | K_10_55 (mM) | $X_{55}$ | 0,05 | 0,9 | 1,1 | 1,01 | 1,01 | 1,01 | 1,01 | 1,01 | 1,01 |
| | K_4_58 | $X_{58}$ | 0,005 | 0,9 | 1,1 | 0,90 | 0,90 | 0,90 | 0,90 | 0,90 | 0,90 |
| | K_9_72 | $X_{72}$ | 0,005 | 0,9 | 1,1 | 1,10 | 1,10 | 1,10 | 1,10 | 1,10 | 1,10 |
| | $(v_{4,11})_{opt}/(v_{4,11})_{basal}$ | | | | | 2,80 | 5,46 | 5,46 | 5,46 | 5,46 | 5,46 |

***a***) Time dependent metabolites ($X_1$,$X_2$, $X_6$,$X_{10}$) ranges between 0.1 and 5.5 their respective basal values and ($X_3$,…,$X_5$, $X_7$,…,$X_9$) are restrained to values between 0.1 and 2.5 their respective basal values.

***b***) External variables ($X_{17}$, $X_{18}$, $X_{22}$, $X_{25}$, and $X_{71}$) are allowed bascule a 10% around their respective basal values

***c*)**The enzymes ($X_{11}$...$X_{16}$, $X_{20}$, $X_{21}$, $X_{23}$, $X_{24}$, $X_{68}$) ranges between 1 and 10 times their basal values

***d*)** The Km's, Kd's, Ki's, and $K_{0,5}$'s variables are not allowed be modified

***e*)** Cellular economy factor p = 2

***f*)** The $v_{9,11}$ basal flux was calculated as:

$$v_{9,11} = \frac{V_{max_{11}} \times X_4 \times X_9}{\left(K_{m_{4,11}} \times X_9 + K_{m_{9,11}} \times X_4 + X_4 \times X_9 + K_{d_{9,11}} \times K_{m_{9,11}}\right)} = 2.83 \times 10^{-4}\ mM \cdot min^{-1}$$

**Table S3.7a (cont…)**

| | | | | | | Opt./Basal | | | | |
|---|---|---|---|---|---|---|---|---|---|---|
| | | | Basal | LB | UB | 7 | 8 | 9 | 10 | 11 |
| Time-dependent | DXP (mM) | $X_1$ | 0,002519342 | 0,1 | **5,5** | 3,63 | 3,63 | 3,63 | 3,63 | 3,63 |
| | CDP_MEP | $X_2$ | 0,002135892 | 0,1 | **5,5** | 4,59 | 4,59 | 4,59 | 4,59 | 4,59 |
| | MEcDP | $X_3$ | 0,001552085 | 0,1 | 2,5 | 1,24 | 1,24 | 1,24 | 1,24 | 2,50 |
| | IPP | $X_4$ | 0,000396505 | 0,1 | 2,5 | 2,24 | 2,24 | 2,24 | 2,24 | 2,24 |
| | DMAPP | $X_5$ | 0,000496462 | 0,1 | 2,5 | 1,19 | 1,19 | 1,19 | 1,19 | 1,19 |
| | MEP | $X_6$ | 0,000125026 | 0,1 | **5,5** | 3,66 | 3,66 | 3,66 | 3,66 | 1,76 |
| | $MEC_{(c)}$ | $X_7$ | 0,000893951 | 0,1 | 2,5 | 0,36 | 0,36 | 0,36 | 0,36 | 0,36 |
| | HMBPP | $X_8$ | 3,84307E-05 | 0,1 | 2,5 | 0,36 | 0,36 | 0,36 | 0,36 | 0,36 |
| | GPP | $X_9$ | 0,000390628 | 0,1 | 2,5 | 2,50 | 2,50 | 2,50 | 2,50 | 2,50 |
| | $MEC_{(e)}$ | $X_{10}$ | 0,02709473 | 0,1 | **5,5** | 5,50 | 5,50 | 5,50 | 5,50 | 5,50 |
| Time-independent | **IspA** | $\boldsymbol{X_{11}}$ | 0,000163286 | 1 | 10 | 1 | 1 | 1 | 1 | 1 |
| | IspE, MCS | $X_{12}$ | 9,59423E-05 | 1 | 10 | 1 | 1 | 1 | 1 | 1 |
| | Effluxpump | $X_{13}$ | 1,47604E-05 | 1 | 10 | 5,64 | 5,64 | 5,64 | 5,64 | 5,64 |
| | IspF, HDS | $X_{14}$ | 5,12E-05 | 1 | 10 | 1 | 1 | 1 | 1 | 1 |
| | IDI | $X_{15}$ | 2,76757E-06 | 1 | 10 | 10 | 10 | 10 | 10 | 10 |
| | **IspA** | $\boldsymbol{X_{16}}$ | 0,000163286 | 1 | 10 | 1 | 1 | 1 | 1 | 1 |
| | IspD, CMK | $X_{20}$ | 2,31E-05 | 1 | 10 | 1 | 1 | 1 | 1 | 2 |
| | IspH | $X_{21}$ | 1,01E-04 | 1 | 10 | 1 | 1 | 1 | 1 | 1 |
| | IspG | $X_{23}$ | 9,40972E-05 | 1 | 10 | 1 | 1 | 1 | 1 | 1 |
| | DXS | $X_{24}$ | 2,81369E-05 | 1 | 10 | 1 | 1 | 1 | 1 | 1 |
| | DXR | $X_{68}$ | 1,47604E-05 | 1 | 10 | 1 | 1 | 1 | 1 | 1 |
| | IspA_11/IspA_16 | $X_{11} / X_{16}$ | 1 | - | - | 1 | 1 | 1 | 1 | 1 |
| | Piruvate (mM) | $X_{17}$ | 2,67 | 0,9 | 1,1 | 0,90 | 0,90 | 0,90 | 0,90 | 0,90 |
| | G3P | $X_{18}$ | 0,041 | 0,9 | 1,1 | 0,90 | 0,90 | 0,90 | 0,90 | 0,90 |
| | ATP | $X_{19}$ | 3 | 0,9 | 1,1 | 0,90 | 0,90 | 0,90 | 0,90 | 0,90 |
| | NADPH | $X_{22}$ | 0,12 | 0,9 | 1,1 | 0,90 | 0,90 | 0,90 | 0,90 | 0,90 |
| | CTP | $X_{25}$ | 2,7 | 0,9 | 1,1 | 1,10 | 1,10 | 1,10 | 1,10 | 1,10 |
| | FldA | $X_{71}$ | 0,003015 | 0,9 | 1,1 | 0,90 | 0,90 | 0,90 | 0,90 | 0,90 |
| | K_10_55 (mM) | $X_{55}$ | 0,05 | 0,9 | 1,1 | 1,01 | 1,01 | 1,01 | 1,01 | 1,01 |
| | K_4_58 | $X_{58}$ | 0,005 | 0,9 | 1,1 | 0,90 | 0,90 | 0,90 | 0,90 | 0,90 |
| | K_9_72 | $X_{72}$ | 0,005 | 0,9 | 1,1 | 1,10 | 1,10 | 1,10 | 1,10 | 1,10 |
| | $(v_{4,11})_{opt} / (v_{4,11})_{basal}$ | | | | | 5,46 | 5,46 | 5,46 | 5,46 | 5,46 |

**Table S3.8a**. Optimization of the kcats and enzymatic levels allowing modifications in the Km's, $K_{0,5}$'s, and Ki's. The basal ratio for IspA ($X_{11}/X_{16}$) is one all along the optimizations.

| | | | | | | Opt./Basal | | | | | | |
|---|---|---|---|---|---|---|---|---|---|---|---|---|
| | | | **Basal** | **LB** | **UB** | **5** | **6** | **7** | **8** | **9** | **10** | **11** |
| Time-dependent | DXP (mM) | $X_1$ | 0,002519342 | 0,8 | **1,2** | 0,80 | 0,80 | 0,80 | 0,80 | 0,80 | 0,80 | 0,80 |
| | CDP_MEP | $X_2$ | 0,002135892 | 0,8 | **1,2** | 1,07 | 1,07 | 1,07 | 1,07 | 1,20 | 1,20 | 0,80 |
| | MEcDP | $X_3$ | 0,001552085 | 0,8 | 1,2 | 0,80 | 1,20 | 1,20 | 1,20 | 1,20 | 1,20 | 0,80 |
| | IPP | $X_4$ | 0,000396505 | 0,8 | 1,2 | 1,20 | 1,20 | 1,20 | 1,20 | 1,20 | 1,20 | 1,20 |
| | DMAPP | $X_5$ | 0,000496462 | 0,8 | 1,2 | 0,80 | 0,80 | 0,80 | 0,80 | 0,80 | 0,80 | 0,80 |
| | MEP | $X_6$ | 0,000125026 | 0,8 | **1,2** | 0,81 | 0,81 | 0,84 | 0,81 | 0,84 | 0,84 | 0,80 |
| | $MEC_{(c)}$ | $X_7$ | 0,000893951 | 0,8 | 1,2 | 0,80 | 0,80 | 0,80 | 0,80 | 0,80 | 0,80 | 0,80 |
| | HMBPP | $X_8$ | 3,84307E-05 | 0,8 | 1,2 | 0,99 | 0,91 | 0,91 | 0,91 | 0,91 | 0,91 | 1,11 |
| | GPP | $X_9$ | 0,000390628 | 0,8 | 1,2 | 1,20 | 1,20 | 1,20 | 1,20 | 1,20 | 1,20 | 1,20 |
| | $MEC_{(e)}$ | $X_{10}$ | 0,02709473 | 0,8 | **1,2** | 1,20 | 1,20 | 1,20 | 1,20 | 1,20 | 1,20 | 1,17 |
| | **IspA** | $X_{11}$ | 0,000163286 | 1 | 10 | 4,03 | 4,07 | 4,07 | 4,07 | 4,07 | 4,07 | 4,07 |
| | IspE, MCS | $X_{12}$ | 9,59423E-05 | 1 | 10 | 1 | 1 | 1 | 1 | 1 | 1 | 1 |
| | Effluxpump | $X_{13}$ | 1,47604E-05 | 1 | 10 | 1 | 1 | 1 | 1 | 1 | 1 | 1 |
| | IspF, HDS | $X_{14}$ | 5,12E-05 | 1 | 10 | 2 | 1 | 1 | 1 | 1 | 1 | 1 |
| | IDI | $X_{15}$ | 2,76757E-06 | 1 | 10 | 5,67 | 4,72 | 4,72 | 4,72 | 4,72 | 4,72 | 1 |
| | **IspA** | $X_{16}$ | 0,000163286 | 1 | 10 | 4,03 | 4,07 | 4,07 | 4,07 | 4,07 | 4,07 | 4,07 |
| | IspD, CMK | $X_{20}$ | 2,31E-05 | 1 | 10 | 1 | 1 | 1 | 1 | 1 | 1 | 1 |
| | IspH | $X_{21}$ | 1,01E-04 | 1 | 10 | 1 | 1 | 1 | 1 | 1 | 1 | 1 |
| | IspG | $X_{23}$ | 9,40972E-05 | 1 | 10 | 1 | 1 | 1 | 1 | 1 | 1 | 1 |
| | DXS | $X_{24}$ | 2,81369E-05 | 1 | 10 | 1 | 1,25 | 1,25 | 1 | 1 | 1 | 1 |
| | DXR | $X_{68}$ | 1,47604E-05 | 1 | 10 | 1 | 1 | 1 | 1 | 1 | 1 | 1 |
| | IspA_11/IspA_16 | $X_{11}/X_{16}$ | 1 | - | - | 1 | 1 | 1 | 1 | 1 | 1 | 1 |
| Time-independent | kcat_11 ($min^{-1}$) | $X_{26}$ | 1,7316 | **0,2** | **1,2** | 1 | 1,20 | 1,20 | 1,20 | 1,20 | 1,20 | 1,20 |
| | kcat_12 | $X_{28}$ | 1010 | 0,2 | 10 | 1 | 1 | 1 | 1 | 0,89 | 0,89 | 0,96 |
| | kcat_13 | $X_{29}$ | 600 | 0,2 | 10 | 0,68 | 0,68 | 0,68 | 0,68 | 0,68 | 0,68 | 0,49 |
| | kcat_14 | $X_{30}$ | 80,7 | 0,2 | 10 | 1 | 1 | 1 | 1 | 1 | 1 | 1,13 |
| | kcat_15 | $X_{31}$ | 19,8 | 0,2 | 10 | 1 | 1 | 1 | 1 | 1 | 1 | 5,20 |
| | kcat_16 | $X_{32}$ | 0,78 | **0,2** | **1,2** | 0,20 | 0,20 | 0,20 | 0,20 | 0,20 | 0,20 | 0,20 |
| | kcat_20 | $X_{33}$ | 1554 | 0,2 | 10 | 1 | 1 | 0,88 | 1 | 0,97 | 0,97 | 0,73 |
| | kcat_21 | $X_{34}$ | 27,6 | 0,2 | 10 | 0,20 | 0,20 | 0,20 | 0,20 | 0,20 | 0,20 | 0,20 |
| | kcat_23 | $X_{35}$ | 23,7 | 0,2 | 10 | 0,24 | 0,20 | 0,20 | 0,22 | 0,20 | 0,20 | 0,22 |
| | kcat_24 | $X_{36}$ | 100,3 | 0,2 | 10 | 0,24 | 0,20 | 0,20 | 0,24 | 0,24 | 0,24 | 0,20 |
| | kcat_68 | $X_{27}$ | 1740 | 0,2 | 10 | 1 | 1 | 1 | 1 | 1 | 1 | 0,72 |
| | kcat_11/ kcat_16 | $X_{26}/X_{32}$ | 2.22 | - | - | 11,10 | 13,32 | 13,32 | 13,32 | 13,32 | 13,32 | 13,32 |
| | Piruvate (mM) | $X_{17}$ | 2,67 | 0,9 | 1,1 | 0,90 | 0,90 | 0,90 | 0,90 | 0,90 | 0,90 | 0,90 |
| | G3P | $X_{18}$ | 0,041 | 0,9 | 1,1 | 0,90 | 0,90 | 0,90 | 0,90 | 0,90 | 0,90 | 0,90 |
| | ATP | $X_{19}$ | 3 | 0,9 | 1,1 | 0,90 | 0,90 | 0,90 | 0,90 | 1,10 | 1,10 | 0,90 |
| | NADPH | $X_{22}$ | 0,12 | 0,9 | 1,1 | 0,90 | 0,90 | 0,90 | 0,90 | 0,90 | 0,90 | 0,90 |

| | | | | | | | | | | | | |
|---|---|---|---|---|---|---|---|---|---|---|---|---|
| | CTP | $X_{25}$ | 2,7 | 0,9 | 1,1 | 0,90 | 0,90 | 0,90 | 0,90 | 0,90 | 0,90 | 0,90 |
| | FldA | $X_{71}$ | 0,003015 | 0,9 | 1,1 | 0,90 | 1,10 | 1,10 | 1,10 | 1,10 | 1,10 | 0,90 |
| Time-independent | Km_1_68 (mM) | $X_{38}$ | 0,045 | 0,9 | 1,1 | 1 | 1 | 1 | 1 | 1 | 1 | 0,90 |
| | Km_2_12 | $X_{39}$ | 0,15 | 0,9 | 1,1 | 1 | 1 | 1 | 1 | 1 | 1 | 0,90 |
| | Km_3_14 | $X_{40}$ | 0,339 | 0,9 | 1,1 | 1 | 1 | 1 | 1 | 1 | 1 | 0,90 |
| | Km_4_15 | $X_{41}$ | 0,0095 | 0,9 | 1,1 | 1 | 1 | 1 | 1 | 1 | 1 | 0,90 |
| | Km_4_16 | $X_{46}$ | 0,0013 | 0,9 | 1,1 | 1,10 | 1,10 | 1,10 | 1,10 | 1,10 | 1,10 | 1,10 |
| | Km_4_11 | $X_{56}$ | 0,0103 | 0,9 | 1,1 | 1 | 0,90 | 0,90 | 0,90 | 0,90 | 0,90 | 0,90 |
| | Km_5_15 | $X_{42}$ | 0,0143 | 0,9 | 1,1 | 1 | 1 | 1 | 1 | 1 | 1 | 0,90 |
| | Km_5_16 | $X_{47}$ | 0,0293 | 0,9 | 1,1 | 1,10 | 1,10 | 1,10 | 1,10 | 1,10 | 1,10 | 1,10 |
| | Km_6_20 | $X_{43}$ | 0,00314 | 0,9 | 1,1 | 1 | 1 | 0,90 | 1 | 1 | 1 | 0,90 |
| | Km_7_13 | $X_{60}$ | 0,00495 | 0,9 | 1,1 | 1 | 1 | 1 | 1 | 1 | 1 | 0,90 |
| | Km_7_23 | $X_{44}$ | 0,56 | 0,9 | 1,1 | 1 | 0,91 | 0,91 | 1 | 0,90 | 0,90 | 0,90 |
| | Km_8_21 | $X_{45}$ | 0,03 | 0,9 | 1,1 | 1 | 1 | 1 | 1 | 1 | 1 | 1,10 |
| | Km_9_11 | $X_{37}$ | 0,0055 | 0,9 | 1,1 | 1 | 0,90 | 0,90 | 0,90 | 0,90 | 0,90 | 0,90 |
| | Km_17_24 | $X_{48}$ | 0,134 | 0,9 | 1,1 | 1 | 1,10 | 1,10 | 1 | 1 | 1 | 1,10 |
| | Km_18_24 | $X_{49}$ | 0,041 | 0,9 | 1,1 | 1 | 1,10 | 1,10 | 1 | 1 | 1 | 1,10 |
| | Km_19_12 | $X_{50}$ | 0,42 | 0,9 | 1,1 | 1 | 1 | 1 | 1 | 1 | 1 | 0,90 |
| | Km_71_21 | $X_{51}$ | 0,012 | 0,9 | 1,1 | 1 | 1 | 1 | 1 | 1 | 1 | 1,10 |
| | Km_71_23 | $X_{52}$ | 0,012 | 0,9 | 1,1 | 1 | 1,10 | 1,10 | 1 | 1,10 | 1,10 | 0,90 |
| | Km_22_68 | $X_{53}$ | 0,0005 | 0,9 | 1,1 | 1 | 1 | 1 | 1 | 1 | 1 | 0,90 |
| | Km_25_20 | $X_{54}$ | 0,76 | 0,9 | 1,1 | 1 | 1 | 1 | 1 | 1 | 1 | 0,90 |
| | K_10_55 (mM) | $X_{55}$ | 0,05 | 0,9 | 1,1 | 1,10 | 1,10 | 1,10 | 1,10 | 1,10 | 1,10 | 0,90 |
| | K_4_58 | $X_{58}$ | 0,005 | 0,9 | 1,1 | 1,10 | 0,90 | 0,90 | 0,90 | 0,90 | 0,90 | 1,10 |
| | K_9_72 | $X_{72}$ | 0,005 | 0,9 | 1,1 | 1,10 | 1,10 | 1,10 | 1,10 | 1,10 | 1,10 | 1,10 |
| | Ki_4_24 | $X_{57}$ | 0,0654 | 1 | 1 | 1 | 1 | 1 | 1 | 1 | 1 | 1 |
| | Ki_5_24 | $X_{59}$ | 0,0813 | 1 | 1 | 1 | 1 | 1 | 1 | 1 | 1 | 1 |
| | Ka_6_14 | $X_{69}$ | 0,00339 | 0,9 | 1,1 | 1 | 1 | 1 | 1 | 1 | 1 | 1 |
| | $\alpha_6$ (adimensional) | $X_{61}$ | 1 | 0,9 | 1,1 | 1 | 1 | 1 | 1 | 1 | 1 | 1 |
| | $\beta_6$(adimensional) | $X_{63}$ | 2000 | 0,9 | 1,1 | 1 | 1 | 1 | 1 | 1 | 1 | 1 |
| | Kd_9_11 | $X_{62}$ | 0,0714 | 0,9 | 1,1 | 1 | 0,90 | 0,90 | 0,90 | 0,90 | 0,90 | 0,90 |
| | Kd_17_24 | $X_{64}$ | 0,0426 | 0,9 | 1,1 | 1 | 1,10 | 1,10 | 1 | 1 | 1 | 1,10 |
| | Kd_22_68 | $X_{65}$ | 0,000333333 | 0,9 | 1,1 | 1 | 1 | 1 | 1 | 1 | 1 | 0,90 |
| | Kd_71_23 | $X_{66}$ | 0,0005 | 0,9 | 1,1 | 1 | 0,90 | 0,90 | 1 | 1,10 | 1,10 | 0,90 |
| | Kd_71_21 | $X_{67}$ | 0,0005 | 0,9 | 1,1 | 1 | 1 | 1 | 1 | 1 | 1 | 1,10 |
| | hill_7_13 (adimensional) | $X_{70}$ | 1 | 0,9 | 1,1 | 1 | 1 | 1 | 1 | 1 | 1 | 1 |
| | $(v_{4,11})_{opt}/(v_{4,11})_{basal}$ | | | | | 5,78 | 8,65 | 8,65 | 8,65 | 8,65 | 8,65 | 8,65 |

***a*)**Time dependent metabolites ($X_1$,$X_2$, $X_6$,$X_{10}$) ranges between 0.8 and 1.2 their respective basal values and ($X_3$,…,$X_5$, $X_7$,…,$X_9$) are restrained to values between 0.1 and 2.5 their respective basal values.

***b*)** External variables ($X_{17}$, $X_{18}$, $X_{22}$, $X_{25}$, and $X_{71}$) are allowed bascule 10% around their respective basal values

***c*)** The enzymes ($X_{11}$...$X_{16}$, $X_{20}$, $X_{21}$, $X_{23}$, $X_{24}$, $X_{68}$) ranges between 1 and 10 times their basal values

***d*)** The kcat's ($X_{27}$...$X_{31}$, $X_{33}$... $X_{36}$) ranges between 0.2 and 10 times their basal values.

***e*)** The Km's, Kd's, Ki's, and $K_{0,5}$'s variables are not allowed be modified

***f*)** Cellular economy factor p = 2

***g*)** The $v_{9,11}$ basal flux was calculated as:

$$v_{9,11} = \frac{V_{max_{11}} \times X_4 \times X_9}{\left(K_{m_{4,11}} \times X_9 + K_{m_{9,11}} \times X_4 + X_4 \times X_9 + K_{d_{9,11}} \times K_{m_{9,11}}\right)} = 2.83 \times 10^{-4}\ mM \cdot min^{-1}$$

**Table S3.8b.** Specific constants (kcat/Km) fold change respect their respective basal values for data from Table S3.8a

| | | | Opt./Basal | | | | | | |
|---|---|---|---|---|---|---|---|---|---|
| **kcat / Km** | **Model symbols** | **Basal** | **5** | **6** | **7** | **8** | **9** | **10** | **11** |
| k_11/Km_4_11 | X26/ X56 | 168,116505 | 1 | 1,33 | 1,33 | 1,33 | 1,33 | 1,33 | 1,33 |
| k_11/Km_9_11 | X26 / X37 | 314,836364 | 1 | 1,33 | 1,33 | 1,33 | 1,33 | 1,33 | 1,33 |
| k_12 / Km_2_12 | X28 / X39 | 6733,33333 | 1 | 1 | 1 | 1 | 0,89 | 0,89 | 1,07 |
| k_12 / Km_19_12 | X28 / X50 | 2404,7619 | 1 | 1 | 1 | 1 | 0,89 | 0,89 | 1,07 |
| k_14 / Km_3_14 | X30 / X40 | 238,053097 | 1 | 1 | 1 | 1 | 1 | 1 | 1,26 |
| k_15 / Km_4_15 | X31 / X41 | 2084,21053 | 1 | 1 | 1 | 1 | 1 | 1 | 5,77 |
| k_15 / Km_5_15 | X31 / X42 | 1384,61538 | 1 | 1 | 1 | 1 | 1 | 1 | 5,77 |
| k_16 / Km_4_16 | X32 / X46 | 600 | 0,18 | 0,18 | 0,18 | 0,18 | 0,18 | 0,18 | 0,18 |
| k_16 / Km_5_16 | X32/ X47 | 26,6211604 | 0,18 | 0,18 | 0,18 | 0,18 | 0,18 | 0,18 | 0,18 |
| k_20 / Km_6_20 | X33 / X43 | 494904,459 | 1 | 1 | 0,97 | 1 | 0,97 | 0,97 | 0,82 |
| k_20 / Km_25_20 | X33 / X54 | 2044,73684 | 1 | 1 | 0,80 | 1 | 0,97 | 0,97 | 0,82 |
| k_21 / Km_8_21 | X34 / X45 | 920 | 0,20 | 0,20 | 0,20 | 0,20 | 0,20 | 0,20 | 0,18 |
| k_21 / Km_71_21 | X34 / X51 | 2300 | 0,20 | 0,20 | 0,20 | 0,20 | 0,20 | 0,20 | 0,18 |
| k_23 / Km_7_23 | X35 / X44 | 42,3214286 | 0,24 | 0,22 | 0,22 | 0,22 | 0,23 | 0,23 | 0,24 |
| k_23 / Km_71_23 | X35 / X52 | 1975 | 0,24 | 0,18 | 0,18 | 0,22 | 0,19 | 0,19 | 0,24 |
| k_24 / Km_17_24 | X36 / X48 | 748,507463 | 0,24 | 0,18 | 0,18 | 0,24 | 0,24 | 0,24 | 0,18 |
| k_24/ Km_18_24 | X36 / X49 | 2446,34146 | 0,24 | 0,18 | 0,18 | 0,24 | 0,24 | 0,24 | 0,18 |
| k_68/ Km_1_68 | X27 / X38 | 38666,6667 | 1 | 1 | 1 | 1 | 1 | 1 | 0,80 |
| k_68/ Km_22_68 | X27 / X53 | 3480000 | 1 | 1 | 1 | 1 | 1 | 1 | 0,80 |

**Table S3.9a**. Optimization of the kcat and enzymatic levels allowing modifications in the Km's, $K_{0,5}$'s, and Ki's. The basal ratio for IspA ($X_{11}/X_{16}$) can change all along the optimizations.

| | | | | | | Opt./Basal | | | | | | |
|---|---|---|---|---|---|---|---|---|---|---|---|---|
| | | | **Basal** | **LB** | **UB** | **5** | **6** | **7** | **8** | **9** | **10** | **11** |
| Time-dependent | DXP (mM) | $X_1$ | 0,002519342 | 0,8 | **1,2** | 0,80 | 0,80 | 0,80 | 0,80 | 0,80 | 1 | 0,80 |
| | CDP_MEP | $X_2$ | 0,002135892 | 0,8 | **1,2** | 1,07 | 1,07 | 1,07 | 1,07 | 1,07 | 1 | 0,80 |
| | MEcDP | $X_3$ | 0,001552085 | 0,8 | 1,2 | 1,20 | 1,20 | 1,20 | 1,20 | 1,20 | 1 | 0,80 |
| | IPP | $X_4$ | 0,000396505 | 0,8 | 1,2 | 1,20 | 1,20 | 1,20 | 1,20 | 1,20 | 1 | 1,20 |
| | DMAPP | $X_5$ | 0,000496462 | 0,8 | 1,2 | 1,20 | 1,20 | 1,20 | 1,20 | 1,20 | 1 | 0,80 |
| | MEP | $X_6$ | 0,000125026 | 0,8 | **1,2** | 0,81 | 0,81 | 0,81 | 0,81 | 0,81 | 1 | 0,80 |
| | $MEC_{(c)}$ | $X_7$ | 0,000893951 | 0,8 | 1,2 | 0,80 | 0,80 | 0,80 | 0,80 | 0,80 | 1 | 0,80 |
| | HMBPP | $X_8$ | 3,84307E-05 | 0,8 | 1,2 | 0,81 | 0,82 | 0,82 | 0,82 | 0,82 | 1 | 1,14 |
| | GPP | $X_9$ | 0,000390628 | 0,8 | 1,2 | 1,20 | 1,20 | 1,20 | 1,20 | 1,20 | 1 | 1,20 |
| | $MEC_{(e)}$ | $X_{10}$ | 0,02709473 | 0,8 | **1,2** | 1,20 | 1,20 | 1,20 | 1,20 | 1,20 | 1 | 1,17 |
| Time-independent | **IspA** | $\mathbf{X_{11}}$ | 0,000163286 | 1 | 10 | 10,00 | 10,00 | 10,00 | 10,00 | 10,00 | 1 | 10,00 |
| | IspE, MCS | $X_{12}$ | 9,59423E-05 | 1 | 10 | 1 | 1 | 1 | 1 | 1 | 1 | 1 |
| | Effluxpump | $X_{13}$ | 1,47604E-05 | 1 | 10 | 1 | 1 | 1 | 1 | 1 | 1 | 1 |
| | IspF, HDS | $X_{14}$ | 5,12E-05 | 1 | 10 | 1,03 | 1,03 | 1,04 | 1,03 | 1,03 | 1 | 1 |
| | IDI | $X_{15}$ | 2,76757E-06 | 1 | 10 | 1,28 | 1,29 | 1 | 1,29 | 1 | 0,77 | 1 |
| | **IspA** | $\mathbf{X_{16}}$ | 0,000163286 | 1 | 10 | 1 | 2,36 | 2,36 | 2,36 | 2,36 | 1 | 1 |
| | IspD, CMK | $X_{20}$ | 2,31E-05 | 1 | 10 | 1 | 1 | 1 | 1 | 1 | 1 | 1 |
| | IspH | $X_{21}$ | 1,01E-04 | 1 | 10 | 1 | 1 | 1 | 1 | 1 | 1 | 1 |
| | IspG | $X_{23}$ | 9,40972E-05 | 1 | 10 | 1 | 1 | 1 | 1 | 1 | 1 | 1 |
| | DXS | $X_{24}$ | 2,81369E-05 | 1 | 10 | 1 | 1,25 | 1 | 1 | 1 | 0,80 | 1 |
| | DXR | $X_{68}$ | 1,47604E-05 | 1 | 10 | 1 | 1 | 1 | 1 | 1 | 1 | 1 |
| | IspA_11/IspA_16 | $X_{11}$ / $X_{16}$ | 2.22 | - | - | 10 | 4,23 | 4,23 | 4,23 | 4,23 | 1 | 10 |
| | kcat_11 ($min^{-1}$) | $X_{26}$ | 1,7316 | **0,2** | **1,2** | 1 | 1,20 | 1,20 | 1,20 | 1,20 | 1 | 1,20 |
| | kcat_12 | $X_{28}$ | 1010 | 0,2 | 10 | 1 | 1 | 1 | 1 | 1 | 1 | 0,96 |
| | kcat_13 | $X_{29}$ | 600 | 0,2 | 10 | 0,68 | 0,68 | 0,68 | 0,68 | 0,68 | 1 | 0,49 |
| | kcat_14 | $X_{30}$ | 80,7 | 0,2 | 10 | 1 | 1 | 1 | 1 | 1 | 1 | 1,13 |
| | kcat_15 | $X_{31}$ | 19,8 | 0,2 | 10 | 1 | 1 | 1,29 | 1 | 2,25 | 2,25 | 5,33 |
| | kcat_16 | $X_{32}$ | 0,78 | **0,2** | **1,2** | 0,42 | 0,20 | 0,20 | 0,20 | 0,20 | 1 | 0,69 |
| | kcat_20 | $X_{33}$ | 1554 | 0,2 | 10 | 1 | 1 | 1 | 1 | 1 | 1 | 0,73 |
| | kcat_21 | $X_{34}$ | 27,6 | 0,2 | 10 | 0,20 | 0,20 | 0,20 | 0,20 | 0,20 | 1 | 0,20 |
| | kcat_23 | $X_{35}$ | 23,7 | 0,2 | 10 | 0,20 | 0,20 | 0,20 | 0,20 | 0,20 | 1 | 0,22 |
| | kcat_24 | $X_{36}$ | 100,3 | 0,2 | 10 | 0,24 | 0,20 | 0,24 | 0,24 | 0,24 | 1,19 | 0,20 |
| | kcat_68 | $X_{27}$ | 1740 | 0,2 | 10 | 1 | 1 | 1 | 1 | 1 | 1 | 0,72 |
| | kcat_11/ kcat_16 | $X_{26}/X_{32}$ | 2.22 | - | - | 5,22 | 13,32 | 13,32 | 13,32 | 13,32 | 2,22 | 3,87 |

| | | | | | | | | | | | |
|---|---|---|---|---|---|---|---|---|---|---|---|
| Piruvate (mM) | $X_{17}$ | 2,67 | 0,9 | 1,1 | 0,90 | 0,90 | 0,90 | 0,90 | 0,90 | 1 | 0,90 |
| G3P | $X_{18}$ | 0,041 | 0,9 | 1,1 | 0,90 | 0,90 | 0,90 | 0,90 | 0,90 | 1 | 0,90 |
| ATP | $X_{19}$ | 3 | 0,9 | 1,1 | 0,90 | 0,90 | 0,90 | 0,90 | 0,90 | 1 | 0,90 |
| NADPH | $X_{22}$ | 0,12 | 0,9 | 1,1 | 0,90 | 0,90 | 0,90 | 0,90 | 0,90 | 1 | 0,90 |
| CTP | $X_{25}$ | 2,7 | 0,9 | 1,1 | 0,90 | 0,90 | 1,10 | 0,90 | 0,90 | 1 | 0,90 |
| FldA | $X_{71}$ | 0,003015 | 0,9 | 1,1 | 1,08 | 1,10 | 1,10 | 1,10 | 1,10 | 1 | 0,90 |
| Km_1_68 (mM) | $X_{38}$ | 0,045 | 0,9 | 1,1 | 1 | 1 | 1 | 1 | 1 | 1 | 0,90 |
| Km_2_12 | $X_{39}$ | 0,15 | 0,9 | 1,1 | 1 | 1 | 1 | 1 | 1 | 1 | 0,90 |
| Km_3_14 | $X_{40}$ | 0,339 | 0,9 | 1,1 | 1 | 1 | 1 | 1 | 1 | 1 | 0,90 |
| Km_4_15 | $X_{41}$ | 0,0095 | 0,9 | 1,1 | 1 | 1 | 1 | 1 | 0,90 | 0,90 | 0,90 |
| Km_4_16 | $X_{46}$ | 0,0013 | 0,9 | 1,1 | 1 | 1,10 | 1,10 | 1,10 | 1,10 | 1 | 0,90 |
| Km_4_11 | $X_{56}$ | 0,0103 | 0,9 | 1,1 | 1 | 0,90 | 0,90 | 0,90 | 0,90 | 1 | 0,90 |
| Km_5_15 | $X_{42}$ | 0,0143 | 0,9 | 1,1 | 1 | 1 | 1 | 1 | 1,10 | 1,10 | 0,90 |
| Km_5_16 | $X_{47}$ | 0,0293 | 0,9 | 1,1 | 1 | 1,10 | 1,10 | 1,10 | 1,10 | 1 | 0,90 |
| Km_6_20 | $X_{43}$ | 0,00314 | 0,9 | 1,1 | 1 | 1 | 1 | 1 | 1 | 1 | 0,90 |
| Km_7_13 | $X_{60}$ | 0,00495 | 0,9 | 1,1 | 1 | 1 | 1 | 1 | 1 | 1 | 0,90 |
| Km_7_23 | $X_{44}$ | 0,56 | 0,9 | 1,1 | 1 | 1 | 1 | 1 | 1 | 1 | 0,90 |
| Km_8_21 | $X_{45}$ | 0,03 | 0,9 | 1,1 | 1 | 1 | 1 | 1 | 1 | 1 | 1,10 |
| Km_9_11 | $X_{37}$ | 0,0055 | 0,9 | 1,1 | 1 | 0,90 | 0,90 | 0,90 | 0,90 | 1 | 0,90 |
| Km_17_24 | $X_{48}$ | 0,134 | 0,9 | 1,1 | 1 | 1,10 | 1 | 1 | 1 | 0,91 | 1,10 |
| Km_18_24 | $X_{49}$ | 0,041 | 0,9 | 1,1 | 1 | 1,10 | 1 | 1 | 1 | 0,91 | 1,10 |
| Km_19_12 | $X_{50}$ | 0,42 | 0,9 | 1,1 | 1 | 1 | 1 | 1 | 1 | 1 | 0,90 |
| Km_71_21 | $X_{51}$ | 0,012 | 0,9 | 1,1 | 1 | 1 | 1 | 1 | 1 | 1 | 1,10 |
| Km_71_23 | $X_{52}$ | 0,012 | 0,9 | 1,1 | 1 | 1 | 1 | 1 | 1 | 1 | 0,90 |
| Km_22_68 | $X_{53}$ | 0,0005 | 0,9 | 1,1 | 1 | 1 | 1 | 1 | 1 | 1 | 0,90 |
| Km_25_20 | $X_{54}$ | 0,76 | 0,9 | 1,1 | 1 | 1 | 1 | 1 | 1 | 1 | 0,90 |
| K_10_55 (mM) | $X_{55}$ | 0,05 | 0,9 | 1,1 | 1,10 | 1,10 | 1,10 | 1,10 | 1,10 | 1 | 0,90 |
| K_4_58 | $X_{58}$ | 0,005 | 0,9 | 1,1 | 0,90 | 0,90 | 0,90 | 0,90 | 0,90 | 1 | 1,10 |
| K_9_72 | $X_{72}$ | 0,005 | 0,9 | 1,1 | **0,90** | **0,90** | **0,90** | **0,90** | **0,90** | **1** | **1,10** |
| Ki_4_24 | $X_{57}$ | 0,0654 | 1 | 1 | 1 | 1 | 1 | 1 | 1 | 1 | 1 |
| Ki_5_24 | $X_{59}$ | 0,0813 | 1 | 1 | 1 | 1 | 1 | 1 | 1 | 1 | 1 |
| Ka_6_14 | $X_{69}$ | 0,00339 | 0,9 | 1,1 | 1 | 1 | 1 | 1 | 1 | 1 | 1 |
| $\alpha_6$ (adimensional) | $X_{61}$ | 1 | 0,9 | 1,1 | 1 | 1 | 1 | 1 | 1 | 1 | 1 |
| $\beta_6$(adimensional) | $X_{63}$ | 2000 | 0,9 | 1,1 | 1 | 1 | 1 | 1 | 1 | 1 | 1 |
| Kd_9_11 | $X_{62}$ | 0,0714 | 0,9 | 1,1 | **1** | **0,90** | **0,90** | **0,90** | **0,90** | **1** | **0,90** |
| Kd_17_24 | $X_{64}$ | 0,0426 | 0,9 | 1,1 | 1 | 1,10 | 1 | 1 | 1 | 0,91 | 1,10 |
| Kd_22_68 | $X_{65}$ | 0,000333333 | 0,9 | 1,1 | 1 | 1 | 1 | 1 | 1 | 1 | 0,90 |
| Kd_71_23 | $X_{66}$ | 0,0005 | 0,9 | 1,1 | 1 | 1 | 1 | 1 | 1 | 1 | 0,90 |
| Kd_71_21 | $X_{67}$ | 0,0005 | 0,9 | 1,1 | 1 | 1 | 1 | 1 | 1 | 1 | 1,10 |
| hill_7_13 (adimensional) | $X_{70}$ | 1 | 0,9 | 1,1 | 1 | 1 | 1 | 1 | 1 | 1 | 1 |
| $(v_{4,11})_{opt}/(v_{4,11})_{basal}$ | | | | | 14,35 | 21,22 | 21,22 | 21,22 | 21,22 | 21,22 | 21,22 |

Time-independent

***a***) Time dependent metabolites ($X_1$,$X_2$, $X_6$,$X_{10}$) ranges between 0.8 and 1.2 their respective basal values and ($X_3$,…,$X_5$, $X_7$,…,$X_9$) are restrained to values between 0.1 and 2.5 their respective basal values.

***b*)** External variables ($X_{17}$, $X_{18}$, $X_{22}$, $X_{25}$, and $X_{71}$) are allowed bascule 10% around their respective basal values

***c*)** The enzymes ($X_{11}$...$X_{16,}$$X_{20,}$ $X_{21,}$ $X_{23,}$ $X_{24,}$ $X_{68}$) ranges between 1 and 10 times their basal values

***d*)** The kcat's ($X_{27}$...$X_{31,}$$X_{33}$... $X_{36}$) ranges between 0.2 and 10 times their basal values.

***e*)** The Km's, Kd's, Ki's, and $K_{0,5}$'s variables are not allowed be modified

***f*)** Cellular economy factor p = 2

***g*)** The $v_{9,11}$ basal flux was calculated as:

$$v_{9,11} = \frac{V_{max_{11}} \times X_4 \times X_9}{\left(K_{m_{4,11}} \times X_9 + K_{m_{9,11}} \times X_4 + X_4 \times X_9 + K_{d_{9,11}} \times K_{m_{9,11}}\right)} = 2.83 \times 10^{-4}\ mM \cdot min^{-1}$$

**Table S3.9b.** Specific constants (kcat/Km) fold change respect their respective basal values for data from Table S3.9a

| | | | Opt./Basal | | | | | | |
|---|---|---|---|---|---|---|---|---|---|
| **kcat / Km** | **Model symbols** | **Basal** | **5** | **6** | **7** | **8** | **9** | **10** | **11** |
| k_11/Km_4_11 | X26/ X56 | 168,116505 | 1 | 1,33 | 1,33 | 1,33 | 1,33 | 1,33 | 1,33 |
| k_11/Km_9_11 | X26 / X37 | 314,836364 | 1 | 1,33 | 1,33 | 1,33 | 1,33 | 1,33 | 1,33 |
| k_12 / Km_2_12 | X28 / X39 | 6733,33333 | 1 | 1 | 1 | 1 | 1 | 0,89 | 1,07 |
| k_12 / Km_19_12 | X28 / X50 | 2404,7619 | 1 | 1 | 1 | 1 | 1 | 0,89 | 1,07 |
| k_14 / Km_3_14 | X30 / X40 | 238,053097 | 1 | 1 | 1 | 1 | 1 | 1 | 1,26 |
| k_15 / Km_4_15 | X31 / X41 | 2084,21053 | 1 | 1 | 1,29 | 1 | 2,50 | 1,29 | 5,93 |
| k_15 / Km_5_15 | X31 / X42 | 1384,61538 | 1 | 1 | 1,29 | 1 | 2,05 | 1,29 | 5,93 |
| k_16 / Km_4_16 | X32 / X46 | 600 | 0,42 | 0,18 | 0,18 | 0,18 | 0,18 | 0,18 | 0,76 |
| k_16 / Km_5_16 | X32/ X47 | 26,6211604 | 0,42 | 0,18 | 0,18 | 0,18 | 0,18 | 0,18 | 0,76 |
| k_20 / Km_6_20 | X33 / X43 | 494904,459 | 1 | 1 | 1 | 1 | 1 | 0,97 | 0,82 |
| k_20 / Km_25_20 | X33 / X54 | 2044,73684 | 1 | 1 | 1 | 1 | 1 | 0,97 | 0,82 |
| k_21 / Km_8_21 | X34 / X45 | 920 | 0,20 | 0,20 | 0,20 | 0,20 | 0,20 | 0,20 | 0,18 |
| k_21 / Km_71_21 | X34 / X51 | 2300 | 0,20 | 0,20 | 0,20 | 0,20 | 0,20 | 0,20 | 0,18 |
| k_23 / Km_7_23 | X35 / X44 | 42,3214286 | 0,20 | 0,20 | 0,20 | 0,20 | 0,20 | 0,20 | 0,25 |
| k_23 / Km_71_23 | X35 / X52 | 1975 | 0,20 | 0,20 | 0,20 | 0,20 | 0,20 | 0,20 | 0,25 |
| k_24 / Km_17_24 | X36 / X48 | 748,507463 | 0,24 | 0,18 | 0,24 | 0,24 | 0,24 | 0,24 | 0,18 |
| k_24/ Km_18_24 | X36 / X49 | 2446,34146 | 0,24 | 0,18 | 0,24 | 0,24 | 0,24 | 0,24 | 0,18 |
| k_68/ Km_1_68 | X27 / X38 | 38666,6667 | 1 | 1 | 1 | 1 | 1 | 1 | 0,80 |
| k_68/ Km_22_68 | X27 / X53 | 3480000 | 1 | 1 | 1 | 1 | 1 | 1 | 0,80 |

### S3.7 - Optimizations equations

Model for Table S3.7a where the enzymes were allowed to vary between 1 and 10 times their nominal value. The software is presented in Lingo input format [1]

```
! Max flux U_4_11: ;

MAX = y11 + y26 - 0.0029399295211171745656475451369261*y37 +
0.99685126702718862556906742611318*y4 - 0.99685126702718851454676496359752*y56 -
0.99142718740203761296214679532568*y62 + 0.99436711692315482569171081195236*y9;

0.03191546745632127535241551186118 2*y17 - 0.94691325449856866836029212208814*y1 +
0.49473931416526567561575689069287*y18 - 0.0028433053581015297305023548091185*y22 + y24 -
y27 + y36 + 0.94691325449856866836029212208814*y38 -
0.0029634277888853960587767222278899*y4 - 0.024145838766709847716951387042172*y48 -
0.49473931416526550908230319691938*y49 - 0.0029848248944612772518003573907208*y5 +
0.00022027695228809667524552673434357*y53 + 0.0029634277888853956250958532336881*y57 +
0.0029848248944612781191620953791244*y59 - 0.0077696286896115586070865610679448*y64 +
0.0026230284058132088964576666967332*y65 - y68 = -0.63939223492636604224501046176572 -
0.000000000000000188163287034859294521993433590 98*K;

y20 - 0.0019616560150146700527984400252 99*y19 - 0.98402651530630902421137307101162*y2 -
y12 + 0.010663710264075591110932705873893*y25 - y28 + y33 +
0.98402651530630913523367553352728*y39 - 0.95145205590302495135546223536949*y43 +
0.0019616560150146678843940950542901*y50 - 0.010663710264075428046925964054026*y54 +
0.95145205590302506237776469788514*y6 = -0.13190177115048234492332979794 91+
0.000000000000000382940207321880166091432329267 26*K;

y12 - y14 + 0.0019616560150146700527984400252 99*y19 +
0.98402651530630902421137307101162*y2 + y28 - 0.99544244307065632426656520692632*y3 - y30
- 0.98402651530630913523367553352728*y39 + 0.99544244307065654631117013195762*y40 -
0.0019616560150146678843940950542901*y50 - 0.95105510287742234254437789786607*y6 +
0.98646207814079334141865729179699*y61 - 0.98662418626324865922327944645076*y63 +
0.95105510287742245356668036038172*y69 = 0.023846158824983119221091751950234 -
0.0001621081224550935916128846514539*K;

0.22249820426599241363074810351463*y15 - 0.012868518561875347261391766551242*y11 -
0.43859130784133681935088588943472*y16 + 0.66108951210732957992632918831077*y21 -
0.012868518561875347261391766551242*y26 + 0.22249820426599231648623344881344*y31 -
0.43859130784133681935088588943472*y32 + 0.66108951210732957992632918831077*y34 +
0.000037832537613101666670433614880764*y37 - 0.98572411499323963485208466295262*y4 +
0.51804988331971001114251151875578*y41 - 0.72474519382444890780448076839093*y42 -
0.65749183019860180099414037613315*y45 + 0.022718342670251568532480135331753*y46 +
0.40894378050542273816958527277166*y47 + 0.31580141331902610718485036045422*y5 -
0.0028752200594027352807680042445782*y51 + 0.012827999033168335785148883587681*y56 -
0.43212788997010975755586059676716*y58 + 0.012758199163830990022217015678052*y62 -
0.093519601646064556832449454759626*y67 + 0.096394821705467315098303515696898*y71 +
0.65749183019860202303874530116445*y8 - 0.012796031701444091424613169749591*y9 = -
0.43604493387723546236870542062071+ -0.0000000000000000626412000210354147355928944 3074*K;

0.85027393600193812428500450550928*y21 - 0.56410327313428376694304233751609*y16 -
0.28617066286765435734196216799319*y15 - 0.28617066286765419080850847421971*y31 -
0.56410327313428376694304233751609*y32 + 0.85027393600193812428500450550928*y34 +
0.63708089597637018730580393821583*y4 - 0.66630056182782237961248483770 76*y41 +
0.93214600635133815220711994697922*y42 - 0.84564670307660272374761234459584*y45 +
0.029219665851451984139863782274915*y46 + 0.52597149324826431016077776803286*y47 -
1.4581174995996024623678977150121*y5 - 0.0036980236896923912408297052678563*y51 -
0.12028216804023779407106786720 76*y67 + 0.12398019172993023084838881686665*y71 +
0.84564670307660294579221726962715*y8 = -0.63598026053481041032333234819626+
```

```
0.00000000000000022594773274597912404715316370l3*K;
```

```
y13-(@log(1.476035e-005*UB13)+K)*W13-(@log(1.476035e-005)+K)*M13 <=0;
y13-(@log(1.476035e-005*LB13)+K)*W13-(@log(1.476035e-005)+K)*M13 >=0;
y14-(@log(5.119995e-005*UB14)+K)*W14-(@log(5.119995e-005)+K)*M14 <=0;
y14-(@log(5.119995e-005*LB14)+K)*W14-(@log(5.119995e-005)+K)*M14 >=0;
y15-(@log(2.767565e-006*UB15)+K)*W15-(@log(2.767565e-006)+K)*M15 <=0;
y15-(@log(2.767565e-006*LB15)+K)*W15-(@log(2.767565e-006)+K)*M15 >=0;
y16-(@log(1.632863e-004*UB16)+K)*W16-(@log(1.632863e-004)+K)*M16 <=0;
y16-(@log(1.632863e-004*LB16)+K)*W16-(@log(1.632863e-004)+K)*M16 >=0;
y17-(@log(2.670000e+000*UB17)+K)*W17-(@log(2.670000e+000)+K)*M17 <=0;
y17-(@log(2.670000e+000*LB17)+K)*W17-(@log(2.670000e+000)+K)*M17 >=0;
y18-(@log(4.100000e-002*UB18)+K)*W18-(@log(4.100000e-002)+K)*M18 <=0;
y18-(@log(4.100000e-002*LB18)+K)*W18-(@log(4.100000e-002)+K)*M18 >=0;
y19-(@log(3*UB19)+K)*W19-(@log(3)+K)*M19 <=0;
y19-(@log(3*LB19)+K)*W19-(@log(3)+K)*M19 >=0;
y20-(@log(2.306304e-005*UB20)+K)*W20-(@log(2.306304e-005)+K)*M20 <=0;
y20-(@log(2.306304e-005*LB20)+K)*W20-(@log(2.306304e-005)+K)*M20 >=0;
y21-(@log(1.005549e-004*UB21)+K)*W21-(@log(1.005549e-004)+K)*M21 <=0;
y21-(@log(1.005549e-004*LB21)+K)*W21-(@log(1.005549e-004)+K)*M21 >=0;
y22-(@log(1.200000e-001*UB22)+K)*W22-(@log(1.200000e-001)+K)*M22 <=0;
y22-(@log(1.200000e-001*LB22)+K)*W22-(@log(1.200000e-001)+K)*M22 >=0;
y23-(@log(9.409721e-005*UB23)+K)*W23-(@log(9.409721e-005)+K)*M23 <=0;
y23-(@log(9.409721e-005*LB23)+K)*W23-(@log(9.409721e-005)+K)*M23 >=0;
y24-(@log(2.813691e-005*UB24)+K)*W24-(@log(2.813691e-005)+K)*M24 <=0;
y24-(@log(2.813691e-005*LB24)+K)*W24-(@log(2.813691e-005)+K)*M24 >=0;
y25-(@log(2.700000e+000*UB25)+K)*W25-(@log(2.700000e+000)+K)*M25 <=0;
y25-(@log(2.700000e+000*LB25)+K)*W25-(@log(2.700000e+000)+K)*M25 >=0;
y26-(@log(1.731600e+000*UB26)+K)*W26-(@log(1.731600e+000)+K)*M26 <=0;
y26-(@log(1.731600e+000*LB26)+K)*W26-(@log(1.731600e+000)+K)*M26 >=0;
y27-(@log(1740*UB27)+K)*W27-(@log(1740)+K)*M27 <=0;
y27-(@log(1740*LB27)+K)*W27-(@log(1740)+K)*M27 >=0;
y28-(@log(1010*UB28)+K)*W28-(@log(1010)+K)*M28 <=0;
y28-(@log(1010*LB28)+K)*W28-(@log(1010)+K)*M28 >=0;
y29-(@log(600*UB29)+K)*W29-(@log(600)+K)*M29 <=0;
y29-(@log(600*LB29)+K)*W29-(@log(600)+K)*M29 >=0;
y30-(@log(8.070000e+001*UB30)+K)*W30-(@log(8.070000e+001)+K)*M30 <=0;
y30-(@log(8.070000e+001*LB30)+K)*W30-(@log(8.070000e+001)+K)*M30 >=0;
y31-(@log(1.980000e+001*UB31)+K)*W31-(@log(1.980000e+001)+K)*M31 <=0;
y31-(@log(1.980000e+001*LB31)+K)*W31-(@log(1.980000e+001)+K)*M31 >=0;
y32-(@log(7.800000e-001*UB32)+K)*W32-(@log(7.800000e-001)+K)*M32 <=0;
y32-(@log(7.800000e-001*LB32)+K)*W32-(@log(7.800000e-001)+K)*M32 >=0;
y33-(@log(1554*UB33)+K)*W33-(@log(1554)+K)*M33 <=0;
y33-(@log(1554*LB33)+K)*W33-(@log(1554)+K)*M33 >=0;
y34-(@log(2.760000e+001*UB34)+K)*W34-(@log(2.760000e+001)+K)*M34 <=0;
y34-(@log(2.760000e+001*LB34)+K)*W34-(@log(2.760000e+001)+K)*M34 >=0;
y35-(@log(2.370000e+001*UB35)+K)*W35-(@log(2.370000e+001)+K)*M35 <=0;
y35-(@log(2.370000e+001*LB35)+K)*W35-(@log(2.370000e+001)+K)*M35 >=0;
y36-(@log(1.003000e+002*UB36)+K)*W36-(@log(1.003000e+002)+K)*M36 <=0;
y36-(@log(1.003000e+002*LB36)+K)*W36-(@log(1.003000e+002)+K)*M36 >=0;
y37-(@log(5.500000e-003*UB37)+K)*W37-(@log(5.500000e-003)+K)*M37 <=0;
y37-(@log(5.500000e-003*LB37)+K)*W37-(@log(5.500000e-003)+K)*M37 >=0;
y38-(@log(4.500000e-002*UB38)+K)*W38-(@log(4.500000e-002)+K)*M38 <=0;
y38-(@log(4.500000e-002*LB38)+K)*W38-(@log(4.500000e-002)+K)*M38 >=0;
y39-(@log(1.500000e-001*UB39)+K)*W39-(@log(1.500000e-001)+K)*M39 <=0;
y39-(@log(1.500000e-001*LB39)+K)*W39-(@log(1.500000e-001)+K)*M39 >=0;
y40-(@log(3.390000e-001*UB40)+K)*W40-(@log(3.390000e-001)+K)*M40 <=0;
y40-(@log(3.390000e-001*LB40)+K)*W40-(@log(3.390000e-001)+K)*M40 >=0;
y41-(@log(9.500000e-003*UB41)+K)*W41-(@log(9.500000e-003)+K)*M41 <=0;
y41-(@log(9.500000e-003*LB41)+K)*W41-(@log(9.500000e-003)+K)*M41 >=0;
y42-(@log(1.430000e-002*UB42)+K)*W42-(@log(1.430000e-002)+K)*M42 <=0;
y42-(@log(1.430000e-002*LB42)+K)*W42-(@log(1.430000e-002)+K)*M42 >=0;
y43-(@log(3.140000e-003*UB43)+K)*W43-(@log(3.140000e-003)+K)*M43 <=0;
y43-(@log(3.140000e-003*LB43)+K)*W43-(@log(3.140000e-003)+K)*M43 >=0;
y44-(@log(5.600000e-001*UB44)+K)*W44-(@log(5.600000e-001)+K)*M44 <=0;
y44-(@log(5.600000e-001*LB44)+K)*W44-(@log(5.600000e-001)+K)*M44 >=0;
y45-(@log(3.000000e-002*UB45)+K)*W45-(@log(3.000000e-002)+K)*M45 <=0;
y45-(@log(3.000000e-002*LB45)+K)*W45-(@log(3.000000e-002)+K)*M45 >=0;
y46-(@log(1.300000e-003*UB46)+K)*W46-(@log(1.300000e-003)+K)*M46 <=0;
```

```
y46-(@log(1.300000e-003*LB46)+K)*W46-(@log(1.300000e-003)+K)*M46 >=0;
y47-(@log(2.930000e-002*UB47)+K)*W47-(@log(2.930000e-002)+K)*M47 <=0;
y47-(@log(2.930000e-002*LB47)+K)*W47-(@log(2.930000e-002)+K)*M47 >=0;
y48-(@log(1.340000e-001*UB48)+K)*W48-(@log(1.340000e-001)+K)*M48 <=0;
y48-(@log(1.340000e-001*LB48)+K)*W48-(@log(1.340000e-001)+K)*M48 >=0;
y49-(@log(4.100000e-002*UB49)+K)*W49-(@log(4.100000e-002)+K)*M49 <=0;
y49-(@log(4.100000e-002*LB49)+K)*W49-(@log(4.100000e-002)+K)*M49 >=0;
y50-(@log(4.200000e-001*UB50)+K)*W50-(@log(4.200000e-001)+K)*M50 <=0;
y50-(@log(4.200000e-001*LB50)+K)*W50-(@log(4.200000e-001)+K)*M50 >=0;
y51-(@log(1.200000e-002*UB51)+K)*W51-(@log(1.200000e-002)+K)*M51 <=0;
y51-(@log(1.200000e-002*LB51)+K)*W51-(@log(1.200000e-002)+K)*M51 >=0;
y52-(@log(1.200000e-002*UB52)+K)*W52-(@log(1.200000e-002)+K)*M52 <=0;
y52-(@log(1.200000e-002*LB52)+K)*W52-(@log(1.200000e-002)+K)*M52 >=0;
y53-(@log(5.000000e-004*UB53)+K)*W53-(@log(5.000000e-004)+K)*M53 <=0;
y53-(@log(5.000000e-004*LB53)+K)*W53-(@log(5.000000e-004)+K)*M53 >=0;
y54-(@log(7.600000e-001*UB54)+K)*W54-(@log(7.600000e-001)+K)*M54 <=0;
y54-(@log(7.600000e-001*LB54)+K)*W54-(@log(7.600000e-001)+K)*M54 >=0;
y55-(@log(5.000000e-002*UB55)+K)*W55-(@log(5.000000e-002)+K)*M55 <=0;
y55-(@log(5.000000e-002*LB55)+K)*W55-(@log(5.000000e-002)+K)*M55 >=0;
y56-(@log(1.030000e-002*UB56)+K)*W56-(@log(1.030000e-002)+K)*M56 <=0;
y56-(@log(1.030000e-002*LB56)+K)*W56-(@log(1.030000e-002)+K)*M56 >=0;
y57-(@log(6.540000e-002*UB57)+K)*W57-(@log(6.540000e-002)+K)*M57 <=0;
y57-(@log(6.540000e-002*LB57)+K)*W57-(@log(6.540000e-002)+K)*M57 >=0;
y58-(@log(5.000000e-003*UB58)+K)*W58-(@log(5.000000e-003)+K)*M58 <=0;
y58-(@log(5.000000e-003*LB58)+K)*W58-(@log(5.000000e-003)+K)*M58 >=0;
y59-(@log(8.130000e-002*UB59)+K)*W59-(@log(8.130000e-002)+K)*M59 <=0;
y59-(@log(8.130000e-002*LB59)+K)*W59-(@log(8.130000e-002)+K)*M59 >=0;
y60-(@log(4.950000e-003*UB60)+K)*W60-(@log(4.950000e-003)+K)*M60 <=0;
y60-(@log(4.950000e-003*LB60)+K)*W60-(@log(4.950000e-003)+K)*M60 >=0;
y61-(@log(1*UB61)+K)*W61-(@log(1)+K)*M61 <=0;
y61-(@log(1*LB61)+K)*W61-(@log(1)+K)*M61 >=0;
y62-(@log(7.140000e-002*UB62)+K)*W62-(@log(7.140000e-002)+K)*M62 <=0;
y62-(@log(7.140000e-002*LB62)+K)*W62-(@log(7.140000e-002)+K)*M62 >=0;
y63-(@log(2000*UB63)+K)*W63-(@log(2000)+K)*M63 <=0;
y63-(@log(2000*LB63)+K)*W63-(@log(2000)+K)*M63 >=0;
y64-(@log(4.260000e-002*UB64)+K)*W64-(@log(4.260000e-002)+K)*M64 <=0;
y64-(@log(4.260000e-002*LB64)+K)*W64-(@log(4.260000e-002)+K)*M64 >=0;
y65-(@log(3.333333e-004*UB65)+K)*W65-(@log(3.333333e-004)+K)*M65 <=0;
y65-(@log(3.333333e-004*LB65)+K)*W65-(@log(3.333333e-004)+K)*M65 >=0;
y66-(@log(5.000000e-004*UB66)+K)*W66-(@log(5.000000e-004)+K)*M66 <=0;
y66-(@log(5.000000e-004*LB66)+K)*W66-(@log(5.000000e-004)+K)*M66 >=0;
y67-(@log(5.000000e-004*UB67)+K)*W67-(@log(5.000000e-004)+K)*M67 <=0;
y67-(@log(5.000000e-004*LB67)+K)*W67-(@log(5.000000e-004)+K)*M67 >=0;
y68-(@log(1.476035e-005*UB68)+K)*W68-(@log(1.476035e-005)+K)*M68 <=0;
y68-(@log(1.476035e-005*LB68)+K)*W68-(@log(1.476035e-005)+K)*M68 >=0;
y69-(@log(3.390000e-003*UB69)+K)*W69-(@log(3.390000e-003)+K)*M69 <=0;
y69-(@log(3.390000e-003*LB69)+K)*W69-(@log(3.390000e-003)+K)*M69 >=0;
y70-(@log(1*UB70)+K)*W70-(@log(1)+K)*M70 <=0;
y70-(@log(1*LB70)+K)*W70-(@log(1)+K)*M70 >=0;
y71-(@log(3.015262e-003*UB71)+K)*W71-(@log(3.015262e-003)+K)*M71 <=0;
y71-(@log(3.015262e-003*LB71)+K)*W71-(@log(3.015262e-003)+K)*M71 >=0;
y72-(@log(5.000000e-003*UB72)+K)*W72-(@log(5.000000e-003)+K)*M72 <=0;
y72-(@log(5.000000e-003*LB72)+K)*W72-(@log(5.000000e-003)+K)*M72 >=0;

! Only to optimize the enzymes: ;
y38-(@log(4.500000e-002*UB38)+K)*W68-(@log(4.500000e-002)+K)*M68 <=0;
y38-(@log(4.500000e-002*LB38)+K)*W68-(@log(4.500000e-002)+K)*M68 >=0;
y53-(@log(5.000000e-004*UB53)+K)*W68-(@log(5.000000e-004)+K)*M68 <=0;
y53-(@log(5.000000e-004*LB53)+K)*W68-(@log(5.000000e-004)+K)*M68 >=0;
y65-(@log(3.333333e-004*UB65)+K)*W68-(@log(3.333333e-004)+K)*M68 <=0;
y65-(@log(3.333333e-004*LB65)+K)*W68-(@log(3.333333e-004)+K)*M68 >=0;
y39-(@log(1.500000e-001*UB39)+K)*W12-(@log(1.500000e-001)+K)*M12 <=0;
y39-(@log(1.500000e-001*LB39)+K)*W12-(@log(1.500000e-001)+K)*M12 >=0;
y50-(@log(4.200000e-001*UB50)+K)*W12-(@log(4.200000e-001)+K)*M12 <=0;
y50-(@log(4.200000e-001*LB50)+K)*W12-(@log(4.200000e-001)+K)*M12 >=0;
y60-(@log(4.950000e-003*UB60)+K)*W13-(@log(4.950000e-003)+K)*M13 <=0;
y60-(@log(4.950000e-003*LB60)+K)*W13-(@log(4.950000e-003)+K)*M13 >=0;
```

```
y40-(@log(3.390000e-001*UB40)+K)*W14-(@log(3.390000e-001)+K)*M14 <=0;
y40-(@log(3.390000e-001*LB40)+K)*W14-(@log(3.390000e-001)+K)*M14 >=0;
y69-(@log(3.390000e-003*UB69)+K)*W14-(@log(3.390000e-003)+K)*M14 <=0;
y69-(@log(3.390000e-003*LB69)+K)*W14-(@log(3.390000e-003)+K)*M14 >=0;
y41-(@log(9.500000e-003*UB41)+K)*W15-(@log(9.500000e-003)+K)*M15 <=0;
y41-(@log(9.500000e-003*LB41)+K)*W15-(@log(9.500000e-003)+K)*M15 >=0;
y42-(@log(1.430000e-002*UB42)+K)*W15-(@log(1.430000e-002)+K)*M15 <=0;
y42-(@log(1.430000e-002*LB42)+K)*W15-(@log(1.430000e-002)+K)*M15 >=0;
y37-(@log(5.500000e-003*UB37)+K)*W16-(@log(5.500000e-003)+K)*M16 <=0;
y37-(@log(5.500000e-003*LB37)+K)*W16-(@log(5.500000e-003)+K)*M16 >=0;
y46-(@log(1.300000e-003*UB46)+K)*W16-(@log(1.300000e-003)+K)*M16 <=0;
y46-(@log(1.300000e-003*LB46)+K)*W16-(@log(1.300000e-003)+K)*M16 >=0;
y47-(@log(2.930000e-002*UB47)+K)*W16-(@log(2.930000e-002)+K)*M16 <=0;
y47-(@log(2.930000e-002*LB47)+K)*W16-(@log(2.930000e-002)+K)*M16 >=0;
y56-(@log(1.030000e-002*UB56)+K)*W16-(@log(1.030000e-002)+K)*M16 <=0;
y56-(@log(1.030000e-002*LB56)+K)*W16-(@log(1.030000e-002)+K)*M16 >=0;
y62-(@log(7.140000e-002*UB62)+K)*W16-(@log(7.140000e-002)+K)*M16 <=0;
y62-(@log(7.140000e-002*LB62)+K)*W16-(@log(7.140000e-002)+K)*M16 >=0;
y43-(@log(3.140000e-003*UB43)+K)*W20-(@log(3.140000e-003)+K)*M20 <=0;
y43-(@log(3.140000e-003*LB43)+K)*W20-(@log(3.140000e-003)+K)*M20 >=0;
y54-(@log(7.600000e-001*UB54)+K)*W20-(@log(7.600000e-001)+K)*M20 <=0;
y54-(@log(7.600000e-001*LB54)+K)*W20-(@log(7.600000e-001)+K)*M20 >=0;
y45-(@log(3.000000e-002*UB45)+K)*W21-(@log(3.000000e-002)+K)*M21 <=0;
y45-(@log(3.000000e-002*LB45)+K)*W21-(@log(3.000000e-002)+K)*M21 >=0;
y51-(@log(1.200000e-002*UB51)+K)*W21-(@log(1.200000e-002)+K)*M21 <=0;
y51-(@log(1.200000e-002*LB51)+K)*W21-(@log(1.200000e-002)+K)*M21 >=0;
y67-(@log(5.000000e-004*UB67)+K)*W21-(@log(5.000000e-004)+K)*M21 <=0;
y67-(@log(5.000000e-004*LB67)+K)*W21-(@log(5.000000e-004)+K)*M21 >=0;
y44-(@log(5.600000e-001*UB44)+K)*W23-(@log(5.600000e-001)+K)*M23 <=0;
y44-(@log(5.600000e-001*LB44)+K)*W23-(@log(5.600000e-001)+K)*M23 >=0;
y52-(@log(1.200000e-002*UB52)+K)*W23-(@log(1.200000e-002)+K)*M23 <=0;
y52-(@log(1.200000e-002*LB52)+K)*W23-(@log(1.200000e-002)+K)*M23 >=0;
y66-(@log(5.000000e-004*UB66)+K)*W23-(@log(5.000000e-004)+K)*M23 <=0;
y66-(@log(5.000000e-004*LB66)+K)*W23-(@log(5.000000e-004)+K)*M23 >=0;
y48-(@log(1.340000e-001*UB48)+K)*W24-(@log(1.340000e-001)+K)*M24 <=0;
y48-(@log(1.340000e-001*LB48)+K)*W24-(@log(1.340000e-001)+K)*M24 >=0;
y49-(@log(4.100000e-002*UB49)+K)*W24-(@log(4.100000e-002)+K)*M24 <=0;
y49-(@log(4.100000e-002*LB49)+K)*W24-(@log(4.100000e-002)+K)*M24 >=0;
y57-(@log(6.540000e-002*UB57)+K)*W24-(@log(6.540000e-002)+K)*M24 <=0;
y57-(@log(6.540000e-002*LB57)+K)*W24-(@log(6.540000e-002)+K)*M24 >=0;
y59-(@log(8.130000e-002*UB59)+K)*W24-(@log(8.130000e-002)+K)*M24 <=0;
y59-(@log(8.130000e-002*LB59)+K)*W24-(@log(8.130000e-002)+K)*M24 >=0;
y64-(@log(4.260000e-002*UB64)+K)*W24-(@log(4.260000e-002)+K)*M24 <=0;
y64-(@log(4.260000e-002*LB64)+K)*W24-(@log(4.260000e-002)+K)*M24 >=0;

! Only to optimize the kcats: ;
y37-(@log(5.500000e-003*UB37)+K)*W26-(@log(5.500000e-003)+K)*M26 <=0;
y37-(@log(5.500000e-003*LB37)+K)*W26-(@log(5.500000e-003)+K)*M26 >=0;
y56-(@log(1.030000e-002*UB56)+K)*W26-(@log(1.030000e-002)+K)*M26 <=0;
y56-(@log(1.030000e-002*LB56)+K)*W26-(@log(1.030000e-002)+K)*M26 >=0;
y62-(@log(7.140000e-002*UB62)+K)*W26-(@log(7.140000e-002)+K)*M26 <=0;
y62-(@log(7.140000e-002*LB62)+K)*W26-(@log(7.140000e-002)+K)*M26 >=0;
y38-(@log(4.500000e-002*UB38)+K)*W27-(@log(4.500000e-002)+K)*M27 <=0;
y38-(@log(4.500000e-002*LB38)+K)*W27-(@log(4.500000e-002)+K)*M27 >=0;
y53-(@log(5.000000e-004*UB53)+K)*W27-(@log(5.000000e-004)+K)*M27 <=0;
y53-(@log(5.000000e-004*LB53)+K)*W27-(@log(5.000000e-004)+K)*M27 >=0;
y65-(@log(3.333333e-004*UB65)+K)*W27-(@log(3.333333e-004)+K)*M27 <=0;
y65-(@log(3.333333e-004*LB65)+K)*W27-(@log(3.333333e-004)+K)*M27 >=0;
y39-(@log(1.500000e-001*UB39)+K)*W28-(@log(1.500000e-001)+K)*M28 <=0;
y39-(@log(1.500000e-001*LB39)+K)*W28-(@log(1.500000e-001)+K)*M28 >=0;
y50-(@log(4.200000e-001*UB50)+K)*W28-(@log(4.200000e-001)+K)*M28 <=0;
y50-(@log(4.200000e-001*LB50)+K)*W28-(@log(4.200000e-001)+K)*M28 >=0;
y60-(@log(4.950000e-003*UB60)+K)*W29-(@log(4.950000e-003)+K)*M29 <=0;
y60-(@log(4.950000e-003*LB60)+K)*W29-(@log(4.950000e-003)+K)*M29 >=0;
y40-(@log(3.390000e-001*UB40)+K)*W30-(@log(3.390000e-001)+K)*M30 <=0;
y40-(@log(3.390000e-001*LB40)+K)*W30-(@log(3.390000e-001)+K)*M30 >=0;
y69-(@log(3.390000e-003*UB69)+K)*W30-(@log(3.390000e-003)+K)*M30 <=0;
```

```
y69-(@log(3.390000e-003*LB69)+K)*W30-(@log(3.390000e-003)+K)*M30 >=0;
y41-(@log(9.500000e-003*UB41)+K)*W31-(@log(9.500000e-003)+K)*M31 <=0;
y41-(@log(9.500000e-003*LB41)+K)*W31-(@log(9.500000e-003)+K)*M31 >=0;
y42-(@log(1.430000e-002*UB42)+K)*W31-(@log(1.430000e-002)+K)*M31 <=0;
y42-(@log(1.430000e-002*LB42)+K)*W31-(@log(1.430000e-002)+K)*M31 >=0;
y46-(@log(1.300000e-003*UB46)+K)*W32-(@log(1.300000e-003)+K)*M32 <=0;
y46-(@log(1.300000e-003*LB46)+K)*W32-(@log(1.300000e-003)+K)*M32 >=0;
y47-(@log(2.930000e-002*UB47)+K)*W32-(@log(2.930000e-002)+K)*M32 <=0;
y47-(@log(2.930000e-002*LB47)+K)*W32-(@log(2.930000e-002)+K)*M32 >=0;
y62-(@log(7.140000e-002*UB62)+K)*W32-(@log(7.140000e-002)+K)*M32 <=0;
y62-(@log(7.140000e-002*LB62)+K)*W32-(@log(7.140000e-002)+K)*M32 >=0;
y43-(@log(3.140000e-003*UB43)+K)*W33-(@log(3.140000e-003)+K)*M33 <=0;
y43-(@log(3.140000e-003*LB43)+K)*W33-(@log(3.140000e-003)+K)*M33 >=0;
y54-(@log(7.600000e-001*UB54)+K)*W33-(@log(7.600000e-001)+K)*M33 <=0;
y54-(@log(7.600000e-001*LB54)+K)*W33-(@log(7.600000e-001)+K)*M33 >=0;
y45-(@log(3.000000e-002*UB45)+K)*W34-(@log(3.000000e-002)+K)*M34 <=0;
y45-(@log(3.000000e-002*LB45)+K)*W34-(@log(3.000000e-002)+K)*M34 >=0;
y51-(@log(1.200000e-002*UB51)+K)*W34-(@log(1.200000e-002)+K)*M34 <=0;
y51-(@log(1.200000e-002*LB51)+K)*W34-(@log(1.200000e-002)+K)*M34 >=0;
y67-(@log(5.000000e-004*UB67)+K)*W34-(@log(5.000000e-004)+K)*M34 <=0;
y67-(@log(5.000000e-004*LB67)+K)*W34-(@log(5.000000e-004)+K)*M34 >=0;
y44-(@log(5.600000e-001*UB44)+K)*W35-(@log(5.600000e-001)+K)*M35 <=0;
y44-(@log(5.600000e-001*LB44)+K)*W35-(@log(5.600000e-001)+K)*M35 >=0;
y52-(@log(1.200000e-002*UB52)+K)*W35-(@log(1.200000e-002)+K)*M35 <=0;
y52-(@log(1.200000e-002*LB52)+K)*W35-(@log(1.200000e-002)+K)*M35 >=0;
y66-(@log(5.000000e-004*UB66)+K)*W35-(@log(5.000000e-004)+K)*M35 <=0;
y66-(@log(5.000000e-004*LB66)+K)*W35-(@log(5.000000e-004)+K)*M35 >=0;
y48-(@log(1.340000e-001*UB48)+K)*W36-(@log(1.340000e-001)+K)*M36 <=0;
y48-(@log(1.340000e-001*LB48)+K)*W36-(@log(1.340000e-001)+K)*M36 >=0;
y49-(@log(4.100000e-002*UB49)+K)*W36-(@log(4.100000e-002)+K)*M36 <=0;
y49-(@log(4.100000e-002*LB49)+K)*W36-(@log(4.100000e-002)+K)*M36 >=0;
y57-(@log(6.540000e-002*UB57)+K)*W36-(@log(6.540000e-002)+K)*M36 <=0;
y57-(@log(6.540000e-002*LB57)+K)*W36-(@log(6.540000e-002)+K)*M36 >=0;
y59-(@log(8.130000e-002*UB59)+K)*W36-(@log(8.130000e-002)+K)*M36 <=0;
y59-(@log(8.130000e-002*LB59)+K)*W36-(@log(8.130000e-002)+K)*M36 >=0;
y64-(@log(4.260000e-002*UB64)+K)*W36-(@log(4.260000e-002)+K)*M36 <=0;
y64-(@log(4.260000e-002*LB64)+K)*W36-(@log(4.260000e-002)+K)*M36 >=0;

!X17(P!yr),X18(G3P),X19(ATP),X22(NADPH),X25(CTP), X71(FldA).+/-10: ;
Const_LB = 0.9;     !0.9;
Const_UB = 1.1;     !1.1;

LB17= Const_LB;
LB18= Const_LB;
LB19= Const_LB;
LB22= Const_LB;
LB25= Const_LB;
LB71= Const_LB;

UB17= Const_UB;
UB18= Const_UB;
UB19= Const_UB;
UB22= Const_UB;
UB25= Const_UB;
UB71= Const_UB;

enz_LB = 1;   ! 1;
enz_UB = 10; ! 5 or 10;

LB11=enz_LB;
LB12=enz_LB;
LB13=enz_LB;
LB14=enz_LB;
LB15=enz_LB;
LB16=enz_LB;
LB20=enz_LB;
LB21=enz_LB;
```

```
LB23=enz_LB;
LB24=enz_LB;
LB68=enz_LB;

UB11=enz_UB;
UB12=enz_UB;
UB13=enz_UB;
UB14=enz_UB;
UB15=enz_UB;
UB16=enz_UB;
UB20=enz_UB;
UB21=enz_UB;
UB23=enz_UB;
UB24=enz_UB;
UB68=enz_UB;

!k_11,k_12,k_13,k_14,k_15,k_16,k_20,k_21,k_23,k_24,k_26,k_71:;
kcat_LB = 0.2;
kcat_UB = 10;

! kcat_11: ;
kcat_LB_26 = 0.2;
kcat_UB_26 = 1.2;

! kcat_16: ;
kcat_LB_32 = 0.2;
kcat_UB_32 = 1.2;

  LB26= kcat_LB_26;
LB27= kcat_LB;
LB28= kcat_LB;
LB29= kcat_LB;
LB30= kcat_LB;
LB31= kcat_LB;
  LB32= kcat_LB_32;
LB33= kcat_LB;
LB34= kcat_LB;
LB35= kcat_LB;
LB36= kcat_LB;

  UB26= kcat_UB_26;
UB27= kcat_UB;
UB28= kcat_UB;
UB29= kcat_UB;
UB30= kcat_UB;
UB31= kcat_UB;
  UB32= kcat_UB_32;
UB33= kcat_UB;
UB34= kcat_UB;
UB35= kcat_UB;
UB36= kcat_UB;

! Km_9_11, Km_1_68, Km_2_12,  Km_3_14, Km_4_15, Km_5_15: ;
Km_LB = 0.9;
Km_UB = 1.1;

LB37=Km_LB;
LB38=Km_LB;
LB39=Km_LB;
LB40=Km_LB;
LB41=Km_LB;
LB42=Km_LB;

UB37=Km_UB;
UB38=Km_UB;
UB39=Km_UB;
UB40=Km_UB;
```

```
UB41=Km_UB;
UB42=Km_UB;

! Km_6_20, Km_7_23, Km_8_21, Km_4_16,  Km_5_16: ;
LB43=Km_LB;
LB44=Km_LB;
LB45=Km_LB;
LB46=Km_LB;
LB47=Km_LB;

UB43=Km_UB;
UB44=Km_UB;
UB45=Km_UB;
UB46=Km_UB;
UB47=Km_UB;

! Km_17_24, Km_18_24, Km_19_12, Km_22_21, Km_22_23, Km_22_68, Km_25_20,Km_4_11,Km_7_13:;
LB48=Km_LB;
LB49=Km_LB;
LB50=Km_LB;
LB51=Km_LB;
LB52=Km_LB;
LB53=Km_LB;
LB54=Km_LB;
LB56=Km_LB;
LB60=Km_LB;

UB48=Km_UB;
UB49=Km_UB;
UB50=Km_UB;
UB51=Km_UB;
UB52=Km_UB;
UB53=Km_UB;
UB54=Km_UB;
UB56=Km_UB;
UB60=Km_UB;

! Ki_4_24, Ki_5_24: ;
Ki_LB = 0.9;
Ki_UB = 1.1;

LB57=Ki_LB;
LB59=Ki_LB;

UB57=Ki_UB;
UB59=Ki_UB;


! alpha_6, betha_6: ;
LB61=9.000000e-001;
LB63=9.000000e-001;

UB61=1.100000e+000;
UB63=1.100000e+000;

!Kd_9_11, Kd_17_24, Kd_22_68, Kd_22_23, Kd_22_21, Ka_6_14 :;
LB62=9.000000e-001;
LB64=9.000000e-001;
LB65=9.000000e-001;
LB66=9.000000e-001;
LB67=9.000000e-001;
LB69=9.000000e-001;

UB62=1.100000e+000;
UB64=1.100000e+000;
UB65=1.100000e+000;
UB66=1.100000e+000;
```

```
UB67=1.100000e+000;
UB69=1.100000e+000;

! hill_7_13:;
LB70=0.9;
UB70=1.1;

! K_10_0, K_4_0, K_5_0: ;
K_LB = 0.9;
K_UB = 1.1;

LB55=K_LB;
LB58=K_LB;
LB72=K_LB;

UB55=K_UB;
UB58=K_UB;
UB72=K_UB;  ! for Kcat;

!metabolites[0.1,5]:;

Met_LB = 0.1;       !0.1 or 0.8;
Met_UB = 2.5;       !1.2 or 2.5;

LB1=Met_LB;
LB2=Met_LB;
LB3=Met_LB;
LB4=Met_LB;
LB5=Met_LB;
LB6=Met_LB;
LB7=Met_LB;
LB8=Met_LB;
LB9=Met_LB;
LB10=Met_LB;


UB1=Met_UB;
UB2=Met_UB;
UB3=Met_UB;
UB4=Met_UB;
UB5=Met_UB;
UB6=Met_UB;
UB7=Met_UB;
UB8=Met_UB;
UB9= Met_UB;
UB10=Met_UB;

! Restrictionminimun flux U_9_11(opt) >= U_9_11(basal):;
 y11 + y26 - 0.0029399295211171745656475451369261*y37 +
0.99685126702718862556906742611318*y4 - 0.99685126702718851454676496359752*y56 -
 0.99142718740203761296214679532568*y62 + 0.99436711692315482569171081195236*y9 >=
 -16.589336311184282353580700053954+ 2.0000000000000001491862189340054*K;

!economy factor:;
p=2 ;!1 or 2;

0.27742946708463949843260188087774*y11 + 0.1630094043887147335423197492163*y12 +
0.025078369905956112852664576802508*y13 +
 0.086990595611285266457680250783699*y14 + 0.0047021943573667711598746081504702*y15 +
0.039184952978056426332288401253919*y20 +
 0.17084639498432601880877742946708*y21 + 0.15987460815047021943573667711599*y23 +
0.047805642633228840125391849529781*y24 +
 0.025078369905956112852664576802508*y68 <= 1.0*K + @log(p) -
9.3790079353335689568877234606994;

!IspA restrictions:;
y16-y11=0;
```

```
! y32-y26+@log(2.22)=0;! maintaining the same relationship than the original model i.e.
kcat_11/kcat_32=2.22 ;
! y26-y32=0;

AA=W11+W12+W14+W15+W16+W20+W21+W23+W24+W68;
AAA=W13;
AAAA=AA+AAA;

BB=W26+W27+W28+W30+W31+W32+W33+W34+W35+W36;
BBB= W29;
BBBB=BB+BBB;

CC=W37+W38+W39+W40+W41+W42+W43+W44+W45+W46+W47+W48+W49+W50+W51+W52+W53+W54+W56+W60;  ! 20
Km's: ;
DD= W17+W18+W19+W22+W25+W71;
EE1= W70;
EE2= W55+W58+W72;
FF= W62+W64+W65+W66+W67;
GG1= W57+W59;
GG2= W69;
HH= W61+W63;

CCCC=AAAA+BBBB;
CC_FF_GG1_GG2= CC+FF+GG1+GG2;


!AA);  AA<=10;
!AA1); AAA<=1;
!AA2); AAAA<=11;

!BB);  BB<=10;
!BB1); BBB<=1;
!BB2); BBBB<=11*0;

!CC);  CC<= 20;
!DD);  DD<= 6*1;
!EE1); EE1<= 1*0;
!EE2); EE2<= 3*1;
!FF);  FF<= 5*1;
!GG1); GG1<= 2*0;
!GG2); GG2<= 1*0;
!HH);  HH<= 2*0;

!JJ);  CCCC<= 22*1;

!KK);  CC_FF_GG1_GG2<= 28*1;

W11+M11=1;W12+M12=1;W13+M13=1;W14+M14=1;W15+M15=1;W16+M16=1;W17+M17=1;W18+M18=1;W19+M19=1;
W20+M20=1;W21+M21=1;W22+M22=1;W23+M23=1;W24+M24=1;W25+M25=1;W26+M26=1;W27+M27=1;W28+M28=1;
W29+M29=1;W30+M30=1;W31+M31=1;W32+M32=1;W33+M33=1;W34+M34=1;W35+M35=1;W36+M36=1;W37+M37=1;
W38+M38=1;W39+M39=1;W40+M40=1;W41+M41=1;W42+M42=1;W43+M43=1;W44+M44=1;W45+M45=1;W46+M46=1;
W47+M47=1;W48+M48=1;W49+M49=1;W50+M50=1;W51+M51=1;W52+M52=1;W53+M53=1;W54+M54=1;W55+M55=1;
W56+M56=1;W57+M57=1;W58+M58=1;W59+M59=1;W60+M60=1;W61+M61=1;W62+M62=1;W63+M63=1;W64+M64=1;
W65+M65=1;W66+M66=1;W67+M67=1;W68+M68=1;W69+M69=1;W70+M70=1;W71+M71=1;W72+M72=1;

@BIN(W11);@BIN(W12);@BIN(W13);@BIN(W14);@BIN(W15);@BIN(W16);@BIN(W17);@BIN(W18);@BIN(W19);@BIN(W20);
@BIN(W21);@BIN(W22);@BIN(W23);@BIN(W24);@BIN(W25);@BIN(W26);@BIN(W27);@BIN(W28);@BIN(W29);@BIN(W30);
@BIN(W31);@BIN(W32);@BIN(W33);@BIN(W34);@BIN(W35);@BIN(W36);@BIN(W37);@BIN(W38);@BIN(W39);@BIN(W40);
@BIN(W41);@BIN(W42);@BIN(W43);@BIN(W44);@BIN(W45);@BIN(W46);@BIN(W47);@BIN(W48);@BIN(W49);@BIN(W50);
@BIN(W51);@BIN(W52);@BIN(W53);@BIN(W54);@BIN(W55);@BIN(W56);@BIN(W57);@BIN(W58);@BIN(W59);@BIN(W60);
@BIN(W61);@BIN(W62);@BIN(W63);@BIN(W64);@BIN(W65);@BIN(W66);@BIN(W67);@BIN(W68);@BIN(W69);@BIN(W70);
@BIN(W71);@BIN(W72);

END
```

### References


[1] I. LINDO Systems, LINGO 11.0.1.6, 1415 North Dayton Street. Chicago, IL 60622, 2008.

[2] L. Fuhrer, C.P. Kubicek, and M. Rohr, Pyridine nucleotide levels and ratios in *Aspergillus niger*. Can J Microbiol 26 (1980) 405-8.

[3] J.L. Galazzo, and J.E. Bailey, Fermentation pathways kinetics and metabolic flux control in suspended and inmobilized Saccharomyces cerevisae. Enzyme Microb. Technol. 12 (1990) 162-172.

[4] L. Guarente, G. Lauer, T.M. Roberts, and M. Ptashne, Improved methods for maximizing expression of a cloned gene: a bacterium that synthesizes rabbit beta-globin. Cell 20 (1980) 543-53.

[5] P. Niederberger, R. Prasad, G. Miozzari, and H. Kacser, A strategy for increasing an in vivo flux by genetic manipulations. The tryptophan system of yeast. Biochem J 287 ( Pt 2) (1992) 473-9.

[6] K. Zhou, R. Zou, G. Stephanopoulos, and H.P. Too, Metabolite profiling identified methylerythritol cyclodiphosphate efflux as a limiting step in microbial isoprenoid production. PLoS One 7 (2012) e47513.

[7] S.P. Lim, and H. Nikaido, Kinetic parameters of efflux of penicillins by the multidrug efflux transporter AcrAB-TolC of *Escherichia coli*. Antimicrob Agents Chemother 54 (2010) 1800-6.

[8] H. Nikaido, and S. Normark, Sensitivity of *Escherichia coli* to various beta-lactams is determined by the interplay of outer membrane permeability and degradation by periplasmic beta-lactamases: a quantitative predictive treatment. Mol Microbiol 1 (1987) 29-36.

[9] A. Sorribas, and M.A. Savageau, Strategies for representing metabolic pathways within biochemical systems theory: reversible pathways. Math Biosci 94 (1989) 239-69.
